# OptiFoot: a method for recording protein footprints on DNA for microscopy and sequence analysis

**DOI:** 10.64898/2026.05.12.724470

**Authors:** Rūta Gerasimaitė, Jonas Bucevičius, Dovilė Bubnytė, Tanja Koenen, Gražvydas Lukinavičius

## Abstract

A comprehensive understanding of protein–nucleic acid interactions in the crowded nuclear milieu is essential for elucidating genome function. State-of-the art methods provide the finest details of genome organization, but lack integration of imaging and sequencing modalities. We introduce OptiFoot, a fluorescence-based platform for recording protein–nucleic acid interactions in cells. OptiFoot employs an engineered sequence-unspecific N6-adenine methyltransferase that attaches diverse functional groups to DNA and RNA. When targeted to protein of interest by genetic fusion or antibody, it generates covalent high contrast fluorescent footprints that can be visualized by super-resolution microscopy and analyzed by optical mapping of single native DNA molecules, allowing complementary spatial and genomic analyses. To illustrate OptiFoot versatility, we imaged lamina-associated domains, DNA replication sites, CTCF binding sites and histone modifications in human cells and produced corresponding genome profiles. We use OptiFoot to demonstrate that majority of nucleoporin NUP153-DNA interactions occur in nucleoplasm, outside nuclear pore.

---

Protein–nucleic acids interactions are essential for controlling transcription, replicating and safeguarding genome. The extremely crowded, intricately organized and dynamic nucleus environment poses many challenges for research and as a result plethora of diverse methods have been developed (reviewed in ^1^). Based on the underlying readout technique, these methods fall into either of the two broad categories: DNA sequencing or microscopy imaging. Sequencing-based approaches, can map protein binding with several base pair resolution ^2, 3^, while the super-resolution fluorescent microscopy can trace chromatin with down to nanometer precision ^4-6^. However, sequencing-based methods lack a simple visual readout and are poorly suited for time-resolved studies of dynamic processes, while imaging approaches struggle to provide genome-wide coverage. A potential solution for filling-in this gap is harnessing enzymes that can covalently modify native DNA in situ thereby creating durable molecular records that can be decoded by complementary readouts.

Prokaryotic AdoMet-dependent *N*6-adenine DNA methyltransferases (MTases) have been used in genome research for over two decades. Because N6-methyl adenine (6mA) is present at extremely low levels in genomes of higher eukaryotes ^7^, it can serve as a minimally invasive mark for chromatin structure. Several sequencing-based approaches to detect protein-DNA interactions rely on 6mA footprints. In DNA adenine methylase identification (DamID), protein of interest (POI) is expressed as a fusion with E.coli DNA adenine methyltransferase (EcoDam) which generates 6mA footprints at nearby G<u>A</u>TC target sites ^8^. EpiDamID extends this approach to chromatin states by fusing EcoDam to chromatin readers or single-chain antibodies targeting histone modifications ^9^. These techniques require few cells, avoid crosslinking and antibodies, and records POI occupancy in vivo ^10, 11^. However, they require genetic modification, have limited temporal resolution and cannot probe GATC-poor regions. Replacing EcoDam with the sequence-unspecific MTase EcoGII enabled unbiased genome coverage in MadID ^12, 13^. Temporal control was improved in pA-DamId, where protein A-EcoDam is targeted to endogenous POIs via antibody in permeabilized cells ^14^. Related strategies have been implemented with unspecific N6-MTases: EcoGII in BIND&MODIFY ^15^ and Hia5 in nanoHiMe-seq and DiMeLo-seq ^16-18^. The power of N6-MTases is maximized when combined with long-read sequencing, such as Oxford Nanopore sequencing and PacBio Single Molecule Real-Time sequencing (SMRT) that directly detect MTase-deposited 6mA footprints and endogenous 5mC on single DNA molecules ^19, 20^. This avoids enrichment bias and preserves long-range connectivity of chromatin fibers ^15, 17, 18, 21^.

These approaches generate linear interaction profiles and do not provide spatial context. In principle, 6mA footprints can be detected by immunostaining with anti-6mA antibody ^13, 22^ or 6mA-binding protein domain 6mA-Tracer ^14, 23^. However, these interactions are non-covalent, often suffer from poor contrast and 6mA remains a poor reporter group for microscopy. Methyltransferase-assisted transfer of activated groups (mTAG) might provide an alternative by enabling covalent installation of fluorophores, affinity tags, and reactive handles directly onto native DNA at short (2–8 bp) specific MTase target sequences (**Fig. 1a**) ^24, 25^. The reaction requires an artificial AdoMet analog with a functionalized transferable chain and an activating propargyl group next to sulfonium center. Often engineering of MTase active site to accommodate these larger transferable groups is needed ^26, 27^. Despite offered versatility, mTAG applications in microscopy so far have been limited to imaging stretched DNA molecules for optical genome mapping (OGM) ^28^.

**Fig. 1.**
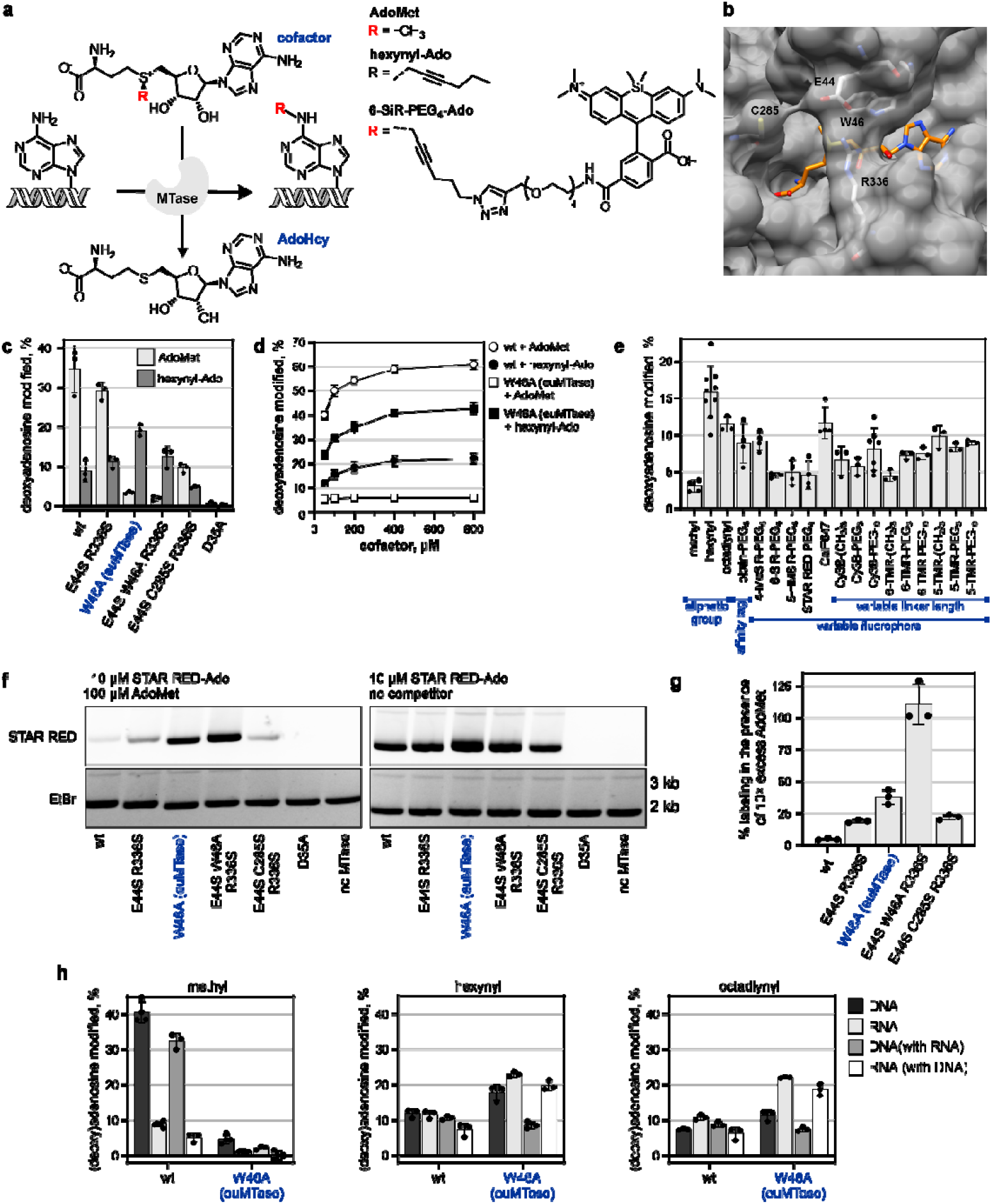
Engineering sequence-unspecific adenine N6-MTase EcoGII to obtain euMTase. **a**, Reaction catalyzed by DNA adenine *N*6-MTases. Transferable group highlighted in red. Structures of the natural cofactor AdoMet, a model activated analog hexynyl-Ado and a cofactor with a fluorescent transferable chain, 6-SiR-PEG_4_-Ado. **b**, Cofactor binding site of M.EcoGII modelled using experimental M.MboIIA-AdoMet structure (1g60) as a template. The protein is rendered as semi-transparent surface, mutagenized amino acids are shown as sticks. AdoMet position (orange) is inferred from template. **c**, Activity of wt M.EcoGII and its mutants with 50 µM AdoMet or hexynyl-Ado. M.EcoGII D35A – catalytic mutant used as a control. **d**, increasing AdoMet concentration does not rescue methylation activity of M.EcoGII W46A (euMTase). Mean ± s.d., n = 3. **e**, Activity of W46A (euMTase) with cofactors bearing transferrable chains of various structure and function. Chemical structures of all AdoMet analogs are in **Supplementary Scheme 1. f-g**, euMTase efficiently labels DNA in the presence of 10× excess of AdoMet. **f**, pUC19 DNA was incubated with M.EcoGII variants and 10 µM STAR RED-Ado in the presence or absence of 100 µM AdoMet for 1 h at 37°C and fractionated on agarose gel. **g**, quantification of fluorescence signal in the agarose gel. Mean ± s.d., n = 3. **h**, Modification of RNA and RNA-DNA mixtures by wt M.EcoGII and euMTase. Percentage of adenine modified was determined by HPLC analysis of nucleoside mixtures (Supplementary **Figs. S5-6**). Mean ± s.d., n = 3. 6mdA – modified deoxyriboadenosine in λ DNA; 6mA – modified riboadenosine in total E.coli RNA; 6mdA (with RNA) and 6mA (with DNA) – modified deoxyriboadenosine and riboadenosine in DNA-RNA mixture.

Here we present OptiFoot (<u>opti</u>cal <u>foot</u>printing of protein-nucleic acids interactions), an integrated platform that combines fluorescence microscopy with optical genome mapping to interrogate protein– nucleic acid interactions in situ inside nucleus by microscopy and downstream sequence analysis. It relies on the engineered sequence-unspecific N6-adenine methyltransferase (euMTase) that preferentially utilizes synthetic AdoMet analogs bearing extended, functionalized reporter groups. The absence of sequence specificity enables dense, high-contrast labeling of nucleic acids, which is critical for resolving defined nuclear features within the crowded chromatin environment. When targeted to a protein of interest by antibody guidance or direct genetic fusion, euMTase covalently marks proximal adenines, thereby recording protein footprints on nucleic acids for visualization by conventional and STED microscopy^29^. Labeled DNA can be isolated and subjected to optical genome mapping to position footprints along genome sequence. We demonstrate this by creating genome-wide interaction profiles of LAMIN B1, CTCF and histone modifications in U-2 OS cells and showing that they agree well with the published data. Imaging nucleoporin NUP153 footprints revealed two functionally distinct NUP153– chromatin subpopulations with different DNA accessibility: one in the nucleoplasm and another associated with the nuclear pore complex.

## Results

### Synthesis of AdoMet analogs with functionalized transferable groups

We synthesized activated AdoMet analogs by regioselective S-alkylation of S-adenosylhomocysteine (AdoHcy) with octa-2,7-diyn-1-yl 4-nitrobenzenesulfonate or 6-azidohex-2-yn-1-yl 4-nitrobenzenesulfonate as alkylating agents (**Supplementary Scheme 1**)^27^. These reactions produced 1:1 mixture of diastereomers at the sulfonium center, from which the active, R-epimer, was purified by preparative reversed-phase high-performance liquid chromatography (RP-HPLC). In the next step, functional moieties – fluorophores or biotin – were attached via copper-catalyzed azide-alkyne cycloaddition (“click”) reactions. Final RP-HPLC purification yielded active cofactors with ≥75%purity. Information about synthesis and characterization of the new cofactors can be found in **Supplementary Methods**.

### Engineering M.EcoGII for large group transfer

mTAG reaction efficiency varies a lot between MTases, and only several wt or engineered enzymes have shown sufficient activity for practical applications ^30^. All published studies employ sequence-specific DNA MTases, leading to labeling patterns that are inherently biased by the uneven genomic distribution and abundance of their 2-to 6-bp recognition sequences. Therefore, we focused on sequence-unspecific DNA and RNA adenine N6-MTase M.EcoGII. High sequence and structure homology of DNA MTases’ family allows building high confidence homology models that provide a good starting point for enzyme engineering.

We used SWISS MODEL ^31^ to build a homology model of M.EcoGII using a structure of M.MboIIA in complex with AdoMet (1g60) as a template ^32^. The model suggested that side chains E44, W46, R336 and S285 might obstruct binding of longer transferable groups (**Fig. 1b**). These positions are conserved among the closest M.EcoGII homologs (**Supplementary Fig. 1**), but are variable across the broader protein family. Notably, they map within or proximal to structurally and functionally conserved regions (**Supplementary Fig. 2**), implying potential participation in the cofactor binding and catalysis. To enlarge the cofactor binding pocket, we mutated these positions into smaller amino acids, either individually or in combination, and compared ability of mutant MTases to transfer methyl and hexynyl groups to λ DNA (**Fig. 1c**). After reaction, DNA was hydrolyzed and the resulting nucleoside mixture was analyzed by LC– MS to identify modified nucleoside and quantify MTase activity (**Supplementary Fig. 3-5**). Similarly to several sequence-specific N6-MTases, wt M.EcoGII showed significant hexynyl transfer activity, although this was approximately 5-fold lower than its methylation activity.The most interesting mutant, W46A, showed ∼2-fold higher activity with the hexynyl-Ado but ∼15-fold reduced activity with AdoMet compared to wt, indicating specificity shift towards AdoMet analogs. Increasing AdoMet concentration did not restore its methylation activity (**Fig. 1d**), indicating non-productive cofactor binding rather than reduced affinity.

Going from hexynyl-Ado to octadiynyl-Ado, W46A activity modestly decreased, but larger groups were well tolerated thereafter (**Fig. 1e**), allowing flexible functional group choice depending on the downstream experimental requirements. Fluorophores spanning positive (4-MeSiR, CalFluor647), zwitterionic (TMR, Cy3B, SiR and HMSiR), and negative (STAR RED) net charges showed broadly similar transfer efficiencies, and PEG linkers had little effect. In further work, we mostly used AdoMet analogs with PEG_4_ linker, as it ensured a better solubility of cofactors with hydrophobic dyes.

In direct competition assay (**Fig. 1f,g**), a 10-fold excess of AdoMet reduced labeling by wt enzyme with STAR RED dye to 4% of “no competitor” level, whereas W46A retained 38% activity. Although the triple mutant E44S W46A R336S was fully resistant to AdoMet competition, W46A showed higher absolute activity and was selected for further studies. Hereafter, M.EcoGII W46A is termed euMTase (<u>e</u>ngineered <u>u</u>nspecific methyltransferase).

Because M.EcoGII also methylates RNA ^12, 33^, we assessed euMTase methylation and alkylation activity on RNA. RP-HPLC readily resolves ribo- and deoxyribonucleosides, which allowed simultaneous quantification of RNA and DNA modification in DNA-RNA mixtures – mimicking conditions in chromatin (**Supplementary Fig. 5**). As reported previously, wt M.EcoGII methylated DNA more efficiently than RNA, but this preference disappeared with AdoMet analogs (**Fig. 1h**). As with DNA, euMTase displayed reduced methylation but enhanced alkylation of RNA. Interestingly, in DNA–RNA mixtures, euMTase preferentially modified RNA (**Fig. 1h**). This suggests that in chromatin context both, DNA and RNA, may serve as labeling substrates.

### OptiFoot targeting by genetic fusion

Using wild-type MTases for DNA labeling in cells is severely limited by the lack of temporal control, due to ever-present AdoMet. Exogenously supplied fluorescent cofactor might be a solution, but requires orthogonal MTase, that would not methylate its targets beforehand.

To investigate orthogonality, we expressed eGFP-tagged euMTase, M.EcoGII wt and catalytic mutant D35A targeted to the nucleus in U-2 OS cells (**Supplementary Fig. 6**). Challenging gDNA isolated from the induced cultures with 6mA-dependent R. DpnI and 6mA-sensitive R.MboI restriction enzymes detected 6mA only in cells expressing wt M.EcoGII, but not euMTase or D35A (**Supplementary Fig. 6d**). Upon permeabilization and incubation with fluorescent cofactor 6-SiR-PEG_4_-Ado, euMTase showed the strongest staining, while staining by wt M.EcoGII was close to background (**Supplementary Fig. 6e-g**). These results indicate that euMTase displays sufficient orthogonality toward AdoMet analogs, enabling efficient labeling in cellular context.

As a proof-of-principle of fluorescent footprinting, we chose to label a prominent nucleus compartment - lamina-associated domains (LADs) (**Fig. 2a**). To this end, we expressed euMTase-lamin B1 fusion in U-2 OS cells using stable episome expression system. To avoid lamin B1 overexpression phenotype ^34^ and limit 6mA accumulation ^35^, expression was kept low by using a doxycycline-inducible promoter and an N-terminal destabilizing domain (DD) ^36^ (**Supplementary Fig. 7**). The immunostaining of induced cells with anti-DD antibody was highly mosaic (**Fig. 2b**): a small subpopulation of cells stained brightly, while majority showed very weak or no staining. Importantly, the bright cells retained normal nucleus morphology. After permeabilization and incubation with 6-SiR-PEG_4_-Ado, a characteristic nuclear rim staining was observed in subpopulation of cells that expressed high levels of transgene (**Fig. 2b**). Super-resolution STED imaging revealed finer structural details of LADs (**Fig. 2c,d**).

**Fig. 2.**
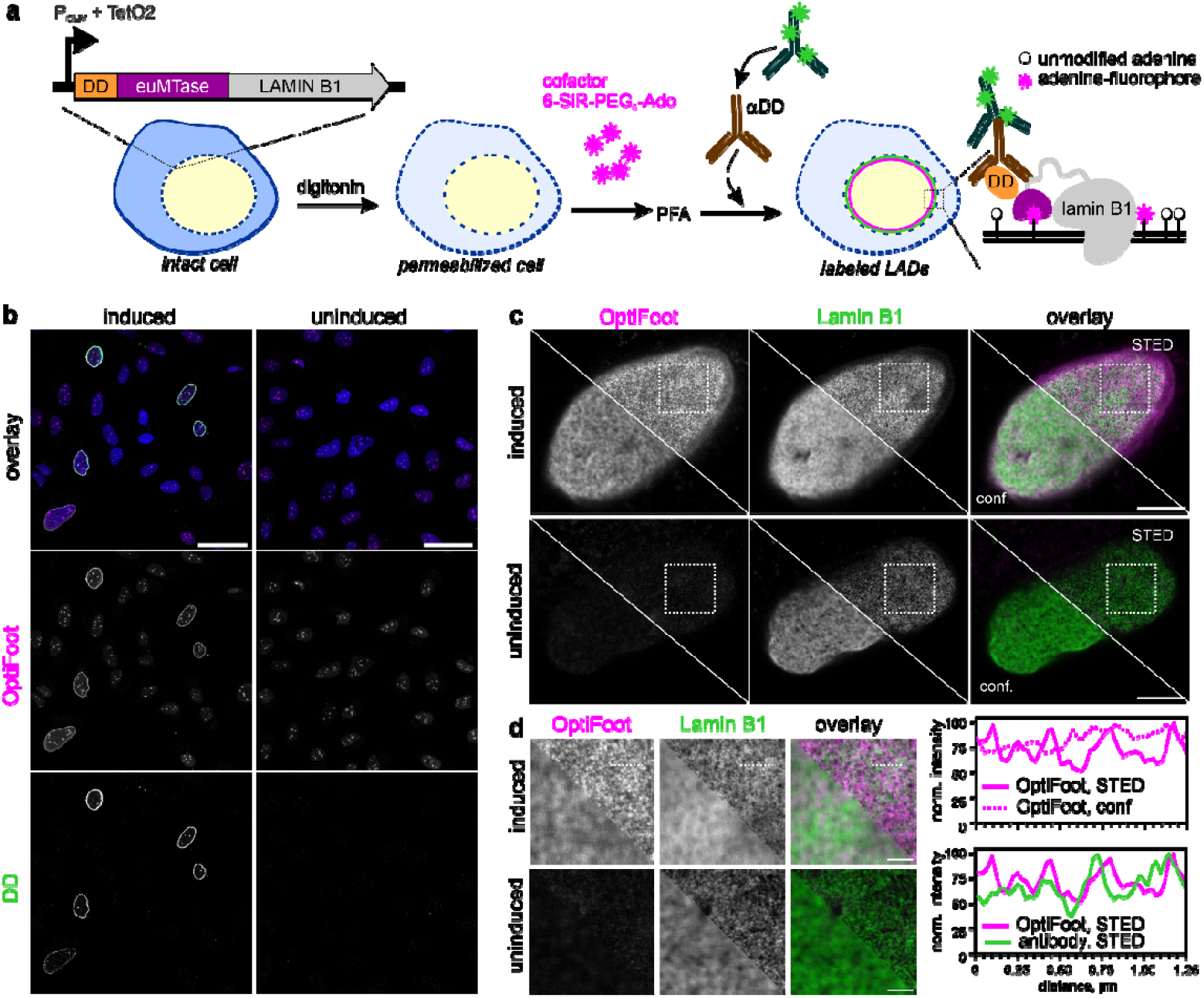
euMTase-hlamin B1 fusion is targeted to nuclear rim and labels lamina associated domains. **a**, schematic representation of expression construct and labeling workflow. DD – degradation domain, stabilized by shield-1 ligand. LAD – lamina-associated domain. **b**, DNA labeling correlates with DD-euMTase-hlamin B1 expression. Confocal slice through cell mid-section is shown. Transgene expression is detected with anti-DD antibody. Scale bar 50 µm. **c**, STED images of LADs in situ. For reference, lamin B1 is visualized with anti-lamin B1 and STAR 580-labelled secondary antibody. STED and confocal slice through bottom of the nucleus are shown for OptiFoot and antibody staining. Note that no LAD staining is visible in the non-induced cell. Scale bar 5 µm. **d**, comparison of confocal and STED image resolution in zoom-ins from c. Line profiles along the dotted line. Scale bar – 1 µm.

### OptiFoot targeting by antibody

To demonstrate the versatility of OptiFoot applications, we employed antibody-mediated targeting of euMTase to the protein of interest (POI). This modular strategy can be readily adapted for imaging any target for which a suitable antibody is available. We fused euMTase with protein A/protein G (pA/G) on N- or C-terminus generating pA/G-L_6_-euMTase and euMTase-L_14_-pA/G, respectively. Both constructs were similarly active as untagged euMTase on λ DNA, retaining ∼5-fold higher activity with hexynyl-Ado than with AdoMet (**Supplementary Fig. 8**).

Again, as a proof-of-principle we labeled LADs (**Fig. 3a**). We permeabilized cells with digitonin, then incubated them with anti-lamin B1 antibody followed by euMTase-L_14_-pA/G and finally labelled, followed by reaction with 3 µM 6-SiR-PEG_4_-Ado for 1h at 37°C. After washing away unreacted cofactor, the samples were fixed with PFA and stained with a secondary antibody. Following this procedure, characteristic nuclear periphery staining was readily detectable (**Fig. 3b**). Consistent with experiments in solution, the staining was reduced by the presence of 10-fold excess and eliminated by 100-fold excess of AdoMet. No LAD staining was observed if antibody or euMTase-L_14_-pA/G was omitted or anti-lamin B1 antibody was substituted with isotype control (rabbit IgG antibody). In the alternative scheme, LAD labeling was also achieved by first introducing biotin in OptiFoot reaction and then staining with fluorescently-labeled streptavidin (**Supplementary Fig. 9**).

**Fig. 3.**
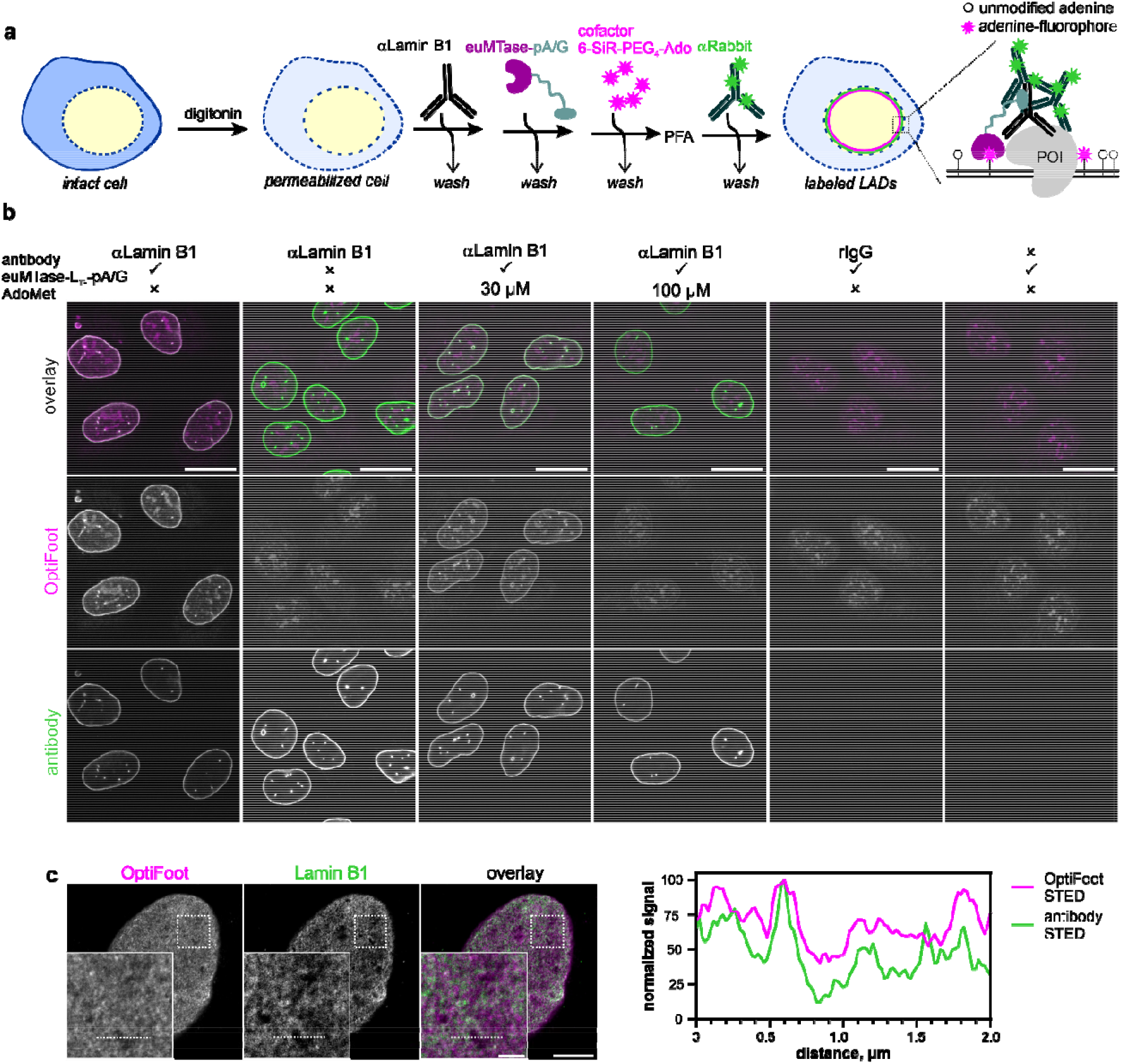
Antibody-targeted labeling of LADs in U-2 OS cells. **a**, antibody-targeted OptiFoot workflow consists of 4 steps: 1) cell permeabilization, 2) binding of antibody, 3) binding of euMTase-L_14_-pA/G and 4) incubation with fluorescent cofactor. After that, the cells can be fixed and stained with antibodies. PFA – paraformaldehyde fixation. **b**, confocal images of labeled LADs. Anti-lamin B1 (mAb) was used for targeting, labeling was performed with 3 µM 6-SiR-PEG_4_-Ado. In the control experiments, the indicated components were omitted or indicated concentration of AdoMet was added during labeling step. rIgG – unspecific rabbit antibody. Anti-lamin B1 antibody was visualized with secondary antibody conjugated with AlexaFluor488. Confocal slices through cell mid-section are shown, scale bar – 20 µm. **c**, STED images of LADs labeled by OptiFoot and the targeting anti-lamin B1 (pAb) stained with anti-rabbit IgG nanobody – Halo fusion labeled with Abberior STAR 580. Scale bars 5 µm, inset – 1 µm. Profiles along the dotted line are shown.

To determine if DNA or RNA is a source of specific signal, we exploited the ability of euMTase to modify DNA in methanol fixed cells. Fixation allowed pretreatment of samples with nucleases without collapse of nucleus structure before carrying on with antibody-targeted OptiFoot labeling (**Supplementary Fig 10**). DNase I treatment abolished both Hoechst33342 and LAD staining in lamin B1–targeted samples, whereas RNase A had minimal effect. Even in RNA-rich structure such as nucleolus, RNase A caused only a modest signal reduction, while DNase I reduced signal to background. Together, these results indicate that DNA is the primary source of the observed fluorescence.

Similarly, we directed OptiFoot labeling to diverse nuclear compartments and chromatin features: nucleolus (antibody against nucleolin), DNA replication sites (PCNA), insulator elements (CTCF), transcriptionally active chromatin (H3K4ac), enhancers (H3K27ac) and heterochromatin (H3K9me2) in U-2 OS and Hela cells and in human fibroblasts (**Fig. 4a, Supplementary Fig. 11**).

**Fig. 4.**
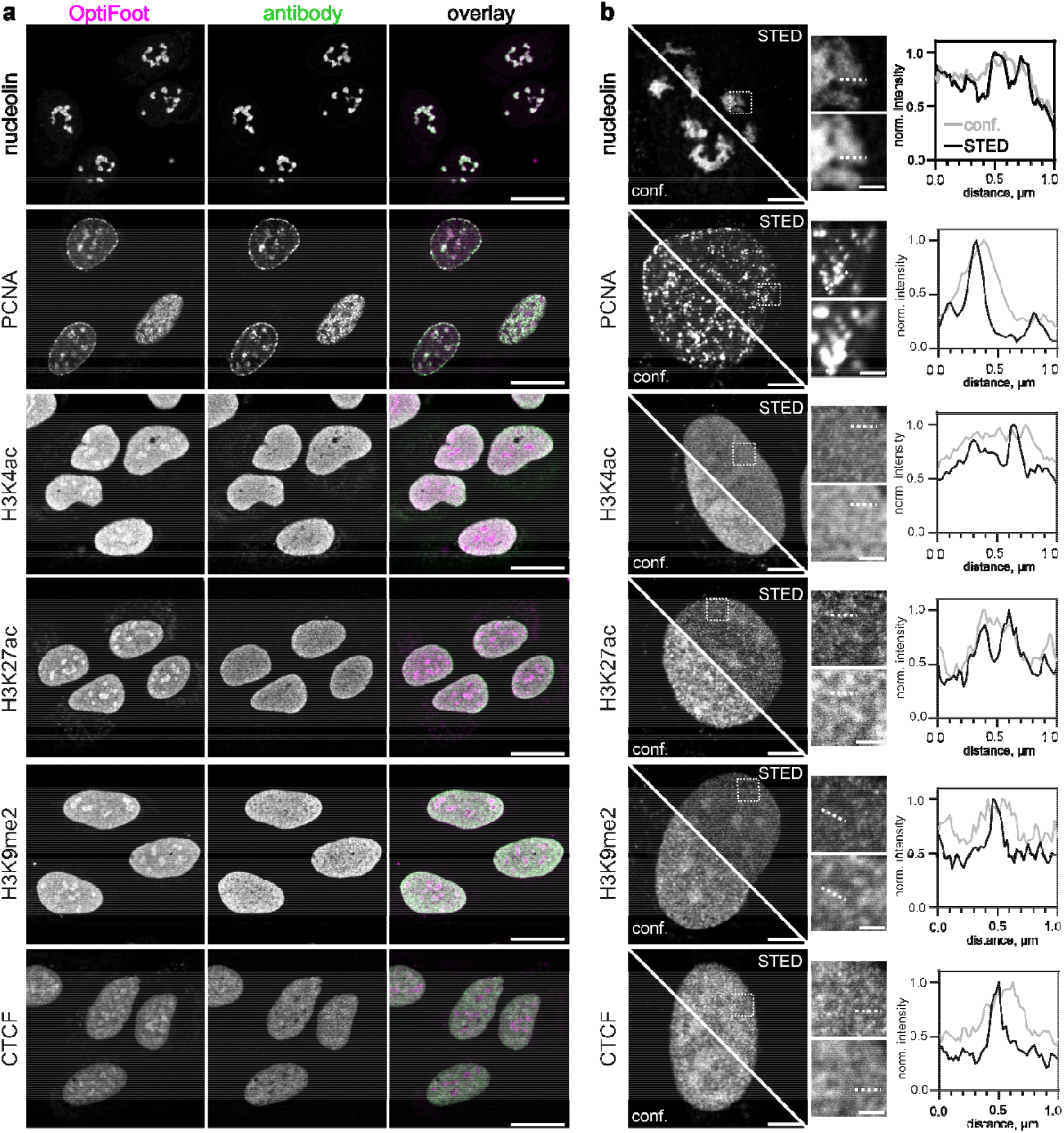
OptiFoot labeling of nuclear compartments, DNA replication sites, histone modifications and CTCF binding sites in U-2 OS cells. **a**, co-localization of antibody-directed OptiFoot staining with secondary antibody staining. A single confocal z-slice through nucleus midsection; scale bar – 25 µm. **b**, Left, comparison of confocal and STED images of OptiFoot-stained nuclei. A Gaussian blur with a standard deviation (σ) of 0.5 was applied using Fiji (ImageJ). Scale bars: 5 µm – whole nucleus, 1 µm – inset. Right, normalized intensity profiles along the dotted lines shown in the insets. Confocal signal – gray, STED – black.

Many levels of chromatin organization are beyond light diffraction limit making super-resolution fluorescent microscopy an essential tool for chromatin studies ^1^. OptiFoot labeling produced sufficient contrast to obtain super-resolution STED microscopy images of all the probed nucleus compartments (LADs and nucleoli), chromatin at CTCF binding sites and different chromatin states (histone modifications), providing access to finer details of their organization (**Fig. 3c, 4b**).

In addition to targeted labeling, we observed low unspecific staining, seemingly enriched in dense nuclear regions such as nucleoli (**Figs. 2b, 3b, Supplementary Fig. 11**). Activity of endogenous MTases with fluorescent cofactors is unlikely to be the main source, as excess AdoMet did not reduce this signal and staining persisted after methanol fixation, when enzymatic activity is largely lost (**Fig. 3b; Supplementary Fig. 10**). Thus, off-target background is most consistent with nonspecific interactions of the cofactor with membranes or hydrophobic protein surfaces. Cofactor chemistry strongly affected this signal (**Supplementary Fig. 12**). Positively charged 4-MeSiR-PEG4-Ado showed prominent cytoplasmic staining, consistent with mitochondrial accumulation and membrane staining with lipophilic cationi dyes ^37, 38^, whereas negatively charged STAR RED-PEG4-Ado reduced this effect. Nucleolar background was lower with CalFluor647-Ado and 6-SiR-PEG4-Ado, likely owing to shielding effects of PEG groups or the fluorogenic properties of SiR ^39, 40^. We therefore used 6-SiR-PEG -Ado for most microscopy experiments because it provided the lowest background with strong targeted signal. Notably, off-target staining is present in the 2-step labeling with biotin-PEG_4_-Ado, but is nearly absent when cofactor is omitted (**Supplementary Fig. 9**). This argues that adenosine part of the cofactor rather than fluorophore is the major contributing factor.

In the previous examples, OptiFoot staining closely follows respective immunostaining. It is not always the case, because euMTase activity is directed related to local DNA accessibility. This was particularly evident for the dynamic nucleoporin NUP153, which is known to interact with chromatin at nuclear pore complex (NPC) and in the nuclear interior ^41, 42^. In line with this, immunostaining was enriched at the nuclear envelope with weaker nucleoplasmic signal (**Fig. 5a**). In contrast, OptiFoot labeling was broadly distributed throughout the nucleoplasm, with little enrichment at the envelope. This suggests that off-pore NUP153 engages more accessible chromatin and therefore generates stronger labeling despite lower abundance. At the nuclear envelope plane, discrete OptiFoot foci colocalized with antibody signal, directly confirming NUP153–DNA interactions at NPCs (**Fig. 5b**). Together, these results highlight the ability of OptiFoot to distinguish functional protein states based on chromatin engagement.

**Fig. 5.**
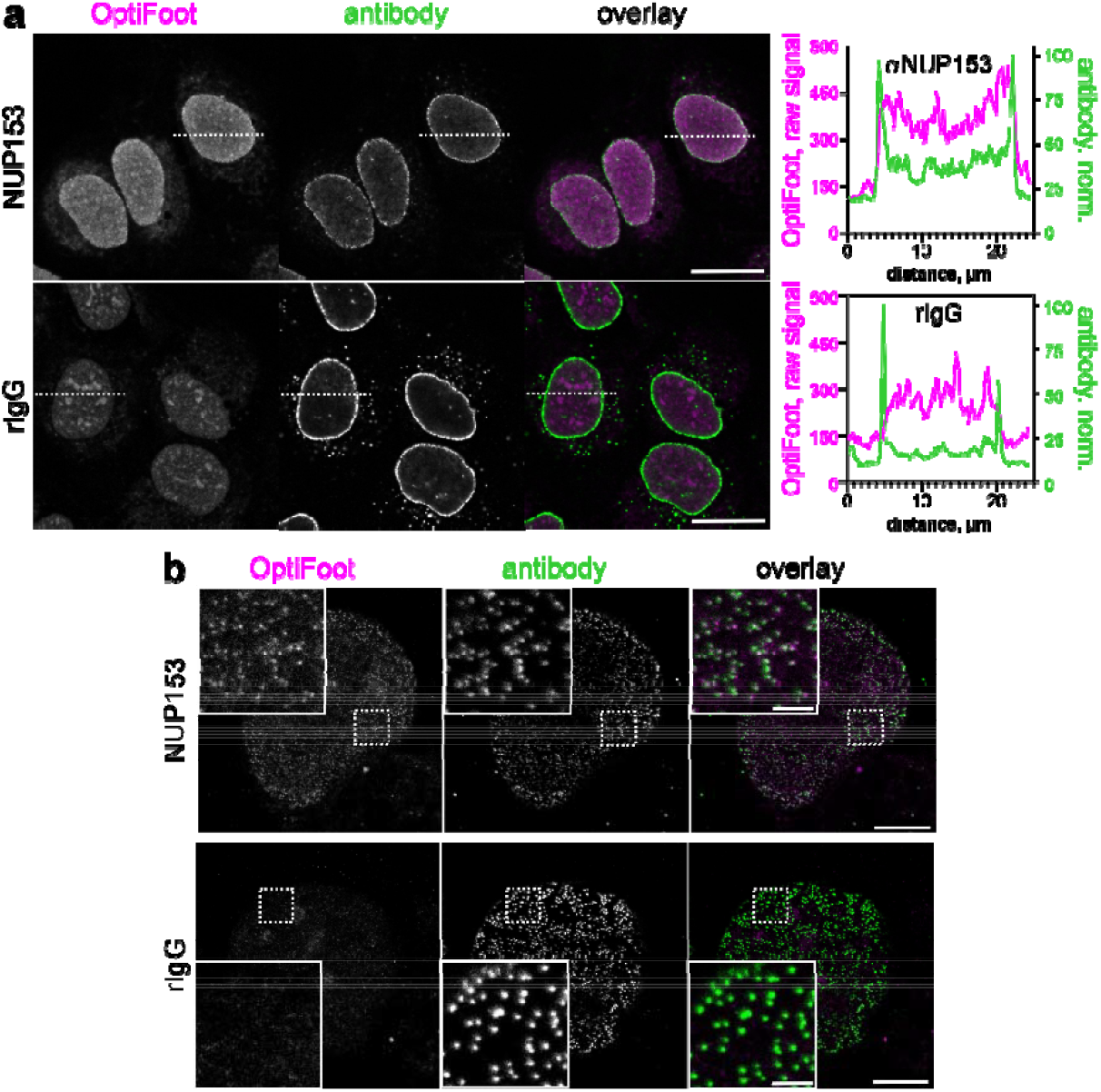
Majority of NUP153-DNA contacts occur in the nucleus interior. **a**, NUP153-DNA interaction in the nucleoplasm. Left, OptiFoot staining targeted by anti-NUP153 antibody or control rIgG. Confocal slice through nucleus midsection is shown. Scale bar 20 µm. Right –profiles along the dotted line. OptiFoot staining is shown as a raw signal to facilitate the comparison between the samples, while antibody signal is normalized. **b**, STED images of NUP153 footprints at the nucleopore. Scale bar – 5 µm, inset – 1 µm. rIgG sample was stained with anti-NUP153 and secondary antibody after OptiFoot labeling and fixation.

### Protein-genome interaction profiling with optical genome mapping

Covalent nature of OptiFoot labels allows isolation and further analysis of modified DNA in vitro. We combined OptiFoot with optical genome mapping (OGM) to generate genome-wide protein occupancy profiles. As a benchmark application, we profiled LADs. Following OptiFoot labeling with red dye (SiR or STAR RED), we isolated ultra-high molecular weight DNA, labeled it at reference sequences with a green commercial dye and analyzed it by OGM, which images stretched fluorescent DNA molecules and aligns them to genome according to reference labeling patterns (**Fig. 6a, Supplementary Fig.13 a**)^43^.

**Fig. 6.**
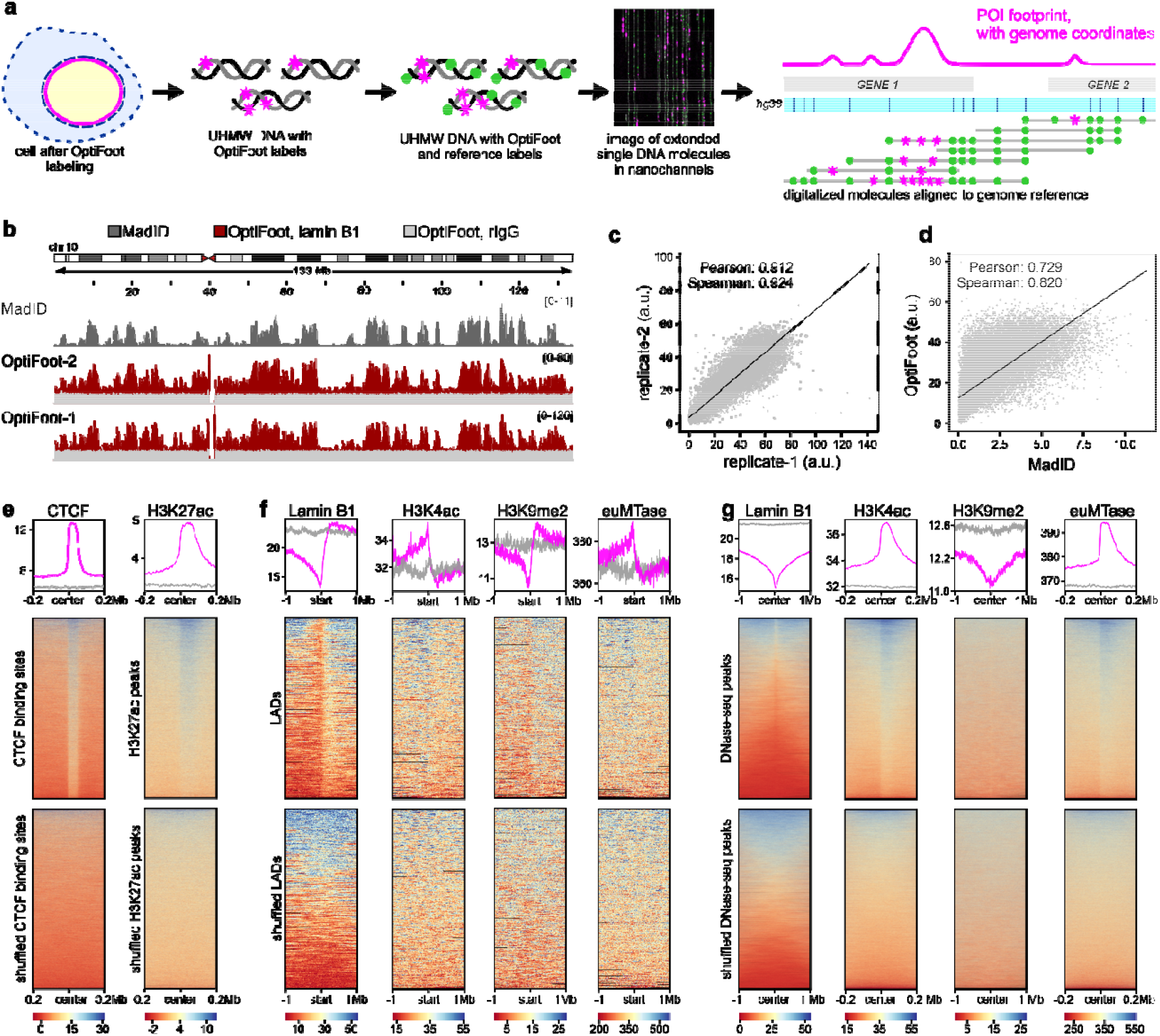
OptiFoot reports interaction profiles of different POIs over the genome. **a**, a workflow of obtaining genome-wide OptiFoot footprints by OGM. **b**, comparison of LAD profiles over chromosome 10 in U-2 OS cells obtained by OptiFoot with those in Hela CCL2 generated with MadID. OptiFoot profiles binned to 100 kb windows to match resolution of MadID dataset ^13^. OptiFoot profile produced with polyclonal lamin B1 antibody is shown in red, background signal is shown in light gray, madID profile is in dark gray. Two OptiFoot replicates are shown with corresponding control traces. **c**, correlation between OptiFoot and MadID signals over the whole genome. **d**, correlation between OptiFoot replicates. Correlations were computed with background-subtracted OptiFoot profiles. **e-g**, profiles and heatmaps of enrichment of OptiFoot signal targeted to various POI across genomic features. Profiles show average OptiFoot signal intensity centered at feature coordinates, with heatmaps visualizing signal across individual features. Enrichment in specific features is in magenta, control in randomized genomic regions is shown in green. List of reference genome features is listed in **Supplementary Table 3. e**, OptiFoot signal obtained with anti-CTCF and anti-H3K27ac antibodies **f**, enrichment of various factors at LAD boundaries. **g**, enrichment in DNase-seq peaks. Enrichment was computed from background-corrected profiles binned to 1 kb.

Two labeling conditions employing different digitonin, antibody, and cofactor concentrations (**Supplementary Table 1**) produced similar LAD profiles despite different label densities (1.17 versus 2.06 labels per 100 kb), with high reproducibility between replicates (**Fig. 6b,c, Supplementary Fig.13 b**). IgG controls showed minimal labeling and no structured signal. OptiFoot lamin B1 profiles in U-2 OS cells correlated well with published MadID data from HeLa cells (Pearson r = 0.7; Spearman r = 0.8)^13^, consistent with conservation of prominent constitutive LADs at 100 kb resolution (**Fig. 6d**). This level of correlation is comparable to that observed in previous studies comparing different methods of POI profiling. ^9, 15, 18^.

We extended OGM profiling to CTCF binding, histone modifications (H3K4ac, H3K27ac, H3K9me2), and DNA accessibility using soluble euMTase (**Fig. 6e,f, Supplementary Table 1**). Because OGM is diffraction-limited to ∼500 bp/pixel, signals were analyzed in 1 kb bins after subtraction of IgG controls, where residual labels appeared random. Resulting profiles showed expected enrichments: CTCF at CTCF sites, H3K27ac at active regulatory regions, and lamin B1 with H3K9me2 at LADs, while active marks and accessibility were enriched at promoters and DNase-accessible regions (**Fig. 6g; Supplementary Fig. 13c**). Together, these data show that OptiFoot faithfully captures genome-wide chromatin interaction landscapes.

## Discussion

Here we introduce OptiFoot, a method for recording POI footprints on genomic DNA in situ by covalent attachment of a range of biorthogonal functional groups, including fluorophores. The latter allows direct imaging of genomic DNA regions in contact with POI in the nucleus with conventional and super-resolution fluorescence microscopy. OptiFoot labeling can be targeted by genetic fusion or via an antibody and thus offers flexible experiment design. The absence of sequence specificity allows for uniform genome-wide coverage and high local labeling density, which supports high contrast. Direct covalent attachment of fluorophores to nucleobases minimizes linkage error, a critical factor for exploiting the spatial resolution of advanced super-resolution techniques such as STED or MINFLUX^44^.

Unlike immunostaining, OptiFoot labeling is directly governed by local DNA accessibility, paving the way to discriminate between different functional states of POI. Thereby, NUP153-targeted OptiFoot directly demonstrated better DNA accessibility in off-pore NUP153-chromatin complexes compared to NPC-bound NUP153. This suggests that the majority of regulatory NUP153-DNA interactions captured by ChIP-seq likely occur in nucleus interior.

OptiFoot relies on the newly developed enzyme, euMTase – a mutant of sequence-unspecific adenine N6-MTase EcoGII. To our best knowledge, this is the first demonstration of sequence-unspecific MTase catalyzing transfer of large functionalized groups to DNA and RNA and the first example of re-engineering cofactor specificity in such enzymes. In contrast to related enzymes, cytosine C5-MTases, that often require cofactor-binding pocket engineering, wild-type adenine N6-MTases often show enough activity with AdoMet analogs for in vitro experiments ^45, 46^, and the only other reported example of adenine N6-MTase engineering is that of a sequence-specific M.TaqI ^47^. However, as we demonstrated here, the reprogrammed cofactor specificity is essential for efficient labeling *in cellulo*, to eliminate competition by AdoMet.

Broad tolerance towards transferrable group structure allows flexible experiment design. The covalent nature of the modification ensures labeling stability and supports downstream analyses of labeled nucleic acids without signal loss or redistribution. Non-fluorescent affinity tags and biorthogonal reactive groups pave the way for pull-down experiments. The ability to modify both RNA and DNA enables either simultaneous or selective analysis of the two nucleic acids. We demonstrate this by enzymatic removal of DNA or RNA using DNase or RNase treatment, prior to imaging.

For genome-wide POI profiling, we chose OGM as a readout because it directly detects OptiFoot fluorescent labels on native DNA molecules bypassing affinity-based enrichment. OGM shares major advantages of long-read sequencing: it retains long-range connectivity on individual DNA molecules, is able to detect large structural variants and differentiate between haplotypes. We demonstrate a fair agreement between OptiFoot genome occupancy profiles of various POIs and published datasets However, OGM resolution is limited to ∼500 bp/px and few tools are available for analyzing continuous fluorescent profiles. Therefore, long-read sequencing platforms (ONT or PacBio), which can distinguish modified nucleobases, would be a more sustainable future option for direct readout of OptiFoot labels. A recent study has demonstrated feasibility to differentiate between DNA methylation and artificial modification with larger groups by ONT sequencing ^48^. euMTase can aid further developments in this direction, as it may be used to easily screen for modifications optimal for detection and, thanks to the absence of sequence specificity, can easily produce datasets for model training. This technology is of particular interest in RNA analysis, where numerous modifications occur naturally and thus possibility to record biorthogonal modifications is essential.

In summary, OptiFoot provides a unique combination of single-cell fluorescence imaging and single-molecule genome mapping, enabling the study of protein-nucleic acid interactions in space of cell nucleus and genomic context. This multimodal capability addresses a critical gap between nuclear architecture and genome-wide protein occupancy, offering a versatile platform for investigating chromatin organization, transcriptional regulation, and dynamic nuclear processes.

## AUTHOR INFORMATION

## Author Contributions

G.L. and R.G. conceived the project; R.G., J.B., D.B., T.G. and G.L. performed the experiments; R.G., J.B., D.B., and G.L. analyzed the data. R.G. wrote the initial draft; all authors contributed to the final version of the manuscript.

## ACKNOWLEDGMENTS

The work was funded by the Max Planck Society. The authors thank dr. Vladimir Belov, Jan Seikowski, Jens Schimpfhauser, and Jürgen Bienert from Facility for Synthetic Chemistry (MPI for Multidisciplinary Sciences) for the NMR/ESI-MS measurements; dr. H. Frauendorf from central analytics’ team (Institute for Organic and Biomolecular Chemistry, Georg-August University, Göttingen) for NMR and HRMS measurements; dr. Peter Lenart and dr. Antonio Politi from Facility for Light Microscopy (MPI for Multidisciplinary Sciences, Fassberg campus) for providing access to a spinning disk confocal microscope; dr. Marco Roose for the help with data analysis. The authors acknowledge ChatGPT (OpenAI) for assistance in polishing the grammar and style of the manuscript.

