## Supplementary information for "OptiFoot: a method for recording protein footprints on DNA for microscopy and sequence analysis"

### Supplementary schemes and figures

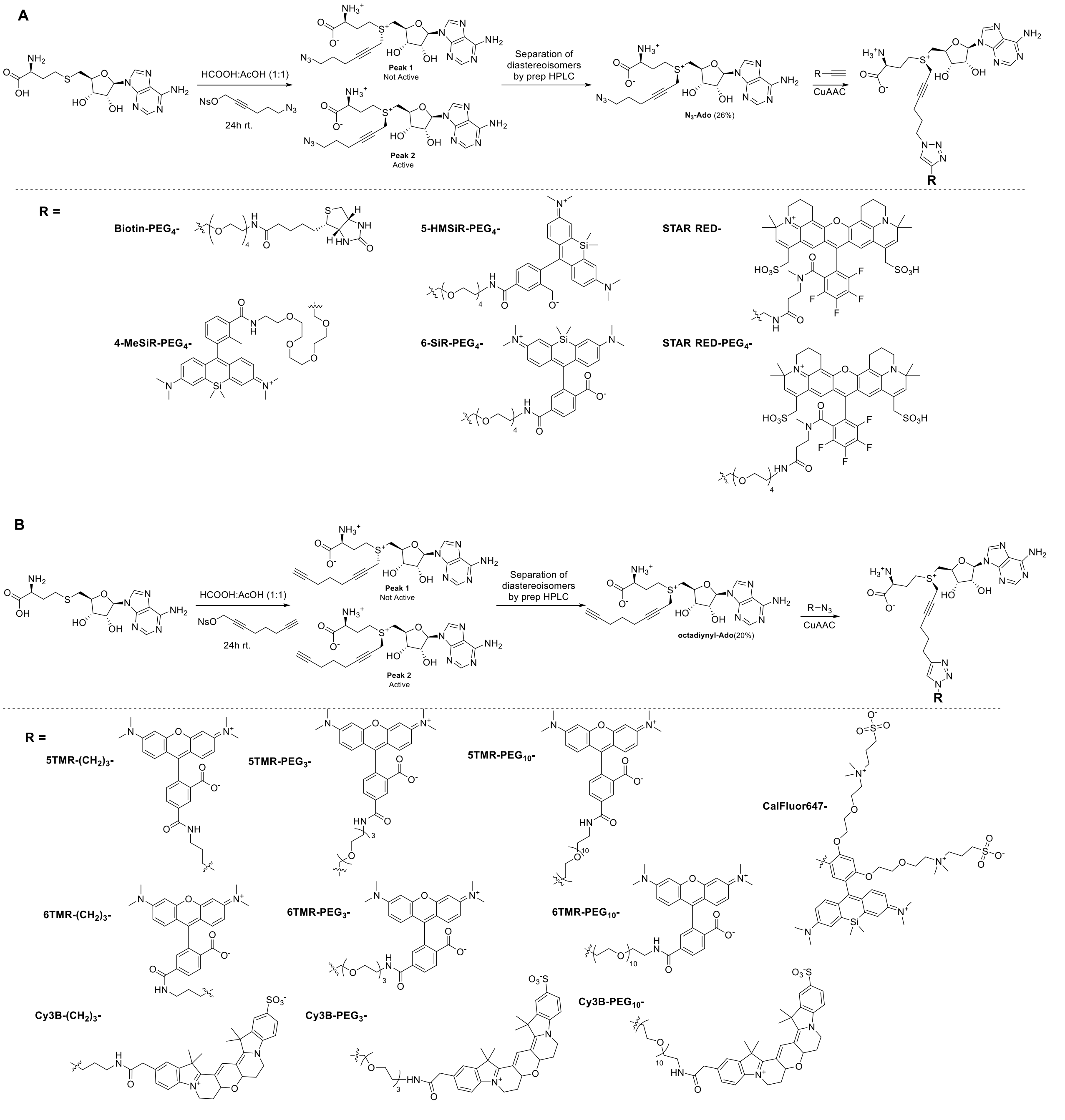

**Scheme S1. Synthesis and structure of functionalized AdoMet analogs.**

**A**, cofactors synthesized from $N_3$-Ado via Copper-catalyzed azide-alkyne cycloaddition (CuAAC) reaction.

**B**, cofactors synthesized from octadiynyl-Ado via CuAAC reaction.

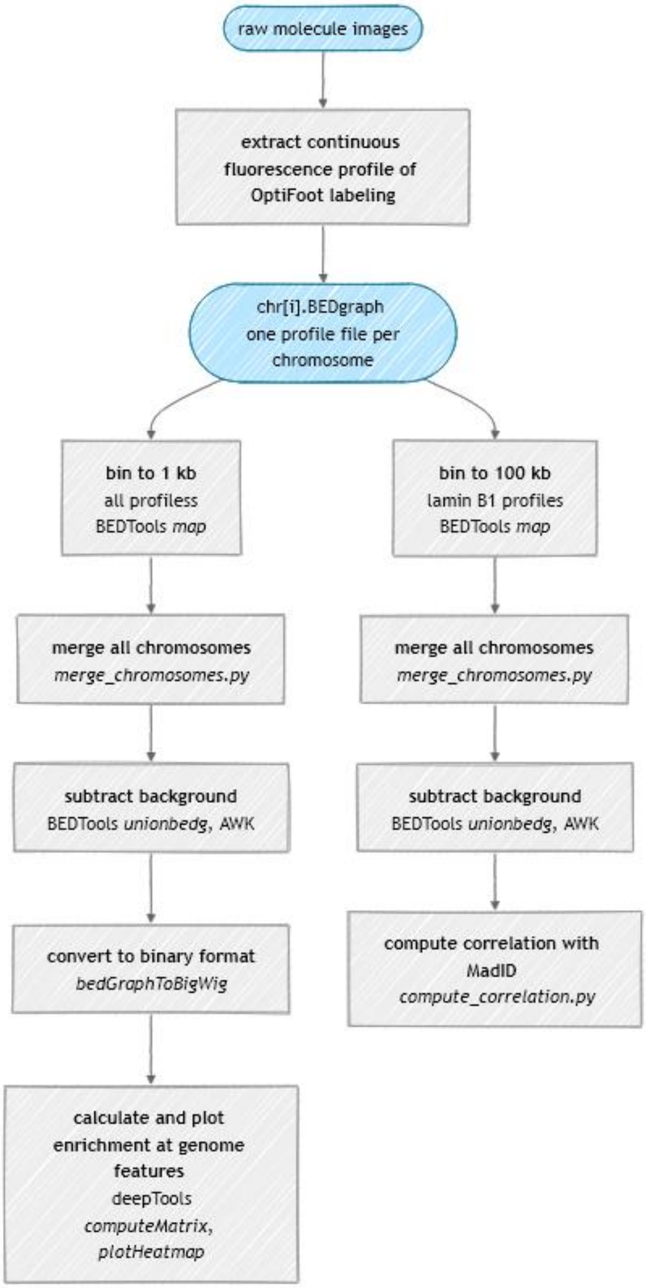

**Scheme S2. Data processing and analysis workflow for protein-genome interaction profiling by optical genome mapping.**

Analysis starts with raw images acquired on Saphyr instrument (Bionano Genomics). The first step, extracting continuous OptiFoot fluorescence profiles of all molecules, aligning them to genome reference and averaging signal mapping to the same genome loci. This is accomplished by a pipeline composed of commercial software and custom scripts that is outlined in **Scheme S2**. Subsequent steps involve custom Python scripts and conventional genome analysis tools – BEDTools and deepTools.

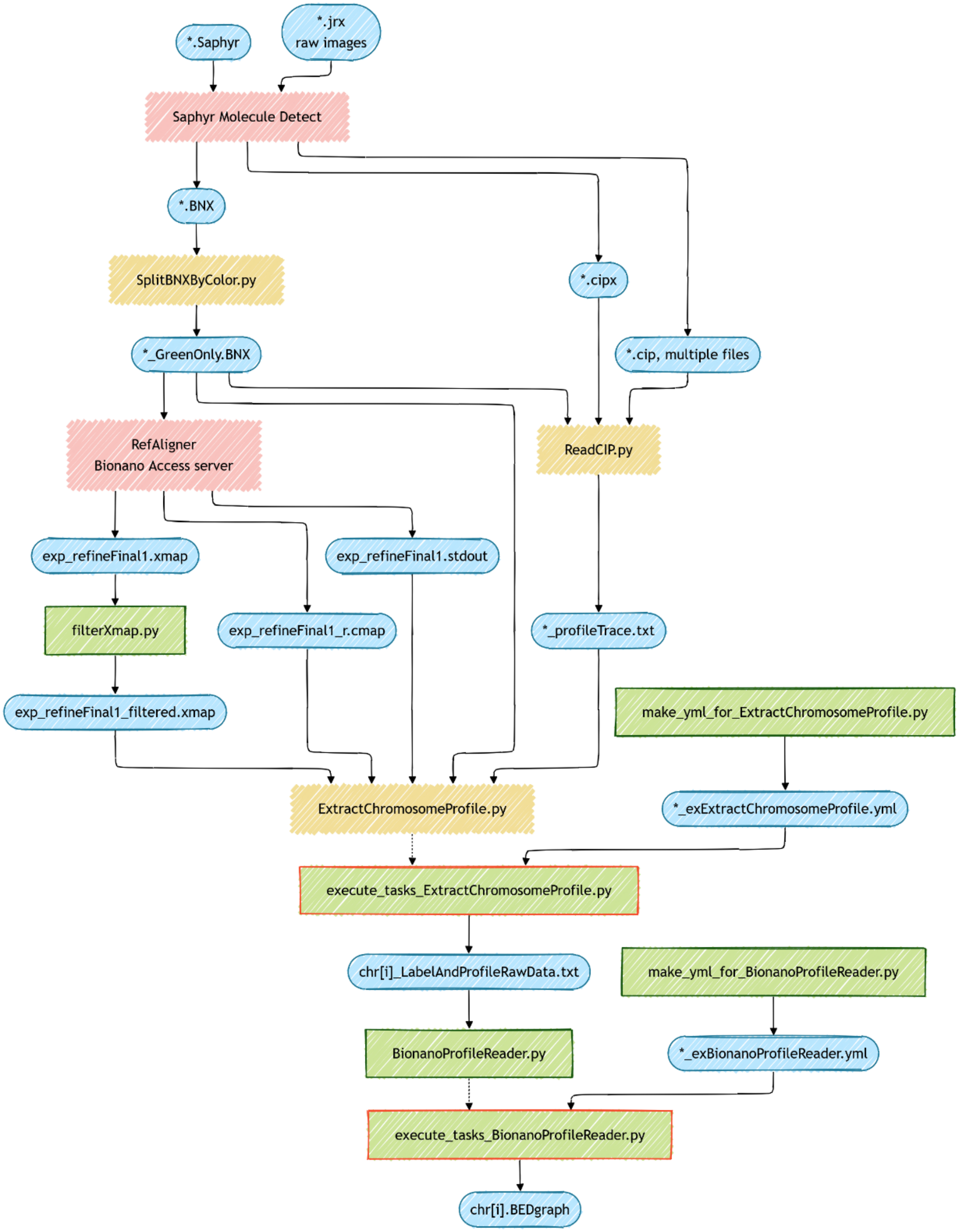

**Scheme S3. Extraction of aggregated continuous OptiFoot labeling profile aligned to genome reference.**

The pipeline starts with 3-color images (*.jrx) captured by Saphyr instrument and database file generated during the run. Software from Bionano Genomics is depicted in red boxes, python scripts originally provided by Bionano Genomics are in yellow boxes and custom python scripts are in green boxes. Red boxes denote scripts for processing individual chromosome files in parallel. Input and output files are in blue rounded boxes. In cases where output consists of multiple files, only the files relevant for this pipeline are shown. Information about the file formats is available from Bionano Genomics.

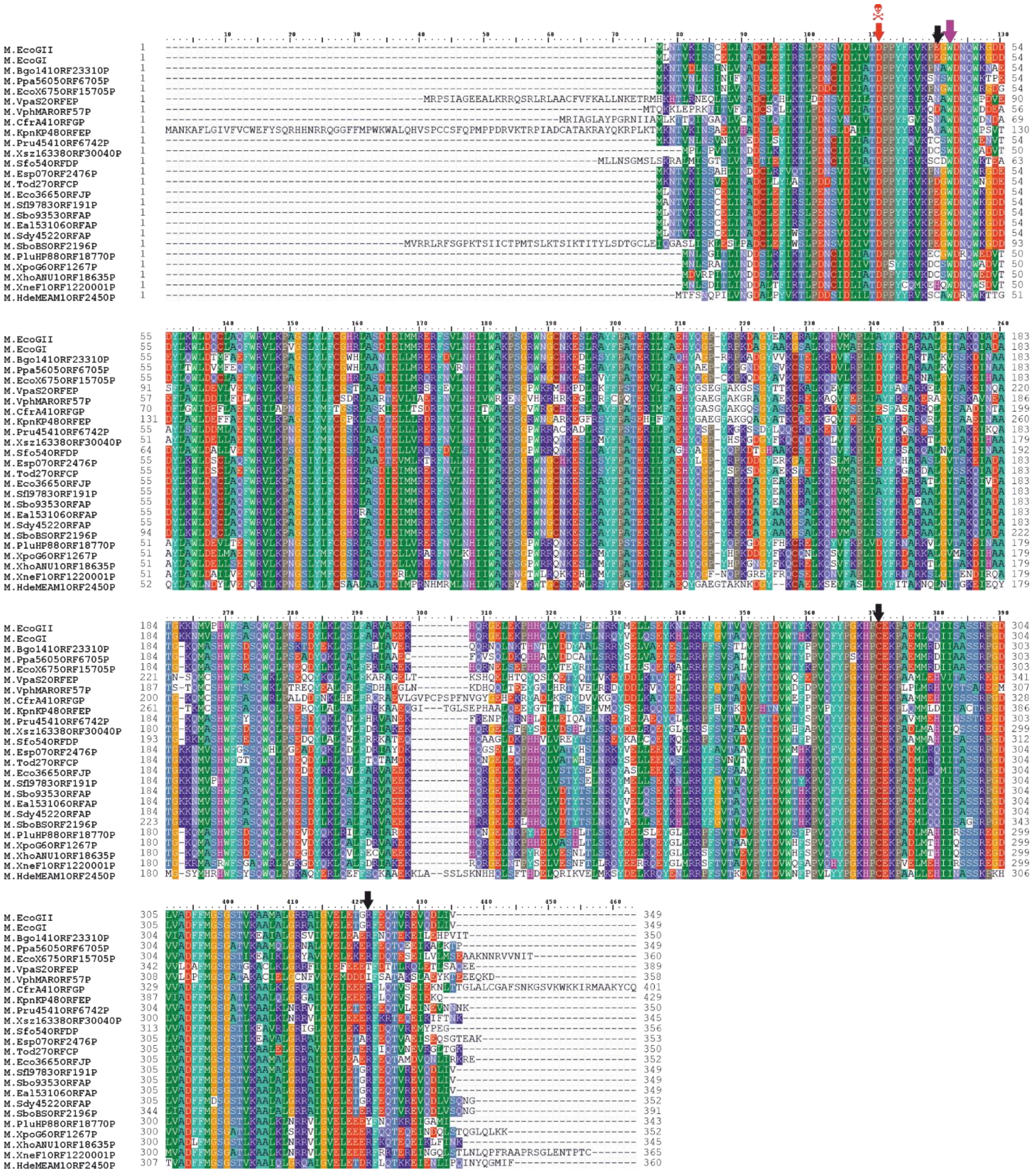

**Fig. S1. Sequence alignment of the closest M.EcoGII homologs.**

Arrows point to the residues mutated in this study: magenta – W46, mutated in euMTase; black – E44, C285 and R336; red – catalytic D35. MTase sequences [1] were downloaded from REBASE and aligned within AlignX (a component of Vector NTI Advance 11.5.0). Residues are shaded at 30% similarity using BLOSUM62 matrix. The figure was prepared with BioEdit v.5.0.9.

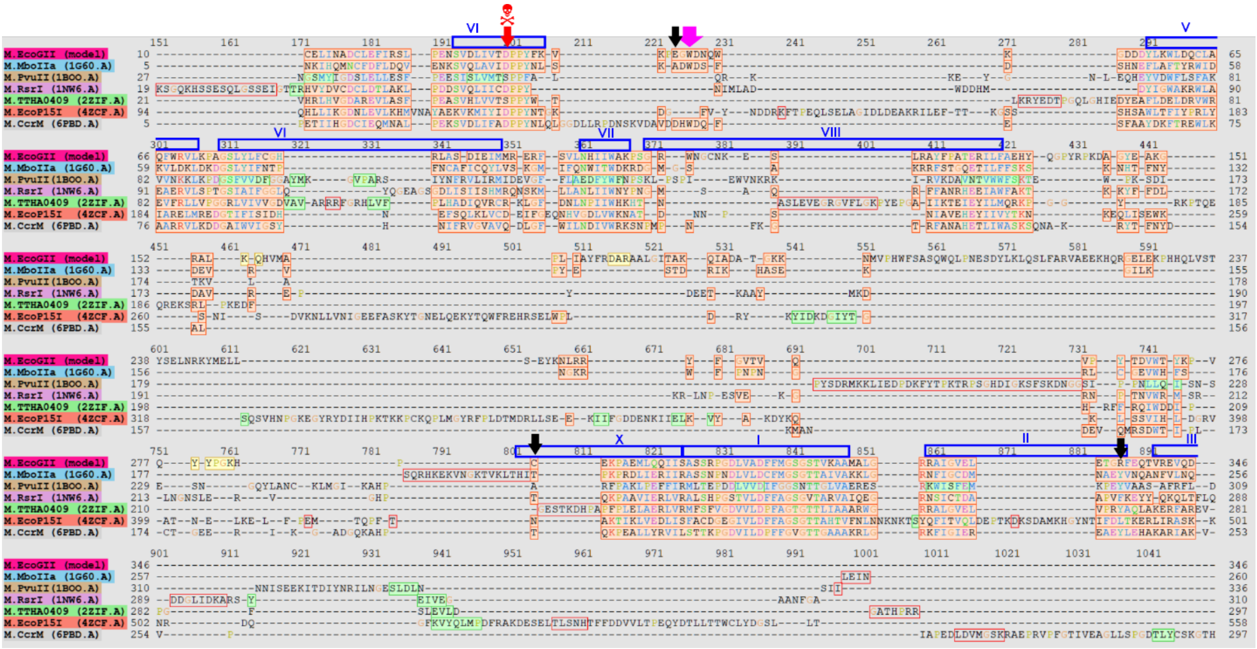

**Fig S2. Identification of structurally conserved regions in M.EcoGII.**
Structural alignment of $\beta$-class DNA methyltransferases with experimentally determined 3D structures. Structural alignment was produced using Pairwise Structure Alignment tool from PDB using TM-align method [2]. Arrows point to the residues mutagenized in this study: magenta – W46, mutated in euMTase, black – E44, C285 and R336, red – catalytic D35. Highly structurally conserved residues are shaded in orange, residues with missing structure are in red boxes. Blue boxes with Roman numbers mark ten structurally and functionally conserved amino acid motifs from DNA MTase family [3]. Alignment visualized with UCSC Chimera 1.13.1 [4].

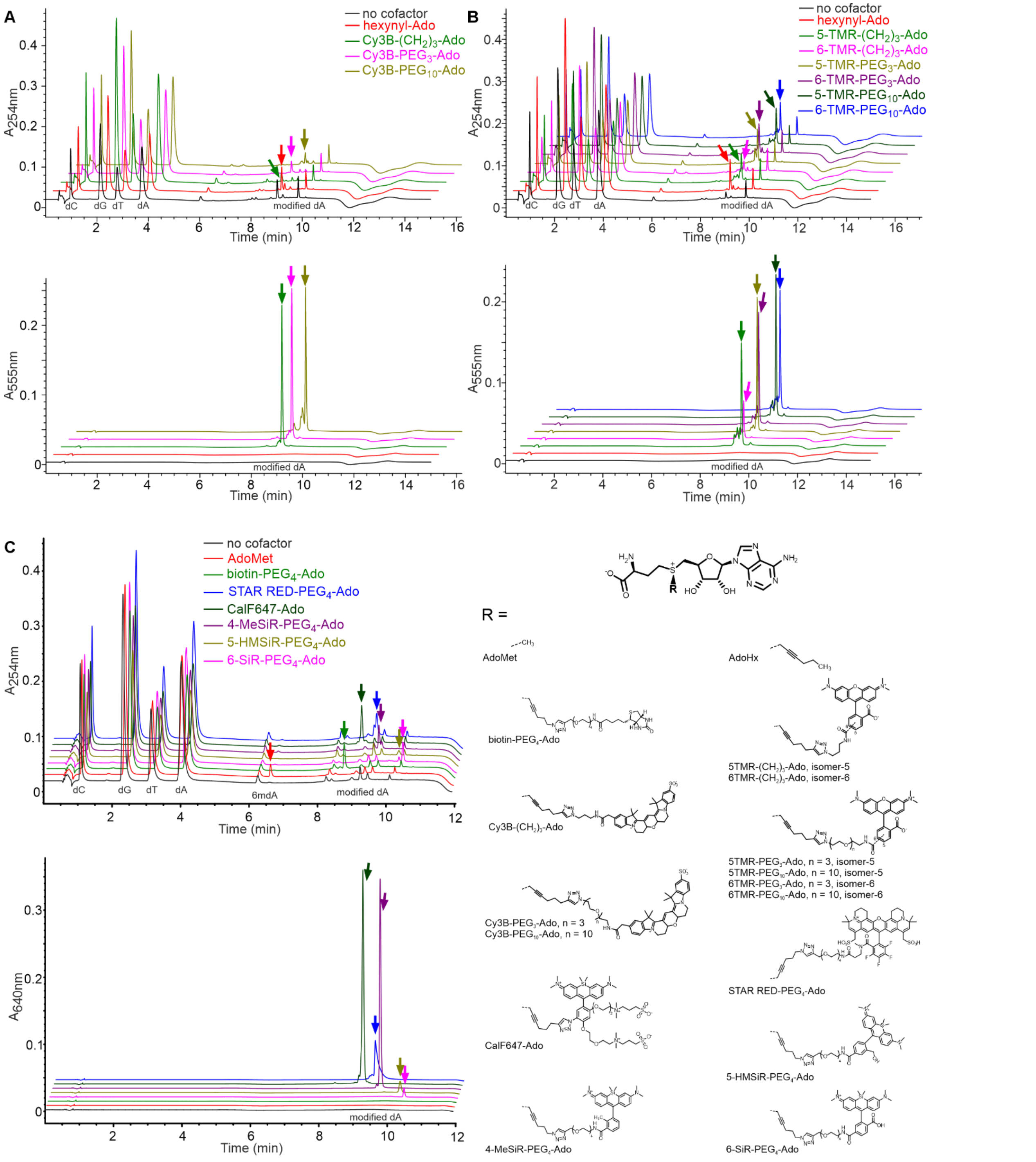

**Fig S3. Modification of λ DNA by euMTase with AdoMet analogs.**

**A-C**, representative analytical HPLC traces. Reactions contained 0.12µg/µl λ DNA, 500 nM euMTase and 100 µM (A, B) or 50 µM (C) of indicated cofactor and were incubated at 37°C for 1h. After reaction, DNA was hydrolized to nucleosides and analyzed by LC-MS. Identity of deoxyadenosine modification was confirmed by mass spectra (**Fig. S4**).

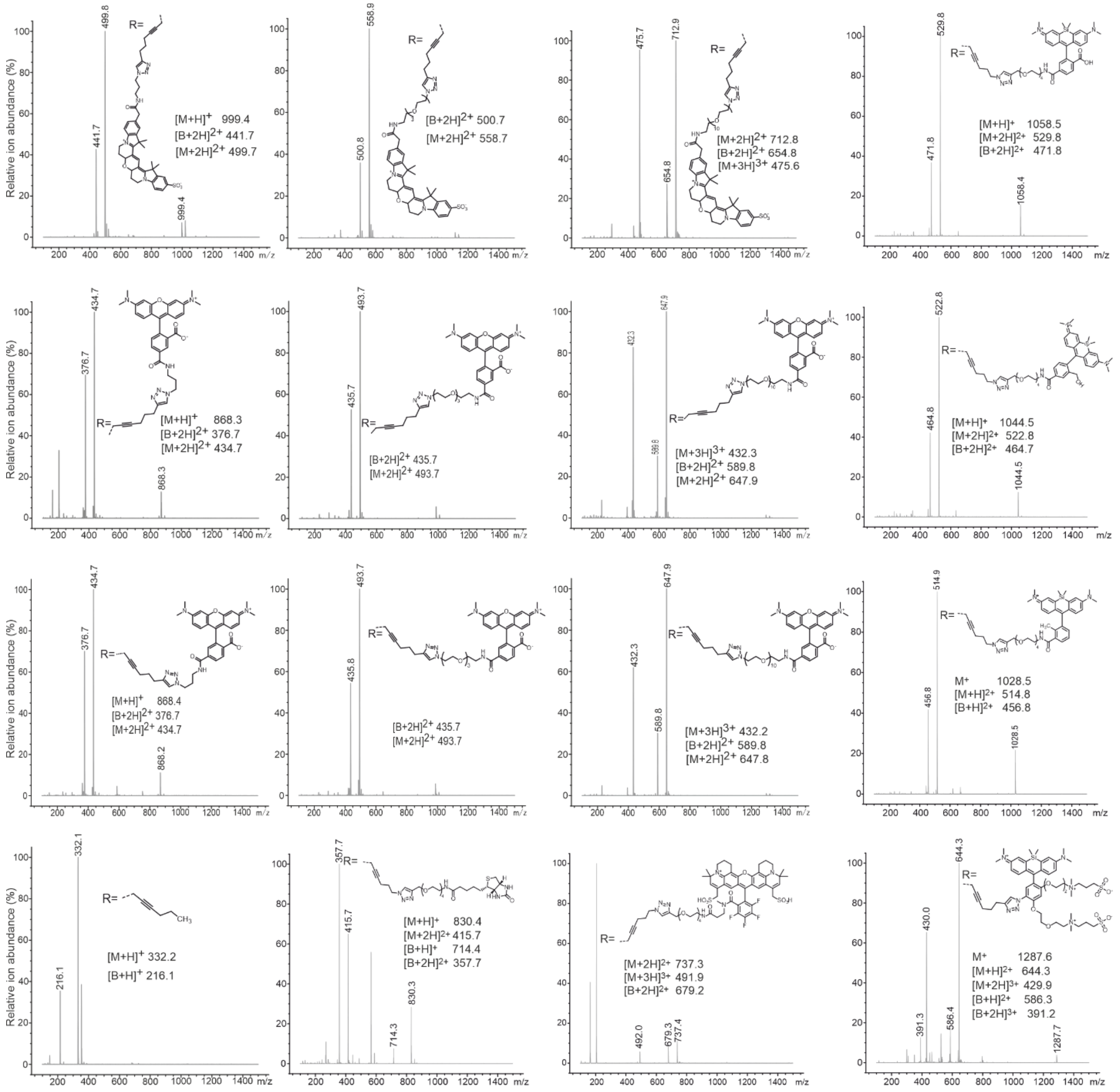

**Fig. S4. MS identification of enzymatic deoxyadenosine modifications produced by euMTase in the presence of AdoMet analogs.**

LC/MS traces are shown in **Fig. S3**. Theoretical m/z values: $[M+H]^+$ – protonated molecular ion, $[B+H]^+$ – protonated nucleobase formed upon fragmentation in the mass spectrometer.

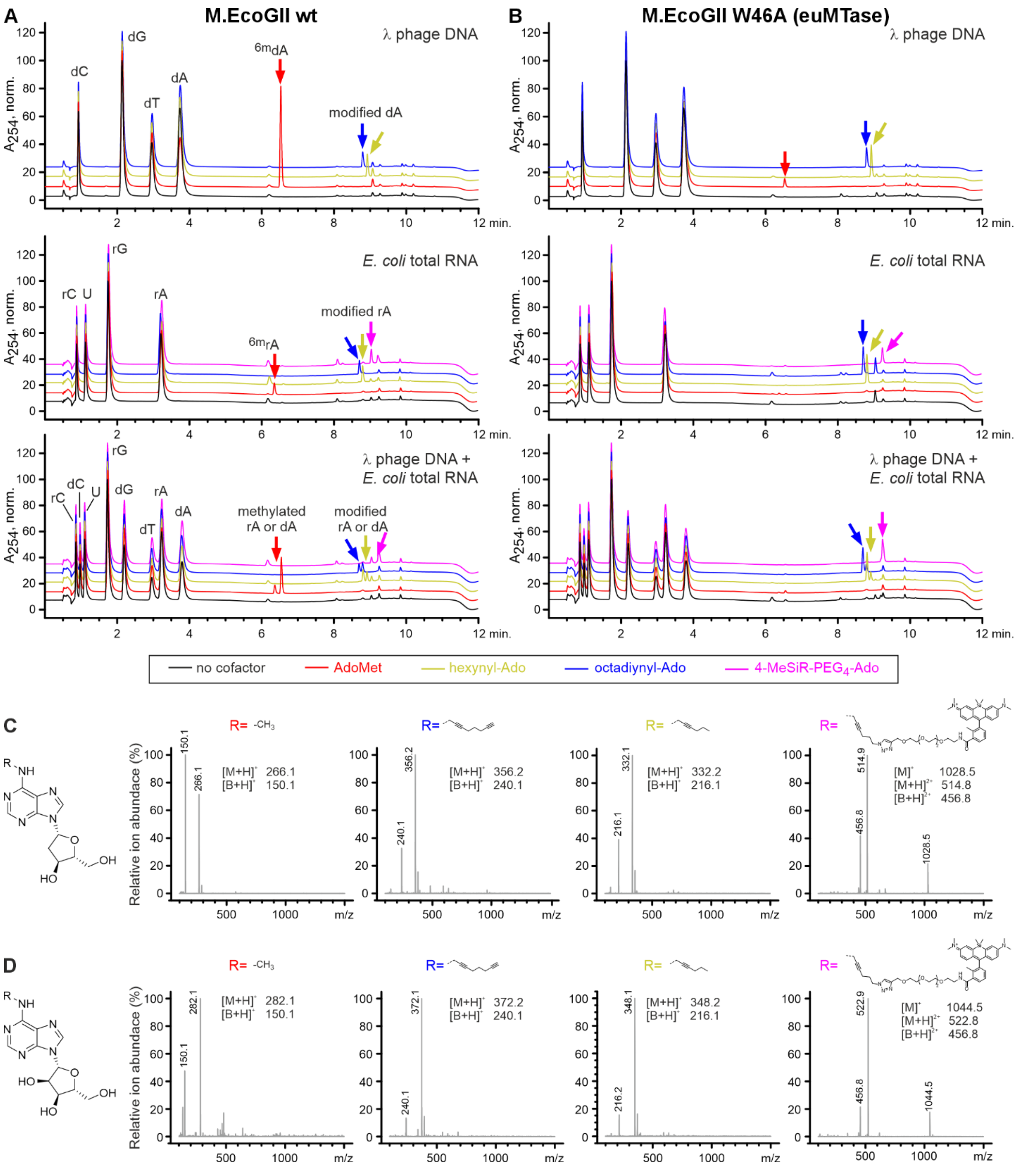

**Fig. S5. Modification of DNA, RNA and DNA-RNA mixture by wt M.EcoGII and euMTase.**
**A-B**, Normalized $A_{254\ nm}$ HPLC traces. Reactions contained 500 nM MTase, 100 µM cofactor and 0.12 µg/µl λ DNA or 0.1 µg/µl *E. coli* total RNA, or mixture of both. **C-D**, MS identification of enzymatic modification products: **C**, modified deoxyadenosines from DNA; **D**, modified adenosines from RNA. R – transferable chain. Theoretical m/z values: $[M+H]^+$ – protonated molecular ion, $[B+H]^+$ – protonated nucleobase formed upon fragmentation in the mass spectrometer.

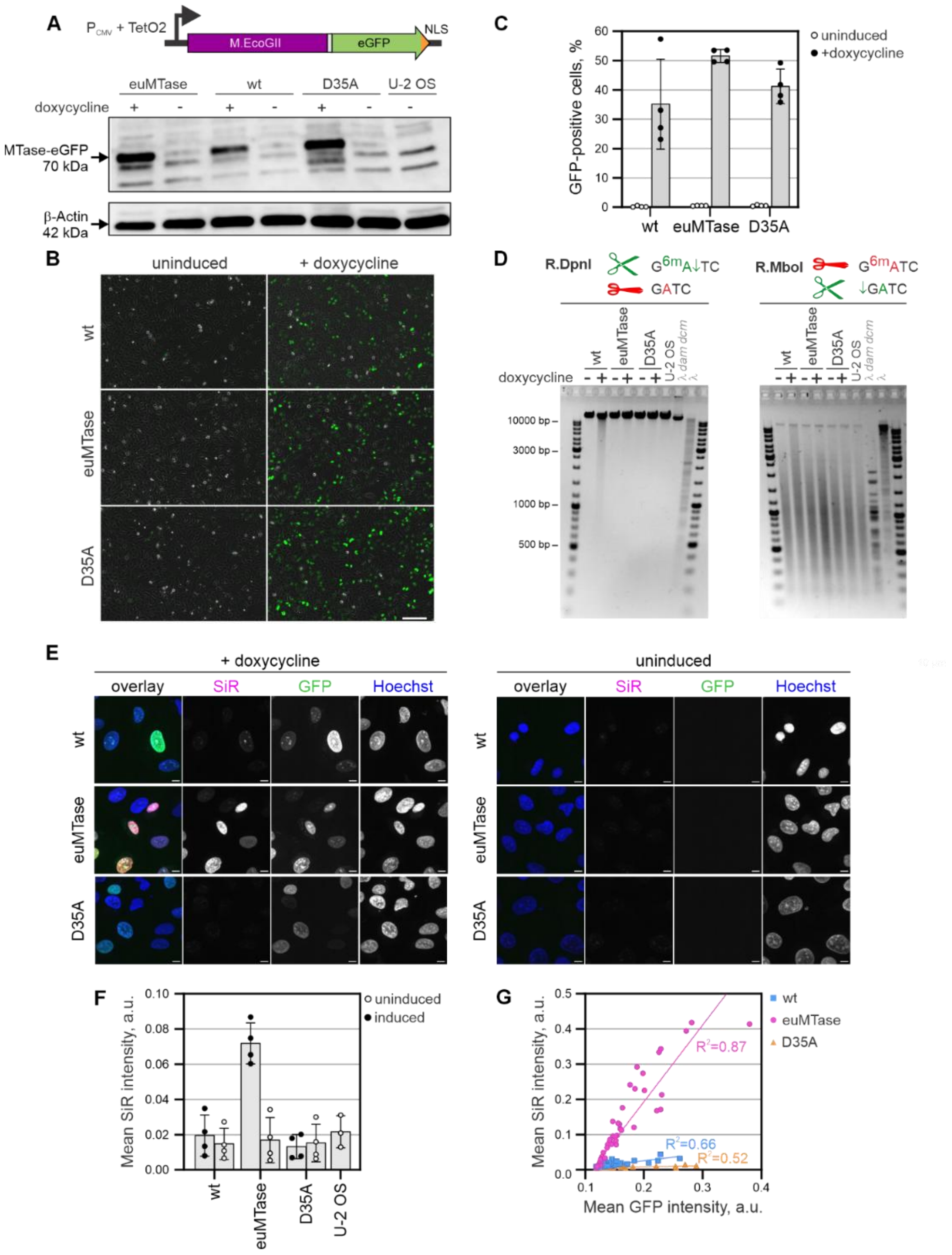

### Fig. S6. euMTase is superior to wt M.EcoGII for DNA labeling in permeabilized cells

**A**, schematic representation of expression construct and confirmation of expression in U-2 OS cells by Western blotting with anti-GFP antibody. **B**-**C,** detection of MTase-GFP-NLS expression by wide field fluorescence microscopy. **B**, Overlay of phase contrast (grayscale) and GFP (green) channels, scale bar 200 µm; **C**, quantification of transgene expressing cells. N=4 independent experiments, errors ± s.d. **D**, 6mA is detected in cells expressing wt M.EcoGII, but not euMTase. A representative image from three independent experiments is shown. **E**, labeling DNA with3 µM 6-SiR-PEG$_4$-Ado in digitonin-permeabilized cells. Sum-intensity z-projections, scale bar 10 µm. **F**, Quantification of MTase staining. Mean signal per nucleus ± s.d. **C**, Correlation between MTase expression (GFP channel) and DNA labeling (SiR channel). A representative example from 4 independent experiments is shown.

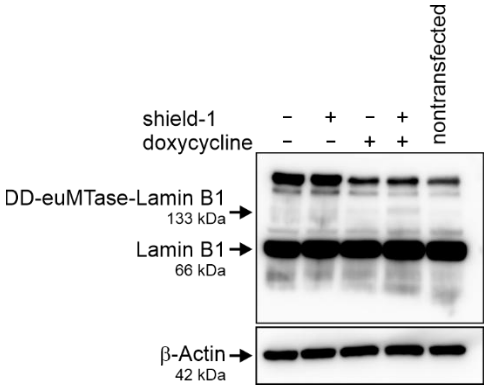

**Fig. S7. Verification of DD-euMTase-hLamin B1 expression by Western blotting.**
To induce transgene expression, the cells were incubated with 0.1 µg/ml doxycycline and 500 nM Shield-1 for 24 h. Low level of tagged protein can be explained by highly mosaic expression in the population (see **Fig. 2b** in the Main text).

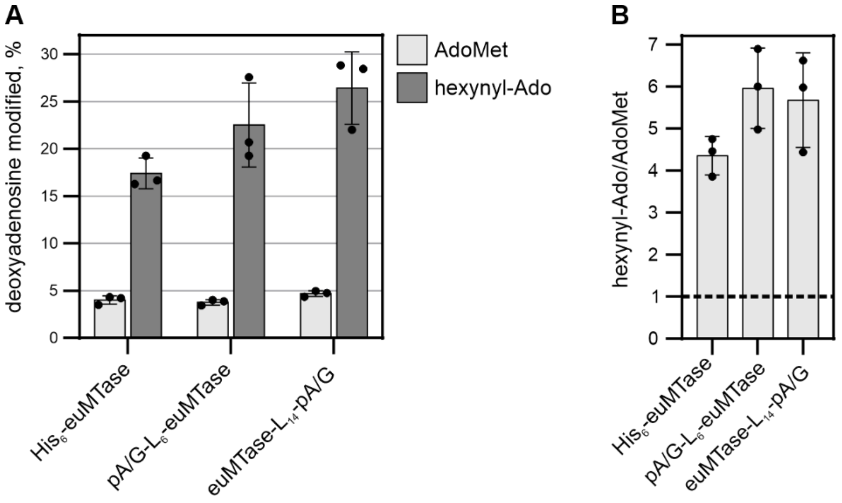

**Fig. S8. Tagging with protein A/G has no effect on euMTase activity.**
**A**, comparison of adenine modification in λ DNA with AdoMet and hexynyl-Ado, as determined by LC/MS analysis of hydrolyzed DNA. **B**, preference of hexynyl-Ado over AdoMet, calculated from data in **A**. Dashed line shows no preference. Reaction containing 0.12 µg/µl λ DNA, 500 nM MTase and 50 µM cofactor were incubated for 1 h at 37°C.

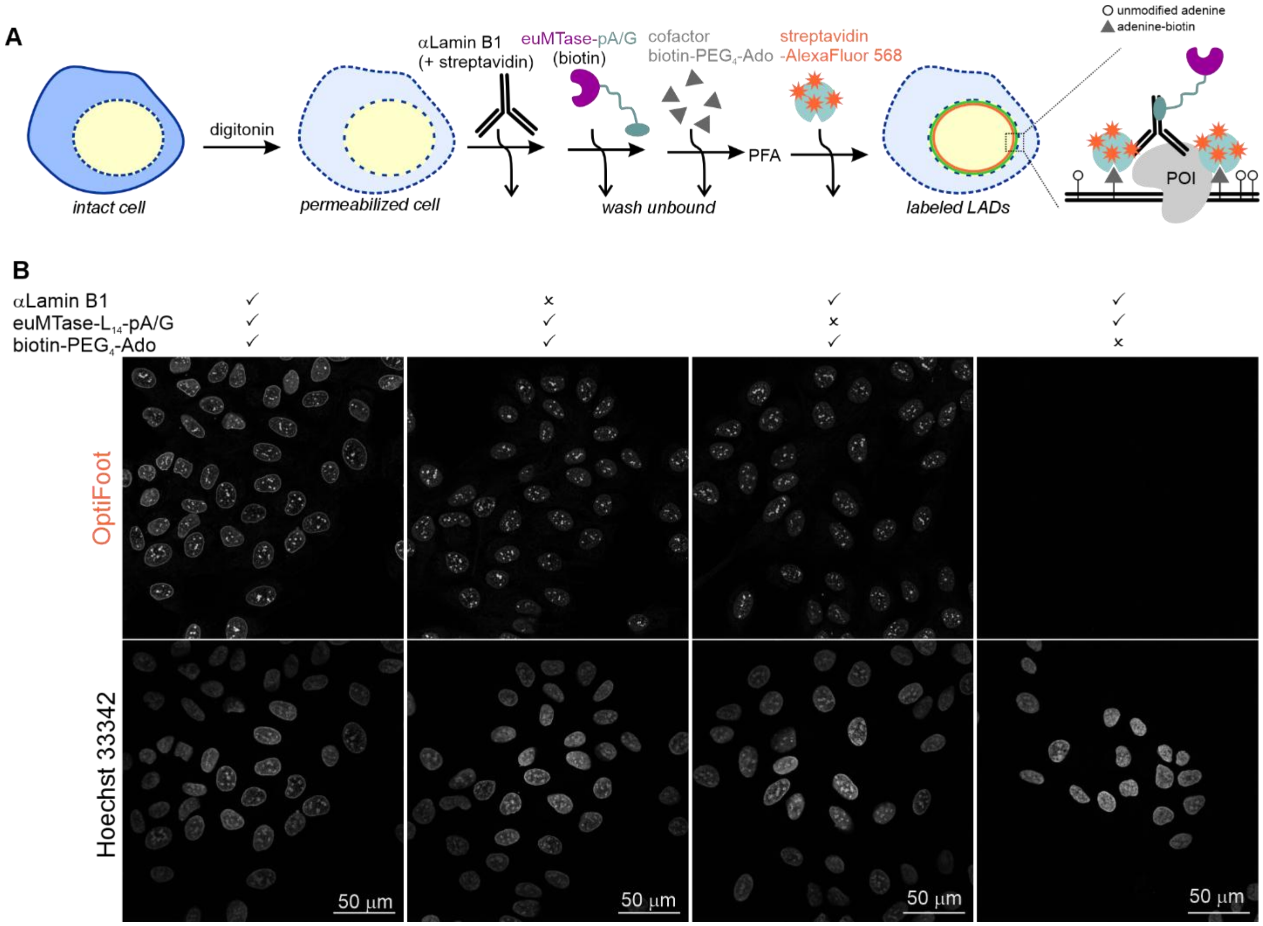

**Fig. S9. LAD labeling through biotin-streptavidin interaction.**

**A**, experiment workflow. Note, that to block endogenous biotin, streptavidin was included during incubation with antibody. Residual streptavidin was quenched by including biotin in the incubation with MTase. **B**, confocal slices through nucleus mid-section. Key components that were excluded in the control reactions are indicated above the images.

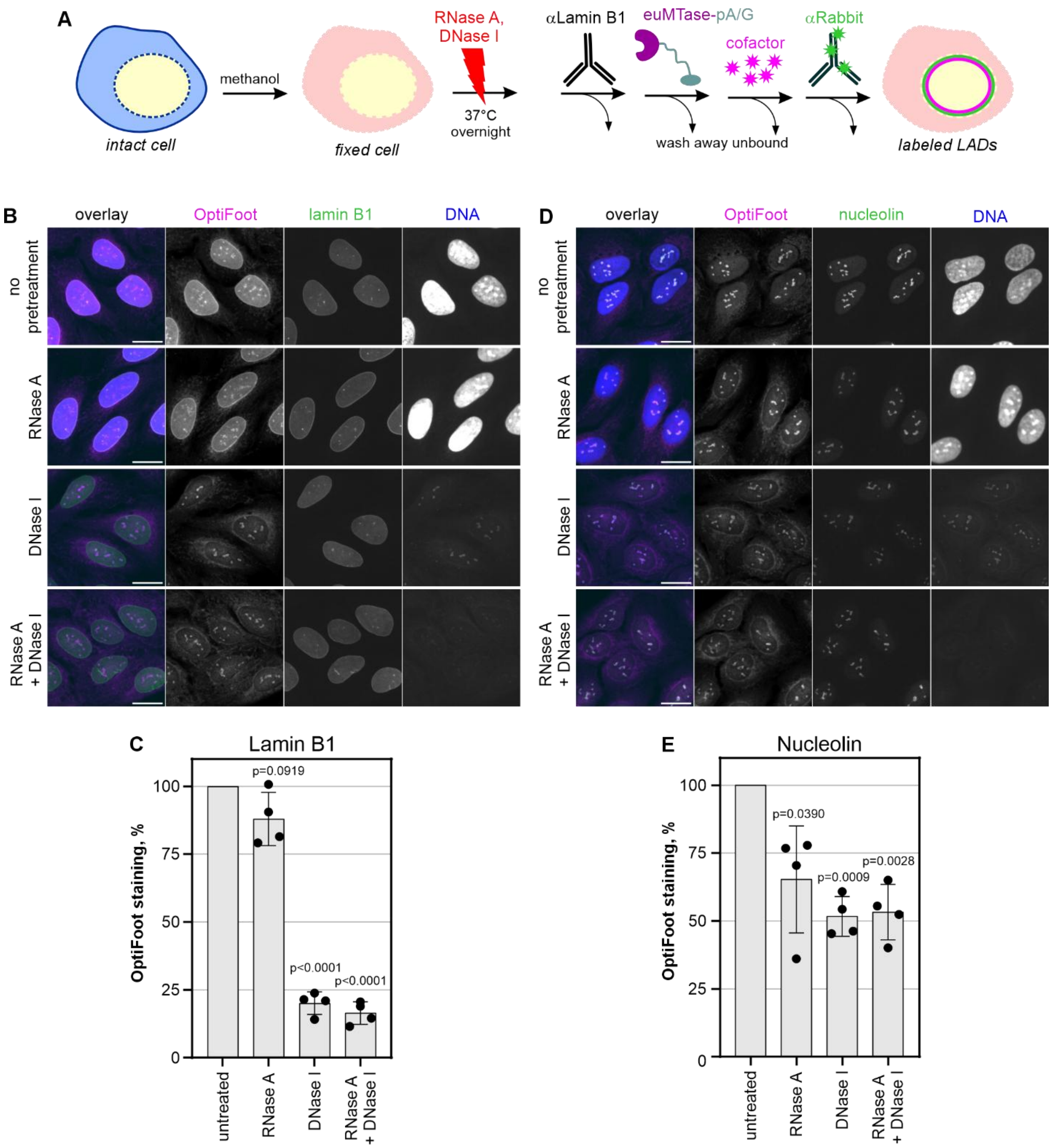

**Fig. S10. OptiFoot staining mainly originates from labeled DNA and not RNA.**

**A**, experimental workflow. **B, C** – example images and quantification of OptiFoot staining targeted to lamin B1 (polyclonal antibody). **D**, **E** – example images and quantification of OptiFoot staining targeted to nucleolin. Images were taken on a spinning disk confocal microscope as z-stacks, sum of slices were used for quantification and are shown. Scale bar – 20 µm. Four independent experiments were performed. OptiFoot staining after treatment (mean gray values) was normalized to a corresponding value measured on untreated cells. One sample t-test was used to assess difference from 1, and *p* values are reported.

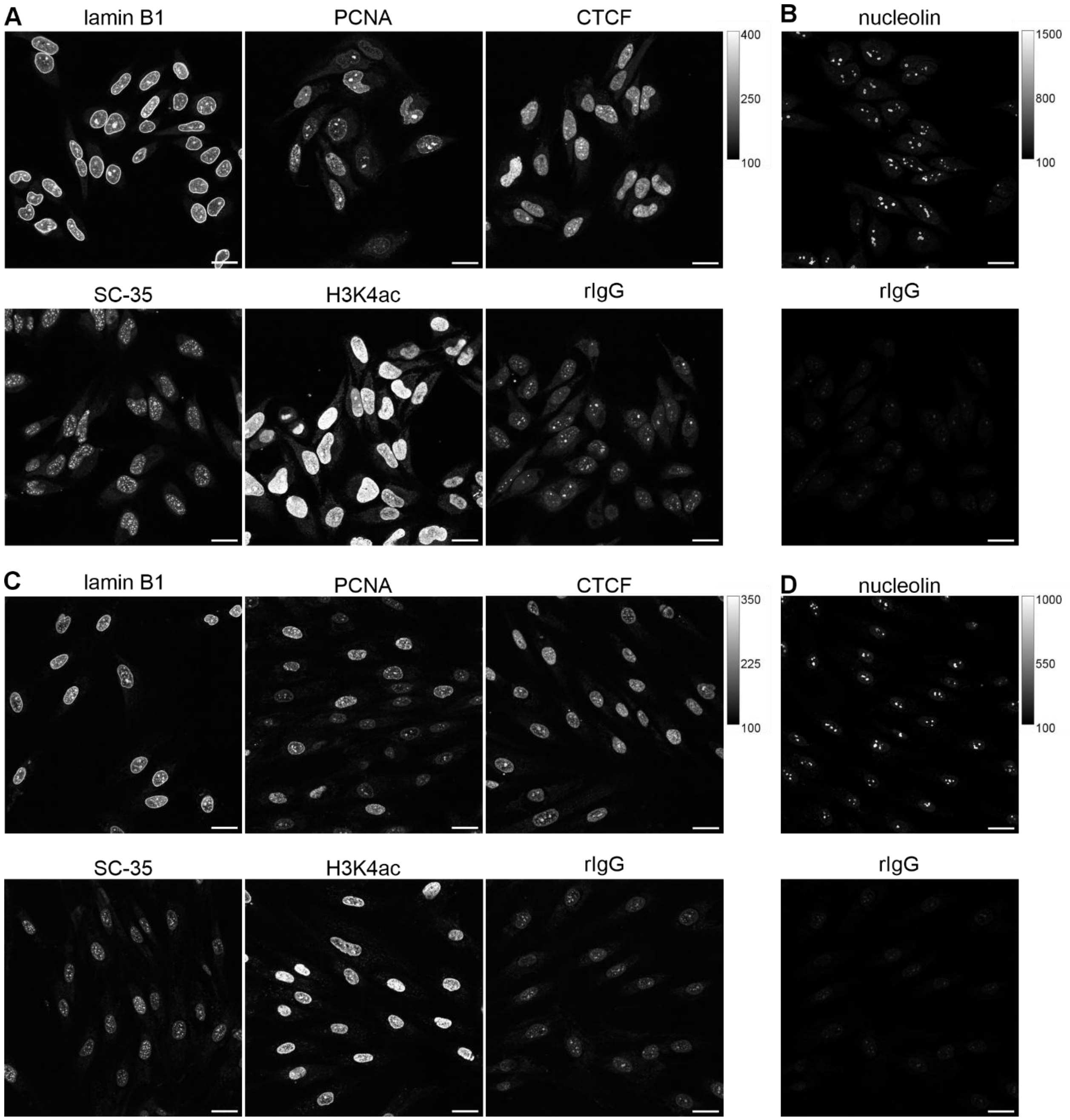

**Fig. S11. OptiFoot staining of POI-genome interactions in Hela (A,B) and human primary fibroblasts (C,D).**

Images were acquired on spinning disk confocal microscope as z-stacks and slices at cell mid-sections are shown, scale bar – 25 µm. All images were acquired with the same microscope settings, only representation settings differ as indicated by the calibration bars. Note, although untargeted staining by cofactor is detectable in nucleoli (**A**, **C**, rIgG sample), the specific OptiFoot staining when targeted with anti-nucleolin antibody is much stronger (**B**, **D**).

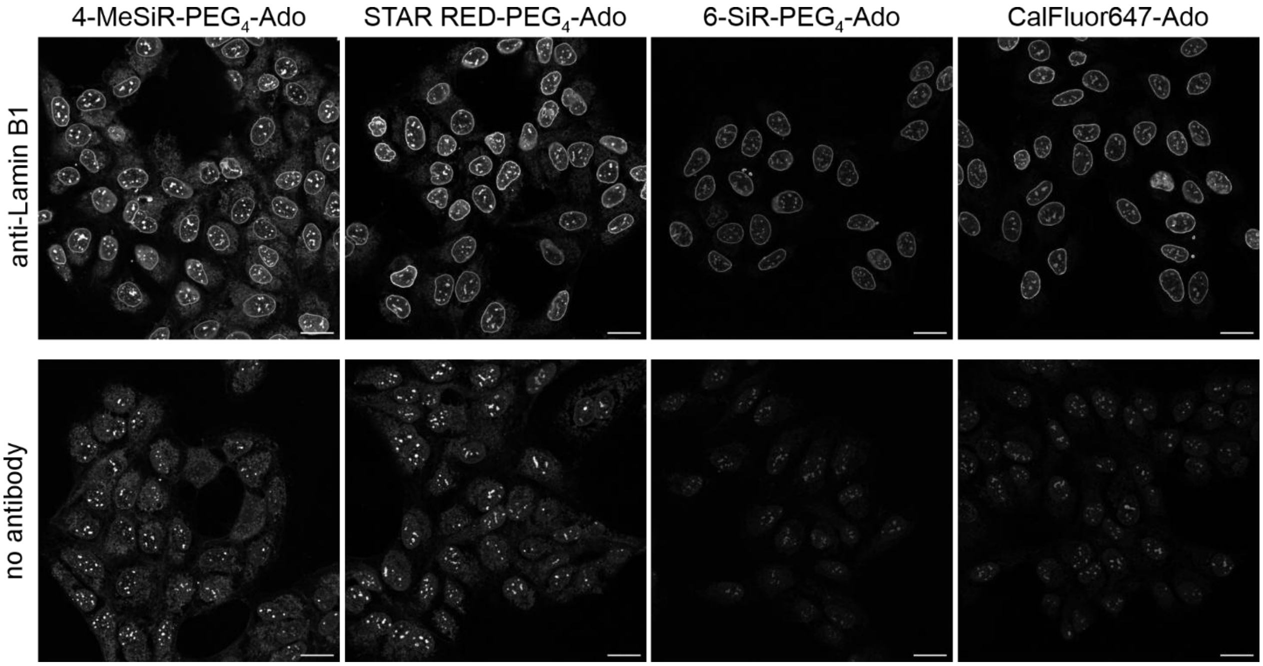

**Fig. S12. Performance of four red fluorophores used in OptiFoot labeling for microscopy.**
U-2 OS cells were permeabilized with 0.02% digitonin and OptiFoot staining was targeted with polyclonal antibody against lamin B1 (1:1000), followed by incubation with 10 nM euMTase-$L_{14}$-pA/G. Staining was performed with 3 µM of indicated cofactor for 1h at 37°C, then the cells were fixed with PFA and imaged. Confocal z-slice through cell mid-section is shown. Scale bar – 25 µm.

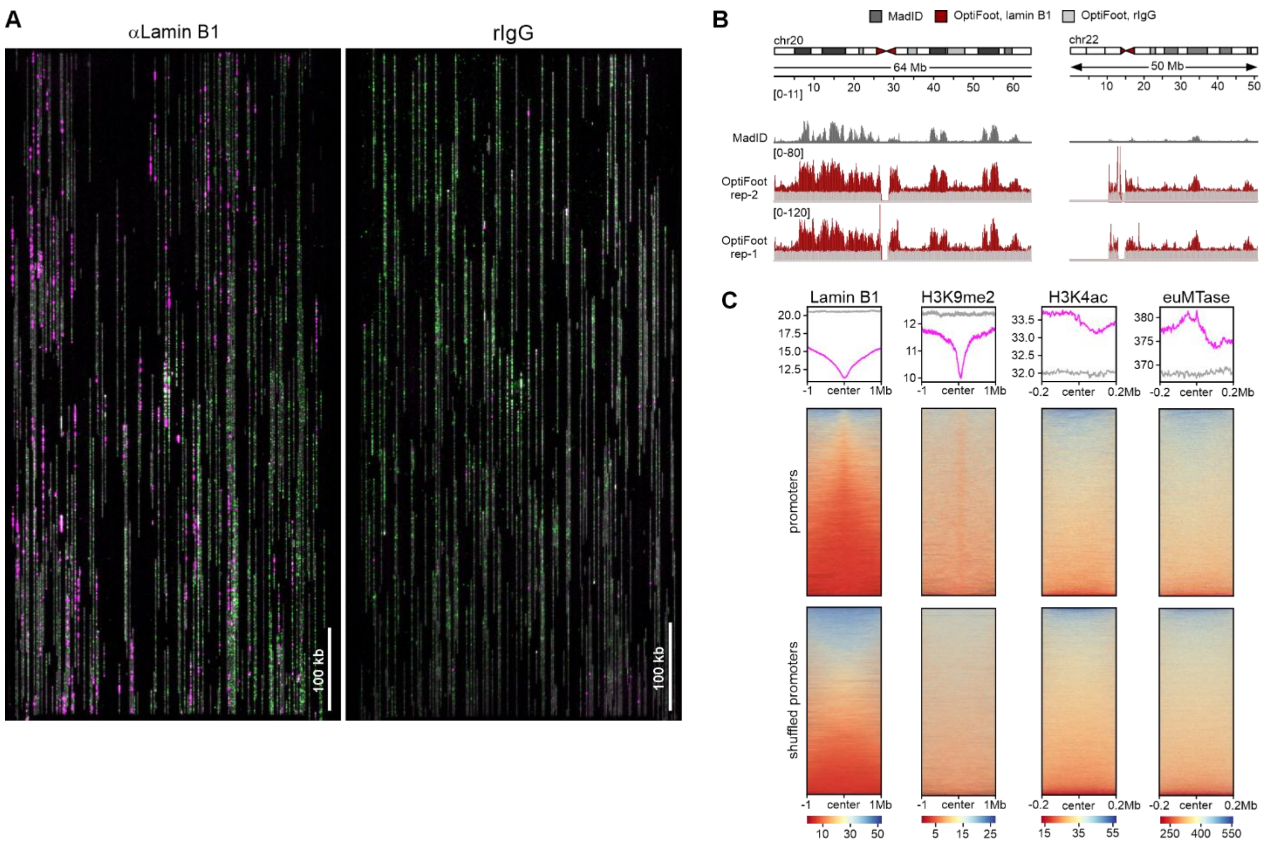

**Fig. S13. Profiling protein interactions over genome with OptiFoot and OGM.**

**A**, raw images of molecules extended in the nanochannels captured by Saphyr instrument. Gray – YOYO staining, green – reference DLE-1 labels, magenta – OptiFoot labels. **B**, Lamin B1 OptiFoot profiles obtained with polyclonal antibody over two small chromosomes with different LAD content: chr20 – LAD-rich, chr22 – LAD-poor. MadID profile of Hela cells [5] is shown for comparison. The profiles are depicted in the same scale as those in the Figure 6. **C**, Enrichment of OptiFoot signal targeted at various POIs in promoters from Homo sapiens (human) curated promoter database.

### Supplementary tables

**Table S1. OptiFoot samples for deriving genome-wide interaction profiles by optical genome mapping.** The Avg. OptiFoot label density /100kbp value is calculated by Bionano Access v1.7.2 (Bionano Genomics) during OGM run.

| | | | OptiFoot labeling conditions | | | | | | |
|---|---|---|---|---|---|---|---|---|---|
| **Sample No.** | **Groups*** | **Unique sample ID** | **Digitonin, %** | **Antibody** | | **10 nM MTase** | **Cofactor** | | **avg. OptiFoot label density /100kbp** |
| 1 | I | **rIgG-01a** | 0.02 | rIgG | 1:1000 | euMTase-$L_{14}$-pAG | STAR RED-$PEG_4$-Ado | 3 µM | 0.05 |
| 2 | | **aLmB1-01a** | 0.02 | Lamin B1 | 1:1000 | euMTase-$L_{14}$-pAG | STAR RED-$PEG_4$-Ado | 3 µM | 1.17 |
| 3 | II | **rIgG-03** | 0.08 | rIgG | 1:250 | euMTase-$L_{14}$-pAG | 4-MeSiR-$PEG_4$-Ado | 30 µM | 0.06 |
| 4 | | **aLmB1-04** | 0.08 | Lamin B1 | 1:250 | euMTase-$L_{14}$-pAG | 4-MeSiR-$PEG_4$-Ado | 30 µM | 2.06 |
| 5 | | **aH3K27ac-01** | 0.08 | H3K27ac | 1:250 | euMTase-$L_{14}$-pAG | 4-MeSiR-$PEG_4$-Ado | 30 µM | 0.37 |
| 6 | III | **rIgG-06** | 0.08 | rIgG | 1:1000 | euMTase-$L_{14}$-pAG | 4-MeSiR-$PEG_4$-Ado | 30 µM | 0.04 |
| 7 | | **aCTCF-01** | 0.08 | CTCF | 1:500 | euMTase-$L_{14}$-pAG | 4-MeSiR-$PEG_4$-Ado | 30 µM | 0.93 |
| 8 | | **aH3K9me2-01** | 0.08 | H3K9me2 | 1:250 | euMTase-$L_{14}$-pAG | 4-MeSiR-$PEG_4$-Ado | 30 µM | 1.57 |
| 9 | | **aH3K4ac-02** | 0.08 | H3K4ac | 1:50 | euMTase-$L_{14}$-pAG | 4-MeSiR-$PEG_4$-Ado | 30 µM | 4.15 |
| 10 | IV | **euMTase-01** | 0.08 | NA | NA | $H_6$-euMTase | STAR RED-$PEG_4$-Ado | 30 µM | 18.0 |
| 11 | | **No MTase** | 0.08 | NA | NA | no MTase | STAR RED-$PEG_4$-Ado | 30 µM | 0.01 |

***** samples were grouped according to rIgG control that was prepared within the same series and was used for background subtraction

**Table S2. Plasmids produced in this study.**
Plasmid sequences available upon request. Protein sequences are provided in **Supplementary sequences** section. $H_{6/14}$ – polyhistidine tag, pA/G – protein A/ G hybrid, DD – Proteotuner$^{TM}$ destabilizing domain. bdSUMO domain is cleaved during protein purification and only the POI (shown in bold) is purified.

| ID | Vector | Expression | Expressed protein |
|---|---|---|---|
| pRG202 | pET15b | E.coli | $H_6$-M.EcoGII wt |
| pRG203 | pET15b | E.coli | $H_6$-M.EcoGII E44S R336S |
| pRG224 | pET15b | E.coli | $H_6$-M.EcoGII W46A |
| pRG225 | pET15b | E.coli | $H_6$-M.EcoGII E44S W46A R336S |
| pRG227 | pET15b | E.coli | $H_6$-M.EcoGII E44S C285S R336S |
| pRG229 | pET15b | E.coli | $H_6$-M.EcoGII D35A |
| pRG274 | pET30a | E.coli | $H_{14}$-bdSUMO-**gly-M.EcoGII W46A-linker$_{14}$-pA/G** |
| pRG280 | pET30a | E.coli | $H_{14}$-bdSUMO-**gly-pA/G-linker$_6$-M.EcoGII W46A** |
| pRG289 | pEBTet | mammalian | DD-$H_6$-M.EcoGII W46A-hLamin B1 |
| pDB009 | pEBTet | mammalian | $H_6$-M.EcoGII W46A-eGFP-NLS |
| pDB012 | pEBTet | mammalian | $H_6$-M.EcoGII wt-eGFP-NLS |
| pDB013 | pEBTet | mammalian | $H_6$-M.EcoGII D35A-eGFP-NLS |

### Table S3. Reference datasets used to benchmark OptiFoot genome profiles.

| Genome feature | Cells | Genome build | Technique | Source | Ref. | remarks |
|---|---|---|---|---|---|---|
| **Lamina-associated domains** | Hela CCL2 | hg38 | MadID | *BioStudies*, E-MTAB-6888 | [5] | experimental |
| **DNAse I hypersensitive sites** | U-2 OS | hg38 | DNAse-seq | GEO, GSM4221655 | [6] | experimental |
| **H3K27ac peaks** | U-2 OS | hg38 | ChIP-seq | GEO, GSM6836721 | [7] | experimental |
| **CTCF sites** | U-2 OS | hg19 | ChiP-seq | GEO, GSE87831 | [8] | experimental, uplifted to hg38 |
| **LBR1-LADs** | U-2 OS | hg19 | DamID | GEO, GSE87831 | [8] | experimental, uplifted to hg38 |
| **Promoters** | NA | hg38 | NA | hsEPDnew, the Homo sapiens (human) curated promoter database | [9] | curated database |

Datasets originally built using hg19 reference, were uplifted to hg38 with *liftOver* utility from UCSC Genome Browser.
Where required, the datasets were cleaned-up to include only the standard chromosomes in order to match OGM output.

### Table S4. Antibodies used in this study.

| Target protein | Supplier | Catalog number, RRID |
|---|---|---|
| **Primary antibodies** | | |
| **Lamin B1 (pAb)** | Abcam | Cat# ab16048, RRID:AB_443298 |
| **Lamin B1 [EPR22165-121] (mAb)** | Abcam | Cat # ab229025, RRID:AB_3083735 |
| **rIgG** | Abcam | Cat# ab171870, RRID:AB_2687657 |
| **Nucleolin** | Abcam | Cat# ab22758, RRID:AB_776878 |
| **PCNA** | Abcam | Cat# ab18197, RRID:AB_444313 |
| **SC-35** | Abcam | Cat# ab11826, RRID:AB_298608 |
| **β-Actin** | Sigma-Aldrich | Cat# A5441, RRID:AB_476744 |
| **AbFlex® CTCF antibody (rAb)** | Active Motif | Cat# 91286, RRID: AB_3216313 |
| **Histone H3K4ac (pAb)** | Active Motif | Cat# 39382, RRID: AB_2793236 |
| **Histone H3K27ac (pAb)** | Active Motif | Cat# 39134, RRID: AB_2722569 |
| **Histone H3K9me2 (pAb)** | Active Motif | Cat# 39753, RRID:AB_2793331 |
| **Nup153 (QE5)** | Novus | Cat# NBP2-89036, RRID:AB_3430447 |
| **GFP** | Abcam | Cat# ab290, RRID:AB_303395 |
| **DD Monoclonal Antibody (ProteoTuner)** | Takara | Cat# 631073 |
| **Secondary antibodies** | | |
| **Donkey anti-Rabbit IgG (H+L) Highly Cross-Adsorbed Secondary Antibody, Alexa Fluor™ Plus 488** | Thermo Fisher Scientific | Cat# A32790, RRID:AB_2762833 |
| **Donkey anti-Mouse IgG (H+L) Highly Cross-Adsorbed Secondary Antibody, Alexa Fluor™ Plus 488** | Thermo Fisher Scientific | Cat# A32766, RRID:AB_2762823 |
| **Goat anti-Rabbit IgG (H+L) Cross-Adsorbed Secondary Antibody, HRP** | Thermo Fisher Scientific | Cat# G-21234, RRID:AB_2536530 |
| **Goat anti-Mouse IgG (H+L) Cross-Adsorbed Secondary Antibody, HRP** | Thermo Fisher Scientific | Cat# G-21040, RRID:AB_2536527 |
| **Donkey IgG anti-Mouse IgG (H+L), in-house labeled with Abberior STAR 580** | Dianova | Cat# 715-005-151, RRID:AB_2340759 |
| **Donkey IgG anti-Rabbit IgG (H+L), in-house labeled with Abberior STAR 580** | Dianova | Cat# 711-005-152, RRID:AB_2340585 |
| **sdAb Anti-Rabbit IgG Halo fusion, labeled with Abberior STAR 580-Halo ligand** | NanoTag | Cat# N2441, RRID:AB_3668679 |

pAb – polyclonal antibody, mAb – monoclonal antibody, rAb – recombinant antibody, sdAb – single domain antibody.

### Supplementary methods

#### Chemical synthesis

*General synthesis procedure A: synthesis of dyes with terminal alkyne linker via peptide coupling*

Fluorescent Dye-COOH (1 eq. of 4-MeSiR-COOH, 6-SiR-COOH, 5-HMSiR-COOH or STAR-RED-COOH) was dissolved in dry DMSO (300-1000 µl) and DIPEA (5 eq) and then a solution of HATU (1.3 eq) in DMSO (300 µl) was added. The reaction mixture was stirred at room temperature for 2-5 minutes, afterwards solution of $NH_2$-$PEG_4$-alkyne or propargyl amino hydrochloride (1.4 eq.) was added and the mixture was stirred for 1h at room temperature. Reaction progress and completion was monitored by LC/MS. Then the reaction mixture was quenched by adding 50 µl of formic acid and diluted with water. The product was purified by preparative HPLC. Fractions containing pure product were merged and solvent was evaporated on rotary evaporator. The solid was dissolved in a MeCN/water 1:2 mixture and lyophilized.

*General synthesis procedure B: synthesis of dyes with terminal azido linker via peptide coupling*

Dye-NHS ester (1eq of 5-TMR-NHS, 6-TMR-NHS or Cy3B-NHS) was dissolved in dry DMSO (800 µL) and then solution of $NH_2$-$(CH_2)_3$-$N_3$, $NH_2$-$PEG_3$-$N_3$ or $NH_2$-$PEG_{10}$-$N_3$ (2 eq) was added and the mixture was stirred for 0.5-1 h at room temperature. Reaction progress and completion was monitored by LC/MS. The product was purified by preparative HPLC. Fractions containing pure product were merged and solvent was evaporated on rotary evaporator followed by evaporation on Biotage® V-10 Touch evaporation system.

*General synthesis procedure C: cofactor synthesis from $N_3$-Ado via CuAAC*

A 6.7 mM solution of $N_3$-Ado[10] in 10mM ammonium formate buffer (pH 3.6) was mixed with a solution (2 eq. in 10mM ammonium formate buffer, pH = 3.6) of 4-MeSiR-$PEG_4$-alkyne, 6-SiR-$PEG_4$-alkyne, 5-HMSiR-$PEG_4$-alkyne, STAR-RED-$PEG_4$-alkyne, STAR-RED-alkyne or Biotin-$PEG_4$-alkyne. The copper (I) containing solution was prepared by mixing fresh solutions of $CuSO_4 \cdot 5H_2O$ (1 mg in 10 µl of AF buffer), ascorbic acid (3 mg in 30 µl of AF buffer) and $(BimC_4A)_3$ (0.5 mg in 5 µl of water, Sigma Aldrich Cat. No.: 696854) followed by immediate color change. The obtained copper (I) solution was sonicated for 1 minute and then 20-30 µl was added to the reaction mixture. The reaction vial was sonicated for 2 min and then left at room temperature for 30-90 min. The progress of the reaction was monitored by LC/MS every 20 min, once the conversion reached >90% the reaction was diluted with water to a volume of 2 ml and the product was purified by preparative HPLC.

*General synthesis procedure D: cofactor synthesis from octadiynyl-Ado via CuAAC*

Octadiynyl-Ado[10] solution in 10 mM ammonium formate buffer (pH 3.5) was mixed with a solution of 5-TMR-$PEG_3$-$N_3$, 6-TMR-$PEG_3$-$N_3$, Cy3B-$PEG_3$-$N_3$, 5-TMR-$PEG_{10}$-$N_3$, 6-TMR-$PEG_{10}$-$N_3$, Cy3B-$PEG_{10}$-$N_3$ or CalFluor647-$N_3$ in 10mM ammonium formate buffer pH = 3.5, in case of 5-TMR-$(CH_2)_3$-$N_3$ or 6-TMR-$(CH_2)_3$-$N_3$ the 1-butanol:methanol:isopropanol mixture (440 µl, ratio of 1:4:4) was added to solubilize the dyes. The copper (I) containing solution was prepared by mixing fresh solutions of sodium ascorbate (100 mM in water, 100 µl), copper (II) acetate (100 mM in water, 100 µl) and BTTAA ligand (50 mM in DMSO, 400 µl, Jena Bioscience Cat. No. CLK-067-100) supplemented to 1 ml with 10mM ammonium formate buffer pH = 3.5. Next, copper (I) solution (450 µl, except in the reaction with CalFluor647-$N_3$ 20 µl of copper (I) solution was used) was added the reaction mixture. The reaction vial was vortexed and left at room temperature for 30 min. The progress of the reaction was monitored by LC/MS, once the conversion reached >90% the reaction product was purified by preparative HPLC.

*General synthesis procedure E: cofactor synthesis from octadiynyl-Ado via CuAAC*

Solution of octadiynyl-Ado in 10 mM ammonium formate buffer (pH 3.5) was mixed with a solution of Cy3B-$(NH_2)_3$-$N_3$ in 10mM ammonium formate buffer pH=3.5. In another Eppendorf, the aqueous solution of Cu (II) was prepared by mixing $CuSO_4{\cdot}5H_2O$ (100 mM in water, 60 µl) and sodium ascorbate (100 mM in water, 20 µL). The aqueous brown mixture then was vortexed for a few seconds and the solution turns orange/yellow. The prepared aqueous solution of Cu (II) (79 µL) was added to the octadiynyl-Ado and Cy3B-$(CH_2)_3$-$N_3$ mixture. The reaction vial was vortexed and left at room temperature for 30 min. The progress of the reaction was monitored by LC/MS, once the conversion reached >90% the reaction product was purified by preparative HPLC.

**Plasmid constructs**

WT and E44S R336S variants of EcoGIIM gene were codon-optimized for bacterial expression and synthesized at GenScript. The genes were subcloned into pET15b under control of IPTG-inducible T7 promoter with N-terminal hexahistidine tag followed by thrombin cleavage site to yield plasmids pRG202 and pRG203, respectively. Other mutations were introduced by QuickChange protocol in combination with Gibson assembly (plasmids pRG224, pRG225, pRG227 and pRG229). pRG274, encoding $H_{14}$-bdSUMO-M.EcoGII W46A-linker$_6$-pAG and plasmid pRG280, encoding $H_{14}$-bdSUMO-pAG-linker$_{14}$-M.EcoGII W46A are based on pET30a vector. $H_{14}$-bdSUMO fragment was from pDG02583 (a gift from Dirk Görlich (Addgene plasmid # 104129 ; http://n2t.net/addgene:104129 ; RRID:Addgene_104129) [11], protein A/G part was from pAG/MNase (a gift from Steven Henikoff (Addgene plasmid # 123461 ; http://n2t.net/addgene:123461 ; RRID:Addgene_123461)[12].

Constructs for eukaryotic expression (pRG289, pDB009, pDB012 and pDB013) are based on pEBTet vector [13]. Proteotuner™ destabilizing domain. (DD) and human *LAMIN B1* gene was from DD-DamWT-LMNB1-IRES2-mCherry (a gift from Aaron Streets (Addgene plasmid # 159601 ; http://n2t.net/addgene:159601 ; RRID:Addgene_159601)) [14], eGFP-NLS fragment was from pcDNA5-MTS-TagBFP-P2AT2A-EGFP-NLS-P2AT2A-mCherry-PTS1 (a gift from Andrea Musacchio (Addgene plasmid # 87829 ; http://n2t.net/addgene:87829 ; RRID:Addgene_87829)) [15]

Desalted DNA oligonucleotides were used as primers and were obtained from Sigma. All cloning steps were performed by Gibson assembly, using Gibson Assembly® Master Mix (New England Biolabs, #E2611L). The final constructs were verified by Sanger sequencing of the complete expression cassette. Plasmid constructs produced in this study are listed in **Supplementary Table 2**.

**Expression and purification of M.EcoGII variants**

The expression plasmids were transformed into *E.coli* BL21 (DE3), the cells were grown in 500 ml LB medium supplemented with Overnight Express™ Autoinduction System 1 – Novagen (Sigma-Aldrich #71300) and 100 µg/ml ampicillin at 21°C overnight, then lysed by sonication in buffer A (50 mM HEPES/NaOH or sodium phosphate pH 7.4 and 500 mM NaCl) and the proteins were purified by affinity chromatography. M.EcoGII wt, E44S R336S, E44S W46A R336S and D35A variants were purified on 5 ml HisTrap HP column (Cytiva, # 17524801) by elution with imidazole gradient (20-500 mM in buffer A1).

M.EcoGII W46A and E44S R336S C285S were batch purified on 3 ml Protino Ni-TED Resin (Macherey-Nagel, #745200.120) using 250 mM imidazole in buffer A for elution.

M.EcoGII fusions with protein A/G were expressed in *E.coli* T7 Express (New England Biolabs, #C2566H). The cells were grown in 500 ml TB medium supplemented with 1% glucose and 25 µg/ml kanamycin and protein expression was induced with 200 µM IPTG at 21°C overnight. The cells were lysed by sonication in a buffer A (50 mM sodium phosphate pH 7.4, 500 mM NaCl and 20 mM imidazole), the cleared lysate was applied on 4.5 ml Ni-NTA Agarose (Qiagen # 30210) resin, washed with 40 ml buffer A in a gravity flow column and eluted by overnight incubation with 30 nM recombinant bdSENP1 [16] in 15 ml of 50 mM sodium phosphate pH 7.4, 500 mM NaCl, 10% sucrose, 10 mM imidazole and 4 mM DTT. The protein was dialyzed against 3 l of 10 mM HEPES/NaOH pH 7.4, 250 mM NaCl and 1 mM DTT for 24 h at 4-8°C, concentrated first by ultrafiltration and then by dialysis against storage buffer (10 mM HEPES/NaOH pH 7.4, 250 mM NaCl, 5 mM DTT and 50% glycerol) and stored at -20°C.

Concentrations of all proteins was determined with Qubit™ Protein Assay Kit (Thermo Fisher Scientific, #Q33212). The working dilutions (typically 10 µM) were made in storage buffer supplemented with 0.2 mg/ml BSA (Sigma-Aldrich, #A7030) and were stored at -20°C.

**HPLC analysis of M.EcoGII activity**

Activity assays contained 50 mM Tris, 50 mM MOPS, pH 8.0, 0.2 mg/ml BSA 0.5 µM methyltransferase and 50 or 100 µM of cofactor. Phage λ DNA (*dam⁻, dcm⁻*) (Thermo Fisher Scientific, #SD0021) at 0.12 µg/µl concentration or *E. coli* total RNA (Thermo Fisher Scientific, #AM7940) at 0.12 µg/µl concentration were used as substrates. The reactions (40 µl) were incubated for 1 h at 37°C, then the methyltransferase was inactivated by heating for 15 min. at 65°C and degraded by incubation with 0.2 - 0.4 mg/ml proteinase K (Thermo Fisher Scientific, #EO0491) for another 1 h at 55°C. The samples were diluted to 100 µl with water, passed through MicroSpin™ Sephadex™ G-25 columns (Cytiva, #27532501) equilibrated with water and mixed with 10 µl of 10× Nucleoside Digestion Mix buffer and 1 µl of Nucleoside Digestion Mix (New England Biolabs, #M0649S). DNA was hydrolyzed at 37°C for 2 h and the resulting nucleoside mixture was analyzed on Agilent 1260 Infinity II LC system using reversed-phase UHPLC column (Supelco Titan™ C18, 1.9 µm, 7.5 cm × 2.1 mm, Sigma-Aldrich, Germany) equipped with a pre-column (Supelco Titan™ C18, 1.9 µm, 0.5 cm × 2.1 mm, Sigma-Aldrich, Germany). Compounds were eluted with methanol (5% for 2 min, followed by linear gradient to 100% in 6 min, hold 100% for 1 min, followed by linear gradient to 5% in 1 min and hold 5% for 5 min) in ammonium formate buffer (25 mM, pH 3.5) at a flow of 0.4 ml/min. and detected with absorbance DAD detector set to 254 nm (scanning 210 – 850 nm range) and single quadruple LC/MSD XT mass spectrometer scanning 100 - 1500 m/z range. An extent of adenosine modification was measured as decrease in deoxyadenosine peak area. In order to account for variable DNA recovery, deoxyguanosine peak area was used for normalization. Mass spectra served to confirm identities of modification products.

**In vitro labeling of pUC19 DNA**

1.5 µg of pUC19 DNA was incubated with 125 nM MTase and 10 µM STAR RED-Ado in 20 µl reaction buffer (50 mM Tris, 50 mM MOPS pH 8.0, 0.2 mg/ml BSA) in the presence or absence of 100 µM AdoMet for 1 h at 37°C. Enzymes were inactivated by heating for 10 min. at 80°C and DNA was purified on MSB® Spin PCRapace columns (Invitek Molecular GmbH, #1020220400) with typical recovery of 60%. 150 ng/well of labeled DNA was fractionated on agarose gel and label incorporation was quantified by measuring in-gel fluorescence in Cy5 channel.

**Maintenance and preparation of cell lines**

All cell lines were maintained in humidified 5% $CO_2$ incubator at 37°C and were split every 3-4 days or at confluence. U-2 OS cells (Sigma-Aldrich, #92022711) were cultured in McCoys 5A medium (Thermo Fisher, #16600082) with 10% FBS (BioSELL, #S0615) supplemented with 1 mM Sodium pyruvate (Sigma, #S8636) and 1% of Penicillin-Streptomycin (Sigma #P0781). Before the experiment, the cells were seeded in DMEM (Dulbecco's Modified Eagle Medium, no phenol red, Thermo Fisher, #31053044) supplemented with 10 % FBS (fetal bovine serum) and grown for 24-48 h. Human primary dermal fibroblasts (NHDF) (Lonza, #CC-2511) were cultured in high-glucose DMEM (no phenol red) with 10% FBS (Thermo Fisher, #10082147) supplemented with 1 mM Sodium pyruvate, 1% GlutaMax (Thermo Fisher, #35050038) and 1% Penicillin-Streptomycin. HeLa (ATCC, # ATCC-CCL2) cells were cultured in high-glucose DMEM (with phenol red, Thermo Fisher, #31966047) with 10% FBS supplemented with 1% Penicillin-Streptomycin.

To generate inducible cell lines, U-2 OS cells were transiently transfected with the pEBTet-vector based plasmids (pRG289, pDB009, pDB012, pDB013) using Lipofectamine 2000 (Thermo Fisher Scientific, #11668027) following manufacturer's recommendations. 48 h after transfection, selective media was applied (DMEM (Thermo Fisher Scientific, #31053028) supplemented with 10% FBS (Thermo Fisher Scientific, #10082139) and 1 µg/ml puromycin (Sigma Aldrich, #P9620)) and the cells were cultivated for 2 weeks to eliminate non-transfected cells. The selected cells were frozen in 10% DMSO and stored in liquid nitrogen. Thereafter, the cells were maintained in the selective media, and seeded in media without antibiotics 24-72 h before the experiment. Expression of transgene was induced with 0.1 µg/ml doxycycline (Sigma Aldrich, #D9891) for 24 – 48 h.

For confocal and STED imaging, cells were grown on µ-Slide 8 Well Glass Bottom chambered coverslips (Ibidi, #80827) with #1.5H glass bottom. Human primary dermal fibroblasts washed away easily by digitonin, therefore for OptiFoot labeling they were seeded on fibronectin-coated slides.

For automated wide-field microscopy, cells were grown in uncoated 12-well No. 1.0, 14 mm glass bottom plates (MatTek, #P12G-1.5-14-F). For genomic DNA isolation and preparation of samples for Western blotting, the cells were grown on 6-well plastic plates (Sarstedt, #83.3920) to 90-100% confluency.

**Western blotting**

The cells grown in 6-well plates were washed twice with room temperature PBS and lysed with 130 µl cold RIPA buffer (25 mM Tris-HCl, pH 7.6; 150 mM NaCl; 1% NP-40; 0.1% SDS) supplemented with 1× protease inhibitor cocktail. The samples were incubated on a Snijders 34528 Test Tube Rotator for 10 min at 4°C, collected into 1.5 ml microcentrifuge tubes and centrifuged for 30 min at 21130 ×g (at 4°C). Protein concentration was determined with the Qubit™ Protein Broad Range Assay Kit (Thermo Fisher Scientific, #A50669). Lysates were adjusted to 1.2 mg/ml protein by adding RIPA buffer and 20 µl of 4x Laemmli sample buffer (Bio-Rad, #1610747, supplemented with 355 mM 2-mercaptoethanol) to a final volume of 80 µl. Samples were denatured at 70°C for 10 min and stored at −20°C.

26 µg per well of protein were fractionated on 10-well Mini-PROTEAN TGX gels (Bio-Rad, #4561083) at 100 V for 1.5 h. Gels were rinsed in 20% ethanol for 10 min and proteins were transferred onto nitrocellulose

membranes using the iBlot 2 Dry Blotting System (Thermo Fisher Scientific, #IB21001) with the default P0 program (7 min). Successful transfer was confirmed by Ponceau S staining of the membrane and the membrane was washed with PBS until the stain was removed. Membrane was rinsed in 1× TBS (25 mM Tris, 150 mM NaCl, pH 7.4) and blocked for 2 h in 5% (w/v) skim milk in TBS-T (TBS supplemented with 0.05% (w/v) Tween20) and washed again for 15 min with TBS-T. Membrane was then incubated overnight at 4°C with an anti-GFP primary antibody (Abcam, #ab290; 1:2000 in 2% skim milk/TBS-T) with gentle shaking. After a brief rinse with TBS-T and three 10 min washes, membrane was incubated for 1 h at room temperature with gentle shaking with HRP-conjugated goat anti-rabbit IgG (H+L) secondary antibody (Thermo Fisher Scientific, #G-21234; 1:5000 in 2% skim milk/TBS-T). Following a brief rinse and three additional 10 min washes in TBS-T, membranes were stored in 1× TBS and imaged using Immobilon Forté Western HRP substrate (Millipore, #WBLUF0100) on an Amersham Imager 600 (Cytiva). β-actin was used as loading control. It was visualized with anti-β-actin antibody (Sigma-Aldrich, Clone AC-15, #A5441; 1:3000 dilution in 2% skim milk/TBS-T) followed by HRP-conjugated goat anti-mouse IgG (H+L) secondary antibody (Thermo Fisher Scientific, #G-31430; 1:5000 dilution in 2% skim milk/TBS-T).

**Labeling DNA by MTases expressed in U-2 OS cells**

Cell lines expressing different EcoGII MTase variants were grown to confluency and permeabilized with digitonin buffer (20 mM HEPES/KOH pH 7.5, 150 mM NaCl, 0.004% digitonin (Sigma-Aldrich, #300410), 0.5 mM spermidine HCl (Sigma-Aldrich, #85558), 1 mg/ml BSA, 1× protease inhibitor cocktail (Roche, #11873580001)) for 10 min on ice. After permeabilization, cells were washed once with cold digitonin buffer and once with cold Tween buffer (0.1% Tween-20, 20 mM HEPES/KOH pH 7.5, 150 mM NaCl, 0.5 mM spermidine HCl, 1 mg/ml BSA, 1× protease inhibitor cocktail). Cells were then incubated for 1 h at 37°C in 5% $CO_2$ with labeling buffer (50 mM Tris/MOPS pH 8.0, 50 µM spermidine HCl, 1 mg/ml BSA, 1× protease inhibitor cocktail and 3 µM 6-SiR-$PEG_4$-Ado). Following incubation, cells were washed twice with cold Tween buffer, twice with PBS (Roth, #9143.1) at room temperature, and fixed with 4% paraformaldehyde (16% PFA diluted in PBS, Thermo Fisher Scientific, #28908) in PBS for 10 min at room temperature. Fixed cells were quenched with 30 mM glycine in PBS for 5 min, washed three times with PBS, and stained with 2 µg/ml Hoechst 33342 (Sigma-Aldrich, # B2261) in PBS. Samples were stored at 4°C and imaged the following day on a spinning-disk confocal microscope.

**Antibody-targeted OptiFoot labeling**

Buffers: Wash buffer: 20 mM HEPES/KOH pH 7.5, 150 mM NaCl, 0.5 mM spermidine hydrochloride, 1 mg/ml BSA, 1× cOmplete™, EDTA-free Protease Inhibitor Cocktail (Roche, #11873580001). Digitonin buffer: wash buffer + 0.08% digitonin, high purity (Sigma, #300410). Tween-20 buffer: wash buffer + 0.1% Tween-20. Labeling buffer: 50 mM Tris, 50 mM MOPS, pH 8.0, 0.1 mg/ml BSA, 50 µM spermidine hydrochloride, 1× protease inhibitor cocktail and 3 - 30 µM cofactor analog. Typically, 3 µM cofactor was used for microscopy and 30 µM for OGM. 0.22 ml and 1.5 ml of reagents was used for 8-well chambered coverslips and 6-well plates, respectively.

To permeabilize cells, the media was replaced with digitonin buffer and the plate was incubated on ice for 10 min. Then, the samples were briefly washed once with digitonin buffer and once with Tween-20 Wash buffer. Appropriate antibody dilution in Tween-20 buffer was added and the samples were incubated for 2 h on ice. Following three washes with Tween-20 buffer, the incubation was continued with 10 nM euMTase-$L_{14}$-pA/G in Tween-20 buffer for another 2 h on ice. Then the samples were washed again three times with Tween-20 buffer and incubated with Labeling buffer for 1 h at 37°C. After that, the samples were washed twice with Tween-20 buffer and twice with PBS. For imaging experiments, the cells were fixed with 4% PFA for 10 min. at room temperature, followed by 5 min incubation with 30 mM glycine in PBS. After final washes with PBS, the samples were stored at 4°C in PBS with 0.1 µg/ml Hoechst 33342.

To perform staining via biotin-streptavidin interaction, endogenous biotin was blocked by supplementing antibody solution (polyclonal $\alpha$Lamin B1, 1:1000) with 0.1 mg/ml streptavidin (Applichem, A1495.0005). Excessive streptavidin was quenched by adding 50 µg/ml D-biotin during the incubation with 10 nM euMTase-$L_{14}$-pA/G, followed by incubation with 10 µM biotin-$PEG_4$-Ado. The cells were fixed with PFA and incubated with 2 µg/ml AlexaFluor™ 568-labeled streptavidin (Invitrogen, S11226) for 1h at room temperature followed by 3 washes with PBS and Hoechst 33342 staining.

To determine, which nucleic acid is labeled in OptiFoot experiment, U-2 OS cells were fixed with methanol + 1 mM EGTA for 5 min at -20°C, washed 4 times with PBS and pretreated with 0.2 mg/ml RNase A (New England Biolabs, #T3018L) or 330 u/ml DNase I (Thermo Fisher Scientific, #18047019) for 2h at 37°C or both in PBS containing 1% BSA. DNase I sample was supplemented with 2.5 mM $MgCl_2$ and 0.5 mM $CaCl_2$. Followed by washes with PBS, the samples were incubated with antibodies against lamin B1 or nucleolin (1:1000) at 4°C overnight followed by 10 nM euMTase-$L_{14}$-pA/G for 3h at room temperature. After washing away unbound MTase, the samples were incubated with 3 µM 6-SiR-$PEG_4$-Ado for 1 h at 37°C in MTase buffer containing 0.2 mg/ml BSA, then decorated with secondary antibody and stained with Hoechst 33342.

**Genomic DNA isolation**

Genomic DNA was isolated with Monarch® HMW DNA Extraction Kit for Cells & Blood (New England Biolabs, #T3050L) following manufacturer's recommendations. To obtain ultra-high molecular weight (UHMW) DNA for optical genome mapping, cell lysis was carried out with 600 rpm at initial lysis step. This yielded highly viscous an inhomogeneous gDNA sample that was left to completely dissolve for ~1 week at 4°C, with occasional mixing by pipetting with wide-bore tips. Presence of UHMW DNA was verified by pulsed-field electrophoresis in 0.75 % agarose gel (SeaKem® Gold Agarose, Lonza # 50150) with Pippin Pulse power supply (Sage Science) using pre-set protocol "5-430kb". Running buffer was 0.5×KBB buffer (51.18 mM Tris (base), 28.81 mM TAPS (free acid), 0.082 mM EDTA (free acid)). The sample was deemed suitable for OGM if molecules larger than 150 kbp were detectable.

For 6mA detection, the same kit was used with 2000 rpm at the initial lysis step. This procedure yielded lower molecular weight DNA that was easier to process due to lower viscosity. gDNA was isolated from cells induced for 48h. To detect 6mA formation, 1 µg of gDNA was incubated in 15 µl of recommended buffer with 3.5 u of R.DpnI (Thermo Fisher Scientific, #ER1705) or R.MboI (Thermo Fisher Scientific, #ER0811) for 1h at 37°C. R.DpnI cleaves DNA only if its target sequence contains N6-methyladenine. In contrast, R.MboI activity is blocked by 6mA in its target sequence.

gDNA was quantified with Qubit™ dsDNA Quantitation kit, Broad Range (Thermo Fisher Scientific, # Q32853).

**Protein-genome interaction profiling with optical genome mapping (OGM)**
Reference labeling of CTTAAG sites with green fluorophore was performed with Bionano Prep DLS-G2 Labeling Kit (Bionano Genomics, #80046) following manufacturer's protocol.

300-1000 Gb of data was acquired on Saphyr instrument and all raw images of molecules were saved on disk. Analyzed samples are listed in **Table S2**.

Data processing and analysis workflow is outlined in **Scheme S2**. In the first step, OptiFoot signal was extracted as continuous fluorescence profile and aligned to hg38 reference genome using previously described pipeline with modifications (github.com/ebensteinLab/DaFCA) [17]. The modifications were aimed at increasing computational speed and streamlining the processing. The complete modified pipeline for this step is outlined in **Scheme S3** and full code is available on GitHub. Through multiple steps, this pipeline yields one BEDgraph file per chromosome (chr[i]_raw.BEDgraph), containing aggregated signal from all the molecules that align to the same genome coordinates. The analysis continued as indicated in **Scheme S3**, using BEDTools (2.31.1) [18], AWK (5.2.1), deepTools (3.5.6) [19] and custom python scripts. All scripts used in this study are available on GitHub (github.com/LabelingAndImaging/OptiFoot).

Single chromosome profiles were binned to non-overlapping 1kb windows to reflect physical resolution limit if OGM (~500 bp/px) using BEDTools *map*, individual chromosomes were merged into a single file to yield one file per dataset with *08_merge_chromosomes.py* and the corresponding background control (profile generated with non-specific antibody rIgG) was subtracted using combination of BEDTools *unionbedg* and AWK. To minimize artefacts of inconsistent coverage between sample and control, the positions where either sample or control is equal to zero were filtered-out. The background-corrected BEDgraph file was sorted lexicographically and converted to a binary bigWig format with *bedGraphToBigWig* utility from the UCSC Genome Browser toolkit. Enrichment of OptiFoot signal at specific genome features (**Supplementary Table 3**) was calculated with deepTools *computeMatrix* and plotted with deepTools *plotHeatmap* functions. To control for specificity, OptiFoot signal enrichment in randomized genomic features was computed and plotted. Shuffled controls were computed with BEDTools *shuffle*. The randomized features were placed on the same chromosome and the regions absent in rIgG-03 sample were excluded, as a close approximation for regions inaccessible to OGM.

To compare OptiFoot and MadID datasets, the OptiFoot profile produced with polyclonal anti-lamin B1 antibody was binned to a matching 100kb resolution, both datasets were aligned using BEDTools *unionbedg* and correlation was computed with *compute_correlatation.py*, employing pandas library's .corr() method. Bins with very high (>150 a.u.), negative and zero values were excluded from calculation. Zero values report bins that lack coverage in either of the samples. High and negative values in OGM experiment appear mostly at the boundaries of repetitive regions that lack reference labels and thus have low and inconsistent coverage. <2% of bins were affected and without affecting reported correlation coefficients.

**Microscopy**
Confocal imaging was performed on Visitron Spinning disk/TIRF/SMLM system (Visitron Systems GmbH, Germany) using Nikon CFI Apo Lambda 60× Oil NA 1.4 objective and Prime BSI sCMOS camera (Teledyne Photometrics) with pixel size of 6.5×6.5 µm, which corresponds to 111×111 nm pixel size on images. Excitation was provided by lasers at 405 nm, 488 nm, 561 nm, and 640 nm, each at 150 mW. The spinning

disk pinhole size was 50 µm. Fluorescence emission was separated using a universal dichroic mirror (405/488/561/640) and detected through the following emission filters: ET460/50m, ET525/50m, ET609/54m, and 665LP. Images were acquired as 61× 200 nm z-stacks.

STED images in **Fig.2b** were acquired on Abberior STED MIRAVA (Abberior Instruments GmbH) equipped with 405, 488, 518, 561, 640 and 700 nm 40 MHz pulsed excitation lasers, a pulsed 775 nm 40 MHz STED laser, an UPlanSApo 60x/1.42 Oil objective (Olympus), three APDs and one MATRIX detector. SiR was excited with 640 nm laser at 10% power and was detected in 650 – 760 nm window; STED laser was 5%. Abberior STAR 580 was excited with 561 nm laser at 20% and emission detected at 571 - 630 nm window; STED laser was 40%. Pixel size was 25 nm, pinhole size - 0.5 AU, line accumulation was 3, pixel dwell time was 20 µs.

STED images in **Fig.3c**, **4a** and **5b** were acquired on Abberior STED 775 QUAD scanning microscope (Abberior Instruments GmbH) equipped with 488 nm, 561 nm and 640 nm 40 MHz pulsed excitation lasers, a pulsed 775 nm 40 MHz STED laser, and an UPlanSApo 100×/1.40 Oil objective. SiR was excited with 640 nm laser at 10-20% power and was detected in 685/70 nm window; STED laser was 40%. Abberior STAR 580 was excited with 561 nm laser at 70-100% and emission detected at 615/20 nm; STED laser was 75%. Pixel size was 20 nm, pinhole size - 0.45 AU, line accumulation was 2, pixel dwell time was 5 µs.

Wide field imaging was performed on Lionheart FX Automated Microscope (BioTek) with Olympus dry 20× NA 0.45 objective, using laser autofocus. 16 fields of view in 3 planes z-stack, spanning 6 µm in thickness, were acquired per well. Images were stitched and focus-stacked with in-built Gene 5 software (BioTek). The final images encompassed 1380 × 950 µm field of view.

**Image analysis**

Basic image processing (z-projections, cropping, brightness adjustment, adding scale bars and choosing z-slice) was done with Fiji[20]. Image segmentation and quantification pipelines were implemented in CellProfiler v4.2.8 [21]. RunCellpose v2 or v3 [22] was run as a CellProfiler plugin in a containerized environment using Docker Desktop v4.49.0.

To count GFP-expressing cells in wide-field fluorescent images, nuclei segmentation of Hoechst 33342 and GFP channels was performed with *RunCellpose* module in nuclei detection mode with expected object diameter of 60, cell probability threshold of 0.0, minimum size of 25 and flow threshold of 0.4. GFP-positive nuclei were identified and counted with *RelateObjects* module.

To avoid bias in selecting a representative z-slice, confocal z-stacks were converted to two-dimensional images using sum-intensity z-projection and saved as 16-bit tiff files.

*Anti-Lamin B1-directed OptiFoot staining*. GFP channel (anti-rabbit-AlexaFluor488 antibody) was smoothed with Median filter with automatically detected artifact diameter and nuclei were identified using *RunCellpose* module in nuclei detection mode with expected object diameter of 200, cell probability threshold of 0.0, minimum size of 150 and flow threshold of 0.5. OptiFoot signal (mean intensity) was measured inside the nuclei in SiR channel (6-SiR-PEG$_4$-Ado cofactor). Untargeted background signal was measured in a 20-pixel-wide region around each identified nucleus (mean intensity) and subtracted from the corresponding nucleus signal.

*Anti-Nucleolin-directed OptiFoot staining*. Nuclei and nucleoli segmentation was performed on GFP channel (anti-rabbit-AlexaFluor488 antibody). First, GFP channel was smoothed using Median filter with automatically detected artifact diameter and nuclei were identified using *RunCellpose* module in nuclei detection mode with expected object diameter of 200, cell probability threshold of 0.0, minimum size of 150 and flow threshold of 0.5. Nucleoli signal was enhanced in *EnhanceOrSupressFeatures* module, using enhance operation, feature type – speckles, feature size – 30, speed and accuracy – fast. The resulting images were smoothed with Median filter with typical artifact diameter of 5. Nucleoli were identified in *IdentifyPrimaryObjects* module with typical object diameter of 10-100, using adaptive Otsu two classes thresholding, with threshold smoothing scale of 1.3488, threshold correction factor of 0.7, size of adaptive window of 20 and method to distinguish clumped objects – none. Nucleoli were related to nuclei in *RelateObjects* module. Background was measured in nucleus area excluding nucleoli (segmented with *IdetifyTertiaryObjects*). OptiFoot signal (mean intensity) was measured in SiR channel within nucleoli and background signal (mean intensity) of the corresponding nucleus was subtracted.

*Labelling by in vivo expressed MTases.* Nuclei were identified in Hoechst 33342 channel using *RunCellpose* module in nuclei detection mode with object diameter of 300, cell probability threshold of 0.0, minimum size of 30 and flow threshold of 0.4. To approximate the perinuclear region, segmented nuclei were expanded using *DilateObjects* module with disk-shaped structuring elements, size 30. Background regions were defined using the *IdentifyTertiaryObjects* module, where larger objects corresponded to the dilated nuclei and smaller objects to the original nuclei identified by *RunCellpose*. MTase signal (mean intensity) was measured in GFP channel, and labeled DNA signal (mean intensity) was measured in SiR channel (6-SiR-$PEG_4$-Ado cofactor), both measured within the nuclear regions. To obtain background corrected values for each nucleus the mean background intensity of the tertiary objects was subtracted from corresponding nuclei mean intensity.

**Supplementary sequences**

- $H_6$-M.**EcoGII** (plasmids pRG202, pRG203, pRG224, pRG225, pRG227, pRG229)

MGSSHHHHHHSSGLVPRGSHMLNTVKISSCELINADCLEFIRSLPENSVDLIVTDPPYFKVKP**E**G**W**DNQWKGDDDYLK WLDQCLAQFWRVLKPAGSLYLFCGHRLASDIEIMMRERFSVLNHIIWAKPSGRWNGCNKESLRAYFPATERILFAEHYQ GPYRPKDAGYEAKGRALKQHVMAPLIAYFRDARAALGITAKQIADATGKKNMVPHWFSASQWQLPNESDYLKLQSLF ARVAEEKHQRGELEKPHHQLVSTYSELNRKYMELLSEYKNLRRYFGVTVQVPYTDVWTYKPVQYYPGKHP**C**EKPAEML QQIISASSRPGDLVADFFMGSGSTVKAAMALGRRAIGVELETG**R**FEQTVREVQDLIV

His-tag and thrombin site are underlined. M.EcoGII wt sequence is shown in blue, mutagenized amino acids (E44, W46, S285 and R336) are in bold and underlined. Numbering ignores parts coming from vector and refers to original M.EcoGII sequence, starting from the first methionine.

- $H_{14}$-bdSUMO-**gly-pA/G-linker$_6$-M.EcoGII W46A** (plasmid pRG274)

MQLSMSKHHHHSGHHHTGHHHHSGSHHHTGSSSAAGGEEDKKPAGGEGGGAHINLKVKGQDGNEVFFRIKRSTQLK KLMNAYCDRQSVDMTAIAFLFDGRRLRAEQTPDELEMEDGDEIDAMLHQTGG🠗GTMITPSLKDDPSQSANLLSEAKK LNESQAPKADNKFNKEQQNAFYEILHLPNLNEEQRNGFIQSLKDDPSQSANLLAEAKKLNDAQAPKADNKFNKEQQNA FYEILHLPNLTEEQRNGFIQSLKDDPSVSKEILAEAKKLNDAQAPKTTYKLVINGKTLKGETTTEAVDAETAERHFKQYAND NGVDGEWTYDDATKTFTVTEKPEVIDASELTPAV*DDDKEF*LNTVKISSCELINADCLEFIRSLPENSVDLIVTDPPYFKVKPE G**A**DNQWKGDDDYLKWLDQCLAQFWRVLKPAGSLYLFCGHRLASDIEIMMRERFSVLNHIIWAKPSGRWNGCNKESL RAYFPATERILFAEHYQGPYRPKDAGYEAKGRALKQHVMAPLIAYFRDARAALGITAKQIADATGKKNMVPHWFSASQ WQLPNESDYLKLQSLFARVAEEKHQRGELEKPHHQLVSTYSELNRKYMELLSEYKNLRRYFGVTVQVPYTDVWTYKPVQ YYPGKHPCEKPAEMLQQIISASSRPGDLVADFFMGSGSTVKAAMALGRRAIGVELETGRFEQTVREVQDLIV

His-tag is underlined. Protein A/G is gold, M.EcoGII W46A sequence is blue, W46A mutation is in bold and underlined, linker is shown in italic, the black arrow denotes bdSENP1 cleavage site.

- $H_{14}$-bdSUMO-**gly-pA/G-linker$_6$-M.EcoGII W46A** (plasmid pRG280)

MSKHHHHSGHHHTGHHHHSGSHHHTGSSSAAGGEEDKKPAGGEGGGAHINLKVKGQDGNEVFFRIKRSTQLKKLMN AYCDRQSVDMTAIAFLFDGRRLRAEQTPDELEMEDGDEIDAMLHQTGG🠗GLNTVKISSCELINADCLEFIRSLPENSVDL IVTDPPYFKVKPEG**A**DNQWKGDDDYLKWLDQCLAQFWRVLKPAGSLYLFCGHRLASDIEIMMRERFSVLNHIIWAKPS GRWNGCNKESLRAYFPATERILFAEHYQGPYRPKDAGYEAKGRALKQHVMAPLIAYFRDARAALGITAKQIADATGKKN MVPHWFSASQWQLPNESDYLKLQSLFARVAEEKHQRGELEKPHHQLVSTYSELNRKYMELLSEYKNLRRYFGVTVQVP YTDVWTYKPVQYYPGKHPCEKPAEMLQQIISASSRPGDLVADFFMGSGSTVKAAMALGRRAIGVELETGRFEQTVREV QDLIV*GSGGSGESGTMITP*SLKDDPSQSANLLSEAKKLNESQAPKADNKFNKEQQNAFYEILHLPNLNEEQRNGFIQSLK DDPSQSANLLAEAKKLNDAQAPKADNKFNKEQQNAFYEILHLPNLTEEQRNGFIQSLKDDPSVSKEILAEAKKLNDAQA PKTTYKLVINGKTLKGETTTEAVDAETAERHFKQYANDNGVDGEWTYDDATKTFTVTEKPEVIDASELTPAV

His-tag is underlined. Protein A/G is gold, M.EcoGII W46A sequence is blue, W46A mutation is in bold and underlined, linker is shown in italic, the black arrow denotes bdSENP1 cleavage site.

- **DD-$H_6$-M.EcoGII W46A-hLamin B1** (plasmid pRG289)

MGVQVETISPGDGRTFPKRGQTCVVHYTGMLEDGKKVDSSRDRNKPFKFMLGKQEVIRGWEEGVAQMSVGQRAKLT ISPDYAYGATGHPGIIPPHATLVFDVELLKPEASATGLRSSHTASATHHHHHHSSGLNTVKISSCELINADCLEFIRSLPENS VDLIVTDPPYFKVKPEG**A**DNQWKGDDDYLKWLDQCLAQFWRVLKPAGSLYLFCGHRLASDIEIMMRERFSVLNHIIWA

KPSGRWNGCNKESLRAYFPATERILFAEHYQGPYRPKDAGYEAKGRALKQHVMAPLIAYFRDARAALGITAKQIADATG
KKNMVPHWFSASQWQLPNESDYLKLQSLFARVAEEKHQRGELEKPHHQLVSTYSELNRKYMELLSEYKNLRRYFGVTV
QVPYTDVWTYKPVQYYPGKHPCEKPAEMLQQIISASSRPGDLVADFFMGSGSTVKAAMALGRRAIGVELETGRFEQTV
REVQDLIV*GSGGSGESGTFELEVLFQGPGTAITSLYKKAGL*MATATPVPPRMGSRAGGPTTPLSPTRLSRLQEKEELRELN
DRLAVYIDKVRSLETENSALQLQVTEREEVRGRELTGLKALYETELADARRALDDTARERAKLQIELGKCKAEHDQLLLNY
AKKESDLNGAQIKLREYEAALNSKDAALATALGDKKSLEGDLEDLKDQIAQLEASLAAAKKQLADETLLKVDLENRCQSLT
EDLEFRKSMYEEEINETRRKHETRLVEVDSGRQIEYEYKLAQALHEMREQHDAQVRLYKEELEQTYHAKLENARLSSEMN
TSTVNSAREELMESRMRIESLSSQLSNLQKESRACLERIQELEDLLAKEKDNSRRMLTDKEREMAEIRDQMQQQLNDYE
QLLDVKLALDMEISAYRKLLQGEEERLKLSPSPSSRVTVSRASSSRSVRTTRGKRKRVDVEESEASSSVSISHSASATGNVCI
EEIDVDGKFIRLKNTSEQDQPMGGWEMIRKIGDTSVSYKYTSRYVLKAGQTVTIWAANAGVTASPPTDLIWKNQNSW
GTGEDVKVILKNSQGEEVAQRSTVFKTTIPEEEEEEEEAAGVVVEEELFHQQGTPRASNRSCAIM

His-tag and precision protease cleavage site are underlined. Proteotuner™ destabilizing domain (DD) is in red, M.EcoGII W46A sequence is blue, W46A mutation is in bold and underlined, linker is shown in italic, hLamin B1 sequence is purple.

- **$H_6$-M.EcoGII-eGFP-NLS** (plasmids pDB009, pDB012 and pDB013)

MGSSHHHHHHSSGLVPRGSHMLNTVKISSCELINADCLEFIRSLPENSVDLIVT**D**PPYFKVKPEG**W**DNQWKGDDDYLK
WLDQCLAQFWRVLKPAGSLYLFCGHRLASDIEIMMRERFSVLNHIIWAKPSGRWNGCNKESLRAYFPATERILFAEHYQ
GPYRPKDAGYEAKGRALKQHVMAPLIAYFRDARAALGITAKQIADATGKKNMVPHWFSASQWQLPNESDYLKLQSLF
ARVAEEKHQRGELEKPHHQLVSTYSELNRKYMELLSEYKNLRRYFGVTVQVPYTDVWTYKPVQYYPGKHPCEKPAEML
QQIISASSRPGDLVADFFMGSGSTVKAAMALGRRAIGVELETGRFEQTVREVQDLIVGSTMVSKGEELFTGVVPILVELD
GDVNGHKFSVSGEGEGDATYGKLTLKFICTTGKLPVPWPTLVTTLTYGVQCFSRYPDHMKQHDFFKSAMPEGYVQERTI
FFKDDGNYKTRAEVKFEGDTLVNRIELKGIDFKEDGNILGHKLEYNYNSHNVYIMADKQKNGIKVNFKIRHNIEDGSVQL
ADHYQQNTPIGDGPVLLPDNHYLSTQSALSKDPNEKRDHMVLLEFVTAAGITLGMDELYKGSPKKKRKVLE

His-tag and SV40 NLS are underlined. M.EcoGII wt sequence is blue, mutagenized D35 and W46 amino acids are in bold and underlined.

### Supplemenatry Notes.

#### Chemical synthesis and characterization of compounds

*General analytical methods*

<u>NMR spectra</u> were recorded at 25°C with an Agilent 400-MR spectrometer at 400.06 MHz ($^1H$) and 100.60 MHz ($^{13}C$) and are reported in ppm. Chemical shifts (δ) are reported in ppm relative to tetramethylsilane (TMS, δ = 0 ppm) and were referenced using the residual solvent signals according to literature values[23]. Multiplicities of signals are described as follows: s = singlet, d = doublet, t = triplet, q = quartet, p = pentet, m = multiplet or overlap of non-equivalent resonances; br = broad signal. Coupling constants (*J*) are given in Hz. All NMR spectra were processed with MestRenova 11.0.4 software.

<u>ESI-MS</u> were recorded on a Varian 500-MS spectrometer (Agilent). <u>ESI-HRMS</u> were recorded on a MICROTOF spectrometer (Bruker) equipped with ESI ion source (Apollo) and direct injector with LC autosampler Agilent RR 1200.

<u>Analytical LC–MS</u> analysis was performed on an Agilent 1260 Infinity II LC/MS system equipped with a binary pump (G7112B), an autosampler (G7129A), temperature-controlled column compartment (G7116A), diode array detector WR (G7115A) and 6100 series quadruple LC/MSD XT (G6135B) with API electrospray. Analysis was performed using either an Ascentis® Express AQ-C18 column (2 µm, 5 cm × 2.1 mm; Sigma cat. no. 567200-U) or a Titan C18 column (1.9 µm, 7.5 cm × 2.1 mm; Sigma cat. no. 557123-U). The mobile phase consisted of A: 25 mM aqueous ammonium formate buffer (pH 3.5) and B: MeOH.

<u>Preparative HPLC</u> was performed on a combined Agilent1260/1290 Infinity II preparative system equipped with 1290 Infinity II open-bed sampler (G7169B)/fraction collector (G7159B), 1290 Infinity II preparative binary pump (G7161A), 1260 Infinity II multiple wavelength detector (G7165A) and with Agilent Pursuit 10 $C_{18}$, 10 µm, 250 × 50 mm preparative column (Agilent, cat. no. A3002250X500).

*Synthesis of cofactors with simple transferable groups*

$N_3$-Ado

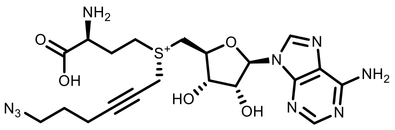

inactive stereoisomer (peak 1)

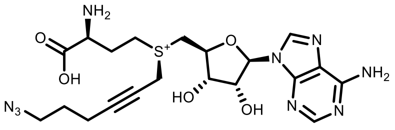

$N_3$-Ado (active, peak 2)

$N_3$-Ado was synthesized as described previously [10]. AdoHcy (100 mg, 0.26 mmol, Sigma Aldrich Cat. Nr. A9384) was dissolved in a 1:1 mixture of formic acid and acetic acid (10 ml) and 6-azidohex-2-yn-1-yl 4-nitrobenzenesulfonate (5 eq., 1.3 mmol) was added at room temperature and was stirred for 24h. The reaction was quenched by adding water (40 ml) and the aqueous phase was washed with diethyl ether (3 × 50 ml). The aqueous phase was concentrated under reduced pressure and redissolved in ammonium formate buffer (20 ml, 10 mM, pH 3.6). The sample was passed through a Dowex 1 column (pre-equilibrated with 10 mM ammonium formate, pH 3.6) to remove 4-nitrobenzensulfonic acid formed during the reaction and the target compound was purified by preparative HPLC (preparative column: Pursuit 10 C18 10 µm, 250 x 50 mm, Agilent, flow rate: 150 ml/min, solvent A: MeOH, solvent B: 10 mM ammonium formate buffer pH 3.5; room temperature, gradient A:B - 2 min 2:98 isocratic, 2-20 min 2:98 to 20:80 gradient and 20-22min 20:80 isocratic). Collected fractions of pure stereoisomers were concentrated and the concentrations of the obtained solutions were determined by their UV absorption ($\varepsilon_{260}$ = 15400 $L{\cdot}mol^{-1}{\cdot}cm^{-1}$). 10ml, 7.9 mM of peak-1 (inactive) in 30% yield, and 10 ml 6.7 mM of peak-2 (active) in 26% yield were obtained. The following analytical data were collected for cofactor and it conforms with previously published results [10]:

$^1H$ NMR (400 MHz, $D_2O$): 1.42-1.56 (m, 2H), 2.0-2.12 (m, 2H), 2.12-2.24 (m, 2H), 3.15 (t, J = 6.3 Hz, 2H), 3.3-3.48 (m, 2H), 3.5-3.8 (m, H, 3H), 4.18-4.21 (m, 2H), 4.41-4.49 (m, 1H), 4.86 (dd, J = 5.7, 5.1 Hz, 1H), 6.01 (d, J = 5.7 Hz, 1H), 8.17 (s, 1H), 8.18 (s, 1H), the signal of C3'-H in the sugar ring overlaps with residual water signal in $D_2O$.

HRMS: m/z $[M]^+$ calculated for $C_{20}H_{28}N_9O_5S$: 506.1934; found: 506.1929

Hexynyl-Ado

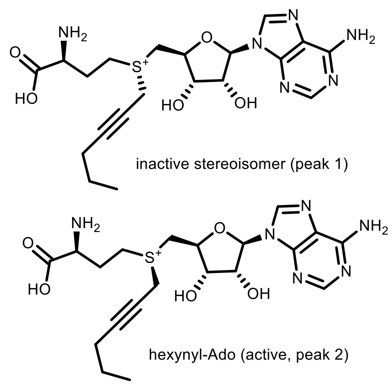
inactive stereoisomer (peak 1)
hexynyl-Ado (active, peak 2)

AdoHcy (10 mg, 0.026 mmol, Sigma Aldrich Cat. Nr. A9384) was dissolved in a (S)-2-chloropropanoic acid (200 µl) and Hex-2-yn-1-yl methanesulfonate (60 µl, 0.34 mmol) was added to the solution. The obtained mixture was stirred at 40°C temperature for 24h, the course of the reaction was monitored by LC/MS. Then a mixture was poured to ammonium formate buffer (25 ml, 10mM) and washed with $Et_2O$ (3x25 ml). The aqueous layer was taken and concentrated to 1-3 ml and the target compound was purified by preparative HPLC: Agilent Pursuit 10 C18 10 µm, 250 x 50 mm (Agilent, cat. No. A3002250X500) preparative column, flow rate: 150 ml/min, solvent A: MeOH, solvent B: 10 mM ammonium formate buffer pH 3.5 at room temperature, gradient A:B - 2 min 2:98 isocratic, 2-20 min 2:98 to 20:80 gradient and 20-22min 20:80 isocratic). Collected fractions of pure stereoisomers were concentrated and concentrations of the obtained solutions were determined by their UV absorption ($\varepsilon_{260}$ = 15400 $L \cdot mol^{-1} \cdot cm^{-1}$). Obtained 0.8 ml, 5.8 mM of peak-1 (inactive) in 18% yield, and 0.8 ml 5.2 mM of peak-2 (active) in 16% yield. The following analytical data collected for cofactor hexynyl-Ado and it conforms with previously published results [24]:

$^1H$ NMR (400 MHz, Deuterium Oxide) δ 8.29 (s, 1H), 8.28 (s, 1H), 6.10 (d, J = 3.9 Hz, 1H), 4.97 (dd, J = 5.5, 3.9 Hz, 1H), 4.68 (d, J = 5.9 Hz, 1H), 4.59 – 4.53 (m, 1H), 4.35 (t, J = 2.3 Hz, 2H), 4.03 – 3.89 (m, 2H), 3.80 (t, J = 6.5 Hz, 1H), 3.65 (dt, J = 13.2, 7.8 Hz, 1H), 3.51 (dt, J = 13.5, 7.6 Hz, 1H), 2.39 – 2.29 (m, 2H), 2.23 (tt, J = 6.5, 2.1 Hz, 2H), 1.52 – 1.42 (m, 2H), 0.89 (t, J = 7.4 Hz, 3H).

HRMS (ESI) calcd. for $C_{20}H_{29}N_6O_5S$ $[M]^+$ 465.1915, found 465.1924.

Octadiynyl-Ado

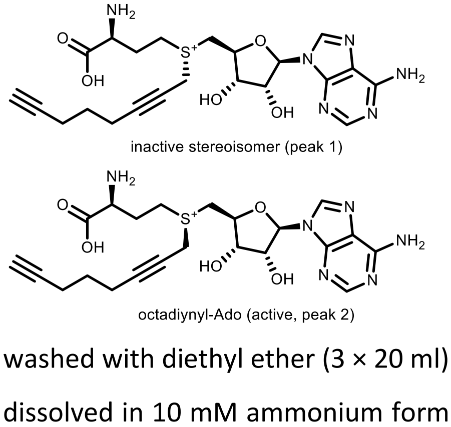
inactive stereoisomer (peak 1)
octadiynyl-Ado (active, peak 2)

Octadiynyl-Ado was synthesized as described previously [10]. In brief, S-Adenosyl-L-homocysteine (50 mg, 0.13 mmol, Sigma Aldrich Cat. Nr. A9384) was dissolved in a 1:1 mixture of formic acid and acetic acid (5 ml) and octa-2,7-diyn-1-yl 4-nitrobenzenesulfonate (4 eq., 160 mg, 0.52 mmol) was added at room temperature and was stirred for 24h.The reaction was quenched by adding water (20 ml) and the aqueous phase was washed with diethyl ether (3 × 20 ml). The aqueous phase was concentrated under reduced pressure and dissolved in 10 mM ammonium formate buffer (5 ml, pH 3.6). The sample was passed through Dowex 1 column (pre-equilibrated with 10 mM ammonium formate, pH 3.6) to remove 4-nitrobenzensulfonic acid and the target compound was purified by the preparative HPLC: Agilent Pursuit 10 C18 10 µm, 250 x 50 mm (Agilent, cat. No. A3002250X500) preparative column, flow rate: 150 ml/min, solvent A: 10mM Amonium formate pH 3.5, solvent B: MeOH; room temperature, gradient A:B - 2 min 98:2 isocratic, 2-20 min 98:2 to 0:100 gradient and 20-22 min 0:100 isocratic. Collected fractions of pure stereoisomers were concentrated and concentrations of the obtained solutions were determined by their UV absorption ($\varepsilon_{260}$ = 15400 $L\cdot mol^{-1}\cdot cm^{-1}$). Obtained peak-1 containing 10.9 mM cofactor solution in 22% yield, and peak-2 containing 10.2 mM cofactor in 20% yield. The peak-2 with longer retention time was determined to be the enzymatically active stereoisomer. The following analytical data collected for cofactor octadiynyl-Ado and it conforms with previously published results [10]:

HRMS: m/z $[M]^+$ calcd for $C_{22}H_{29}N_6O_5S^+$: 489.1915; found 489.1917

*Synthesis of dyes with reactive linker*

MeSiR-PEG$_4$-alkyne

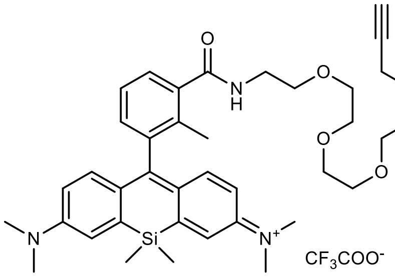

Reaction was performed following general synthesis procedure A from 4-MeSiR-COOH[25] (25mg) and $NH_2$-PEG$_4$-alkyne using HATU as peptide coupling reagent. The compound was purified by preparative HPLC (preparative column: Pursuit 10 C18 10 µm, 250 x 50 mm, Agilent, flow rate: 150 ml/min, solvent A: MeOH, solvent B: $H_2O$ + 0.1% TFA; room temperature, gradient A:B - 2 min 30:70 isocratic, 2-20 min 30:70 to 100:0 gradient and 20-22 min 100:0 isocratic). Obtained 30 mg (70% yield) of a dark blue solid.

$^1$H NMR (400 MHz, Methanol-d4) δ 7.55 (d, J = 6.9 Hz, 1H, Ar-H), 7.45 (t, J = 7.6 Hz, 1H, Ar-H), 7.37 (d, J = 2.8 Hz, 2H, Xan-H), 7.21 (d, J = 7.2 Hz, 1H, Ar-H), 7.11 (d, J = 9.6 Hz, 2H, Xan-H), 6.77 (dd, J = 9.6, 2.8 Hz, 2H, Xan-H), 4.15 (d, J = 2.4 Hz, 2H, Alkyne-$CH_2$), 3.70 – 3.57 (m, 16H, $CH_2$ PEG), 3.36 (s, 12H, $N(CH_3)_2$), 2.84 (t, J = 2.3 Hz, 1H, Alkyne-H), 2.07 (s, 3H, -$CH_3$), 0.62 (s, 3H, Si-$CH_3$), 0.61 (s, 3H, Si-$CH_3$). NH proton not observed due to exchange with $CD_3OD$ solvent.

ESI-MS, positive mode: m/z = 656.4 $[M]^+$.

HRMS (ESI) calculated for $C_{38}H_{50}N_3O_5Si$ $[M]^+$ 656.3539, found 656.3514.

Copy of HRMS:

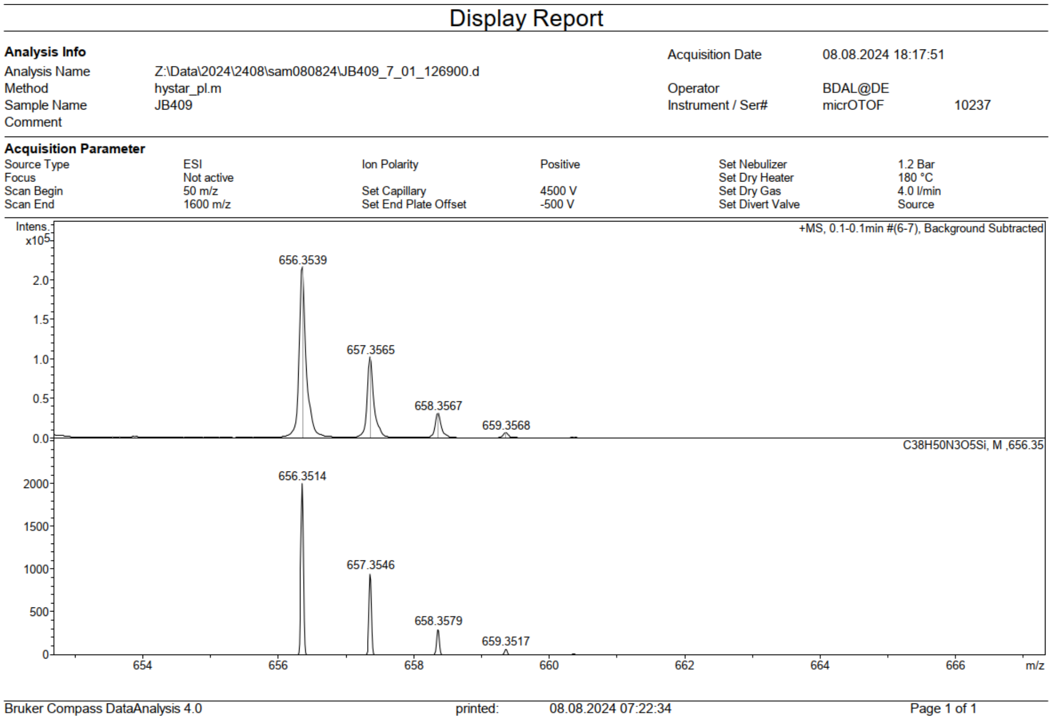

6-SiR-PEG$_4$-alkyne

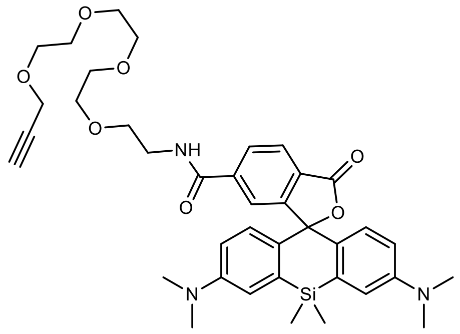

Reaction was performed following general synthesis procedure A from 6-SiR-COOH[26] (10 mg) and $NH_2$-PEG$_4$-alkyne using HATU as peptide coupling reagent. The compound was purified by preparative HPLC (preparative column: Pursuit 10 C18 10 µm, 250 x 50 mm, Agilent, flow rate: 150 ml/min, solvent A: MeOH, solvent B: $H_2O$ + 0.1% TFA; room temperature, gradient A:B - 2 min 30:70 isocratic, 2-20 min 30:70 to 100:0 gradient and 20-22 min 100:0 isocratic). Obtained 9.5 mg (65% yield) of a light blue solid.

$^1$H NMR (400 MHz, Methanol-$d_4$) δ 8.29 (d, *J* = 8.2 Hz, 1H, Ar-H), 8.13 (dd, *J* = 8.2, 1.8 Hz, 1H, Ar-H), 7.73 (dd, *J* = 1.8, 0.6 Hz, 1H, Ar-H), 7.32 (d, *J* = 2.8 Hz, 2H, Xan-H), 6.97 (d, *J* = 9.5 Hz, 2H, Xan-H), 6.75 (dd, *J* = 9.5, 2.9 Hz, 2H, Xan-H), 4.12 (d, *J* = 2.3 Hz, 2H, Alkyne-$CH_2$), 3.65 – 3.51 (m, 16H, $CH_2$ PEG), 3.28 (s, 12H, -$N(CH_3)_2$), 2.82 (t, *J* = 2.4 Hz, 1H, Alkyne-H), 0.64 (s, 3H, -Si-$CH_3$), 0.59 (s, 3H, -Si-$CH_3$). NH proton not observed due to exchange with $CD_3OD$ solvent.

ESI-MS, positive mode: m/z = 708.3 $[M+Na]^+$.

HRMS (ESI) calculated for $C_{38}H_{47}N_3O_7Si$ $[M+Na]^+$ 708.3075 found 708.3079.

Copy of HRMS:

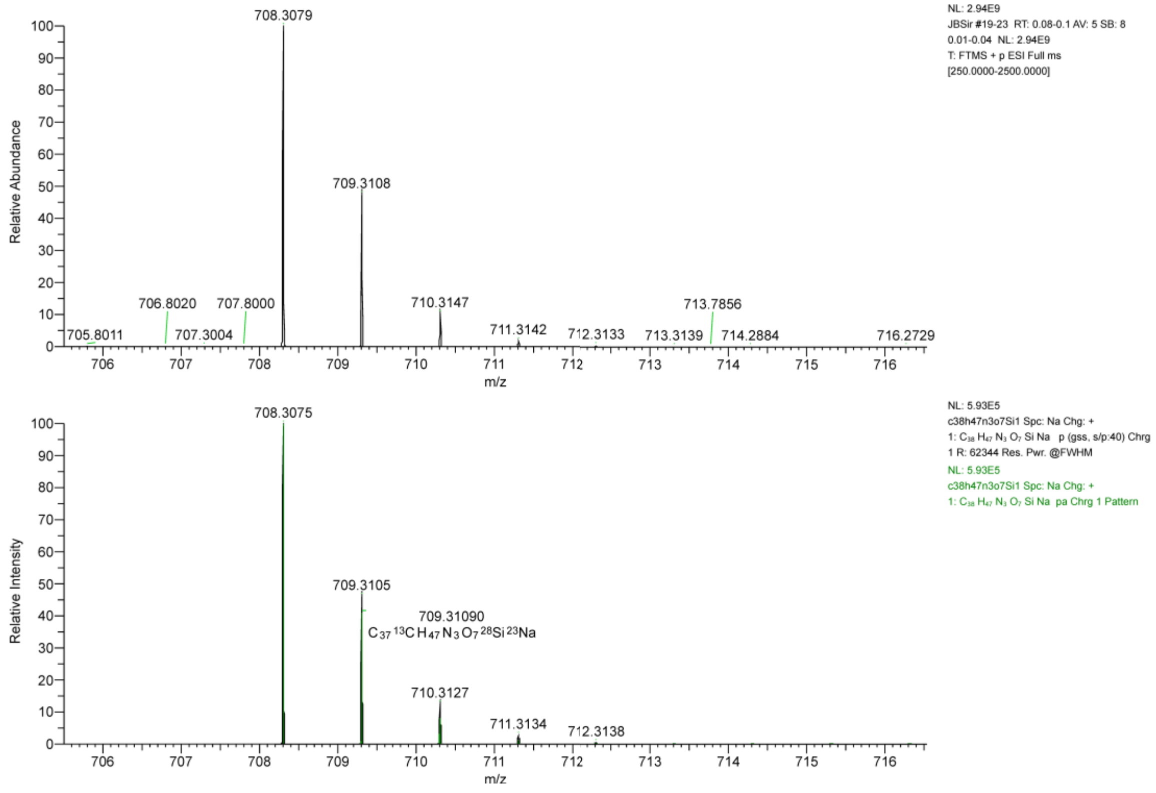

5-HMSiR-PEG$_4$-alkyne

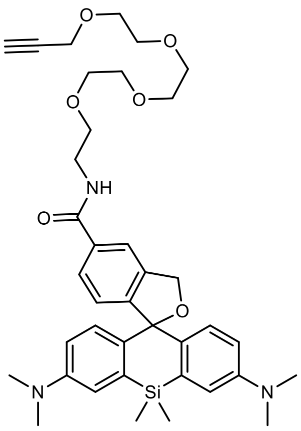

Reaction was performed following general synthesis procedure A from 5-HMSiR-COOH[27] (5 mg) and $NH_2$-PEG$_4$-alkyne using HATU as peptide coupling reagent. The compound was purified by preparative HPLC (preparative column: Pursuit 10 C18 10 µm, 250 x 50 mm, Agilent, flow rate: 150 ml/min, solvent A: MeOH, solvent B: $H_2O$ + 0.1% TFA; room temperature, gradient A:B - 2 min 30:70 isocratic, 2-20 min 30:70 to 100:0 gradient and 20-22 min 100:0 isocratic). Obtained 5.7 mg (77% yield) of a light blue solid.

$^1$H NMR (400 MHz, Methanol-$d_4$) δ 8.63 (t, $J$ = 5.5, 1H, -NH), 8.22 (s, 1H, Ar-H), 7.92 (d, $J$ = 8.1 Hz, 1H, Ar-H), 7.37 (d, $J$ = 2.8 Hz, 2H, Xan-H), 7.27 (d, $J$ = 7.9 Hz, 1H, Ar-H), 7.05 (d, $J$ = 9.6 Hz, 2H, Xan-H), 6.77 (dd, $J$ = 9.6, 2.9 Hz, 2H), Xan-H, 4.36 (s, 2H, Ar-$CH_2$-O-), 4.14 (d, $J$ = 2.4 Hz, 2H, -$CH_2$-Alkyne), 3.76 – 3.60 (m, 16H, $CH_2$ PEG), 3.35 (s, 12H, $N(CH_3)_2$), 2.82 (t, $J$ = 2.4 Hz, 1H, Alkyne-H), 0.62 (s, 3H, -Si-$CH_3$), 0.61 (s, 3H, -Si-$CH_3$).

ESI-MS, positive mode: m/z = 694.3 $[M+Na]^+$.

HRMS (ESI) calculated for $C_{38}H_{49}N_3O_6Si$ $[M+Na]^+$ 694.3283 found 694.3286.

HRMS:

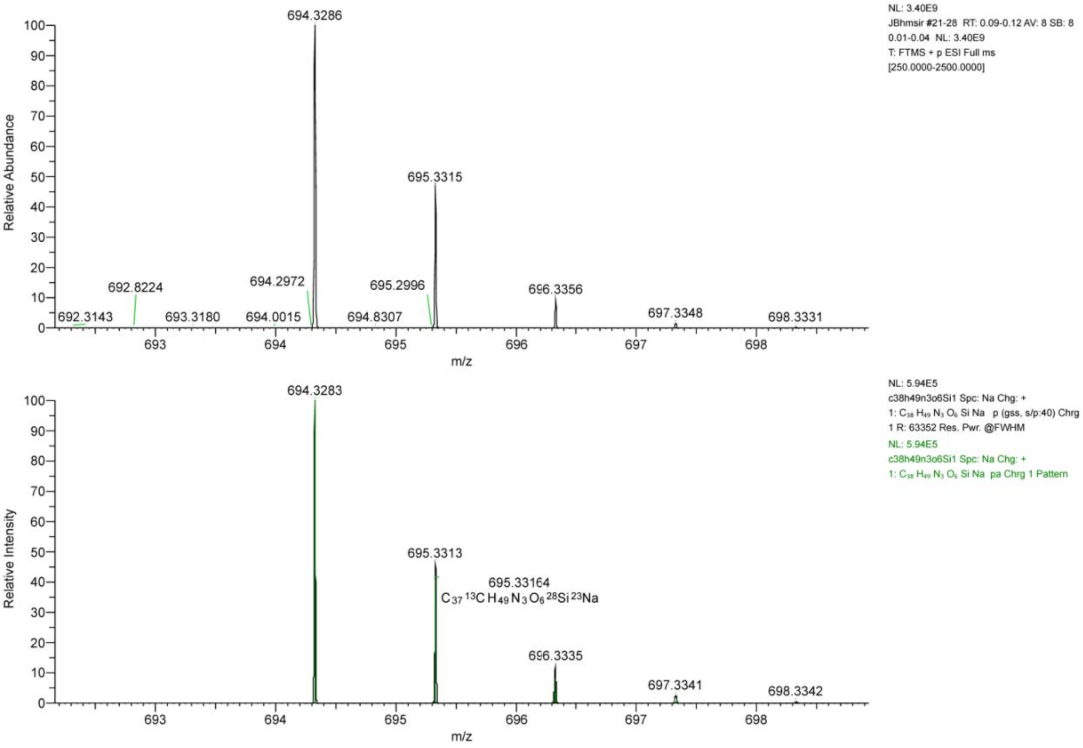

STAR-RED-alkyne

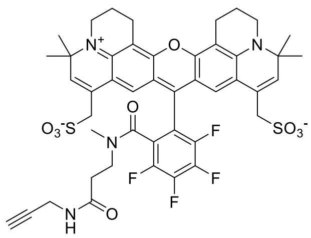

Reaction was performed following general synthesis procedure A from STAR-RED-COOH (1 mg, Abberior GmbH, product Nr.: STRED-0001) and propargyl amino hydrochloride using HATU as peptide coupling reagent. The compound was purified by preparative HPLC (preparative column: preparative column: Agilent 5 Prep-C18, 5 µm, 100 x 50 mm, flow rate: 40 ml/min, solvent A: MeCN, solvent B: $H_2O$ + 0.1% TFA; room temperature, gradient A:B - 2 min 30:70 isocratic, 2-20 min 30:70 to 100:0 gradient and 20-22 min 100:0 isocratic). Obtained 0.7 mg (67% yield) of a dark blue solid. The amount obtained was too small to characterize the compound by NMR.

ESI-MS, negative mode: m/z = 923.2[M]$^-$.

HRMS (ESI) calculated for $C_{45}H_{43}F_4N_4O_9S_2$ [M]$^-$ 923.2413 found 923.2407.

Copy of HRMS:

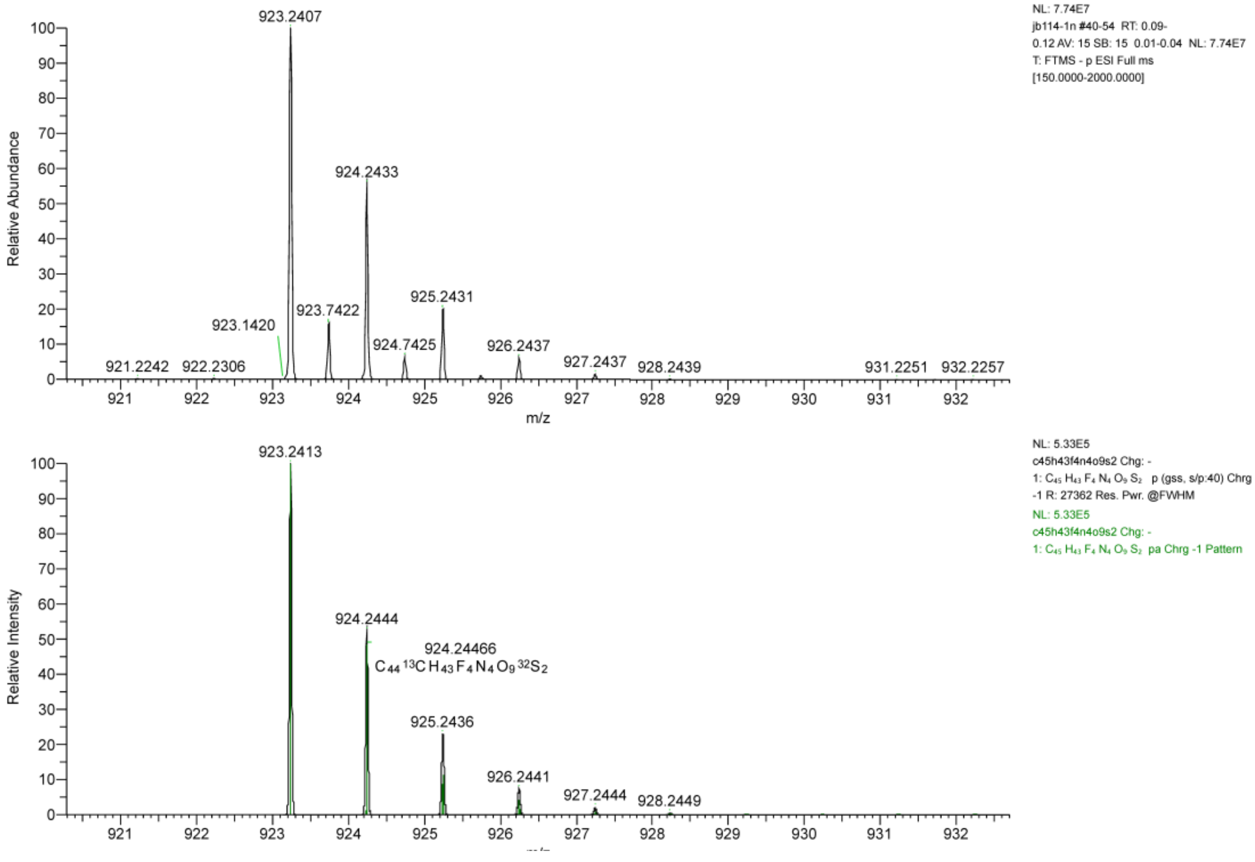

LC/MS:

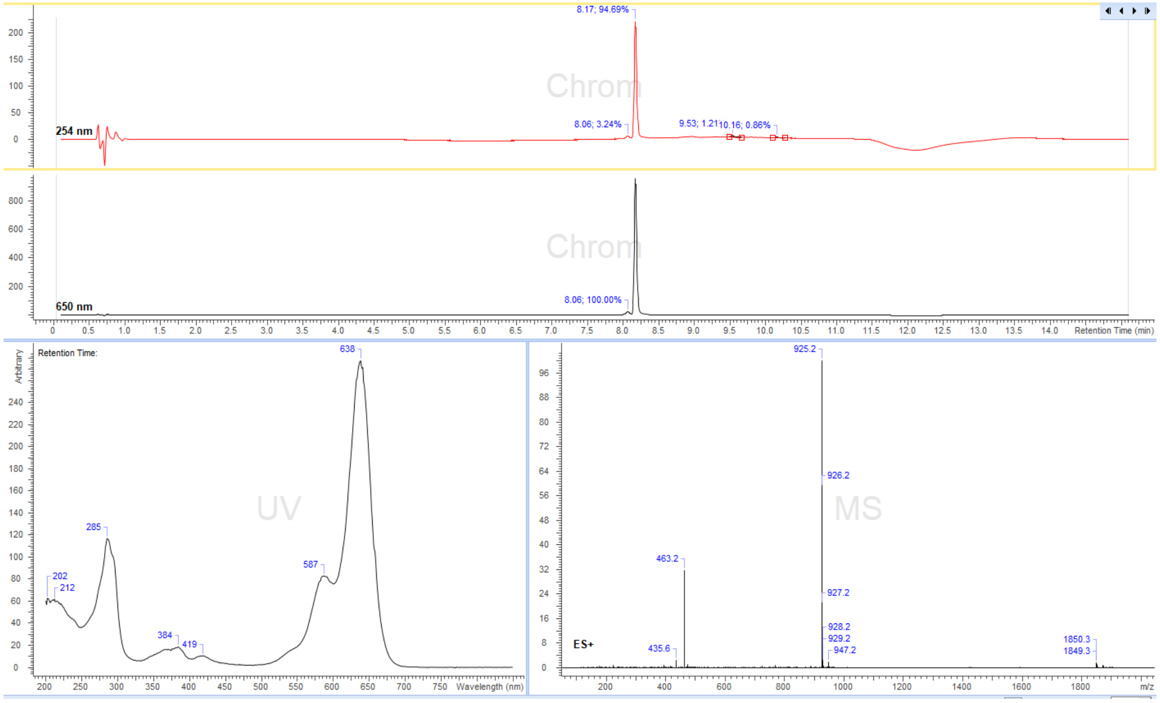

8.17; 94.69%
254 nm
Chrom
8.06; 3.24%
9.53; 1.21
10.16; 0.86%
650 nm
8.06; 100.00%
Retention Time (min)
Retention Time:
UV
638
285
587
202
212
384
419
Wavelength (nm)
MS
925.2
926.2
463.2
927.2
928.2
929.2
947.2
435.6
ES+
1850.3
1849.3
m/z

STAR-RED-PEG$_4$-alkyne

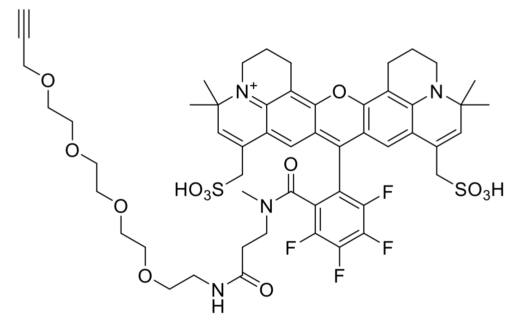

Reaction was performed following general synthesis procedure A from STAR-RED-COOH (1 mg, Abberior GmbH, product Nr.: STRED-0001) and $NH_2$-PEG$_4$-alkyne using HATU as peptide coupling reagent. The compound was purified by preparative HPLC (preparative column: preparative column: Agilent 5 Prep-C18, 5 µm, 100 x 50 mm, flow rate: 40 ml/min, solvent A: MeCN, solvent B: $H_2O$ + 0.1% TFA; room temperature, gradient A:B - 2 min 30:70 isocratic, 2-20 min 30:70 to 100:0 gradient and 20-22 min 100:0 isocratic). Obtained 0.9 mg (73% yield) of a dark blue solid. The amount obtained was too small to characterize the compound by NMR.

ESI-MS, negative mode: m/z = 1099.3$[M]^-$.

HRMS (ESI) calculated for $C_{53}H_{59}F_4N_4O_{13}S_2$ $[M]^-$ 1099.3462 found 1099.3459.

Copy of HRMS:

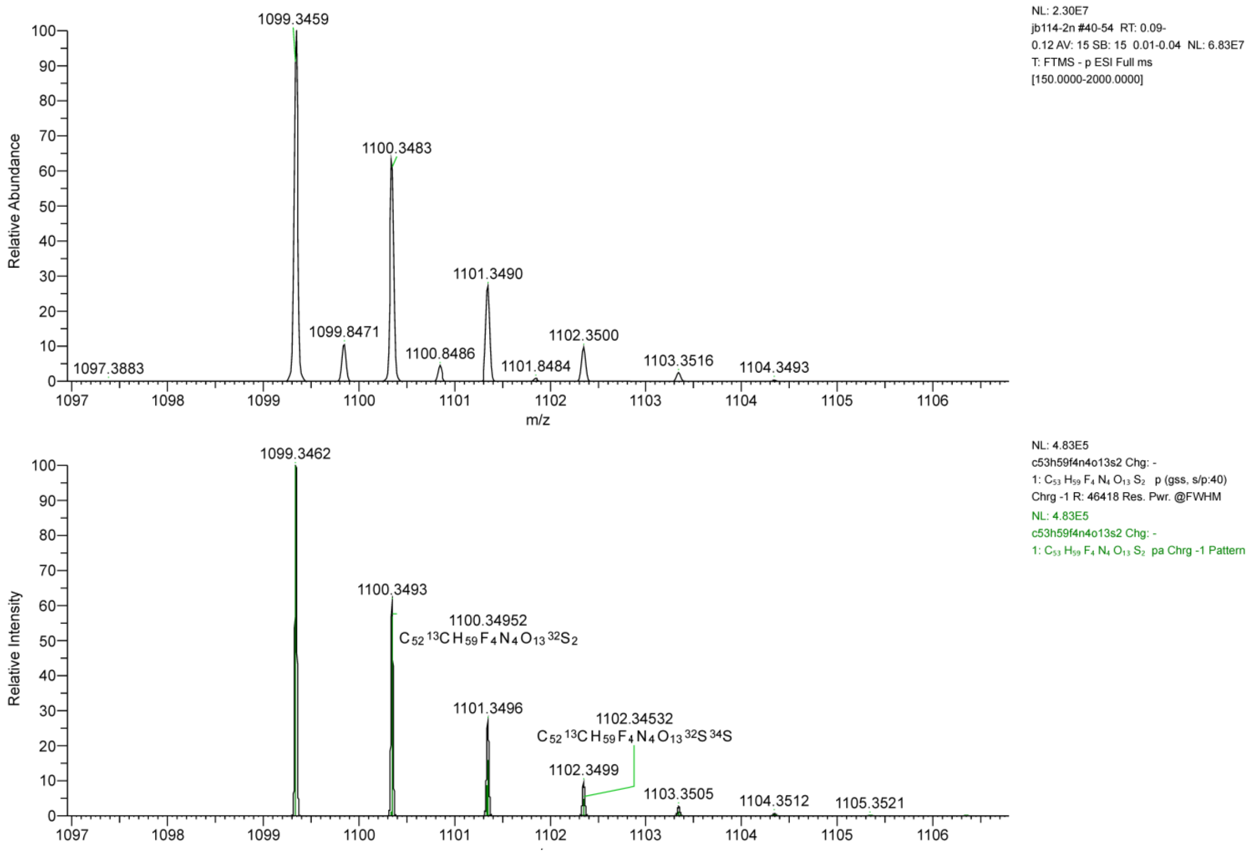

LC/MS:

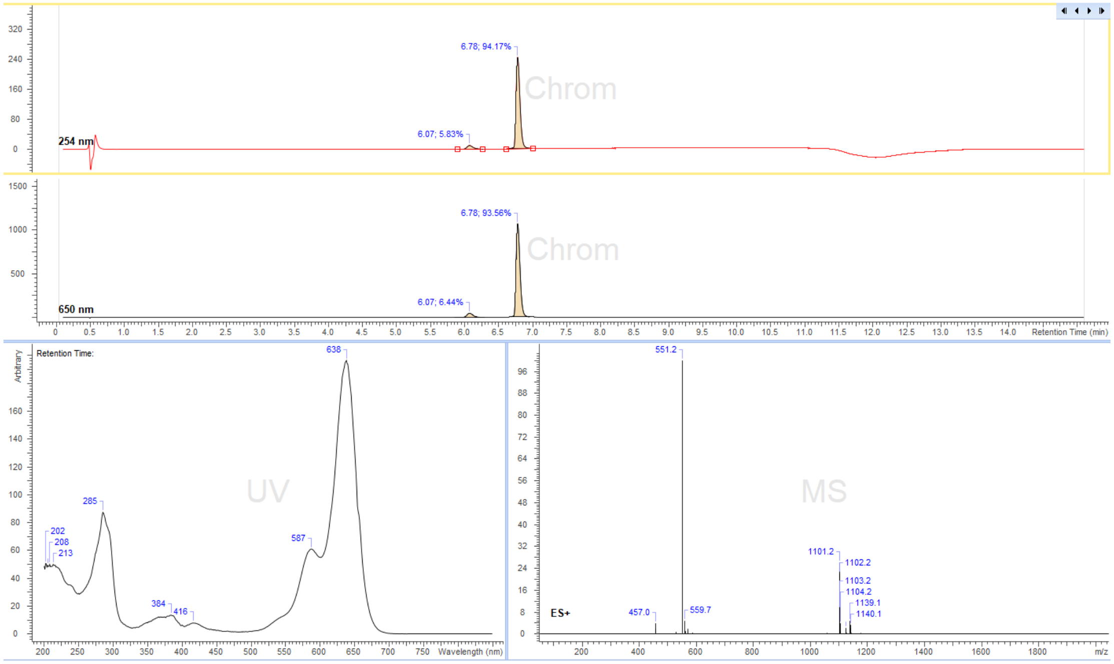

6.78; 94.17%
6.07; 5.83%
254 nm
Chrom
6.78; 93.56%
6.07; 6.44%
650 nm
Retention Time (min)
Retention Time:
UV
638
587
285
202
208
213
384
416
Wavelength (nm)
MS
551.2
559.7
457.0
1101.2
1102.2
1103.2
1104.2
1139.1
1140.1
ES+
m/z

5-TMR-$(CH_2)_3$-$N_3$

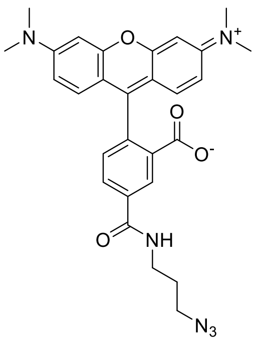

Reaction was performed following general synthesis procedure B from 5-TMR-NHS ester (5 mg, Lumiprobe Cat. Nr. 27120) and $NH_2$-$(CH_2)_3$-$N_3$ in dry DMSO. The product was purified by preparative HPLC (preparative column: Eurospher II 100-5 CN Column 250x30 mm, 5 µm, flow rate: 35 mL/min, solvent A: 10 mM ammonium formate (pH 3.5), solvent B: MeOH; room temperature, gradient A:B- 2 min 80:20 isocratic, 2-12 min 80:20 to 0:100 gradient,12-15 min 0:100 isocratic, 15-16 min 0:100 to 80:20 gradient and16-20 min 80:20 isocratic). Obtained 4.5 mg (92% yield) of a dark pink solid.

$^1$H NMR (400 MHz, $CD_3OD$) δ 8.53 (dd, J = 1.9, 0.5 Hz, 1H, Ar-H), 8.06 (dd, J = 7.9, 1.9 Hz, 1H, Ar-H), 7.37 (dd, J = 7.9, 0.5 Hz, 1H, Ar-H), 7.24 (d, J = 9.5 Hz, 2H, Xan-H), 7.02 (dd, J = 9.5, 2.5 Hz, 2H, Xan-H), 6.93 (d, J = 2.5 Hz, 2H, Xan-H), 3.54 (t, J = 6.8 Hz, 2H, CH2N3), 3.47 (t, J = 6.7 Hz, 2H, CH2NH), 3.28 (s, 12H, N(CH3)2), 1.94 (p, J = 6.8 Hz, 2H, CH2). NH proton not observed (exchange with solvent $CD_3OD$).

ESI-MS, positive mode: m/z = 513.1 $[M]^+$.

HRMS (ESI) calculated for $C_{28}H_{28}N_6O_4$ $[M]^+$ 513.2245, found 513.2246.

Copy of HRMS:

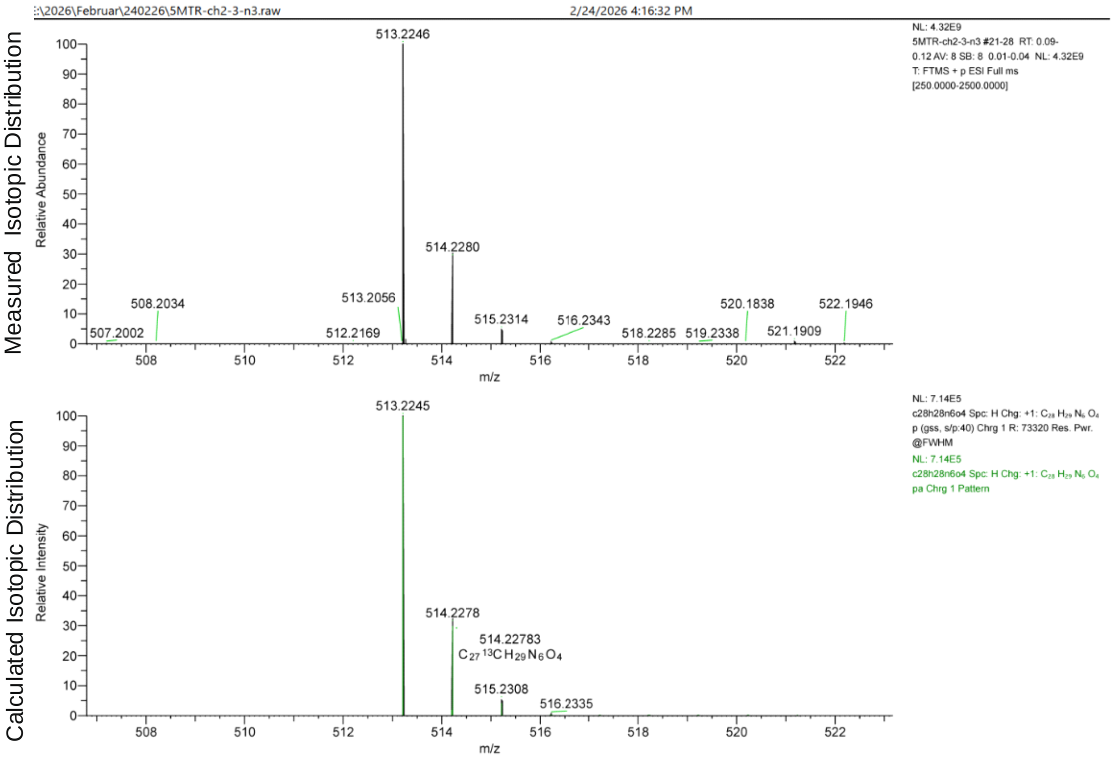

LC/MS:

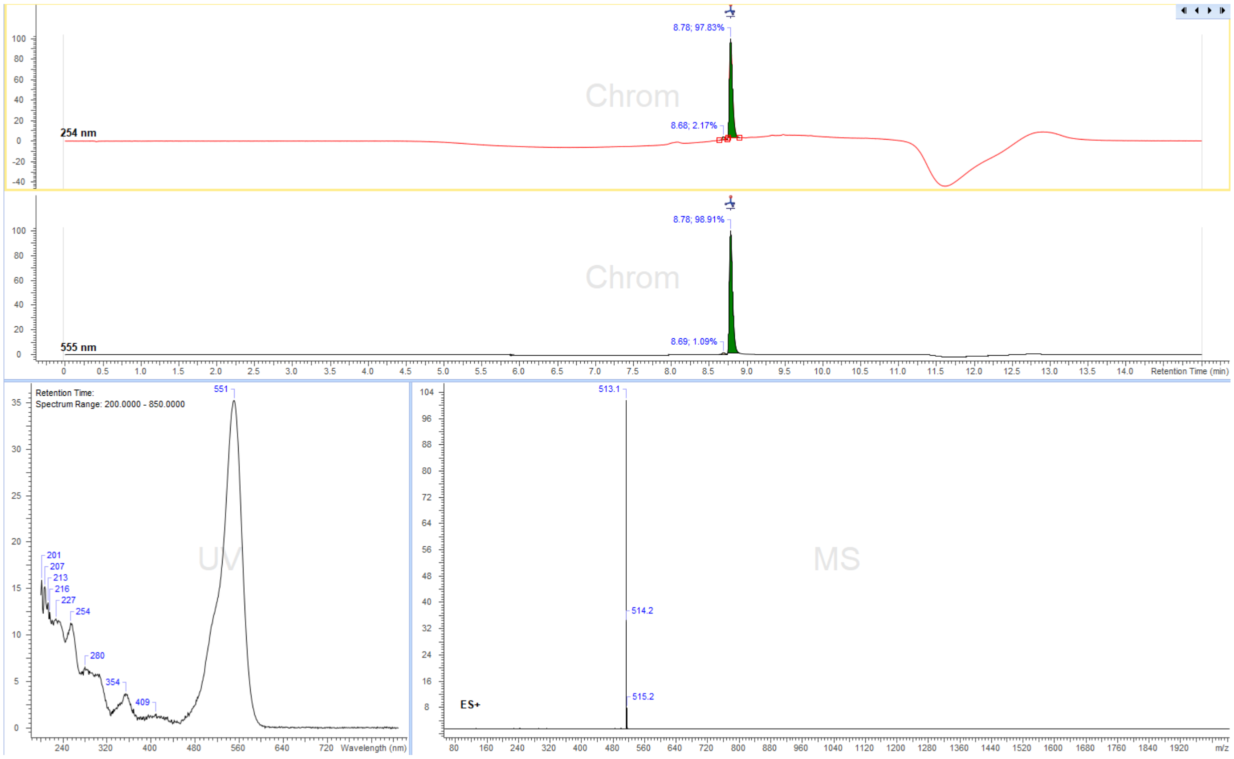

8.78; 97.83%
8.68; 2.17%
254 nm
8.78; 98.91%
8.69; 1.09%
555 nm
Retention Time (min)
Retention Time:
Spectrum Range: 200.0000 - 850.0000
551
201
207
213
216
227
254
280
354
409
Wavelength (nm)
513.1
514.2
515.2
ES+
m/z

6-TMR-$(CH_2)_3$-$N_3$

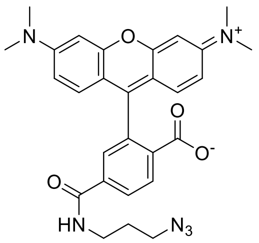

Reaction was performed following general synthesis procedure B from 6-TMR-NHS ester (5 mg, Lumiprobe, Cat. Nr. 28120) and $NH_2$-$(CH_2)_3$-$N_3$ in dry DMSO. The product was purified by preparative HPLC (preparative column: Eurospher II 100-5 CN Column 250x30 mm, 5 µm, flow rate: 35 mL/min, solvent A: 10 mM ammonium formate (pH 3.5), solvent B: MeOH; room temperature, gradient A:B-2 min 80:20 isocratic, 2-12 min 80:20 to 0:100 gradient, 12-15 min 0:100 isocratic, 15-16 min 0:100 to 80:20 gradient and 16-20 min 80:20 isocratic). Obtained 3 mg (61% yield) of a dark pink solid.

$^1$H NMR (400 MHz, $CD_3OD$) δ 8.19 – 8.15 (m, 1H, Ar-H), 8.11 – 8.07 (m, 1H, Ar-H), 7.72 (dd, J = 1.8, 0.5 Hz, 1H, Ar-H), 7.25 (d, J = 9.5 Hz, 2H, Xan-H), 7.02 (dd, J = 9.5, 2.5 Hz, 2H, Xan-H), 6.93 (d, J = 2.5 Hz, 2H, Xan-H), 3.46 (t, J = 6.9 Hz, 2H, CH2N3), 3.40 (t, J = 6.7 Hz, 2H, CH2NH), 3.28 (s, 12H, N(CH3)2), 1.87 (p, J = 6.7 Hz, 2H, CH2). NH proton not observed (exchange with solvent $CD_3OD$).

ESI-MS, positive mode: m/z = 513.2 $[M]^+$.

HRMS (ESI) calculated for $C_{28}H_{28}N_6O_4$ $[M]^+$513.2245, found 513.2248.

Copy of HRMS:

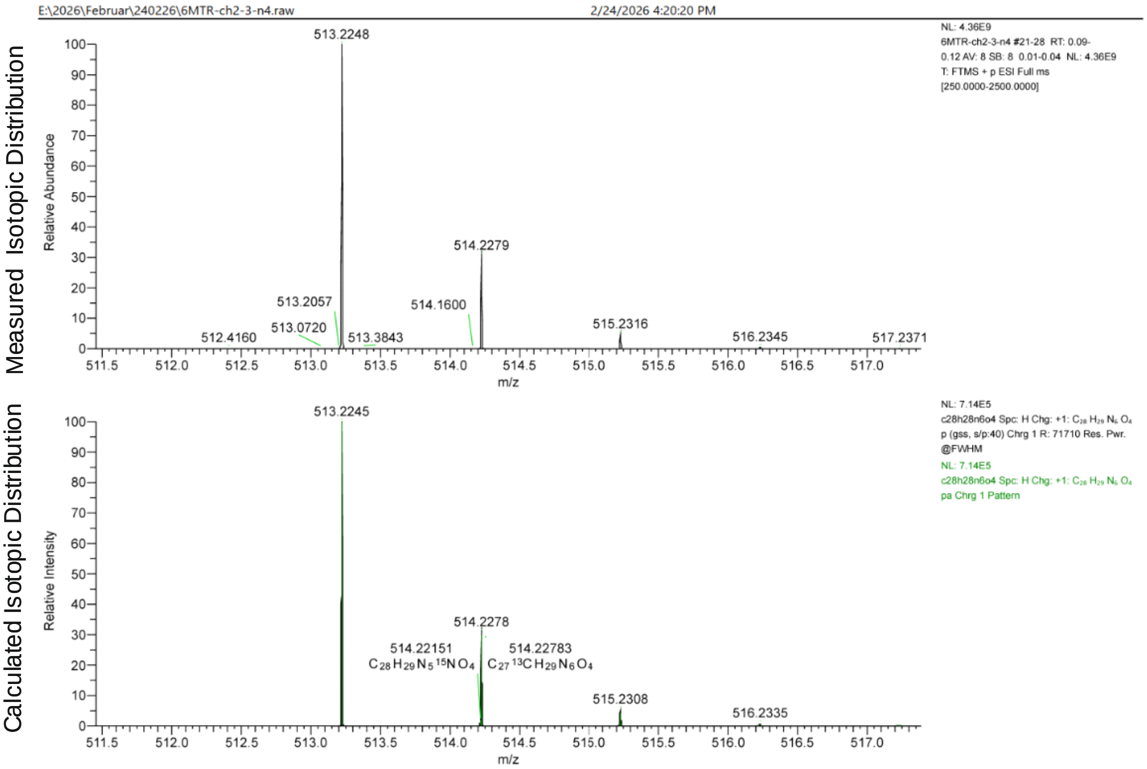

LC/MS:

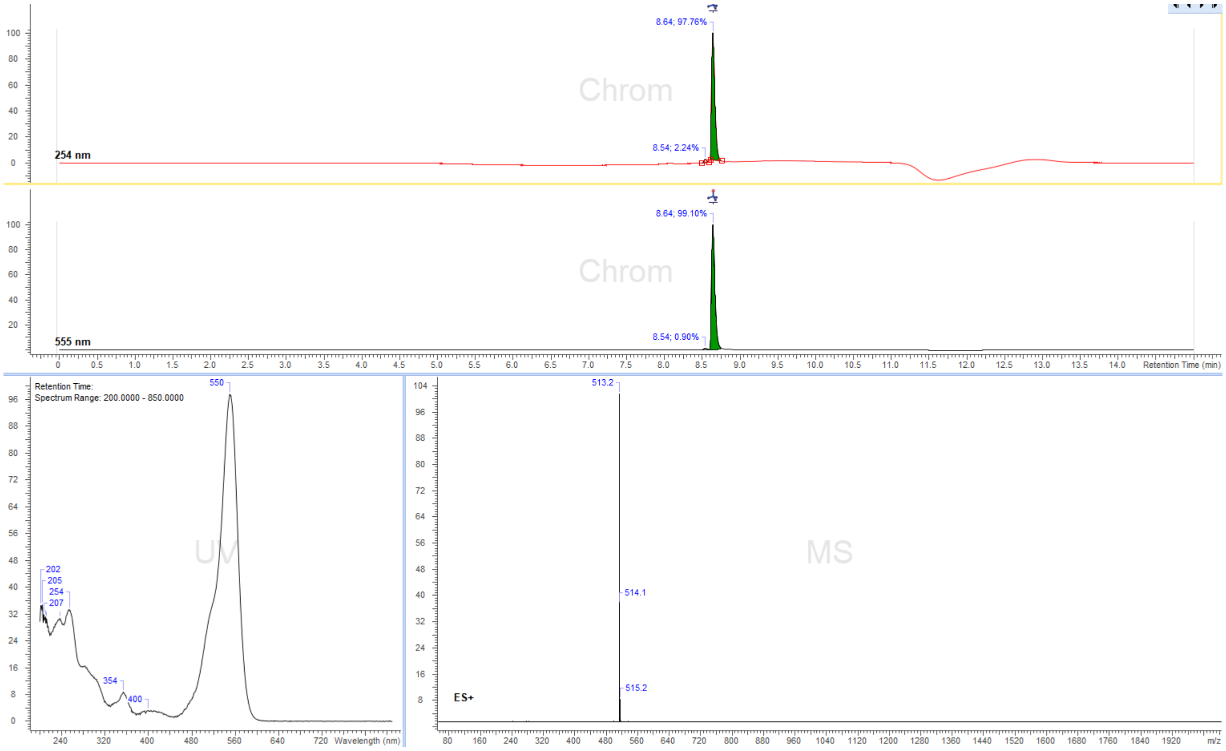
8.64; 97.76%
Chrom
8.54; 2.24%
254 nm
8.64; 99.10%
Chrom
555 nm
8.54; 0.90%
Retention Time (min)
Retention Time:
Spectrum Range: 200.0000 - 850.0000
550
202
205
254
207
354
400
UV
Wavelength (nm)
513.2
514.1
515.2
MS
ES+
m/z

Cy3B-$(CH_2)_3$-$N_3$

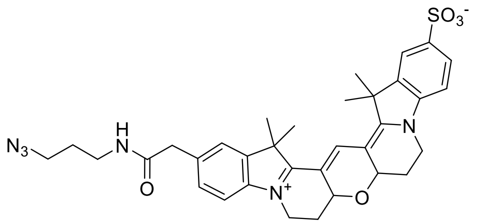

Reaction was performed following general synthesis procedure B from Cy3B-NHS ester (5 mg, Lumiprobe Cat. Nr. 29320) and $NH_2$-$(CH_2)_3$-$N_3$ in dry DMSO. The product was purified by preparative HPLC (preparative column: Eurospher II 100-5 CN Column 250x30 mm, 5 µm, flow rate: 30 mL/min, solvent A: 10 mM ammonium formate (pH 3.5), solvent B: MeOH; room temperature, gradient A:B-2 min 80:20 isocratic, 2-16 min 80:20 to 0:100 gradient, 16-19 min 0:100 isocratic, 19-20 min 0:100 to 80:20 gradient and 20-24 min 80:20 isocratic). Obtained 4.2 mg (86% yield) of a dark pink solid.

$^1H$ NMR (400 MHz, $(CD_3)_2SO$) δ 8.15 (t, J = 5.6 Hz, 1H, CONH), 7.99 (s, 1H, Ar-H), 7.80 (d, J = 1.6 Hz, 1H, Ar-H), 7.67 (dd, J = 8.2, 1.6 Hz, 1H, Ar-H), 7.49 (d, J = 1.1 Hz, 1H, Ar-H), 7.35 – 7.26 (m, 3H, Ar-H), 4.64 (dd, J = 11.4, 5.1 Hz, 2H, CH2-bridge), 4.31 (d, J = 14.1 Hz, 2H, CH2-bridge), 3.95 – 3.85 (m, 2H, CH2N), 3.48 (s, 2H, CH2-linker), 3.33 (t, J = 6.8 Hz, 2H, CH2N3), 3.17 – 3.06 (m, 2H, CH2NH), 1.96 (qd, J = 12.3, 5.3 Hz, 2H, CH2-ring), 1.70 (dd, J = 5.3, 1.7 Hz, 12H, 4 x CH3), 1.64 (q, J = 6.8 Hz, 2H, CH2). According COSY $H_2$ protons are under solvent $(CD_3)_2SO$.

ESI-MS, positive mode: m/z = 643.1 $[M]^+$.

HRMS (ESI) calculated for $C_{34}H_{38}N_6O_5S$ $[M+Na]^+$ 665.2517, found 665.2524.

Copy of HRMS:

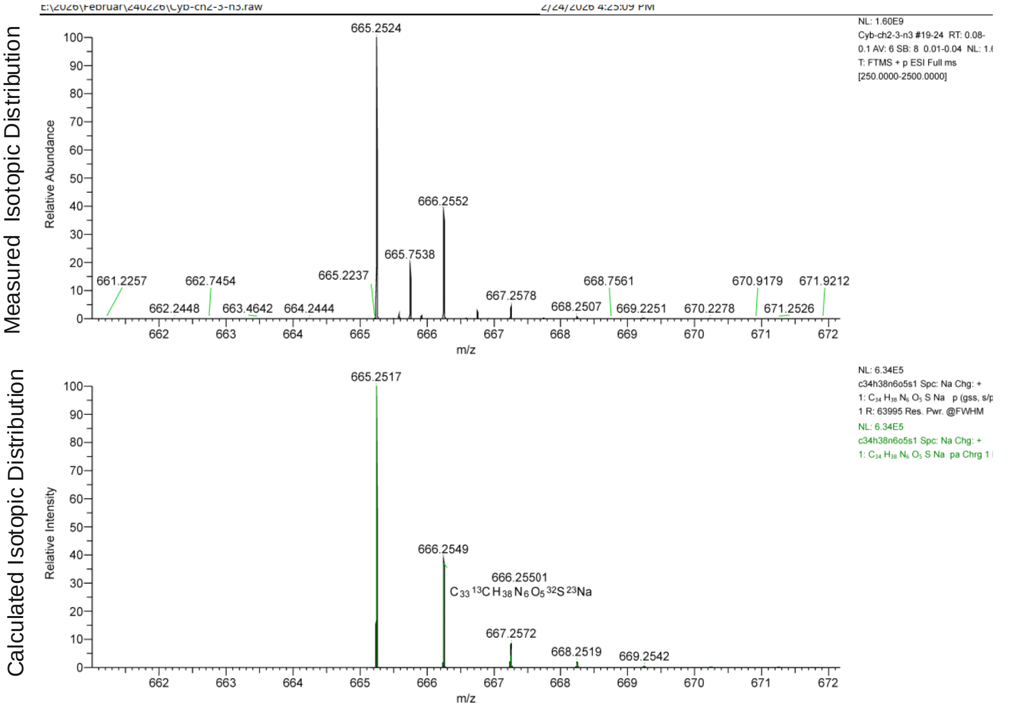

LC/MS:

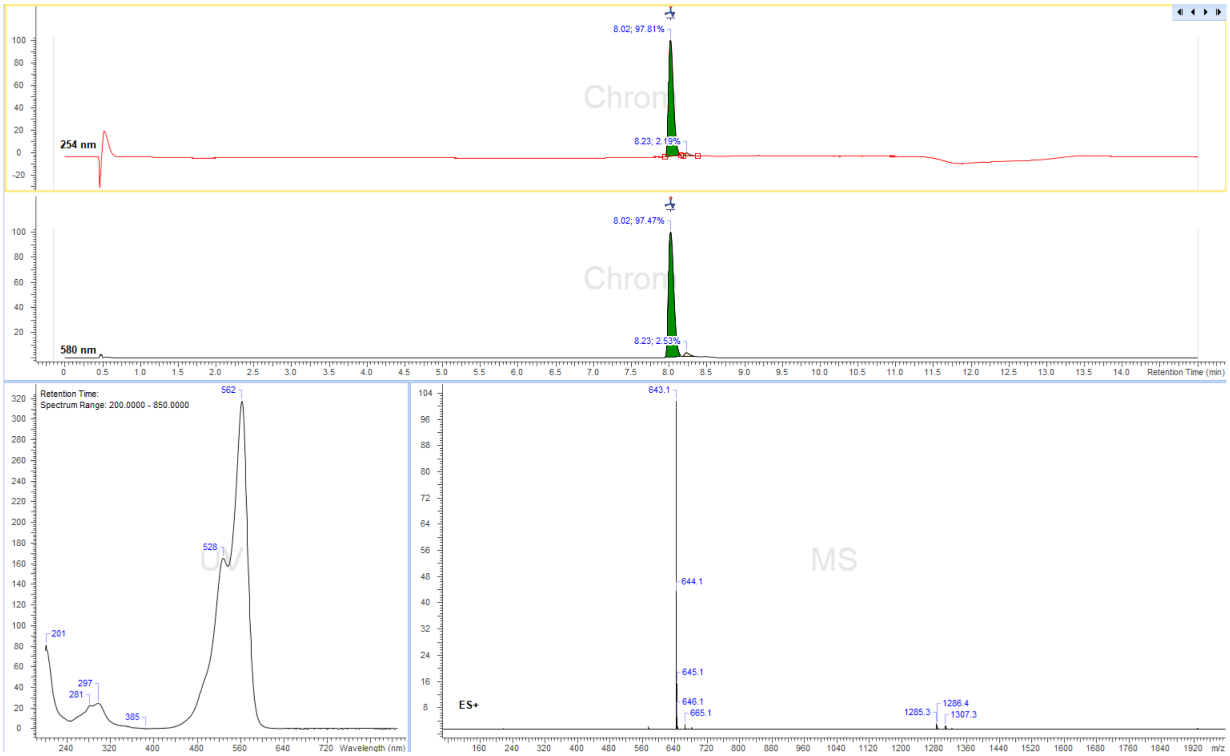
8.02; 97.81%
8.23; 2.19%
254 nm
Chron
8.02; 97.47%
8.23; 2.53%
580 nm
Retention Time (min)
Retention Time:
Spectrum Range: 200.0000 - 850.0000
562
528
201
297
281
385
UV
Wavelength (nm)
643.1
644.1
645.1
646.1
665.1
1285.3
1286.4
1307.3
ES+
MS
m/z

5-TMR-$PEG_3$-$N_3$

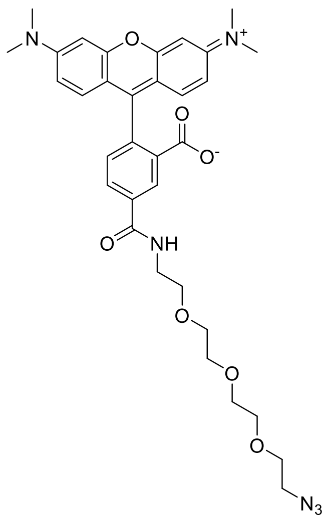

Reaction was performed following general synthesis procedure B 5-TMR-NHS ester (5 mg, Lumiporbe Cat. Nr. 27120) and $NH_2$-$PEG_3$-$N_3$ in dry DMSO. The product was purified by preparative HPLC (preparative column: Eurospher II 100-5 CN Column 250x30 mm, 5 µm, flow rate: 35 mL/min, solvent A: 10 mM ammonium formate (pH 3.5), solvent B: MeOH; room temperature, gradient A:B-2 min 80:20 isocratic, 2-12 min 80:20 to 0:100 gradient, 12-15 min 0:100 isocratic, 15-16 min 0:100 to 80:20 gradient and 16-20 min 80:20 isocratic). Obtained 4.1 mg (70% yield) of a dark pink solid.

$^1H$ NMR (400 MHz, $CD_3OD$) δ 8.56 (dd, J = 1.9, 0.5 Hz, 1H, Ar-H), 8.08 (dd, J = 7.9, 1.9 Hz, 1H, Ar-H), 7.41 – 7.34 (m, 1H, Ar-H), 7.25 (d, J = 9.5 Hz, 2H, Xan-H), 7.02 (dd, J = 9.5, 2.5 Hz, 2H, Xan-H), 6.93 (d, J = 2.5 Hz, 2H, Xan-H), 3.75 – 3.63 (m, 14H, CH2 PEG), 3.37 – 3.33 (m, 2H, CH2N3), 3.28 (s, 12H, N(CH3)2). NH proton not observed (exchange with solvent $CD_3OD$).

ESI-MS, positive mode: m/z = 631.3 $[M]^+$.

HRMS (ESI) calculated for $C_{33}H_{38}N_6O_7$ $[M]^+$ 631.2875, found 631.2874.

Copy of HRMS:

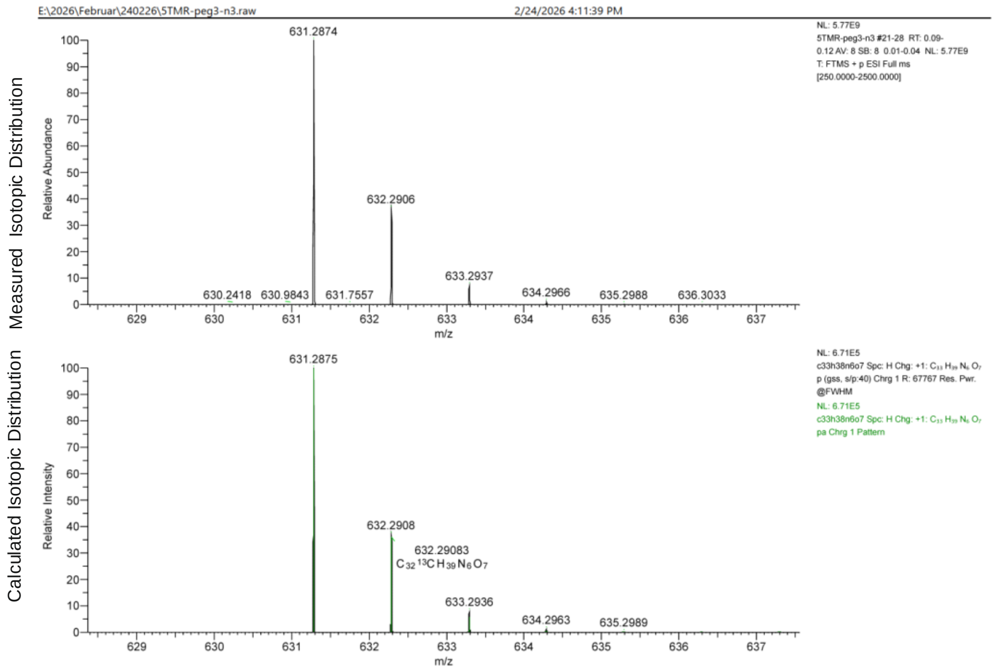

LC/MS:

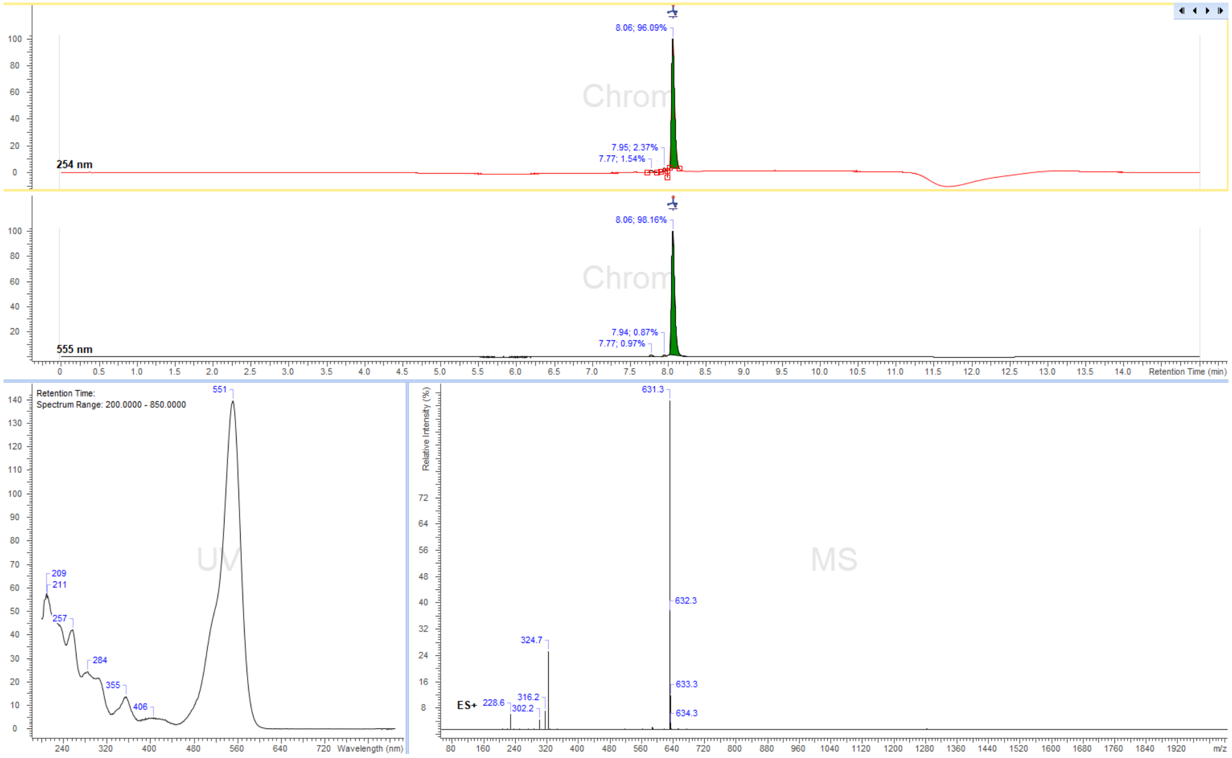

8.06; 96.09%
7.95; 2.37%
7.77; 1.54%
254 nm
Chrom
8.06; 98.16%
7.94; 0.87%
7.77; 0.97%
555 nm
Retention Time (min)
Retention Time:
Spectrum Range: 200.0000 - 850.0000
551
209
211
257
284
355
406
UV
Wavelength (nm)
Relative Intensity (%)
631.3
632.3
633.3
634.3
324.7
316.2
302.2
228.6
ES+
MS
m/z

6-TMR-PEG$_3$-N$_3$

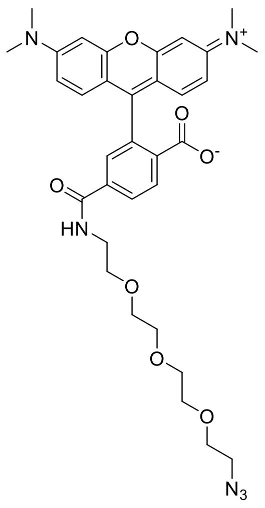

Reaction was performed following general synthesis procedure B from 6-TMR-NHS ester (5 mg, Lumiprobe, Cat. Nr. 28120) and NH$_2$-PEG$_3$-N$_3$ in dry DMSO. The product was purified by preparative HPLC (preparative column: Eurospher II 100-5 CN Column 250x30 mm, 5 µm, flow rate: 35 mL/min, solvent A: 10 mM ammonium formate (pH 3.5), solvent B: MeOH; room temperature, gradient A:B-2 min 80:20 isocratic, 2-12 min 80:20 to 0:100 gradient, 12-15 min 0:100 isocratic, 15-16 min 0:100 to 80:20 gradient and 16-20 min 80:20 isocratic). Obtained 3.6 mg (61% yield) of a dark pink solid.

$^1$H NMR (400 MHz, CD3OD) δ 8.17 (dd, J = 8.1, 0.6 Hz, 1H, Ar-H), 8.11 (dd, J = 8.1, 1.8 Hz, 1H, Ar-H), 7.73 (dd, J = 1.8, 0.5 Hz, 1H, Ar-H), 7.26 (d, J = 9.5 Hz, 2H, Xan-H), 7.02 (dd, J = 9.5, 2.5 Hz, 2H, Xan-H), 6.93 (d, J = 2.5 Hz, 2H, Xan-H), 3.68 – 3.54 (m, 16H, 8 x CH2 PEG), 3.29 (s, 12H, N(CH3)2). NH proton not observed (exchange with solvent CD$_3$OD).

ESI-MS, positive mode: m/z = 631.3 $[M]^+$.

HRMS (ESI) calculated for $C_{33}H_{38}N_6O_7$ $[M]^+$ 631.2875, found 631.2878.

Copy of HRMS:

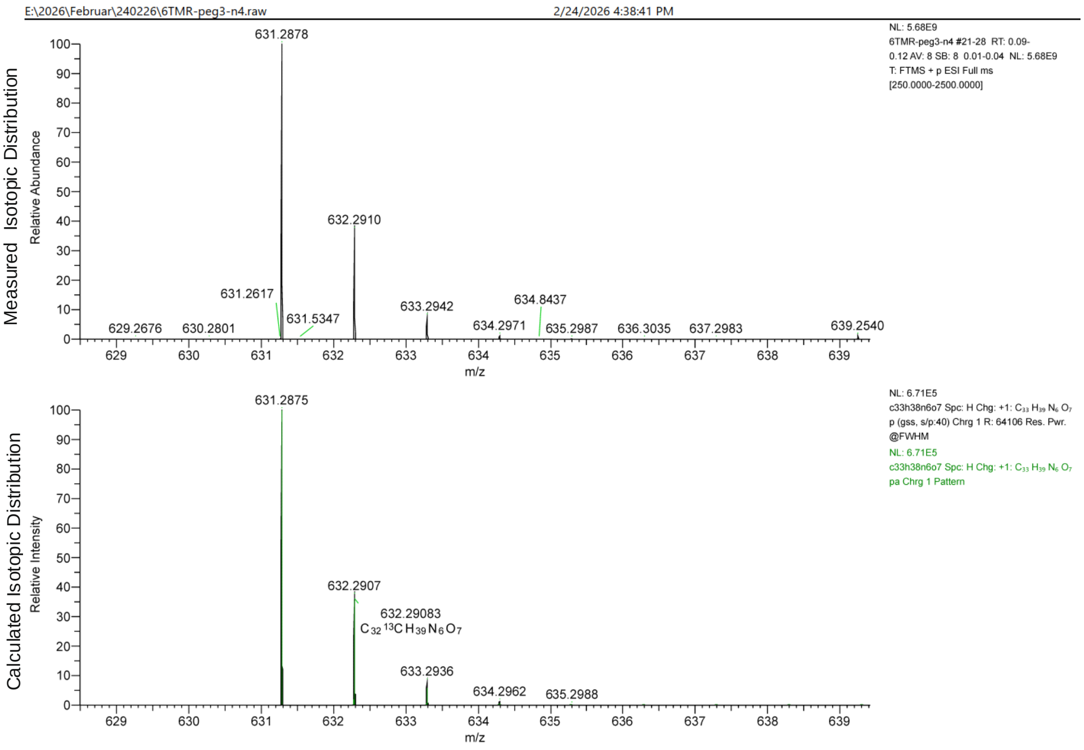

LC/MS:

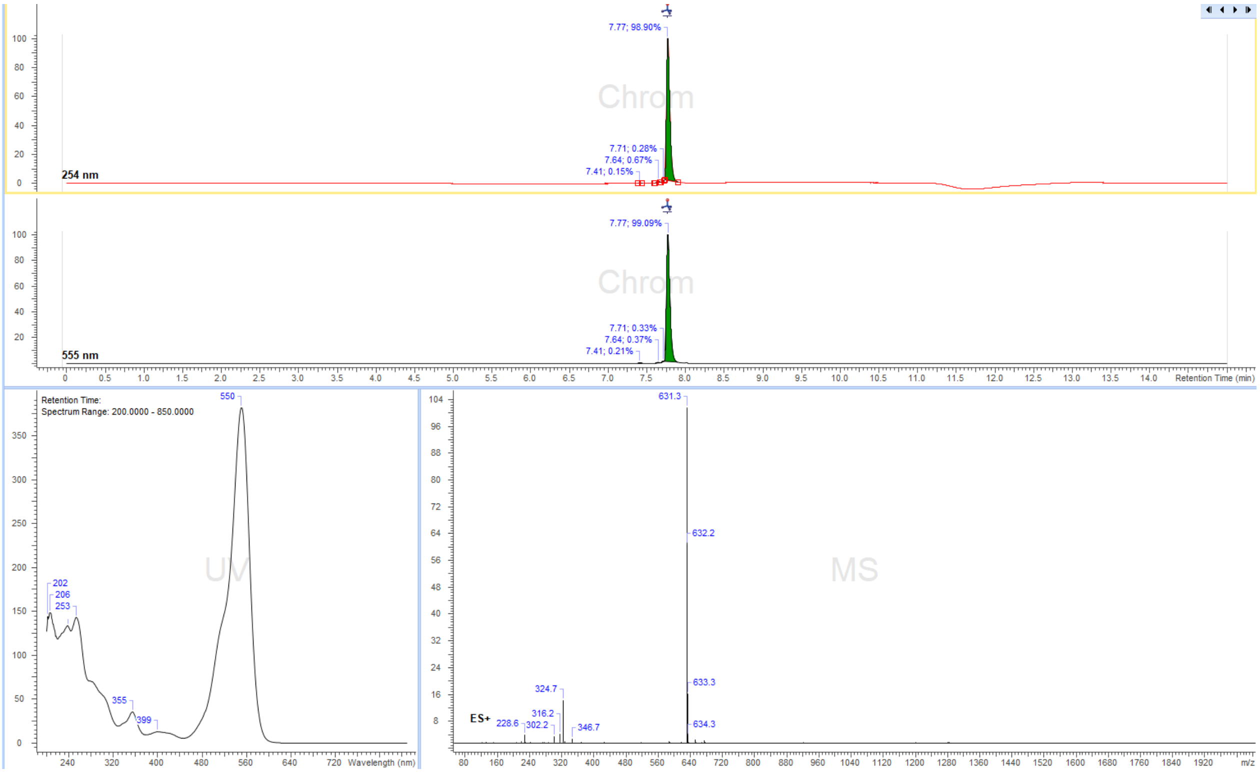

7.77; 98.90%
7.71; 0.28%
7.64; 0.67%
7.41; 0.15%
254 nm
7.77; 99.09%
7.71; 0.33%
7.64; 0.37%
7.41; 0.21%
555 nm
Retention Time (min)
Retention Time:
Spectrum Range: 200.0000 - 850.0000
550
202
206
253
355
399
Wavelength (nm)
631.3
632.2
633.3
634.3
324.7
316.2
302.2
228.6
346.7
ES+
m/z

Cy3B-PEG$_3$-N$_3$

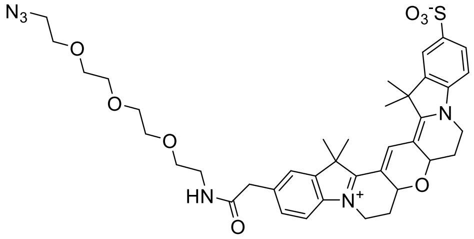

Reaction was performed following general synthesis procedure B from Cy3B-NHS ester (5 mg, Lumiprobe, Cat. Nr. 29320) and NH$_2$-PEG$_3$-N$_3$ in dry DMSO. The product was purified by preparative HPLC (preparative column: Eurospher II 100-5 CN Column 250x30 mm, 5 µm, flow rate: 35 mL/min, solvent A: 10 mM ammonium formate (pH 3.5), solvent B: MeOH; room temperature, gradient A:B-2 min 80:20 isocratic, 2-12 min 80:20 to 0:100 gradient, 12-15 min 0:100 isocratic, 15-16 min 0:100 to 80:20 gradient and 16-20 min 80:20 isocratic). Obtained 2.7 mg (47% yield) of a dark pink solid.

$^1$H NMR (400 MHz, CD3OD) δ 8.19 (s, 1H, meso-CH), 7.93 – 7.88 (m, 2H, Ar-H), 7.50 (d, J = 1.2 Hz, 1H, Ar-H), 7.40 (dd, J = 8.2, 1.6 Hz, 1H, Ar-H), 7.29 (dd, J = 8.3, 0.6 Hz, 2H, Ar-H), 4.69 (dd, J = 11.6, 5.0 Hz, 2H, bridge-CH2), 4.42 – 4.31 (m, 2H, bridge-CH2), 3.95 (dtd, J = 17.4, 14.7, 4.1 Hz, 2H, CH2N-ring), 3.69 – 3.60 (m, 12H, CH2 PEG), 3.56 (dd, J = 5.8, 5.1 Hz, 2H, CH2NH), 3.38 (dt, J = 6.7, 5.2 Hz, 4H, CH2N3 and CH2 PEG), 2.65 – 2.56 (m, 2H, CH2-ring), 2.03 (qd, J = 12.3, 5.2 Hz, 2H, CH2-ring), 1.78 (s, 12H, 4 x CH3). ). NH proton not observed (exchange with solvent CD$_3$OD).

ESI-MS, positive mode: m/z = 761.3 [M]$^+$.

HRMS (ESI) calculated for $C_{39}H_{48}N_6O_8S$ [M]$^+$ 761.3327, found 761.3329.

Copy of HRMS:

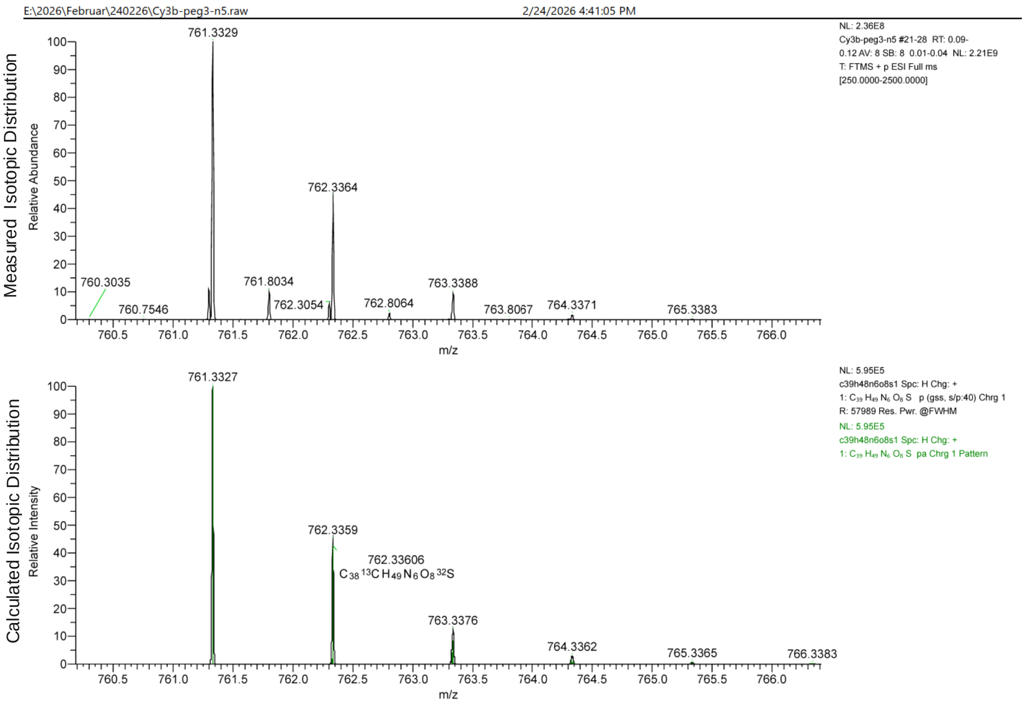

LC/MS:

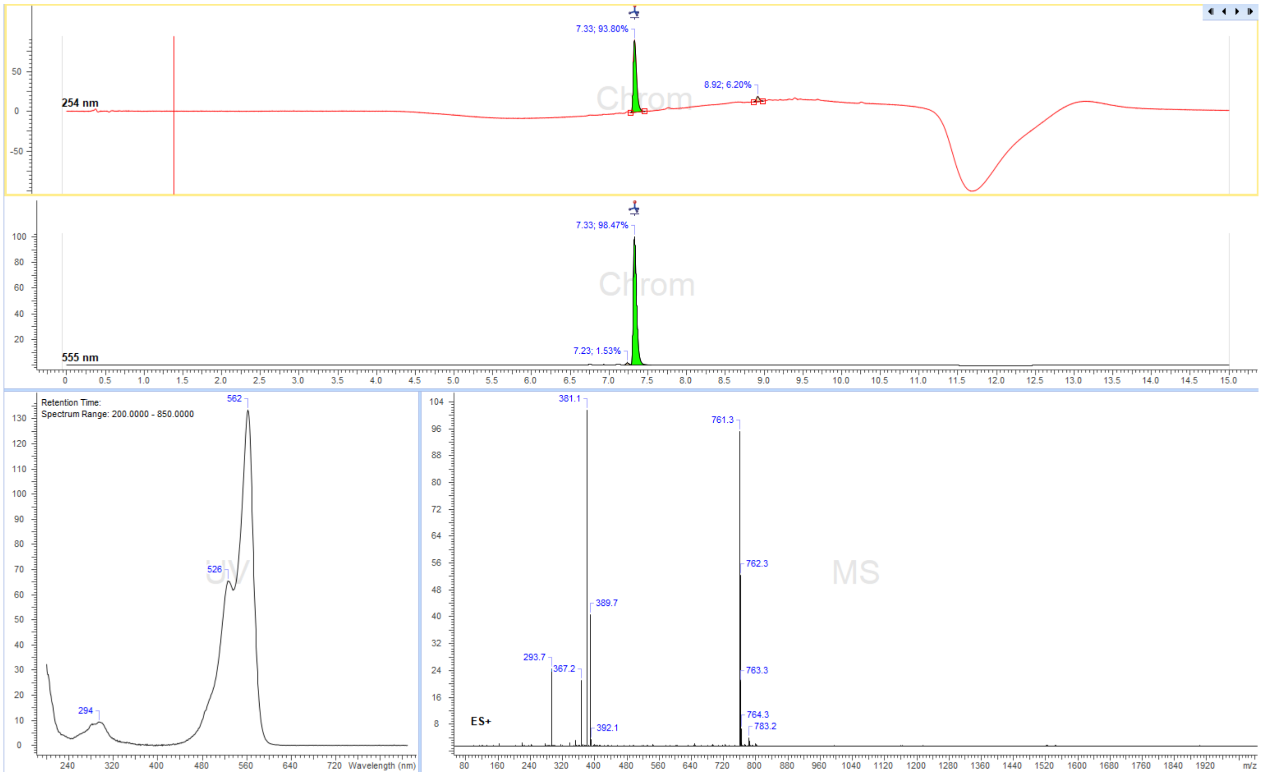
7.33; 93.80%
8.92; 6.20%
254 nm
7.33; 98.47%
7.23; 1.53%
555 nm
Retention Time:
Spectrum Range: 200.0000 - 850.0000
562
526
294
Wavelength (nm)
381.1
761.3
762.3
389.7
293.7
367.2
763.3
764.3
783.2
392.1
ES+
m/z

5-TMR-PEG$_{10}$-N$_3$

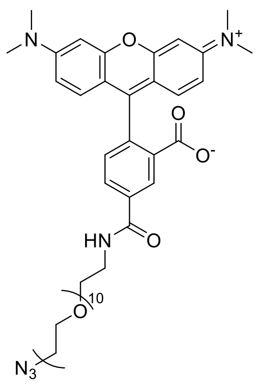

Reaction was performed following general synthesis procedure B from 5-TMR-NHS ester (5 mg, Lumiprobe, Cat. Nr. 27120) and NH$_2$-PEG$_{10}$-N$_3$ in dry DMSO. The product was purified by preparative HPLC (preparative column: Eurospher II 100-5 CN Column 250x30 mm, 5 µm, flow rate: 35 mL/min, solvent A: 10 mM ammonium formate (pH 3.5), solvent B: MeOH; room temperature, gradient A:B-2 min 80:20 isocratic, 2-12 min 80:20 to 0:100 gradient, 12-15 min 0:100 isocratic, 15-16 min 0:100 to 80:20 gradient and 16-20 min 80:20 isocratic). Obtained 7.6 mg (85% yield) of a dark pink solid.

$^{1}$H NMR (400 MHz, CD$_3$OD) δ 8.59 (dd, J = 1.9, 0.5 Hz, 1H, Ar-H), 8.11 (dd, J = 7.9, 1.9 Hz, 1H, Ar-H), 7.40 (dd, J = 7.9, 0.5 Hz, 1H, Ar-H), 7.23 (d, J = 9.5 Hz, 2H, Xan-H), 7.03 (dd, J = 9.5, 2.5 Hz, 2H, Xan-H), 6.94 (d, J = 2.5 Hz, 2H, Xan-H), 3.75 – 3.60 (m, 42H, 20 x CH2O, CH2N3), 3.38 – 3.35 (m, 2H, CH2NH), 3.29 (s, 12H, N(CH3)2). NH proton not observed (exchange with solvent CD$_3$OD).

ESI-MS, positive mode: m/z = 939.4 [M]$^+$.

HRMS (ESI) calculated for $C_{47}H_{66}N_6O_{14}$ [M]$^+$ 939.4710, found 939.4699.

Copy of HRMS:

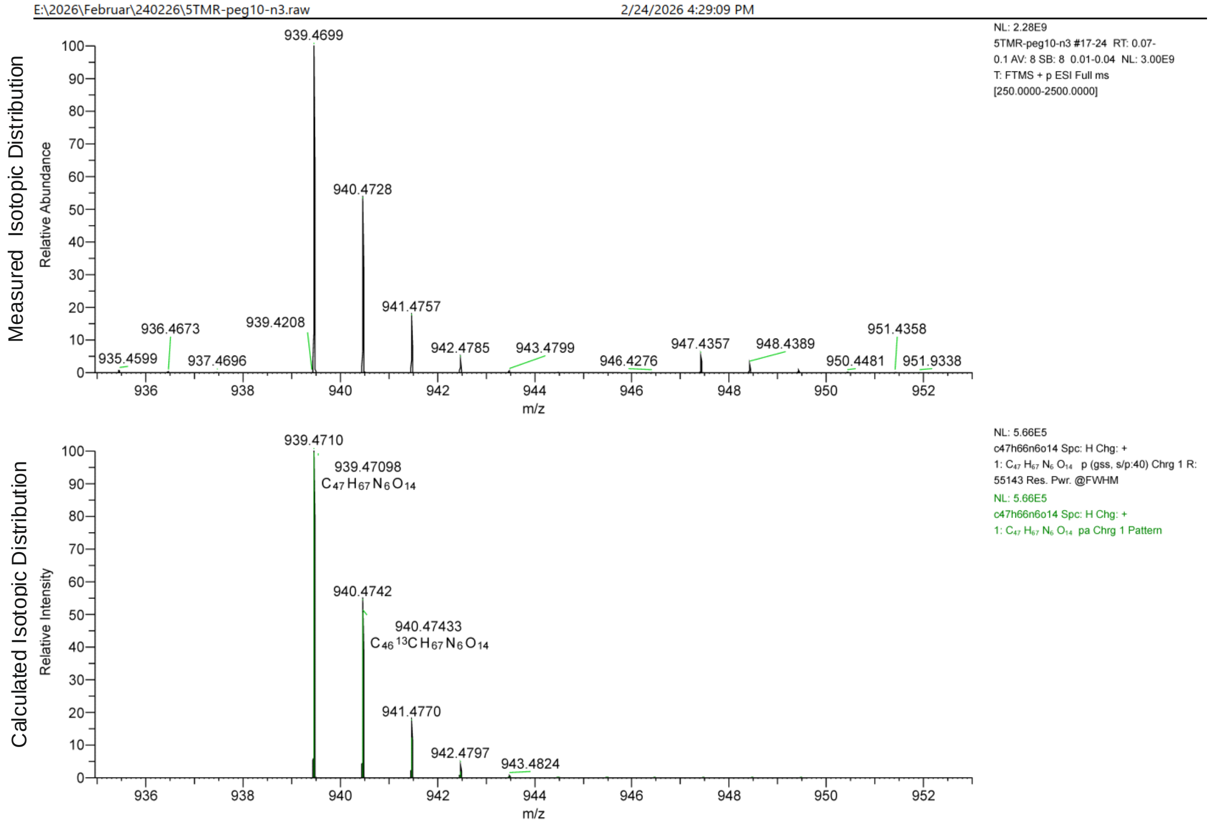

LC/MS:

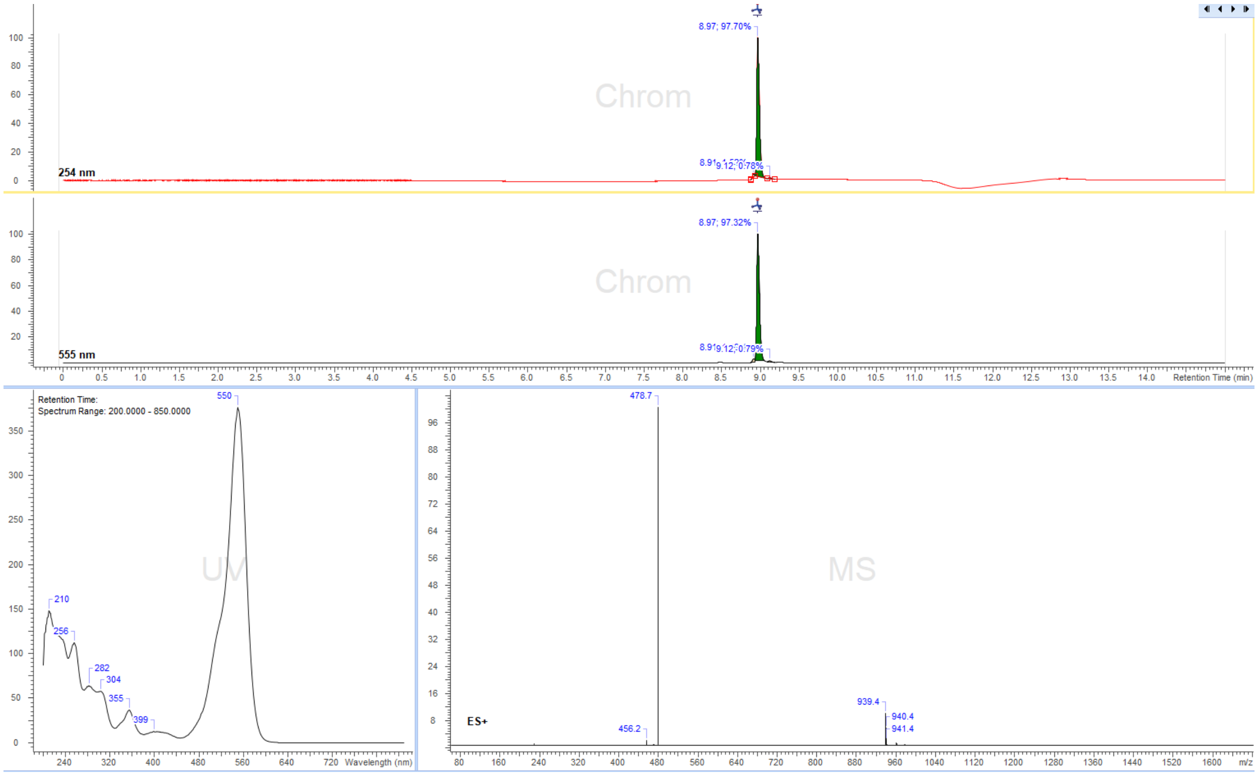

8.97; 97.70%
Chrom
254 nm
8.97; 97.32%
Chrom
555 nm
Retention Time (min)
Retention Time:
Spectrum Range: 200.0000 - 850.0000
550
210
256
282
304
355
399
UV
Wavelength (nm)
478.7
MS
939.4
940.4
941.4
456.2
ES+
m/z

6-TMR-PEG$_{10}$-N$_3$

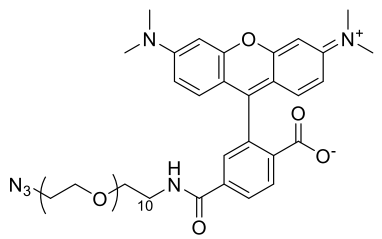

Reaction was performed following general synthesis procedure B from 6-TMR-NHS ester (5 mg, Lumiprobe, Cat. Nr. 28120) and NH$_2$-PEG$_{10}$-N$_3$ in dry DMSO. The product was purified by preparative HPLC (preparative column: Eurospher II 100-5 CN Column 250x30 mm, 5 µm, flow rate: 35 mL/min, solvent A: 10 mM ammonium formate (pH 3.5), solvent B: MeOH; room temperature, gradient A:B-2 min 80:20 isocratic, 2-12 min 80:20 to 0:100 gradient, 12-15 min 0:100 isocratic, 15-16 min 0:100 to 80:20 gradient and 16-20 min 80:20 isocratic). Obtained 7.1 mg (80% yield) of a dark pink solid.

$^1$H NMR (400 MHz, CD3OD) δ 8.18 (dd, J = 8.1, 0.5 Hz, 1H, Ar-H), 8.12 (dd, J = 8.1, 1.8 Hz, 1H, Ar-H), 7.74 (dd, J = 1.8, 0.6 Hz, 1H, Ar-H), 7.26 (d, J = 9.5 Hz, 2H, Xan-H), 7.03 (dd, J = 9.5, 2.5 Hz, 2H, Xan-H), 6.94 (d, J = 2.5 Hz, 2H, Xan-H), 3.68 – 3.53 (m, 42H, 20 x CH2O, CH2N3), 3.38 – 3.35 (m, 2H, CH2NH), 3.29 (s, 12H, N(CH3)2). NH proton not observed (exchange with solvent CD3OD).

ESI-MS, positive mode: m/z = 939.4 [M]$^+$.

HRMS (ESI) calculated for $C_{47}H_{66}N_6O_{14}$ [M]$^+$ 939.4710, found 939.4714.

Copy of HRMS:

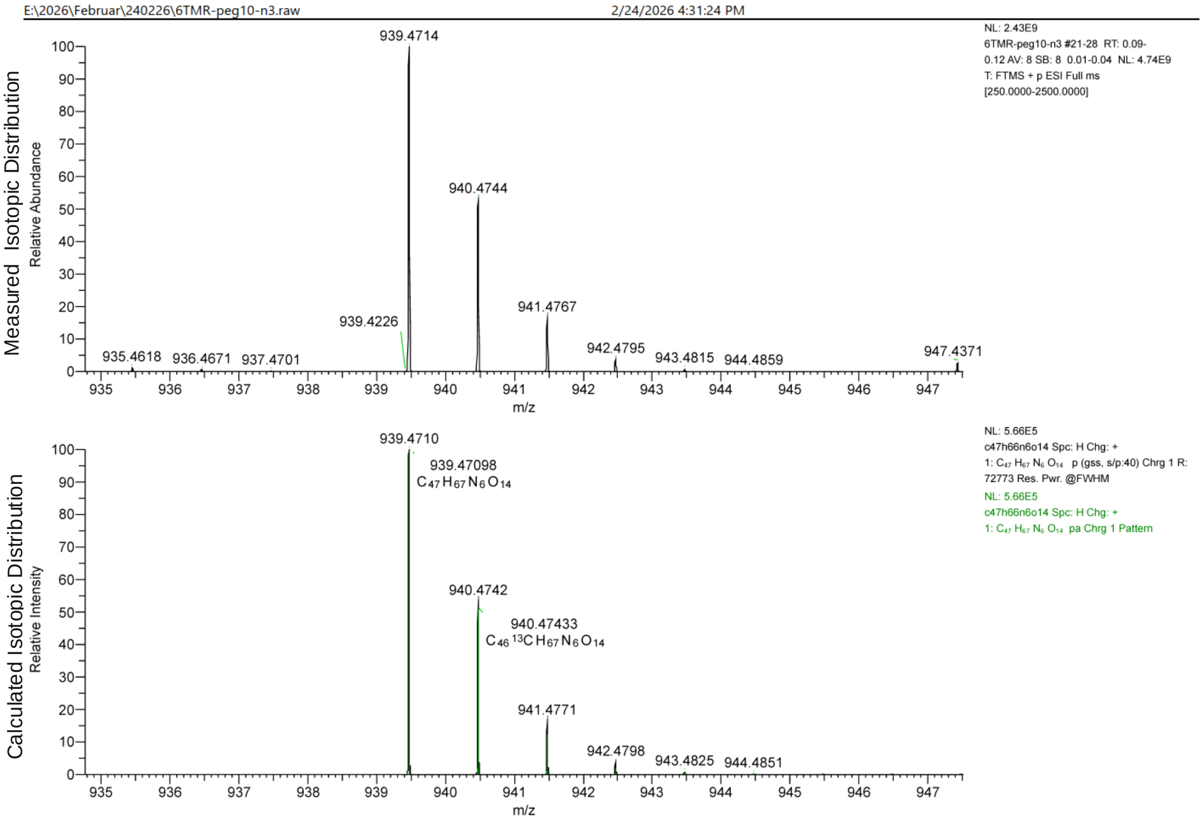

LC/MS:

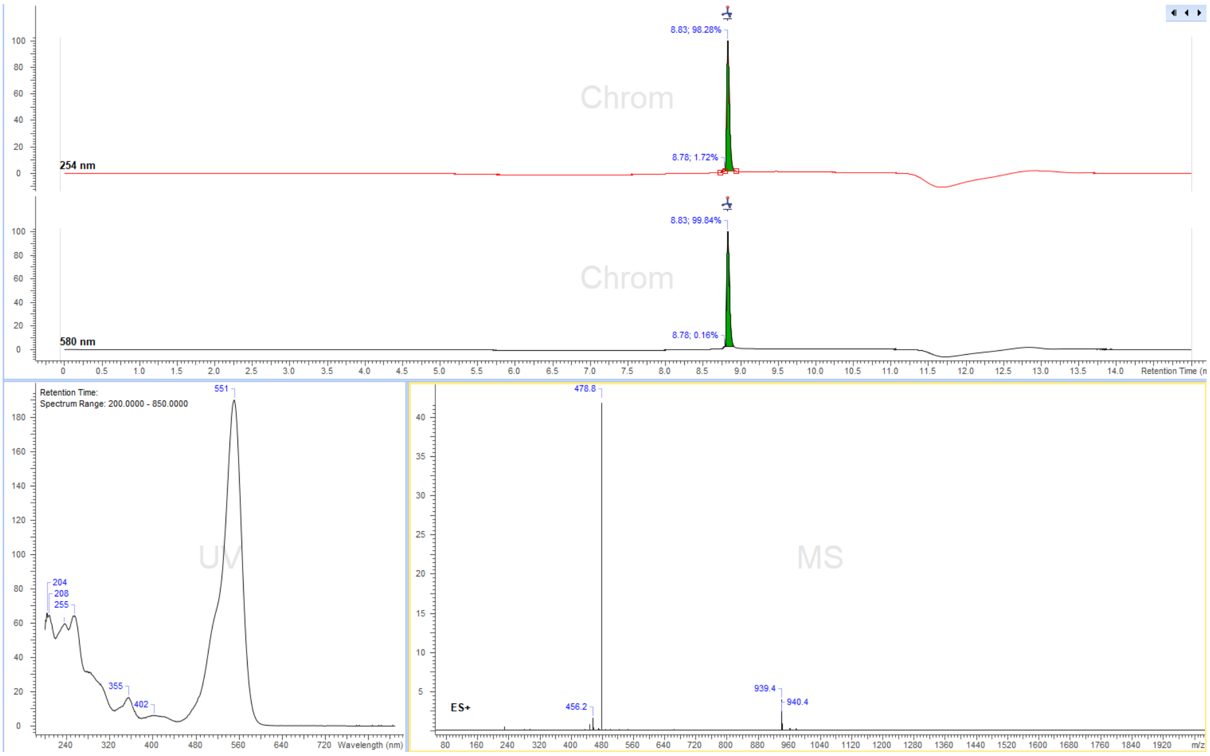

8.83; 98.28%
Chrom
8.78; 1.72%
254 nm
8.83; 99.84%
Chrom
8.78; 0.16%
580 nm
Retention Time (min)
Retention Time:
Spectrum Range: 200.0000 - 850.0000
551
204
208
255
355
402
UV
Wavelength (nm)
478.8
MS
456.2
939.4
940.4
ES+
m/z

Cy3B-PEG$_{10}$-N$_3$

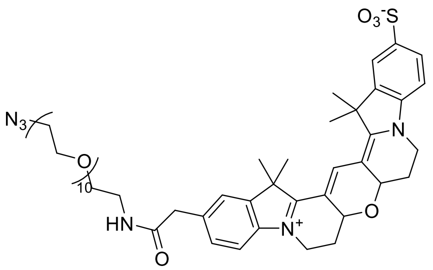

Reaction was performed following general synthesis procedure B from Cy3B-NHS ester (5 mg, Lumiprobe, Cat. Nr. 29320) and NH$_2$-PEG$_{10}$-N$_3$ in dry DMSO. The product was purified by preparative HPLC (preparative column: Eurospher II 100-5 CN Column 250x30 mm, 5 µm, flow rate: 30 mL/min, solvent A: 10 mM ammonium formate (pH 3.5), solvent B: MeOH; room temperature, gradient A:B-2 min 80:20 isocratic, 2-16 min 80:20 to 0:100 gradient, 16-19 min 0:100 isocratic, 19-20 min 0:100 to 80:20 gradient and 20-24 min 80:20 isocratic). Obtained 6.4 mg (79% yield) of a dark pink solid.

$^1$H NMR (400 MHz, CD$_3$OD) d 8.19 (s, 1H, Ar-H), 7.93 – 7.88 (m, 2H, methine-H), 7.51 (d, J = 1.6 Hz, 1H, Ar-H), 7.40 (dd, J = 8.1, 1.6 Hz, 1H, Ar-H), 7.30 (dd, J = 8.2, 2.8 Hz, 2H, Ar-H), 4.69 (dd, J = 11.6, 5.0 Hz, 2H, CH2-bridge), 4.37 (ddd, J = 14.3, 10.4, 4.2 Hz, 2H, CH2-bridge), 3.95 (dtd, J = 17.8, 13.7, 4.1 Hz, 2H, CH2N-ring), 3.68 – 3.61 (m, 40H, CH2 PEG), 3.56 (t, J = 5.5 Hz, 2H, CH2N3), 3.38 (q, J = 5.3 Hz, 4H, CH2NH and CH2O PEG), 2.61 (dd, J = 11.7, 5.1 Hz, 2H, CH2-ring), 2.10 – 1.98 (m, 2H, CH2-ring), 1.80 – 1.77 (m, 12H, 4 x CH3). NH proton not observed (exchange with solvent CD3OD).

ESI-MS, positive mode: m/z = 1069.3 [M]$^+$.

HRMS (ESI) calculated for $C_{53}H_{76}N_6O_{15}S$ [M]$^+$ 1069.5162, found 1069.5167.

Copy of HRMS:

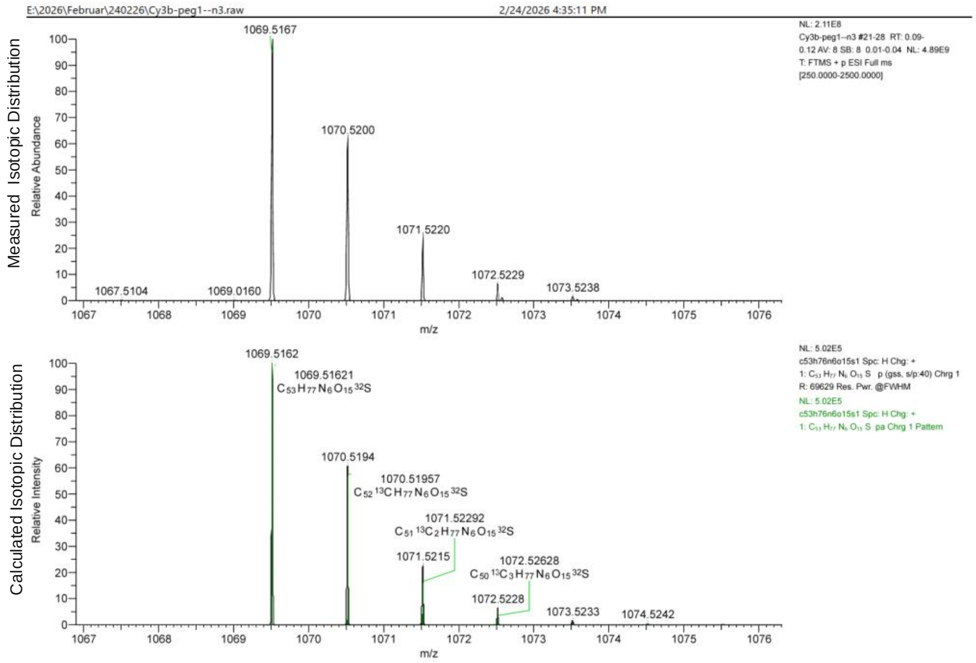

LC/MS:

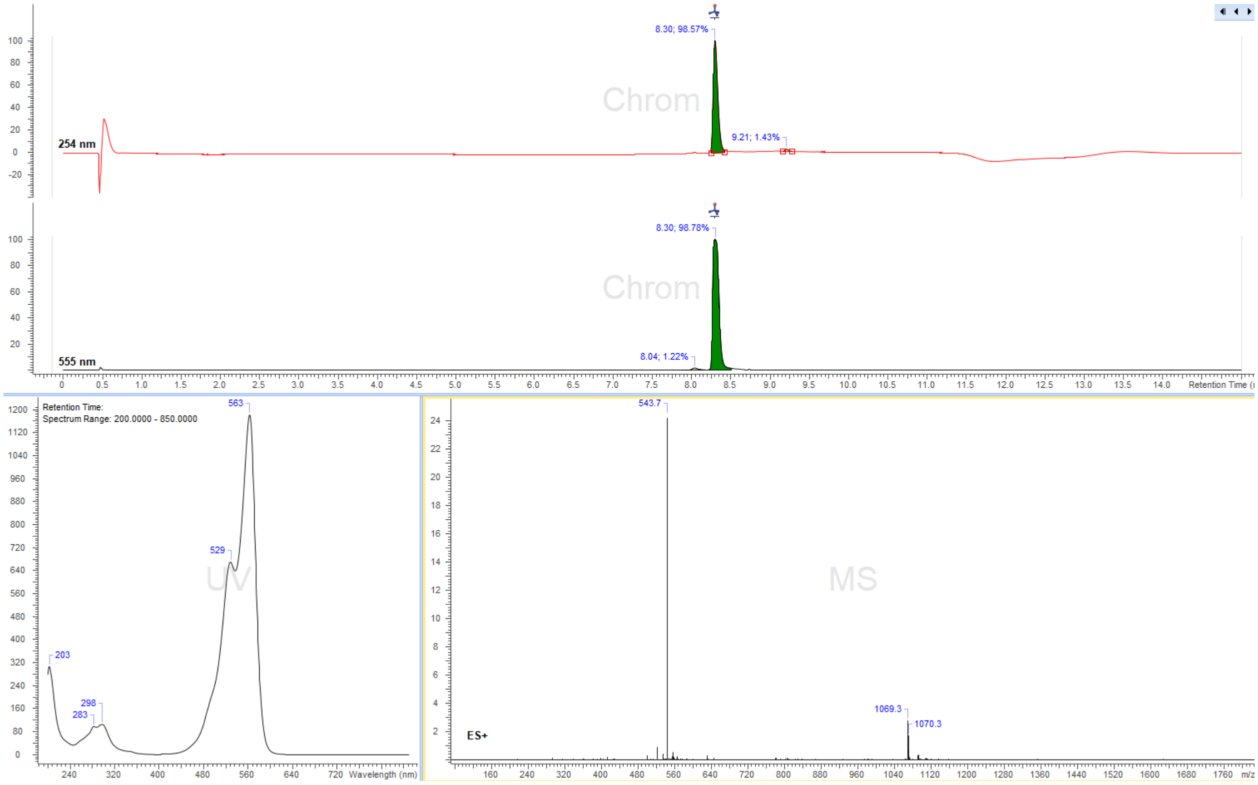
8.30; 98.57%
9.21; 1.43%
254 nm
Chrom
8.30; 98.78%
8.04; 1.22%
555 nm
Retention Time
Retention Time:
Spectrum Range: 200.0000 - 850.0000
563
529
203
298
283
UV
Wavelength (nm)
543.7
1069.3
1070.3
ES+
MS
m/z

*Synthesis of fluorescent cofactors*

4-MeSiR-PEG$_4$-Ado

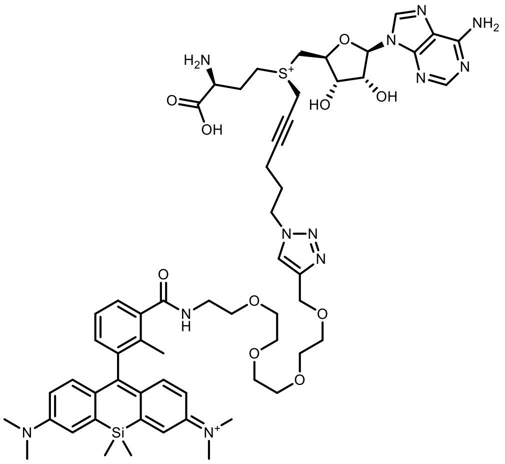

4-MeSiR-PEG$_4$-Ado

Reaction was following general synthesis procedure C from 500µl of 6.7 mM **N$_3$-Ado** and 500µl of 10 mM 4-MeSiR-PEG$_4$-Alkyne. The product was purified by preparative HPLC: Agilent Pursuit 10 C18 10 µm, 250 x 50 mm (Agilent, cat. No. A3002250X500), flow rate: 150 ml/min, solvent A: MeOH, solvent B: 10 mM ammonium formate buffer pH 3.5 at room temperature, gradient A:B - 2 min 30:70 isocratic, 2-20 min 30:70 to 100:0 gradient and 20-22min 100:0 isocratic). Collected fractions of pure product were concentrated to a volume of ~1ml and concentration of the obtained stock solutions was determined with nanodrop based on the absorption at λ= 640nm ($\varepsilon_{640}$ = 150 000 L·mol$^{-1}$·cm$^{-1}$). Obtained 0.8 ml, 2.8 mM of stock solution in 67% yield.

ESI-MS, positive mode: m/z = 581.3 $[M]^{2+}$.

HRMS (ESI) calculated for $C_{58}H_{78}N_{12}O_{10}SSi$ $[M]^{2+}$ 581.2721, found 581.2730.

Copy of HRMS:

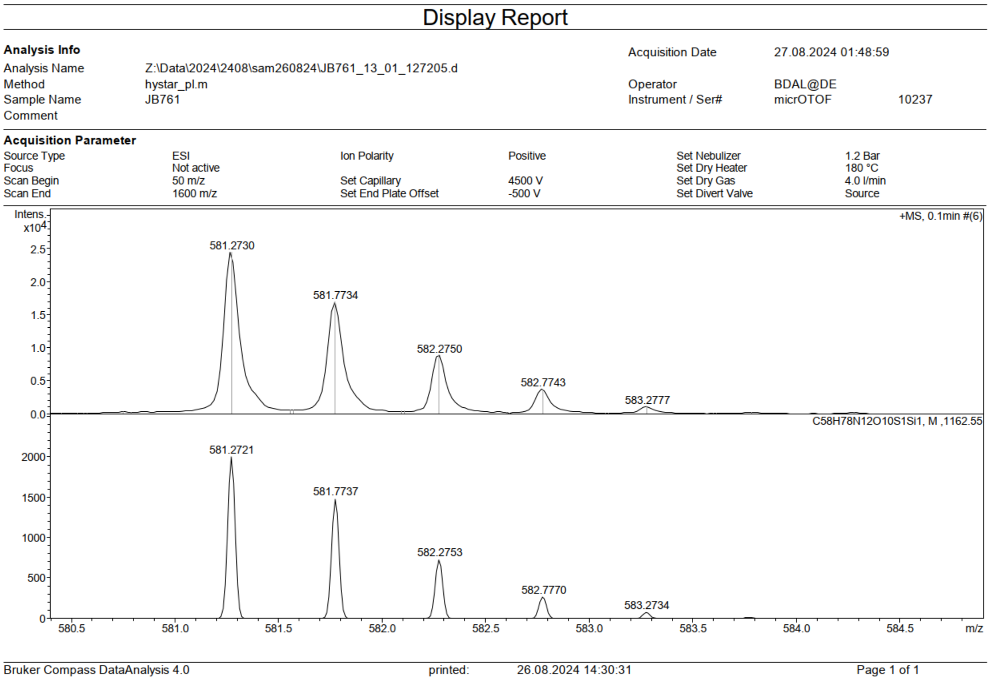

LC/MS:

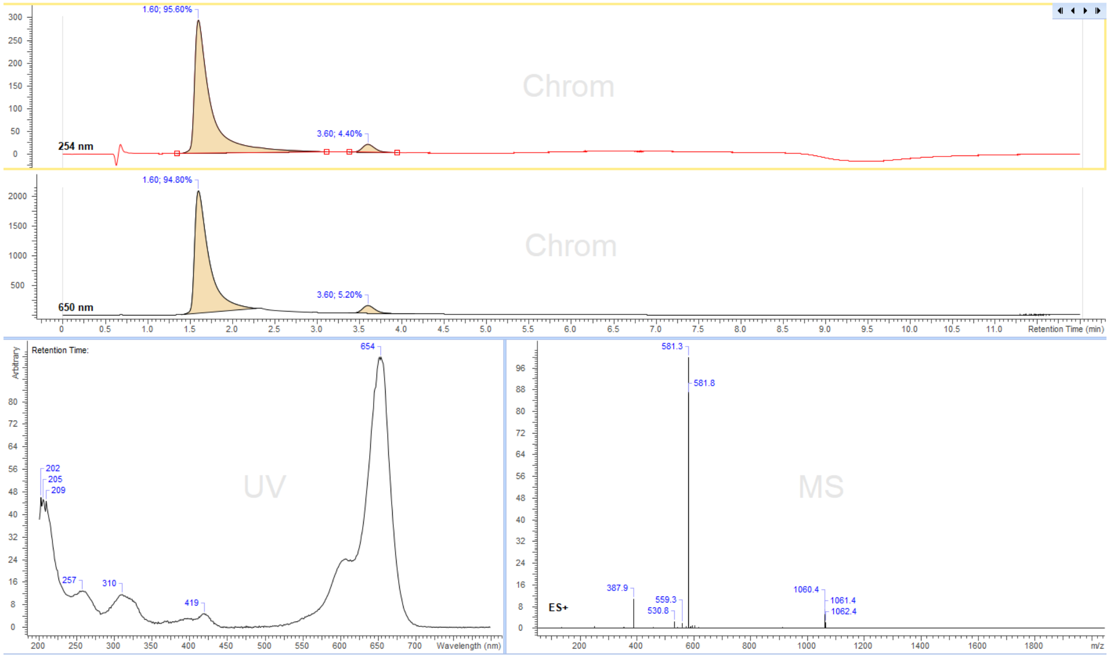

1.60; 95.60%
3.60; 4.40%
254 nm
Chrom
1.60; 94.80%
3.60; 5.20%
650 nm
Retention Time (min)
Retention Time:
Arbitrary
654
202
205
209
257
310
419
UV
Wavelength (nm)
581.3
581.8
387.9
559.3
530.8
1060.4
1061.4
1062.4
ES+
MS
m/z

6-SiR-PEG$_4$-Ado

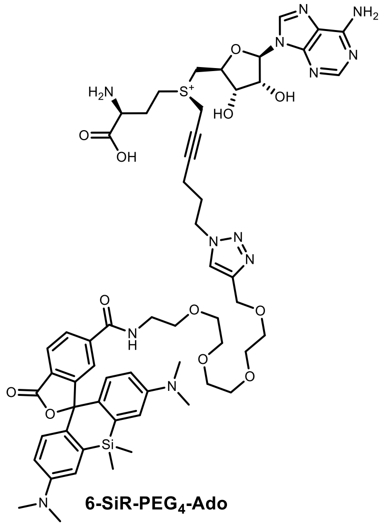

6-SiR-PEG$_4$-Ado

Reaction was performed following general synthesis procedure C from 300 µl of 6.7mM **N$_3$-Ado** and 300µl of 10 mM 6-SiR-PEG$_4$-alkyne (50 µl of formic acid was added to solution of dye for the solubilization purposes, the addition of formic acid and pH change did not have any negative effect towards the completion of the reaction). The product was purified by preparative HPLC: Agilent Pursuit 10 C18 10 µm, 250 x 50 mm (Agilent, cat. No. A3002250X500), flow rate: 150 ml/min, solvent A: MeOH, solvent B: 10 mM ammonium formate buffer pH 3.5 at room temperature, gradient A:B - 2 min 40:60 isocratic, 2-20 min 40:60 to 100:0 gradient and 20-22min 100:0 isocratic). Collected fractions of pure product were concentrated to a volume of ~1ml and concentration of the obtained stock solutions was determined with nanodrop based on the absorption at λ= 640nm ($\varepsilon_{640}$ = 100 000 L·mol$^{-1}$·cm$^{-1}$, dilution was done in PBS + 0.1% SDS in ordert to bring the dye fully in zwiterionic state). Obtained 0.9 ml,1.7 mM of stock solution in 76% yield.

ESI-MS, positive mode: m/z = 1191.5 $[M]^{2+}$.

HRMS (ESI) calculated for $C_{58}H_{75}N_{12}O_{12}SSi$ $[M]^+$ 1191.5112, found 1191.5114.

Copy of HRMS:

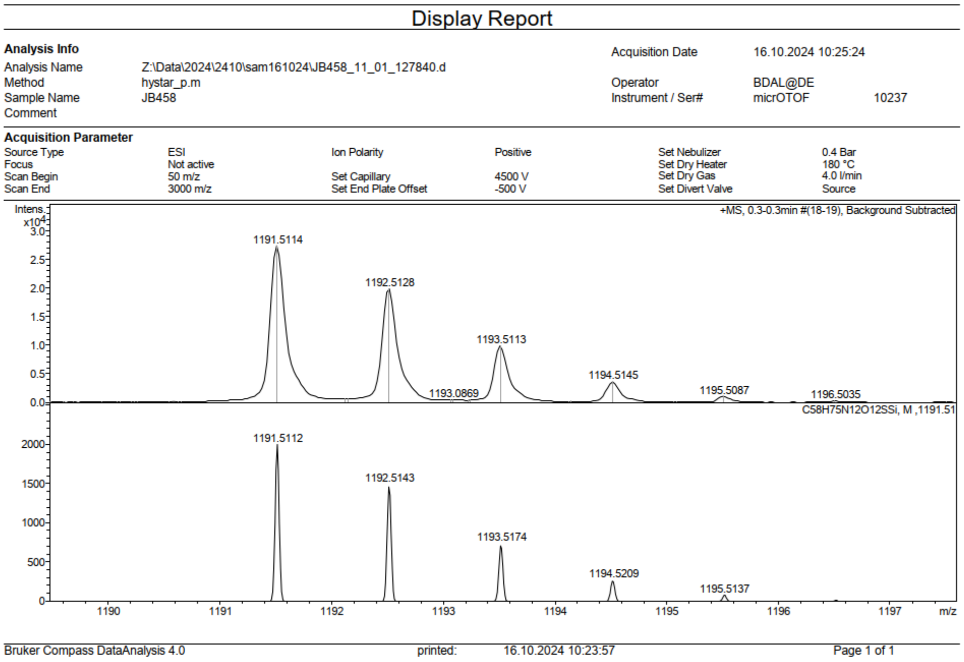

LC/MS:

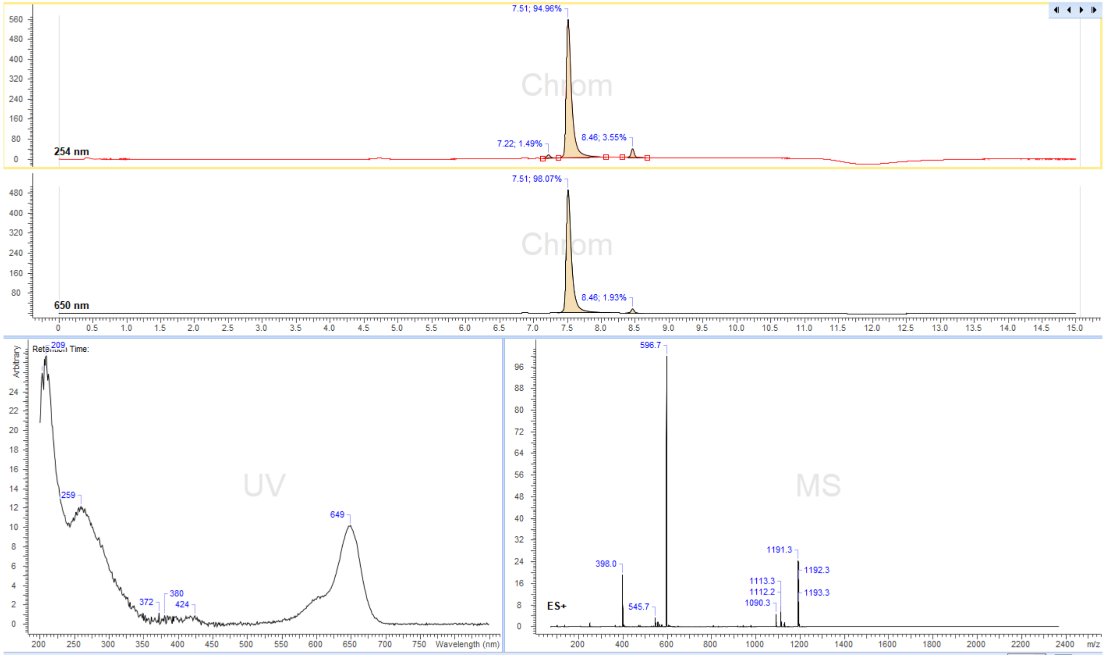

7.51; 94.96%
254 nm
7.22; 1.49%
8.46; 3.55%
7.51; 98.07%
650 nm
8.46; 1.93%
UV
209
259
372
380
424
649
Wavelength (nm)
MS
596.7
398.0
545.7
1191.3
1192.3
1193.3
1113.3
1112.2
1090.3
ES+
m/z

5-HMSiR-PEG$_4$-Ado

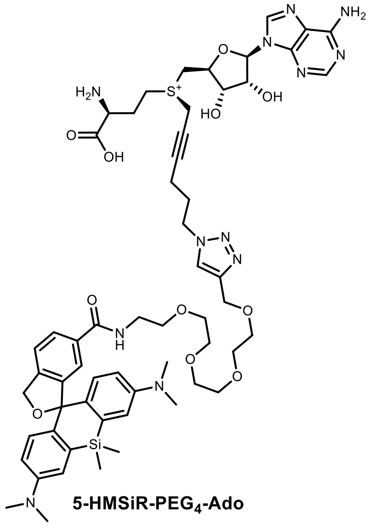

5-HMSiR-PEG$_4$-Ado

Reaction was performed following general synthesis procedure C from 300 µl of 6.7mM **$N_3$-Ado** and 300µl of 10mM 5-HMSiR-PEG$_4$-alkyne (50 µl of formic acid was added to solution of dye for the solubilization purposes, the addition of formic acid and pH change did not have any negative effect towards the completion of the reaction). The product was purified by preparative HPLC: Agilent Pursuit 10 C18 10 µm, 250 x 50 mm (Agilent, cat. No. A3002250X500), flow rate: 150 ml/min, solvent A: MeOH, solvent B: 10 mM ammonium formate buffer pH 3.5 at room temperature, gradient A:B - 2 min 40:60 isocratic, 2-20 min 40:60 to 100:0 gradient and 20-22min 100:0 isocratic). Collected fractions of pure product were concentrated to a volume of ~1ml and concentration of the obtained stock solutions was determined with nanodrop based on the absorption at λ= 640nm ($\varepsilon_{640}$ = 63 000 L·mol$^{-1}$·cm$^{-1}$, dilution was done in PBS + 0.1% SDS in order to bring the dye fully in zwiterionic state). Obtained 0.8 ml, 1.5 mM of stock solution in 60% yield.

ESI-MS, positive mode: m/z = 1177.5 $[M]^+$

HRMS (ESI) calculated for $C_{58}H_{77}N_{12}O_{11}SSi$ $[M]^+$ 1177.5319, found 1177.5323.

Copy of HRMS:

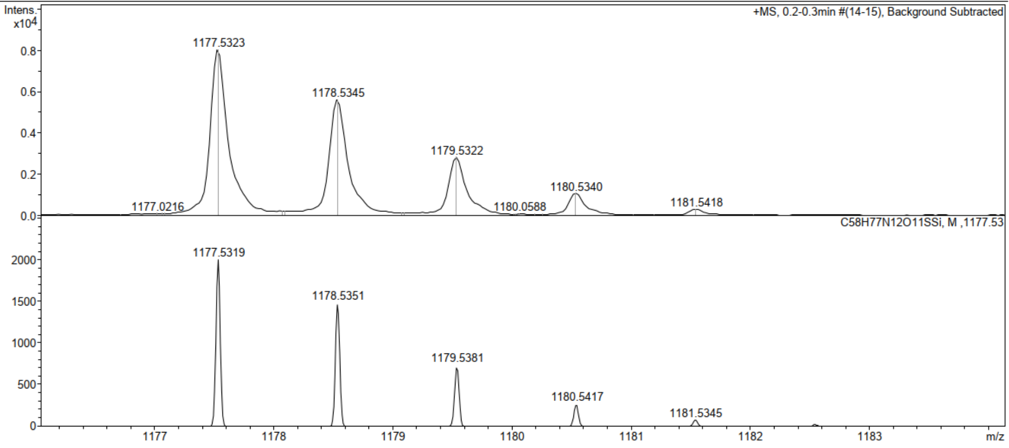

Display Report

**Analysis Info**

| | | | |
|---|---|---|---|
| Analysis Name | Z:\Data\2024\2410\sam161024\JB740c_12_01_127841.d | Acquisition Date | 16.10.2024 10:30:02 |
| Method | hystar_p.m | Operator | BDAL@DE |
| Sample Name | JB740c | Instrument / Ser# | micrOTOF 10237 |
| Comment | | | |

**Acquisition Parameter**

| | | | | | |
|---|---|---|---|---|---|
| Source Type | ESI | Ion Polarity | Positive | Set Nebulizer | 0.4 Bar |
| Focus | Not active | | | Set Dry Heater | 180 °C |
| Scan Begin | 50 m/z | Set Capillary | 4500 V | Set Dry Gas | 4.0 l/min |
| Scan End | 3000 m/z | Set End Plate Offset | -500 V | Set Divert Valve | Source |

Bruker Compass DataAnalysis 4.0    printed: 16.10.2024 10:29:15    Page 1 of 1

LC/MS:

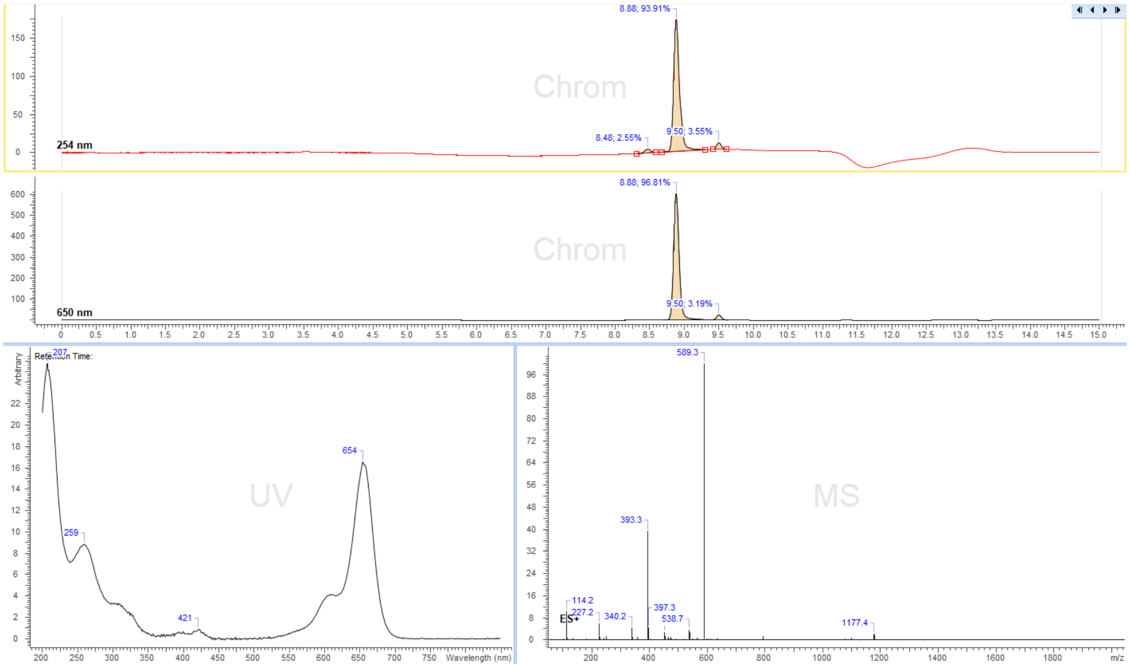

8.88; 93.91%
Chrom
8.48; 2.55%
9.50; 3.55%
254 nm
8.88; 96.81%
Chrom
9.50; 3.19%
650 nm
UV
207
259
421
654
Wavelength (nm)
MS
589.3
393.3
114.2
227.2
340.2
397.3
538.7
1177.4
ES+
m/z

Biotin-$PEG_4$-Ado

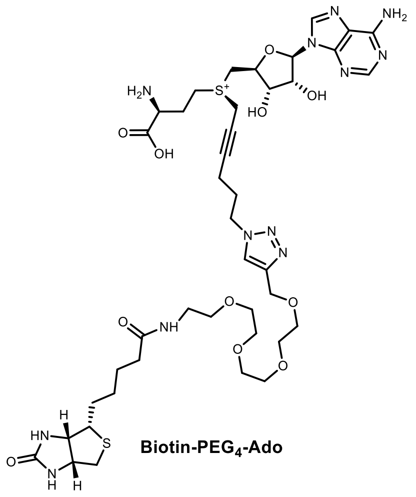

Biotin-$PEG_4$-Ado

Reaction was performed following general synthesis procedure C from 500 µl of 6.7 mM $N_3$-Ado and 200µl of 20 mM Biotin-$PEG_4$-alkyne (Lumiprobe, Cat. Nr.: 2615). The product was purified by preparative HPLC: Agilent Pursuit 10 C18 10 µm, 250 x 50 mm (Agilent, cat. No. A3002250X500), flow rate: 150 ml/min, solvent A: MeOH, solvent B: 10 mM ammonium formate buffer pH 3.5 at room temperature, gradient A:B - 2 min 10:90 isocratic, 2-20 min 10:90 to 100:0 gradient and 20-22min 100:0 isocratic). Collected fractions of pure product were concentrated to a volume of ~1ml and concentration of the obtained stock solutions was determined with nanodrop based on the absorption at λ= 254 nm ($\varepsilon_{640}$ = 15 400 L·mol$^{-1}$·cm$^{-1}$). Obtained 0.9 ml, 2.8 mM of stock solution in 74% yield.

ESI-MS, positive mode: m/z = 963.4 $[M]^+$.

HRMS (ESI) calculated for $C_{41}H_{63}N_{12}O_{11}S_2$ $[M]^+$ 963.4175, found 963.4172.

Copy of HRMS:

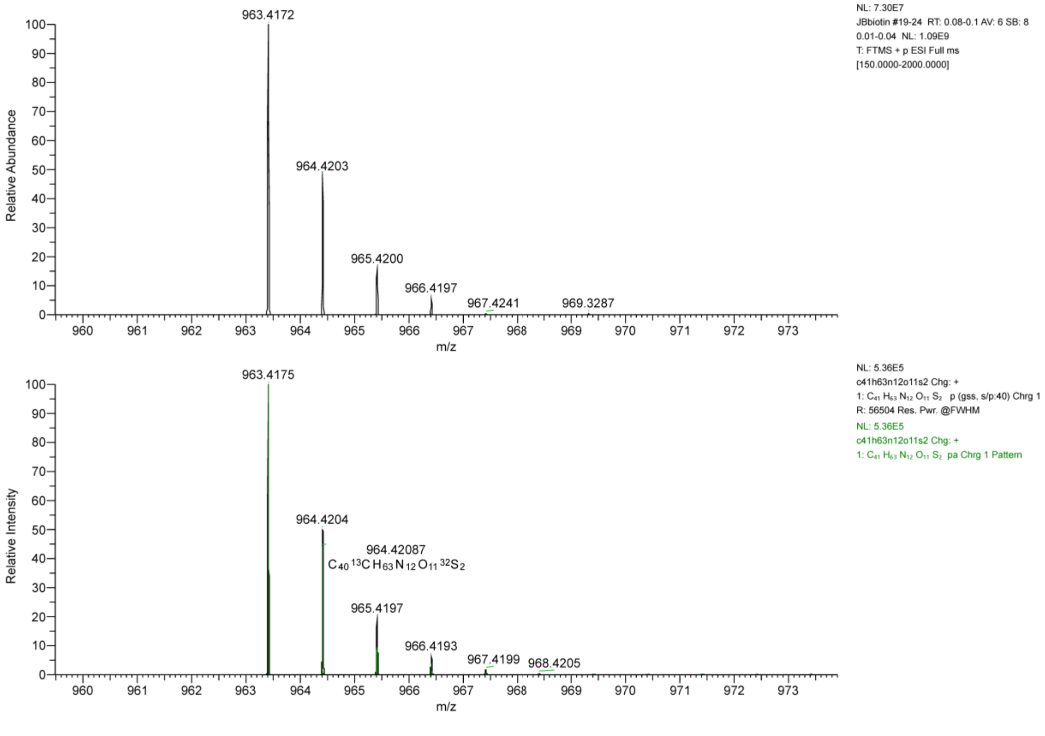

LC/MS:

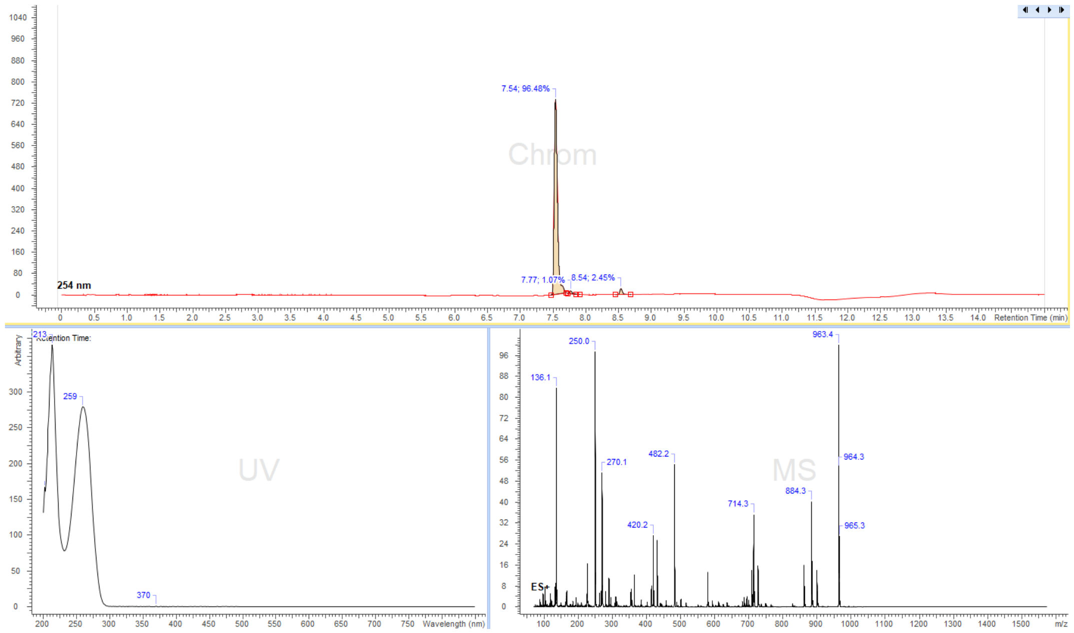
7.54; 96.48%
Chrom
254 nm
7.77; 1.07%
8.54; 2.45%
Retention Time (min)
213
Retention Time:
Arbitrary
259
UV
370
Wavelength (nm)
250.0
136.1
270.1
482.2
420.2
MS
963.4
964.3
884.3
714.3
965.3
ES+
m/z

STAR-RED-Ado

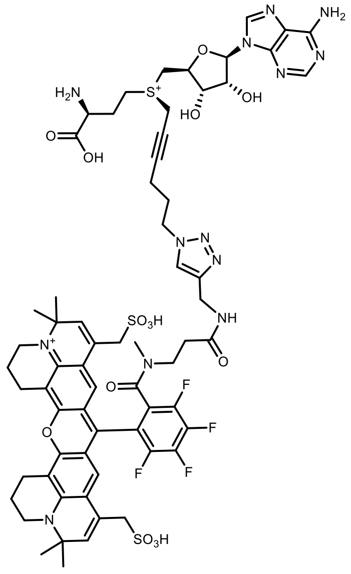

**STAR-RED-Ado**

Reaction was performed following general synthesis procedure C from (in this case ~2eq excess of **$N_3$-Ado** was used) 500 µl of 6.7mM **$N_3$-Ado** and 200µl of 3.8 mM **STAR-RED-alkyne**. The product was purified by preparative HPLC: Agilent 5 Prep-C18, 5 µm, 100 x 50 mm, flow rate: 40 ml/min, solvent A: MeCN, solvent B: 10 mM ammonium formate buffer pH 3.5; room temperature, gradient A:B - 2 min 20:80 isocratic, 2-20 min 20:80 to 100:0 gradient and 20-22 min 100:0 isocratic). Collected fractions of pure product were concentrated to a volume of ~1ml and concentration of the obtained stock solutions was determined with nanodrop based on the absorption at λ= 640 nm ($\varepsilon_{640}$ = 120 000 $L\cdot mol^{-1}\cdot cm^{-1}$). Obtained 0.35 ml, 1.3 mM of stock solution in 59% yield.

ESI-MS, positive mode: m/z =715.7 $[M]^{2+}$

HRMS (ESI) calculated for $C_{65}H_{73}F_4N_{13}O_{14}S_3$ $[M]^{2+}$ 715.7724, found 715.7262

Copy of HRMS:

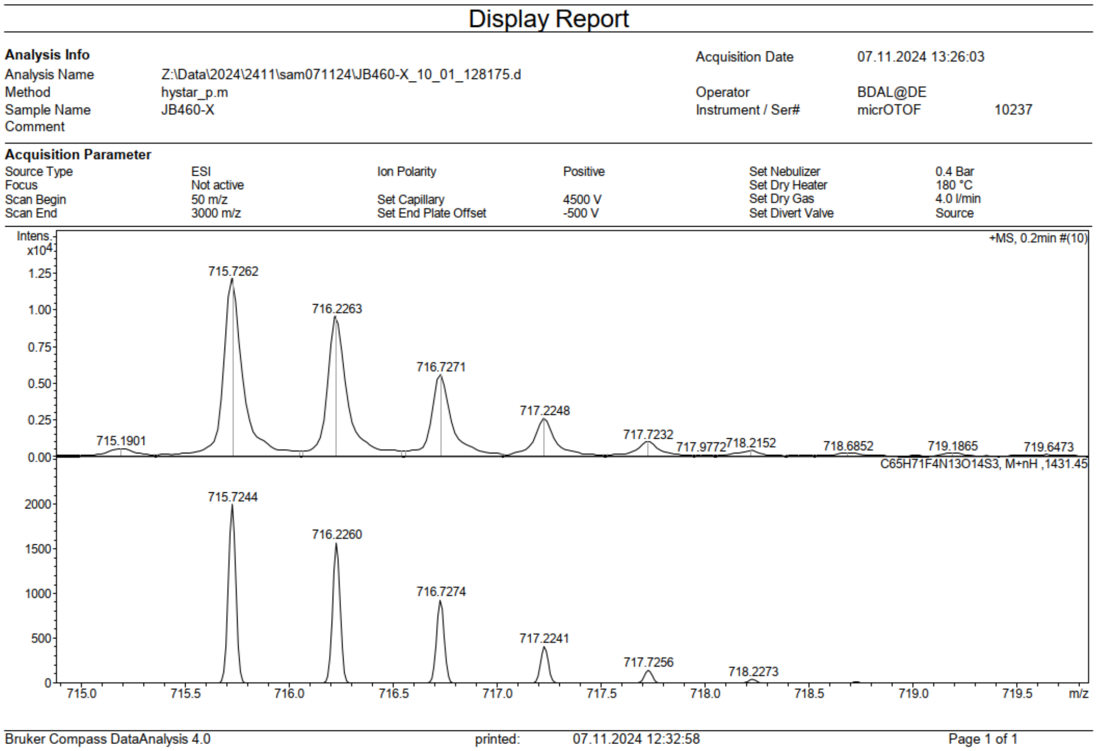

LC/MS:

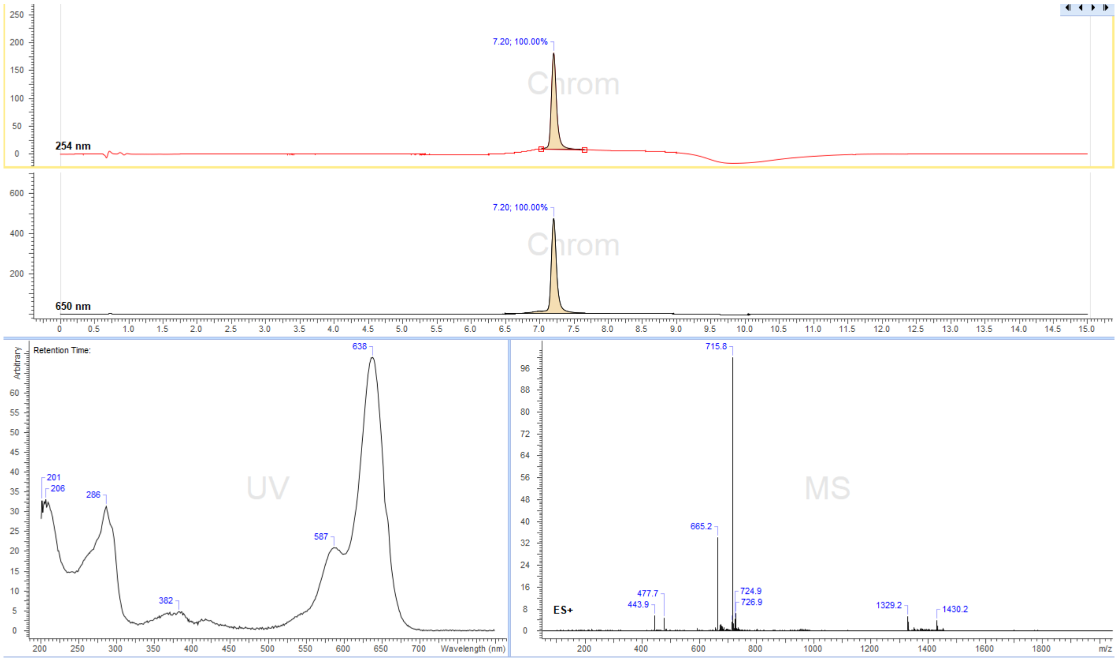

7.20; 100.00%
254 nm
7.20; 100.00%
650 nm
Retention Time:
638
201
206
286
382
587
Wavelength (nm)
715.8
665.2
477.7
443.9
724.9
726.9
1329.2
1430.2
ES+
m/z

STAR-RED-PEG$_4$-Ado

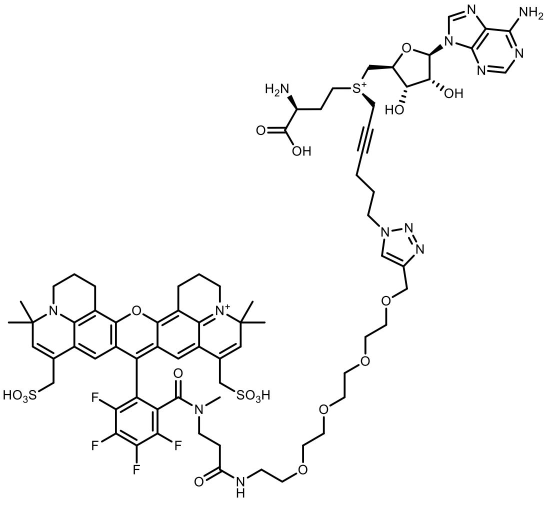

STAR-RED-PEG$_4$-Ado

Reaction was performed following general synthesis procedure C from (in this case ~2eq excess of **$N_3$-Ado** was used ) 500 µl of 6.7mM **$N_3$-Ado** and 200µl of 4.0 mM **STAR-RED-PEG$_4$-alkyne**. The product was purified by preparative HPLC: Agilent 5 Prep-C18, 5 µm, 100 x 50 mm, flow rate: 40 ml/min, solvent A: MeCN, solvent B: 10 mM ammonium formate buffer pH 3.5; room temperature, gradient A:B - 2 min 40:60 isocratic, 2-20 min 40:60 to 100:0 gradient and 20-22 min 100:0 isocratic). Collected fractions of pure product were concentrated to a volume of ~1ml and concentration of the obtained stock solutions was determined with nanodrop based on the absorption at λ= 640 nm ($\varepsilon_{640}$ = 120 000 L·mol$^{-1}$·cm$^{-1}$). Obtained 0.4 ml, 1.2 mM of stock solution in 58% yield.

ESI-MS, positive mode: m/z = 803.8 $[M]^{2+}$

HRMS (ESI) calculated for $C_{73}H_{89}F_4N_{13}O_{18}S_{32}$ $[M]^{2+}$ 803.7768, found 803.7768.

Copy of HRMS:

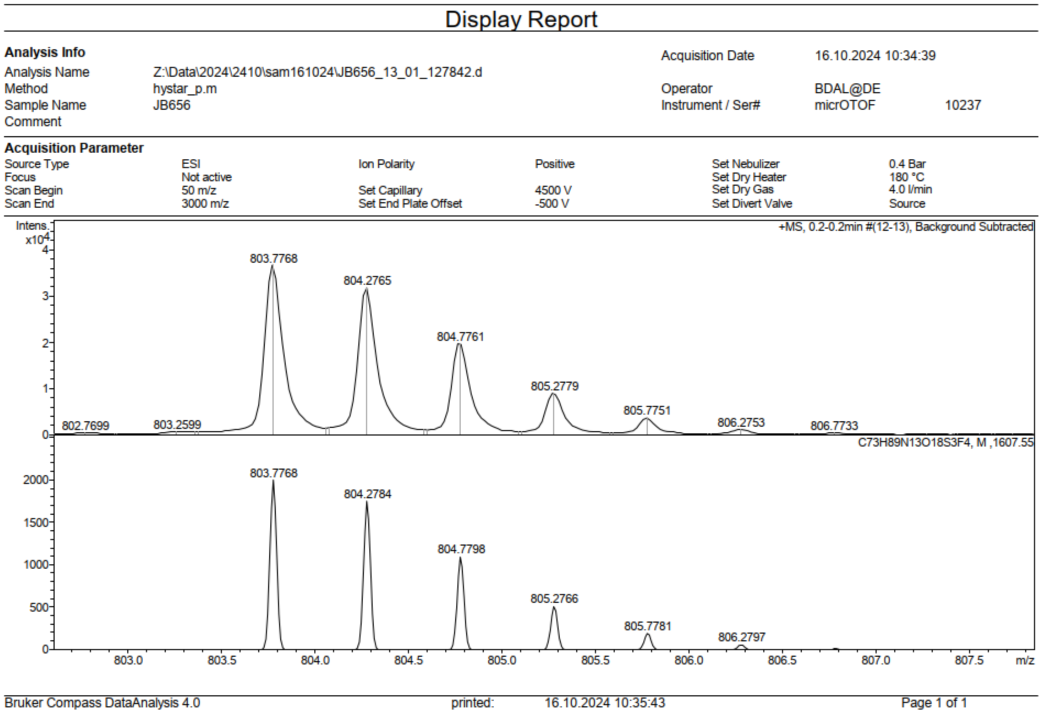

LC/MS:

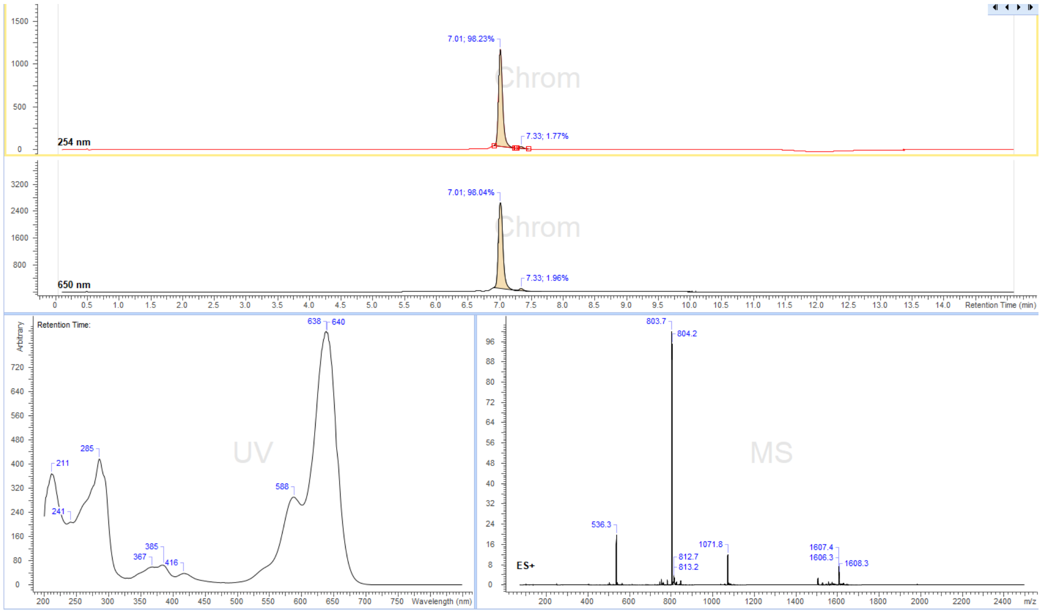
7.01; 98.23%
7.33; 1.77%
254 nm
7.01; 98.04%
7.33; 1.96%
650 nm
Retention Time (min)
Retention Time:
Arbitrary
211
241
285
367
385
416
588
638
640
UV
Wavelength (nm)
803.7
804.2
536.3
812.7
813.2
1071.8
1607.4
1606.3
1608.3
ES+
MS
m/z

5-TMR-$(CH_2)_3$-Ado

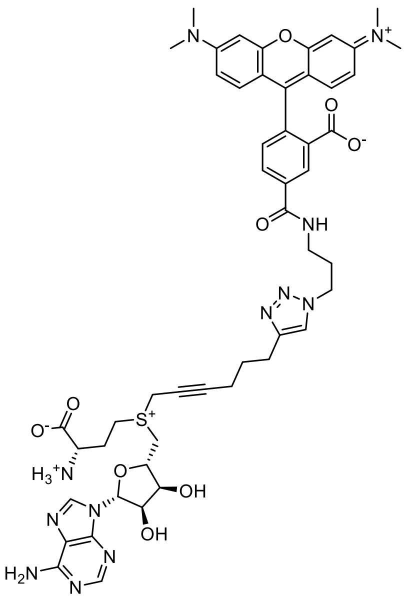

Reaction was performed following general synthesis procedure D from 450µL of 12mM **octadiynyl-Ado** and 800µL of 4mM **5-TMR-$(CH_2)_3$-$N_3$**. The product was purified by preparative HPLC: Eurospher II 100-5 CN Column 250x30 mm, 5 µm (Knauer, Cat. No. 25ME200E2J) column, flow rate: 35 mL/min, solvent A: 10 mM ammonium formate (pH 3.5) and solvent B: MeOH at room temperature, gradient A:B-2 min 80:20 isocratic, 2-12 min 80:20 to 0:100 gradient, 12-15min 0:100 isocratic, 15-16 min 0:100 to 80:20 gradient and 16-20 min isocratic 80:20. Collected fractions of pure product were concentrated and concentration of the obtained stock solution was determined with nanodrop based on the absorption at λ= 555nm ($\varepsilon_{555}$ =83600 $L{\cdot}mol^{-1}{\cdot}cm^{-1}$). Obtained 1.46 mL, 0.63 mM of stock solution in 29% yield.

ESI-MS, positive mode: m/z = 1001.3 $[M]^+$ and 501.20 $[M]^{2+}$.

HRMS (ESI) calculated for $C_{50}H_{57}N_{12}O_9S$ $[M]^+$ 1001.4087, found 1001.4090.

Copy of HRMS:

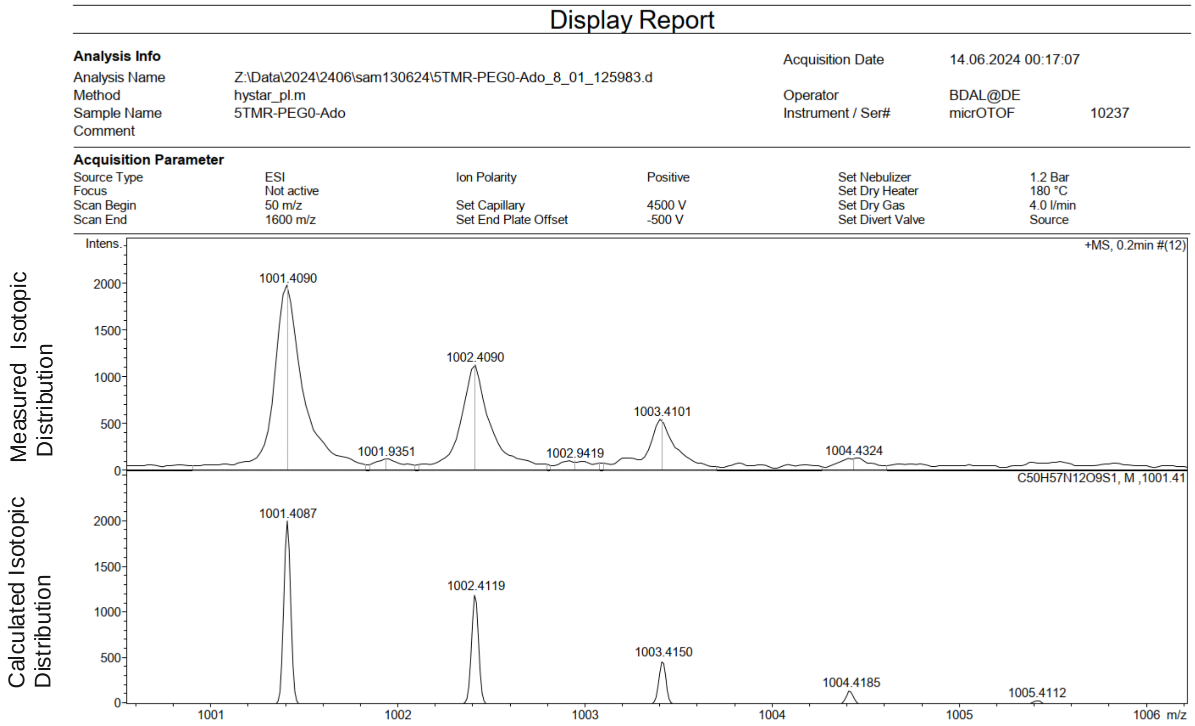

LC/MS:

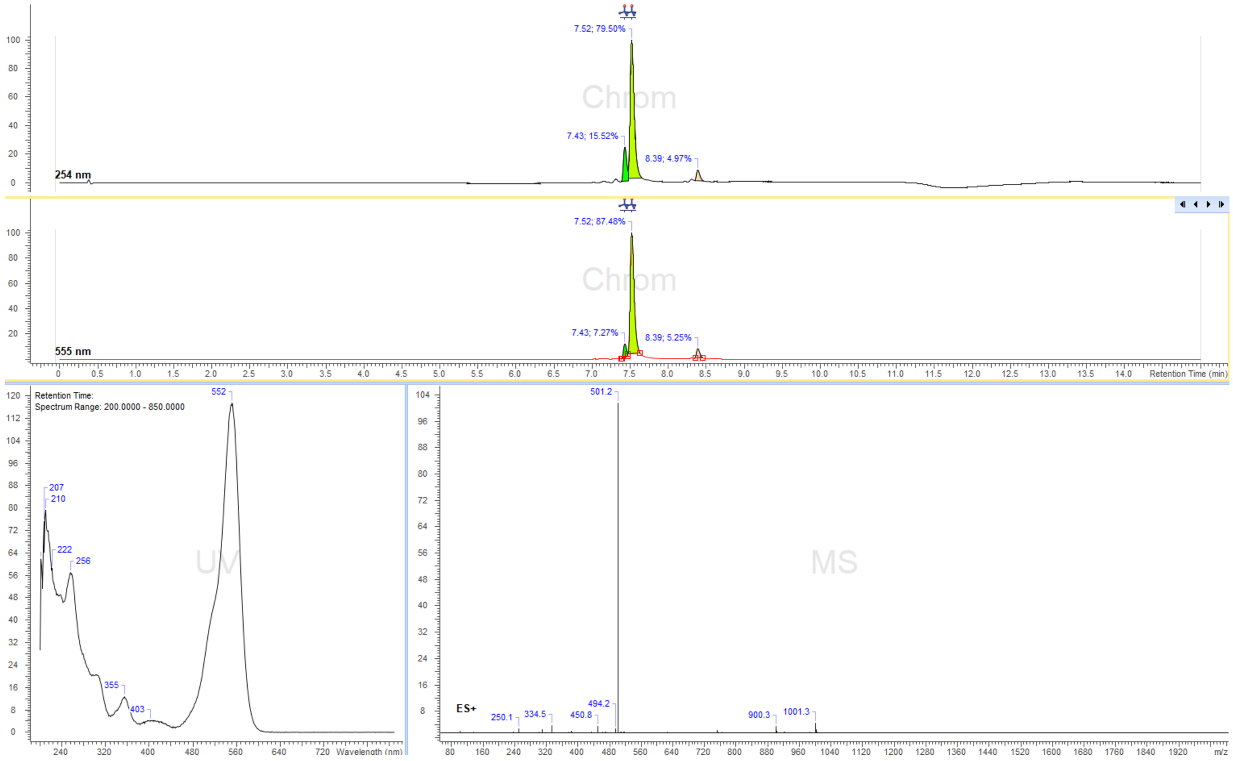

7.52; 79.50%
7.43; 15.52%
8.39; 4.97%
254 nm
7.52; 87.48%
7.43; 7.27%
8.39; 5.25%
555 nm
Retention Time (min)
Retention Time:
Spectrum Range: 200.0000 - 850.0000
552
207
210
222
256
355
403
Wavelength (nm)
ES+
501.2
494.2
450.8
334.5
250.1
900.3
1001.3
m/z

6-TMR-$(CH_2)_3$-Ado

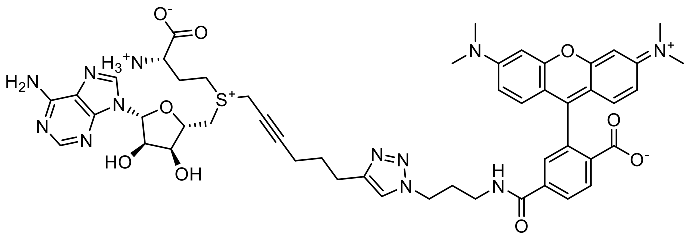

Reaction was performed following general synthesis procedure D from 450µL of 12mM **octadiynyl-Ado** and 650µL of 4.4mM **5-TMR-$(CH_2)_3$-$N_3$**. The product was purified by preparative HPLC: Eurospher II 100-5 CN Column 250x30 mm, 5 µm (Knauer, Cat. No. 25ME200E2J) column, flow rate: 35 mL/min, solvent A: 10 mM ammonium formate (pH 3.5) and solvent B: MeOH at room temperature, gradient A:B-2 min 80:20 isocratic, 2-12 min 80:20 to 0:100 gradient, 12-15min 0:100 isocratic, 15-16 min 0:100 to 80:20 gradient and 16-20 min isocratic 80:20. Collected fractions of pure product were concentrated and concentration of the obtained stock solution was determined with nanodrop based on the absorption at λ= 555nm ($\varepsilon_{555}$ =86600 $L\cdot mol^{-1}\cdot cm^{-1}$). Obtained 1.63 mL, 1.68 mM of stock solution in 75% yield.

ESI-MS, positive mode: m/z = 501.20 $[M]^{2+}$.

HRMS (ESI) calculated for $C_{50}H_{57}N_{12}O_9S$ $[M]^+$ 1001.4087, found 1001.4084.

Copy of HRMS:

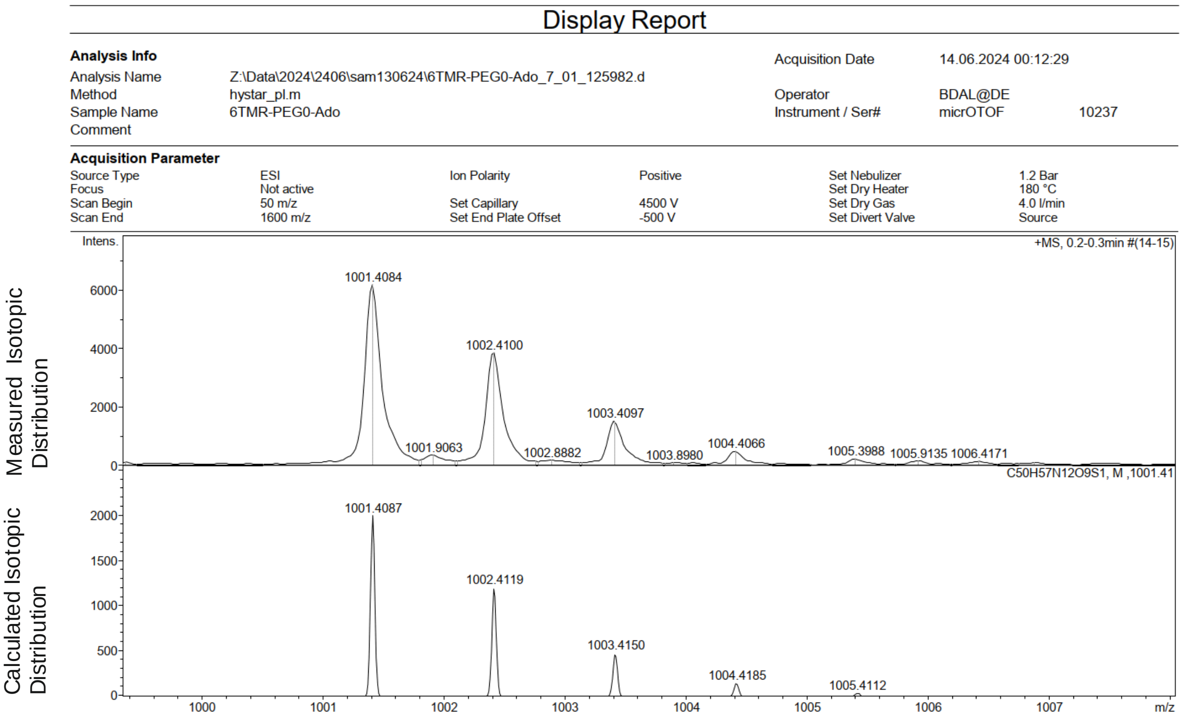

LC/MS:

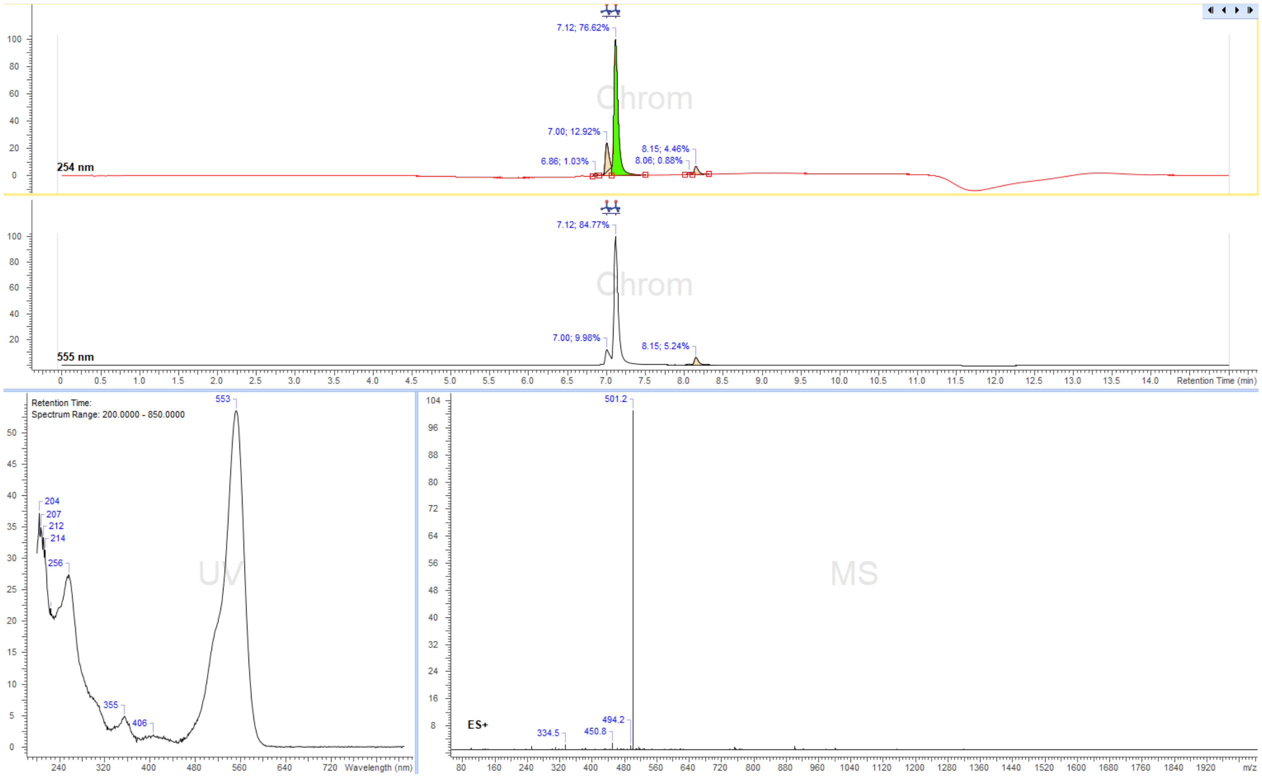

7.12; 76.62%
7.00; 12.92%
6.86; 1.03%
8.15; 4.46%
8.06; 0.88%
254 nm
7.12; 84.77%
7.00; 9.98%
8.15; 5.24%
555 nm
Retention Time (min)
Retention Time:
Spectrum Range: 200.0000 - 850.0000
553
204
207
212
214
256
355
406
Wavelength (nm)
501.2
494.2
450.8
334.5
ES+
m/z

5-TMR-PEG$_3$-Ado

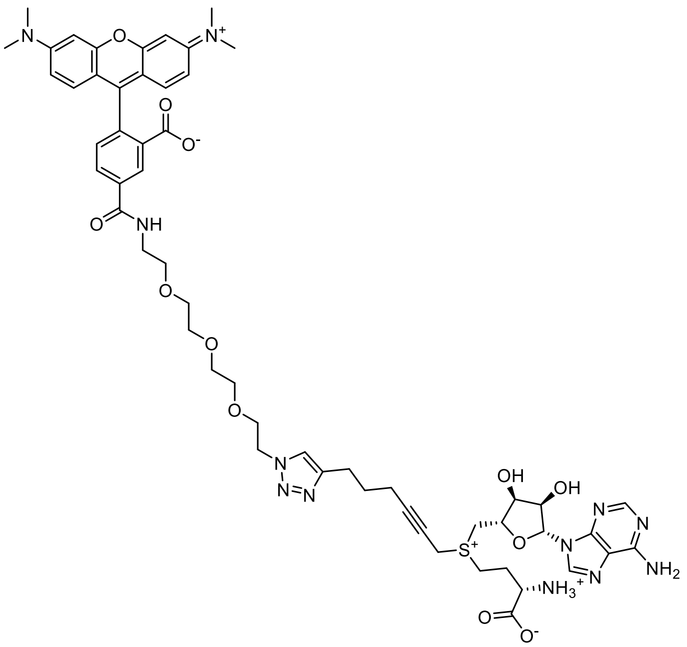

Reaction was performed following general synthesis procedure D from 500 µL of 10.14 mM **octadiynyl-Ado**, 500 µL of 7 mM **5-TMR-PEG$_3$-N$_3$** and 500 µL copper (I) solution. The product was purified by preparative HPLC: Eurospher II 100-5 CN Column 250x30 mm, 5 µm (Knauer, Cat. No. 25ME200E2J) column, flow rate: 35 mL/min, solvent A: 10 mM ammonium formate (pH 3.5) and solvent B: MeOH at room temperature, gradient A:B-2 min 80:20 isocratic, 2-12 min 80:20 to 0:100 gradient, 12-15min 0:100 isocratic, 15-16 min 0:100 to 80:20 gradient and 16-20 min isocratic 80:20. Collected fractions of pure product were concentrated and concentration of the obtained stock solution was determined with nanodrop based on the absorption at λ=555nm ($\varepsilon_{555}$=83600 L·mol$^{-1}$·cm$^{-1}$). Obtained 0.97 mL, 2.17 mM of stock solution in 59% yield.

ESI-MS, positive mode: m/z = 560.3 $[M]^{2+}$.

HRMS (ESI) calculated for $C_{55}H_{67}N_{12}O_{12}S$ 1119.4717 $[M]^+$, found 1119.4700.

Copy of HRMS:

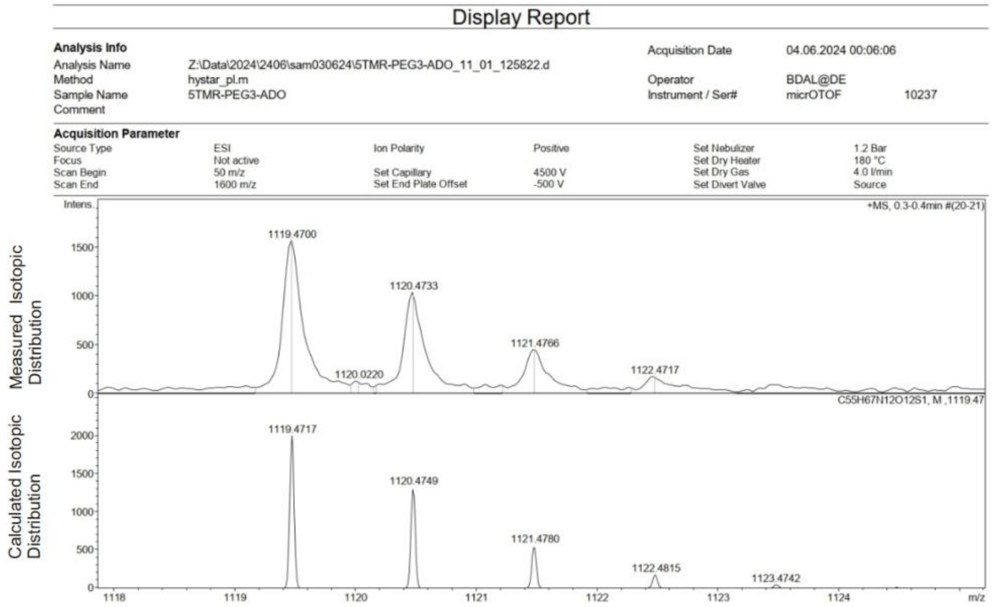

LC/MS:

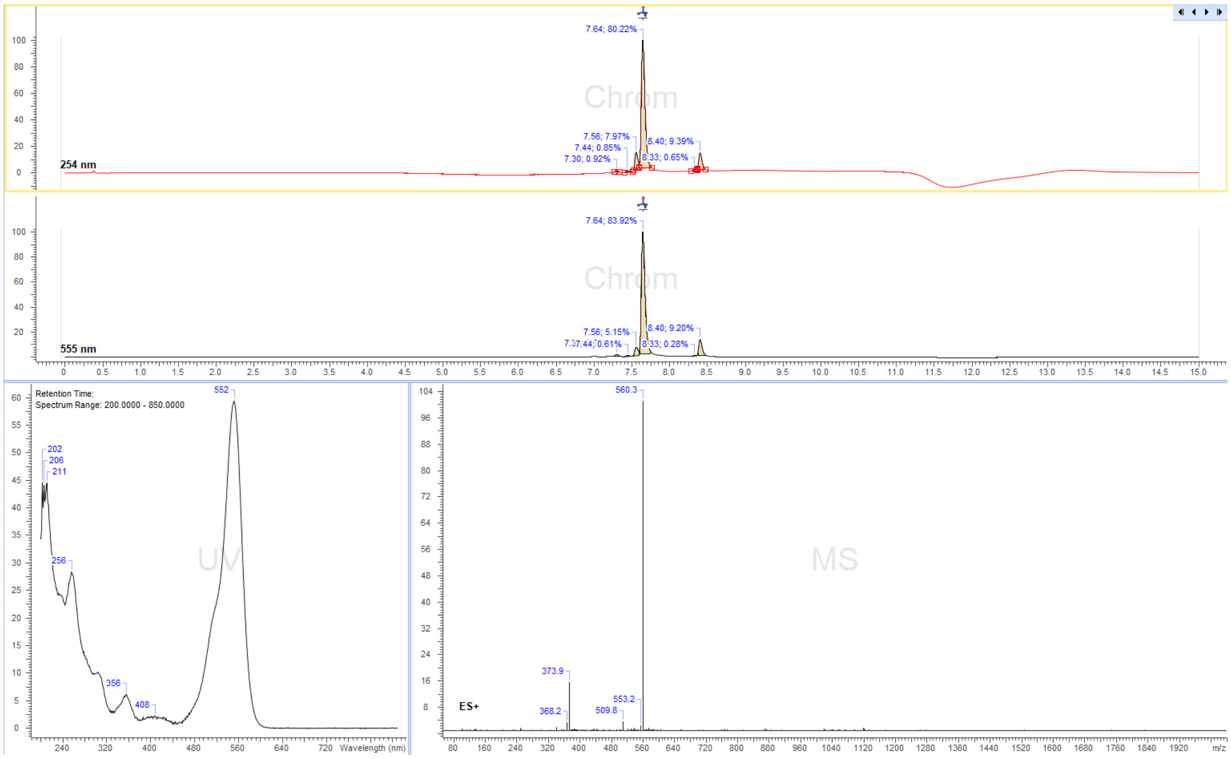
7.64; 80.22%
7.56; 7.97%
7.44; 0.85%
7.30; 0.92%
8.40; 9.39%
8.33; 0.65%
254 nm
7.64; 83.92%
7.56; 5.15%
8.40; 9.20%
8.33; 0.28%
555 nm
Retention Time:
Spectrum Range: 200.0000 - 850.0000
552
202
206
211
256
356
408
Wavelength (nm)
560.3
373.9
553.2
509.8
368.2
ES+
m/z

6-TMR-PEG$_3$-Ado

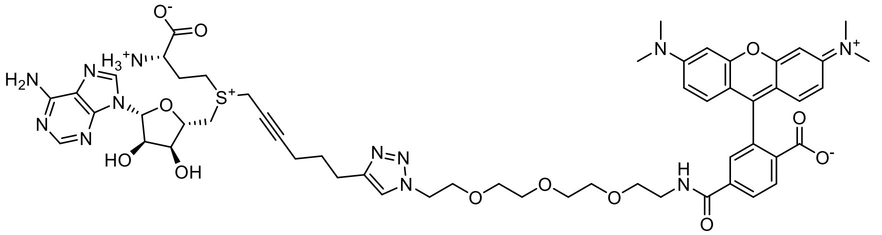

Reaction was performed following general synthesis procedure D from 500 µL of 10.14 mM **octadiynyl-Ado**, 500 µL of 6.3 mM **6-TMR-PEG$_3$-N$_3$** and 500 µL copper (I) solution. The product was purified by preparative HPLC: Eurospher II 100-5 CN Column 250x30 mm, 5 µm (Knauer, Cat. No. 25ME200E2J) column, flow rate: 35 mL/min, solvent A: 10 mM ammonium formate (pH=3.5) and solvent B: MeOH at room temperature, gradient A:B-2 min 80:20 isocratic, 2-12 min 80:20 to 0:100 gradient, 12-15min 0:100 isocratic, 15-16 min 0:100 to 80:20 gradient and 16-20 min isocratic 80:20. Collected fractions of pure product were concentrated and concentration of the obtained stock solution was determined with nanodrop based on the absorption at λ=555nm ($\varepsilon_{555}$=86600 L·mol$^{-1}$·cm$^{-1}$). Obtained 0.88 mL, 2.27 mM of stock solution in 63% yield.

ESI-MS, positive mode: m/z = 560.3 $[M]^{2+}$.

HRMS (ESI) calculated for $C_{55}H_{67}N_{12}O_{12}S$ $[M]^+$ 1119.4717, found 1119.4700.

Copy of HRMS:

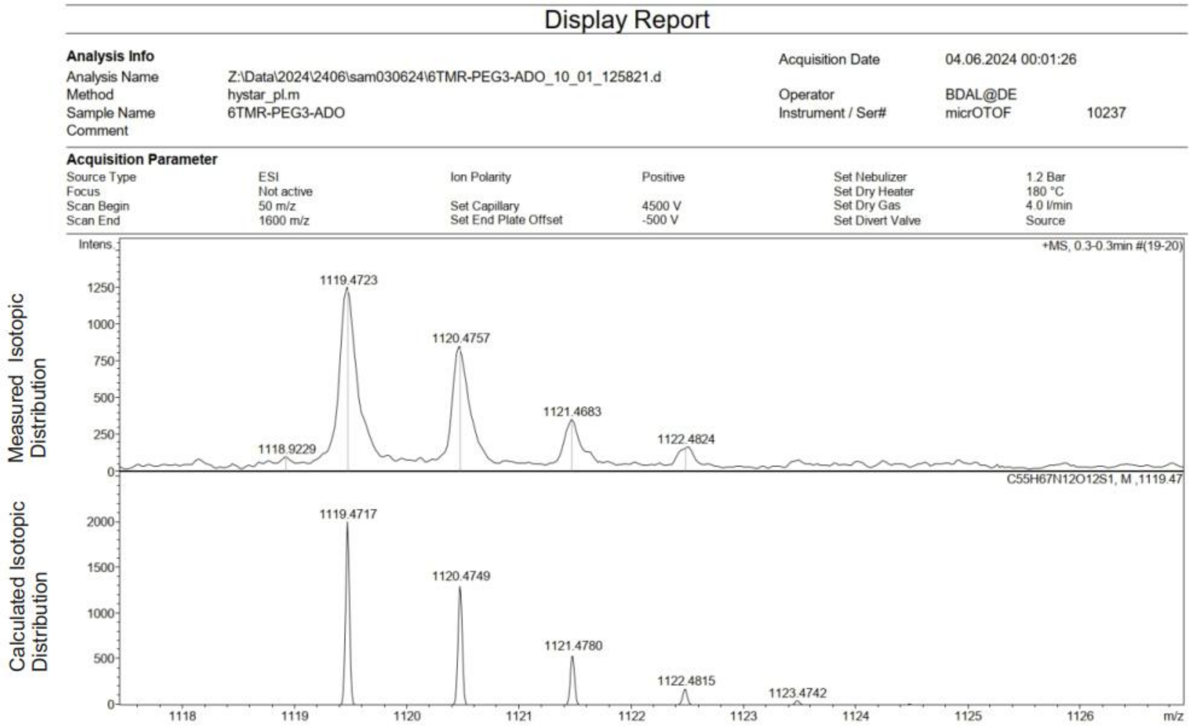

LC/MS:

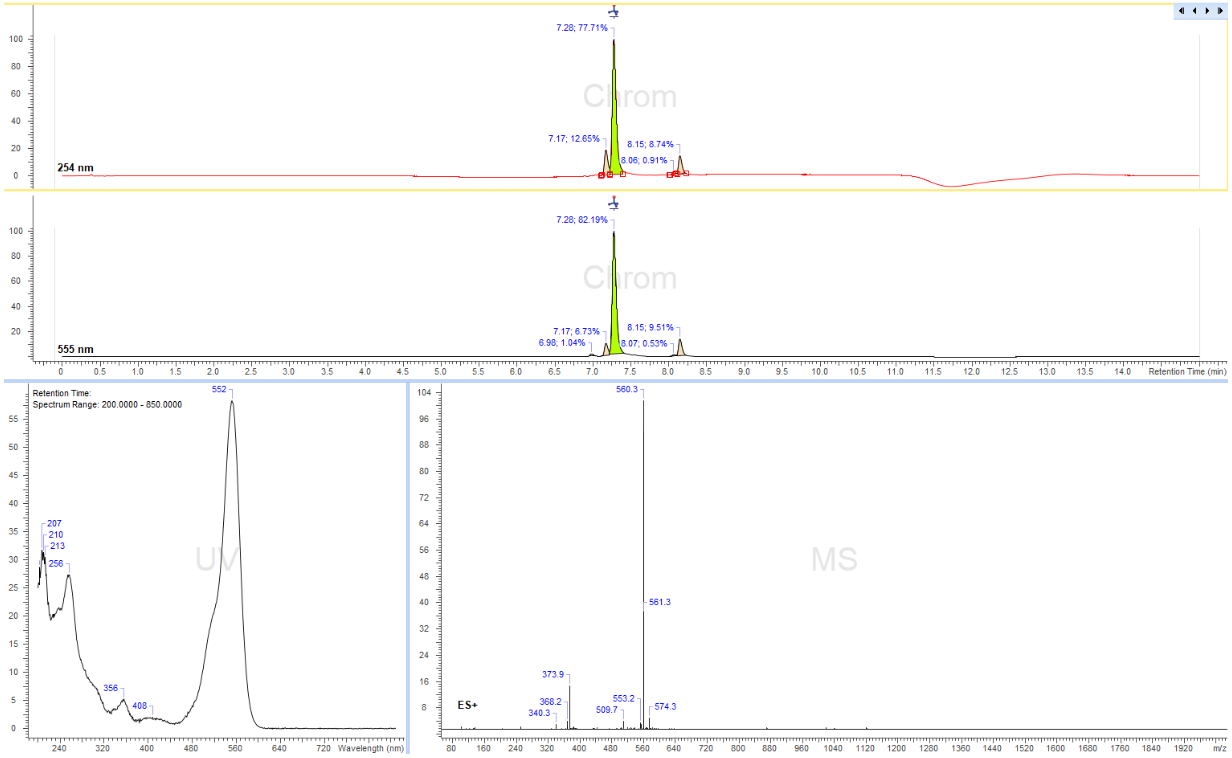

7.28; 77.71%
7.17; 12.65%
8.15; 8.74%
8.06; 0.91%
254 nm
7.28; 82.19%
7.17; 6.73%
6.98; 1.04%
8.15; 9.51%
8.07; 0.53%
555 nm
Retention Time (min)
Retention Time:
Spectrum Range: 200.0000 - 850.0000
552
207
210
213
256
356
408
Wavelength (nm)
560.3
561.3
373.9
368.2
340.3
553.2
509.7
574.3
ES+
m/z

Cy3B-PEG$_3$-Ado

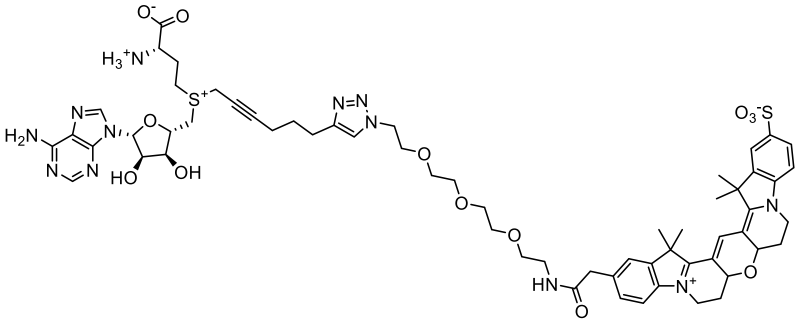

Reaction was performed following general synthesis procedure D from 495 µL of 8.11 mM **octadiynyl-Ado**, 500 µL of 6.5 mM **Cy3B-PEG$_3$-N$_3$** and 500 µL copper (I) solution. The product was purified by preparative HPLC: Eurospher II 100-5 CN Column 250x30 mm, 5 µm (Knauer, Cat. No. 25ME200E2J) column, flow rate: 35 mL/min, solvent A: 10 mM ammonium formate (pH=3.5) and solvent B: MeOH at room temperature, gradient A:B-2 min 80:20 isocratic, 2-12 min 80:20 to 0:100 gradient, 12-15 min 0:100 isocratic, 15-16 min 0:100 to 80:20 gradient and 16-20 min isocratic 80:20. Collected fractions of pure product were concentrated and concentration of the obtained stock solution was determined with nanodrop based on the absorption at λ=559 nm ($\varepsilon_{559}$=121000 L·mol$^{-1}$·cm$^{-1}$). Obtained 0.93 mL, 2.65 mM of stock solution in 75% yield.

ESI-MS, positive mode: m/z= 1249.5 $[M]^+$.

HRMS (ESI) calculated for $C_{61}H_{77}N_{12}O_{13}S$ $[M]^+$ 1249.5169, found 1249.5142.

Copy of HRMS:

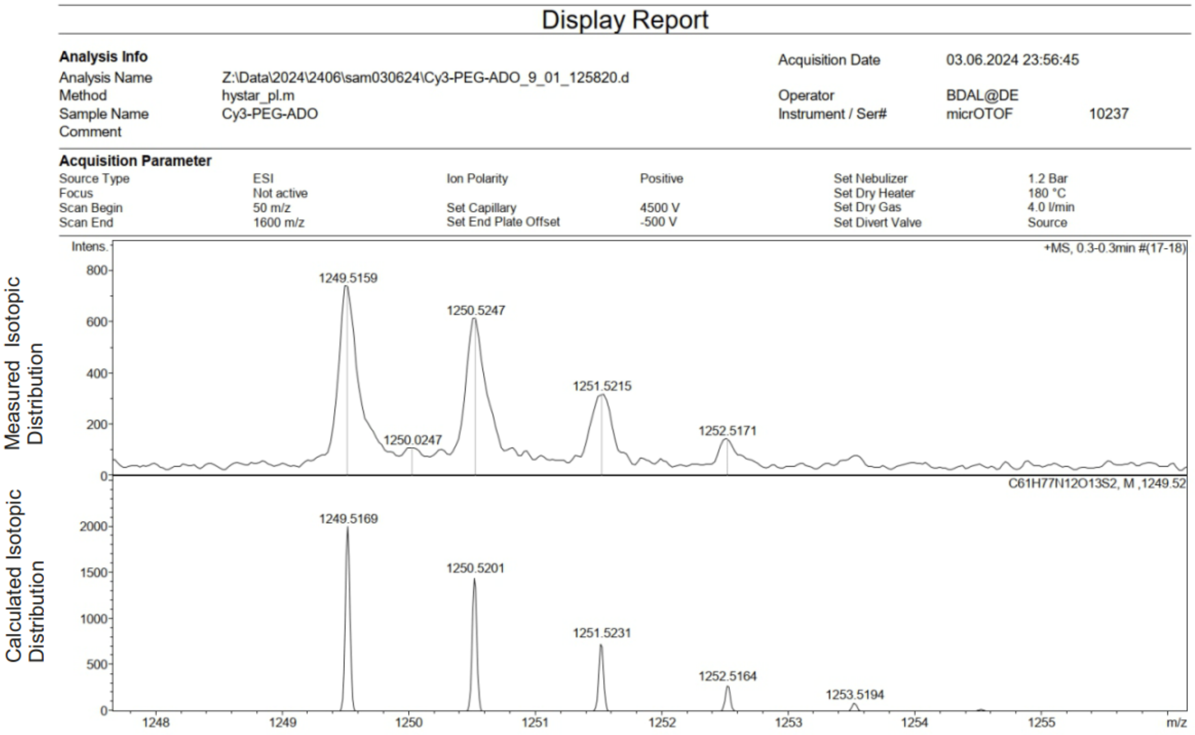

LC/MS:

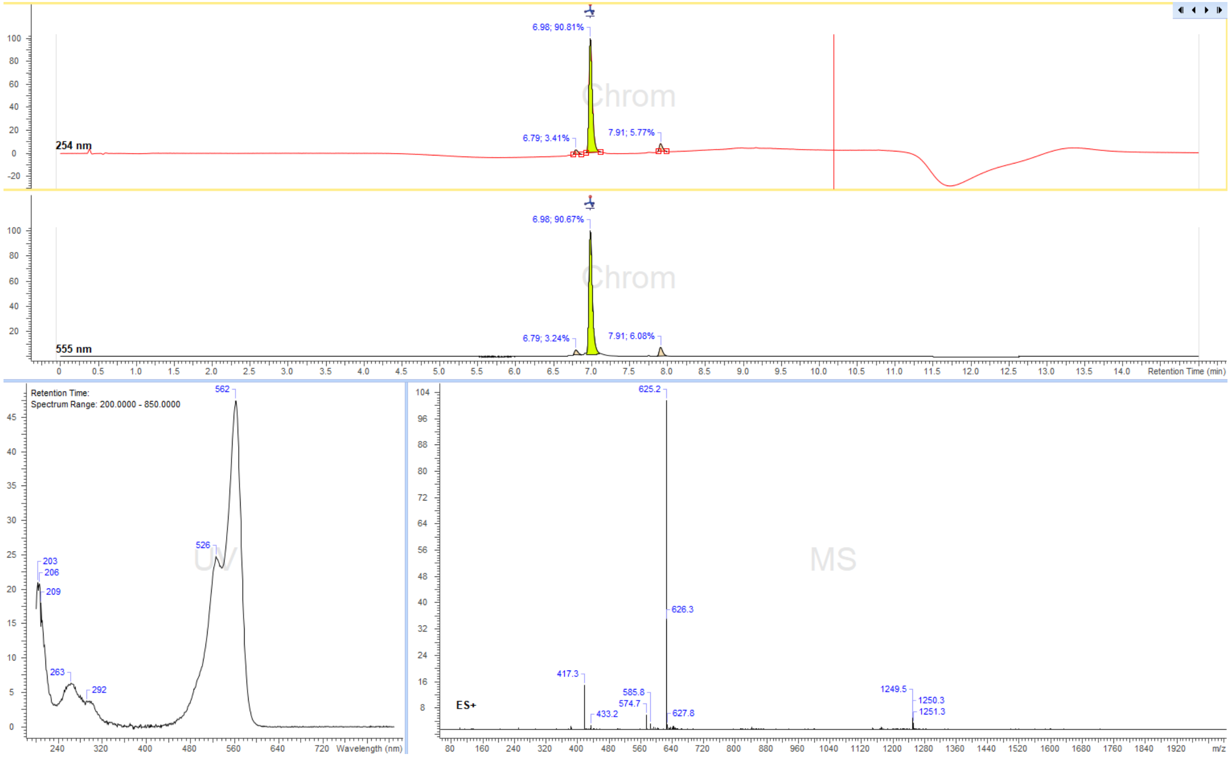

6.98; 90.81%
6.79; 3.41%
7.91; 5.77%
254 nm
Chrom
6.98; 90.67%
6.79; 3.24%
7.91; 6.08%
555 nm
Retention Time (min)
Retention Time:
Spectrum Range: 200.0000 - 850.0000
562
526
203
206
209
263
292
UV
Wavelength (nm)
625.2
626.3
417.3
433.2
585.8
574.7
627.8
1249.5
1250.3
1251.3
ES+
MS
m/z

5-TMR-PEG$_{10}$-Ado

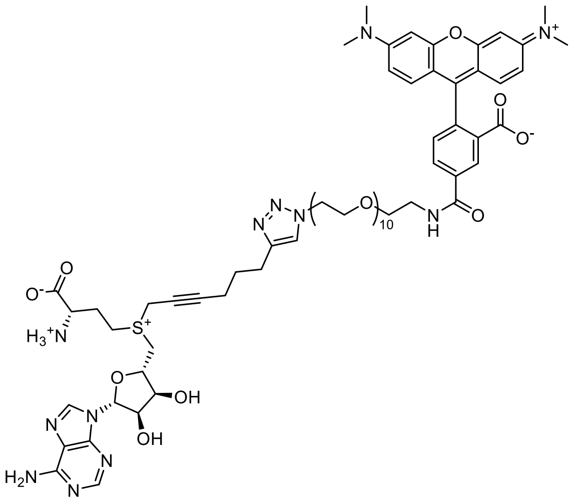

Reaction was performed following general synthesis procedure D from450 µL of 12 mM **octadiynyl-Ado**, 450 µL of 7.4 mM **5-TMR-PEG$_{10}$-N$_3$** and 450 µL copper (I) solution. The product was purified by preparative HPLC: Eurospher II 100-5 CN Column 250x30 mm, 5 µm (Knauer, Cat. No. 25ME200E2J) column, flow rate: 35 mL/min, solvent A: 10 mM ammonium formate (pH 3.5) and solvent B: MeOH at room temperature, gradient A:B-2 min 80:20 isocratic, 2-12 min 80:20 to 0:100 gradient, 12-15min 0:100 isocratic, 15-16 min 0:100 to 80:20 gradient and 16-20 min isocratic 80:20. Collected fractions of pure product were concentrated and concentration of the obtained stock solution was determined with nanodrop based on the absorption at λ=555nm ($\varepsilon_{555}$=83600 L·mol$^{-1}$·cm$^{-1}$). Obtained 0.72mL, 1.78 mM of stock solution in 38% yield.

ESI-MS, positive mode: m/z= 714.7 $[M]^{2+}$.

HRMS (ESI) calculated for $C_{69}H_{95}N_{12}O_9S$ $[M]^{2+}$ 714.3312, found 714.3321.

Copy of HRMS:

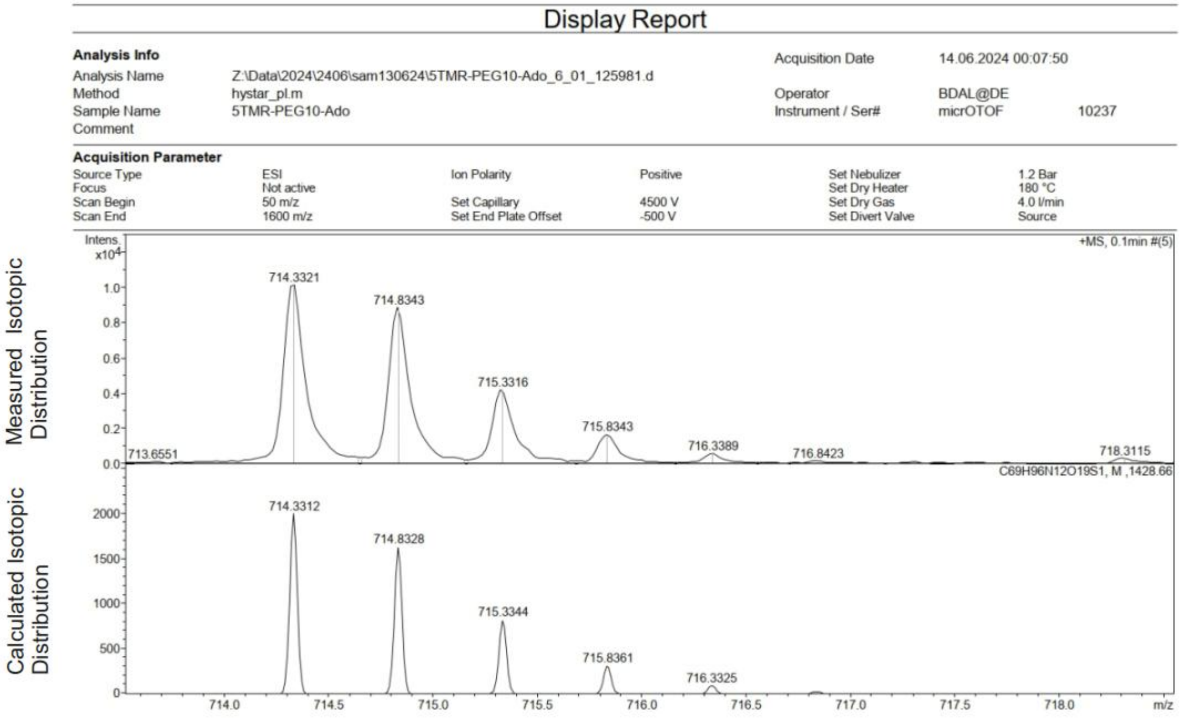

LC/MS:

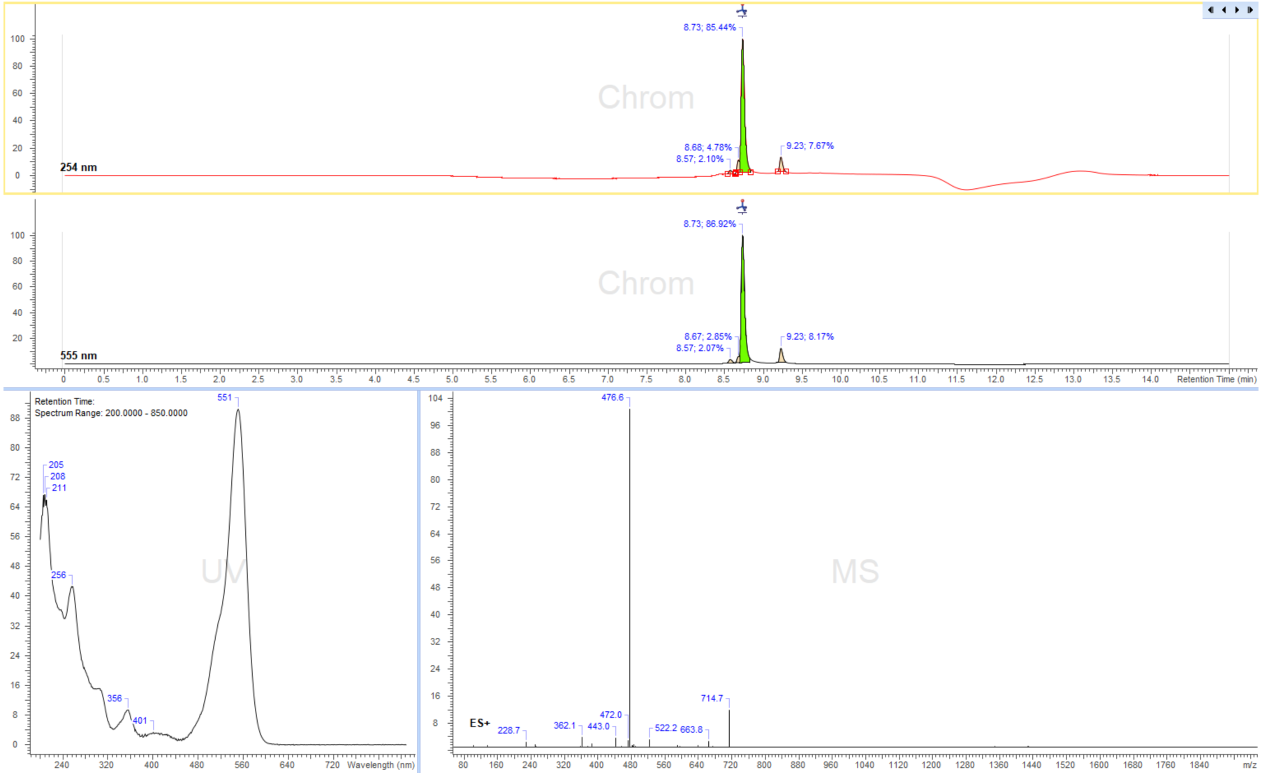
8.73; 85.44%
8.68; 4.78%
8.57; 2.10%
9.23; 7.67%
254 nm
8.73; 86.92%
8.67; 2.85%
8.57; 2.07%
9.23; 8.17%
555 nm
Retention Time (min)
Retention Time:
Spectrum Range: 200.0000 - 850.0000
551
205
208
211
256
356
401
Wavelength (nm)
476.6
ES+
228.7
362.1
443.0
472.0
522.2
663.8
714.7
m/z

6-TMR-PEG$_{10}$-Ado

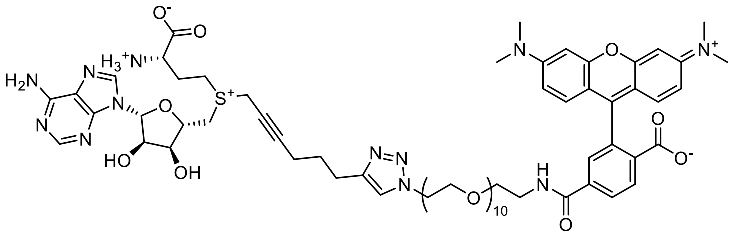

Reaction was performed following general synthesis procedure D from 450 µL of 12 mM **octadiynyl-Ado**, 450 µL of 6 mM **6-TMR-PEG$_{10}$-N$_3$** and 450 µL copper (I) solution. The product was purified by preparative HPLC: Eurospher II 100-5 CN Column 250x30 mm, 5 µm (Knauer, Cat. No. 25ME200E2J) column, flow rate: 35 mL/min, solvent A: 10 mM ammonium formate (pH=3.5) and solvent B: MeOH at room temperature, gradient A:B-2 min 80:20 isocratic, 2-12 min 80:20 to 0:100 gradient, 12-15 min 0:100 isocratic, 15-16 min 0:100 to 80:20 gradient and 16-20 min isocratic 80:20. Collected fractions of pure product were concentrated and concentration of the obtained stock solution was determined with nanodrop based on the absorption at λ=555 nm ($\varepsilon_{555}$=86600 L·mol$^{-1}$·cm$^{-1}$). Obtained 0.9 mL, 1.77 mM of stock solution in 59% yield.

ESI-MS, positive mode: m/z= 714.3 $[M]^{2+}$.

HRMS (ESI) calculated for $C_{69}H_{95}N_{12}O_{19}S$ $[M]^{2+}$ 714.3312, found 714.3324.

Copy of HRMS:

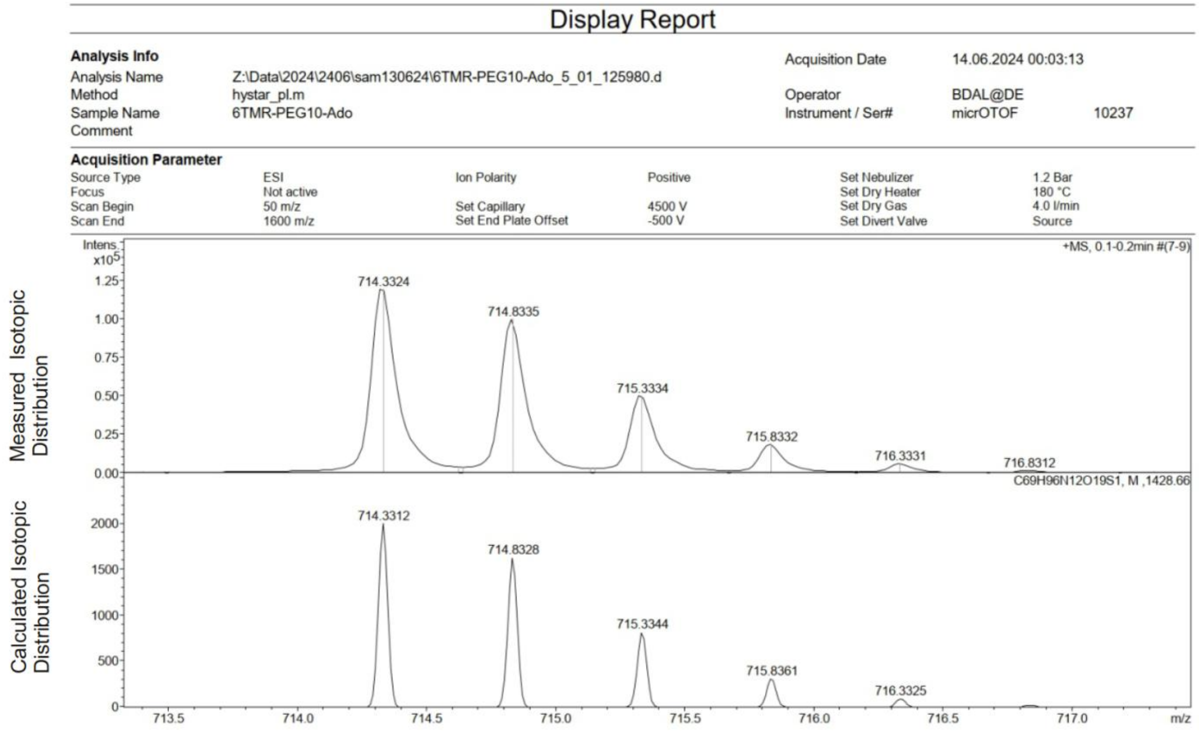

LC/MS:

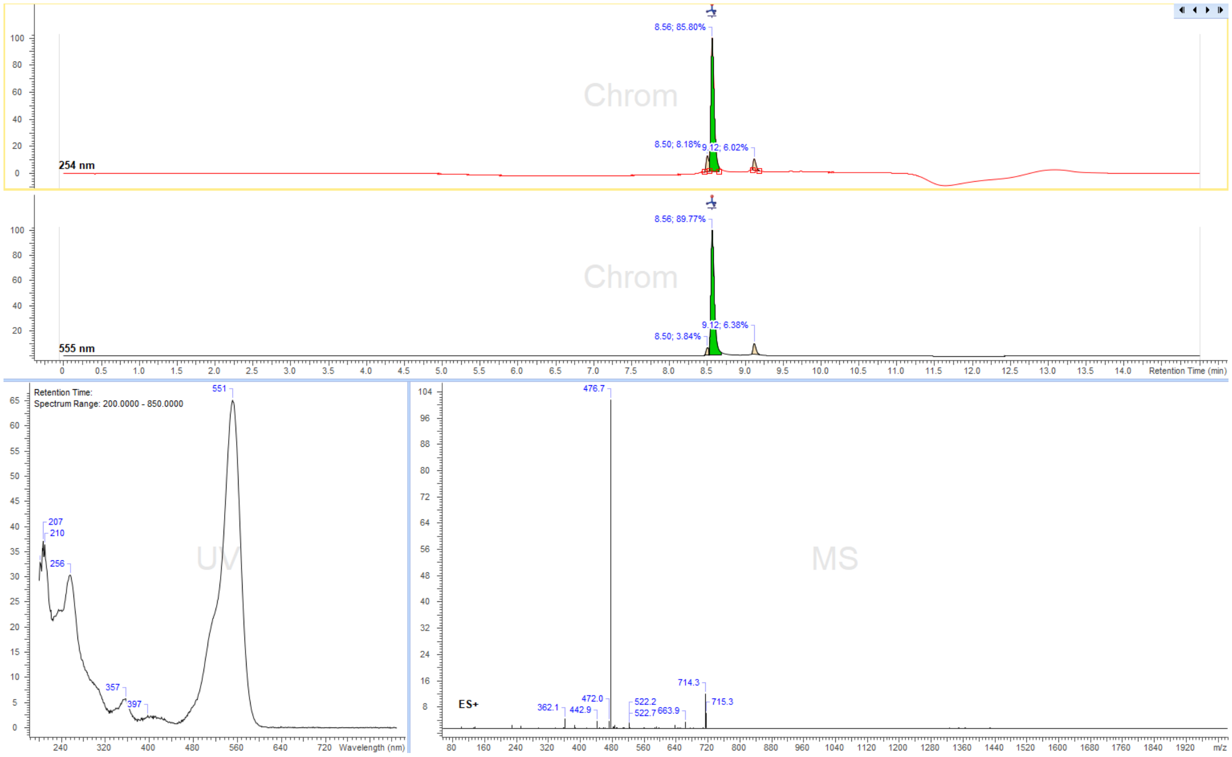
Chrom
8.56; 85.80%
8.50; 8.18%
9.12; 6.02%
254 nm
8.56; 89.77%
9.12; 6.38%
8.50; 3.84%
555 nm
Retention Time (min)
Retention Time:
Spectrum Range: 200.0000 - 850.0000
551
207
210
256
357
397
UV
Wavelength (nm)
476.7
MS
714.3
715.3
472.0
522.2
522.7
663.9
362.1
442.9
ES+
m/z

Cy3B-PEG$_{10}$-Ado

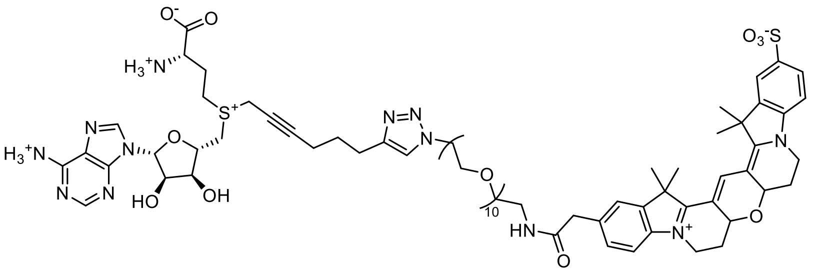

Reaction was performed following general synthesis procedure D from 200 µL of 10.4 mM **octadiynyl-Ado**, 150 µL of 9.46 mM **Cy3B-PEG$_{10}$-N$_3$** and 200 µL copper (I) solution. The product was purified by preparative HPLC: Eurospher II 100-5 CN Column 250x30 mm, 5 µm (Knauer, Cat. No. 25ME200E2J) column, flow rate: 35 mL/min, solvent A: 10 mM ammonium formate (pH=3.5) and solvent B: MeOH at room temperature, gradient A:B-2 min 80:20 isocratic, 2-12 min 80:20 to 0:100 gradient, 12-15 min 0:100 isocratic, 15-16 min 0:100 to 80:20 gradient and 16-20 min isocratic 80:20. Collected fractions of pure product were concentrated and concentration of the obtained stock solution was determined with nanodrop based on the absorption at λ=559 nm ($\varepsilon_{559}$=121000 L·mol$^{-1}$·cm$^{-1}$). Obtained 1.32 mL, 0.50 mM of stock solution in 47% yield.

ESI-MS, positive mode: m/z= 779.6 $[M]^{2+}$.

HRMS (ESI) calculated for $C_{75}H_{105}N_{12}O_{20}S_2$ $[M]^{2+}$ 779.3539, found 779.3522.

Copy of HRMS:

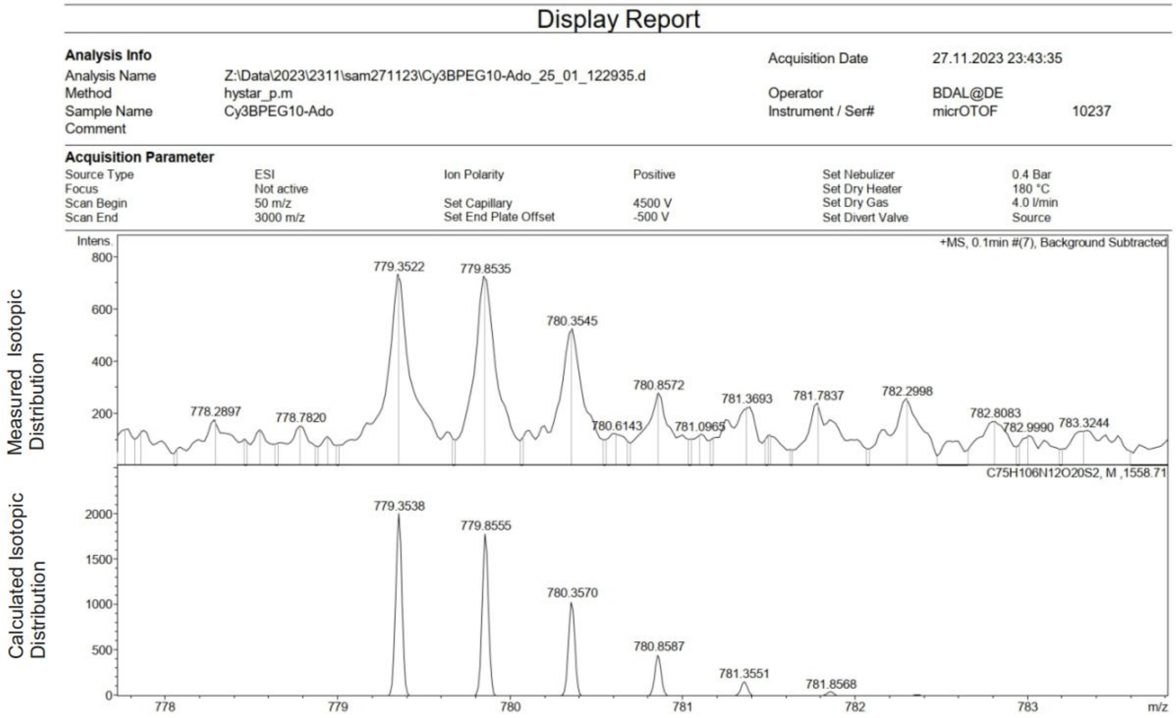

LC/MS:

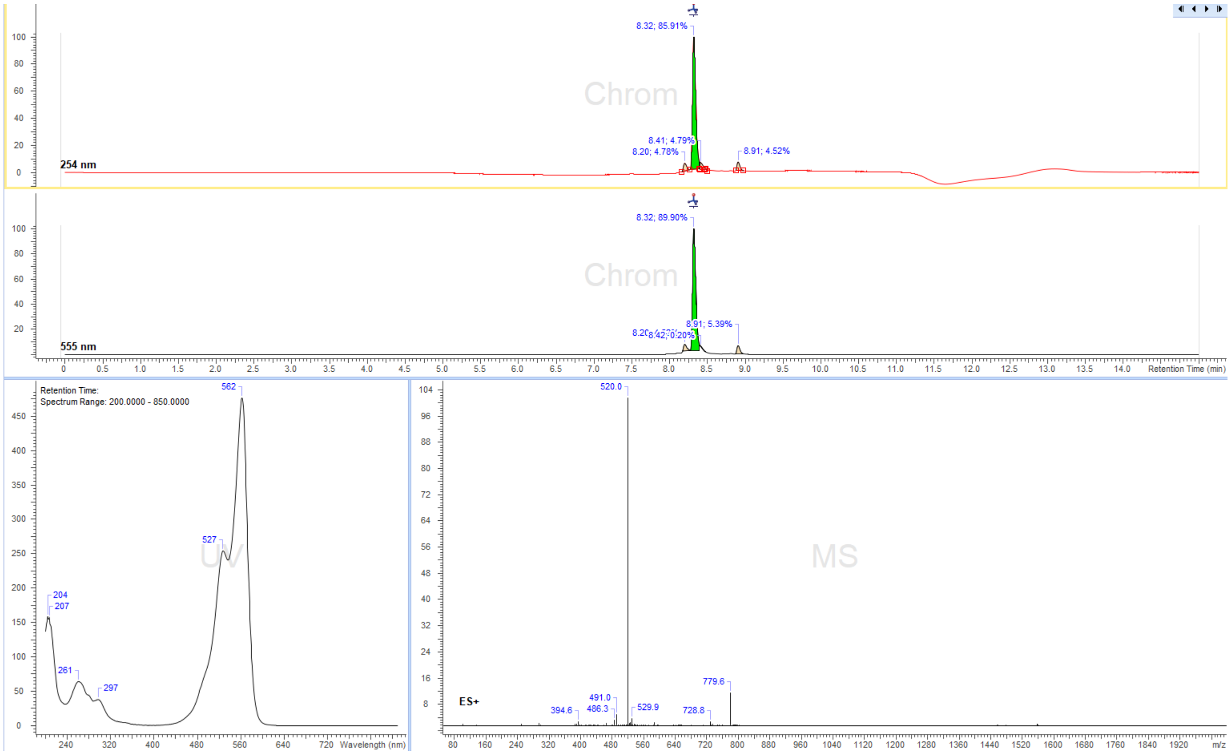
Chrom
254 nm
8.32; 85.91%
8.41; 4.79%
8.20; 4.78%
8.91; 4.52%
Chrom
555 nm
8.32; 89.90%
8.91; 5.39%
8.42; 0.20%
Retention Time (min)
Retention Time:
Spectrum Range: 200.0000 - 850.0000
UV
562
527
204
207
261
297
Wavelength (nm)
MS
ES+
520.0
779.6
491.0
486.3
529.9
394.6
728.8
m/z

Cy3B-$(CH_2)_3$-Ado

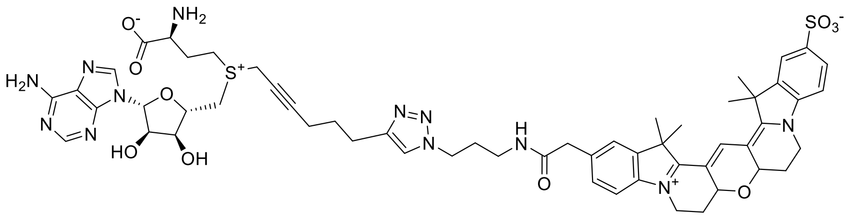

Reaction was performed following general synthesis procedure D from 148 µL of 8.11 mM **octadiynyl-Ado** and 176 µL of 13.86 mM **Cy3B-$(CH_2)_3$-$N_3$**. The product was purified by preparative HPLC: Eurospher II 100-5 CN Column 250x30 mm, 5 µm (Knauer, Cat. No. 25ME200E2J) column, flow rate: 35 mL/min, solvent A: 10 mM ammonium formate (pH=3.5) and solvent B: MeOH at room temperature, gradient A:B-2 min 80:20 isocratic, 2-12 min 80:20 to 0:100 gradient, 12-15 min 0:100 isocratic, 15-16 min 0:100 to 80:20 gradient and 16-20 min isocratic 80:20. Collected fractions of pure product were concentrated and concentration of the obtained stock solution was determined with nanodrop based on the absorption at λ=559 nm ($\varepsilon_{559}$=121000 L·mol$^{-1}$·cm$^{-1}$). Obtained 1.4 mL, 0.22 mM of stock solution in 69% yield.

ESI-MS, positive mode: m/z= 1131.4 $[M]^+$and 566.2 $[M]^{2+}$.

HRMS (ESI) calculated for $C_{56}H_{67}N_{12}O_{10}S_2$ 566:2306 $[M]^{2+}$, found 566.2312.

Copy of HRMS:

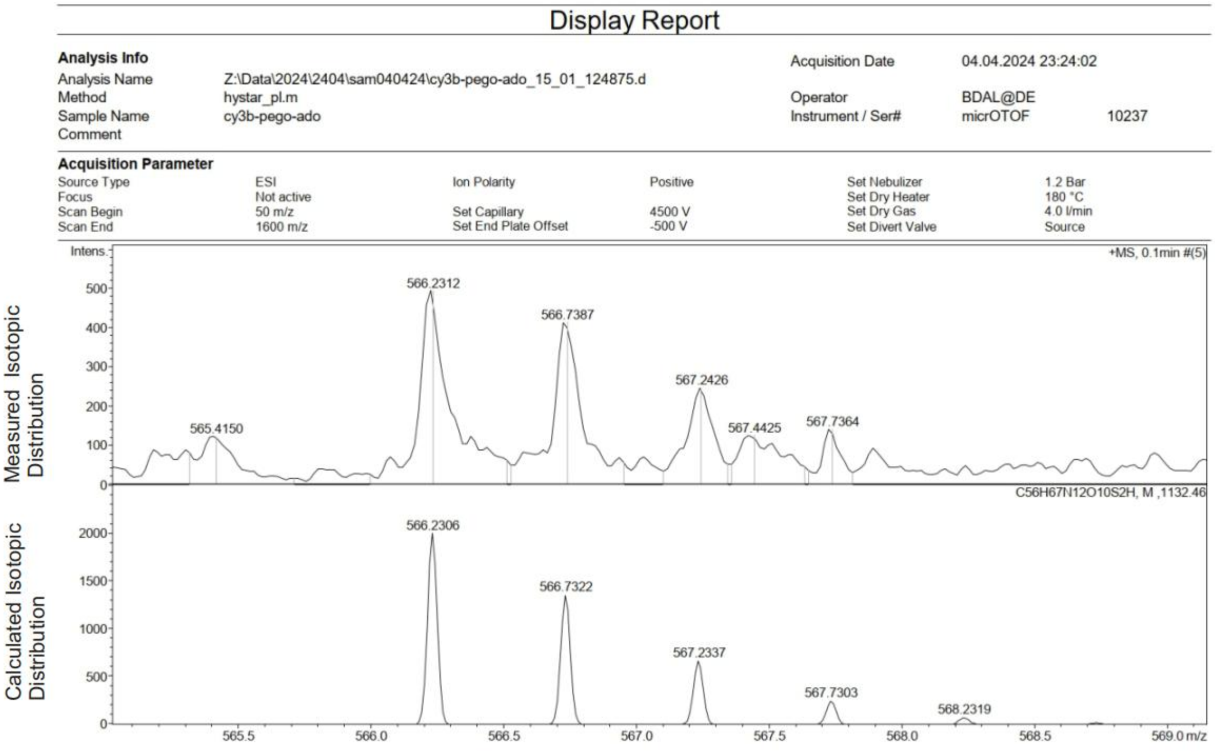

LC/MS:

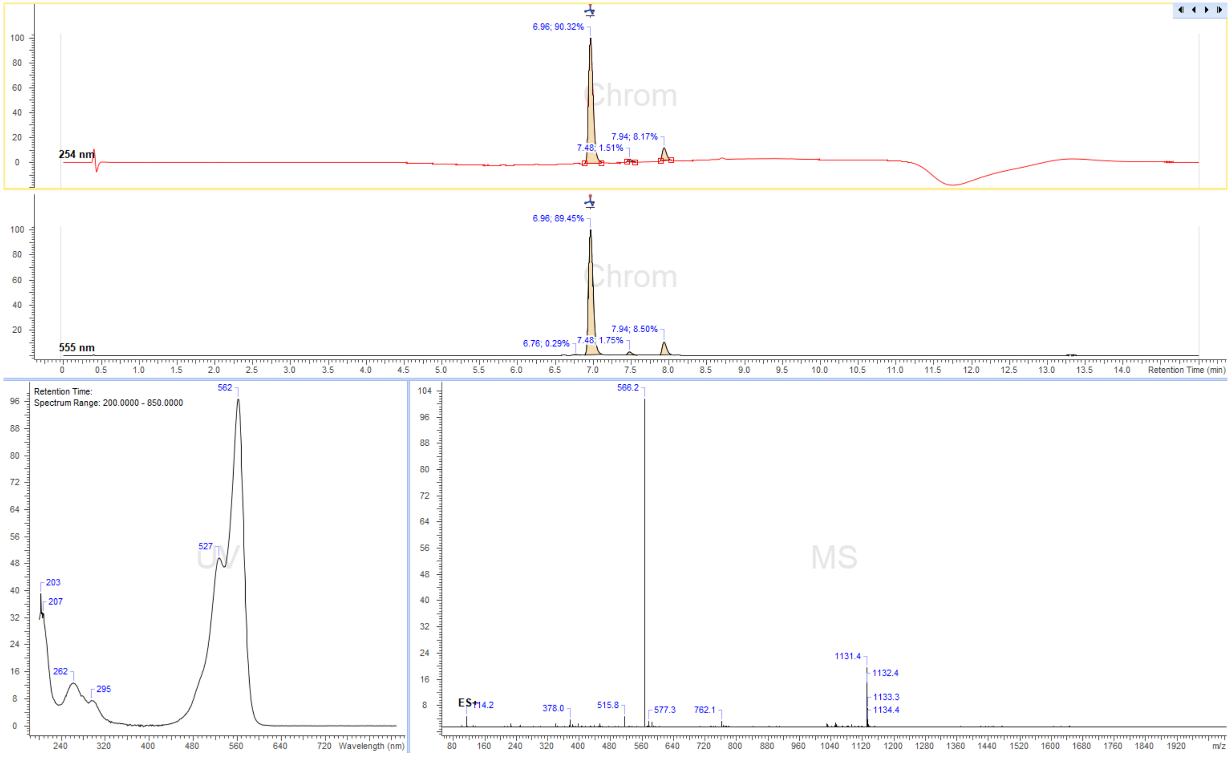

6.96; 90.32%
Chrom
7.94; 8.17%
7.48; 1.51%
254 nm
6.96; 89.45%
7.94; 8.50%
6.76; 0.29%
7.48; 1.75%
555 nm
Retention Time (min)
Retention Time:
Spectrum Range: 200.0000 - 850.0000
562
527
203
207
262
295
Wavelength (nm)
566.2
MS
1131.4
1132.4
1133.3
1134.4
ES+
114.2
378.0
515.8
577.3
762.1
m/z

CalFluor647-Ado

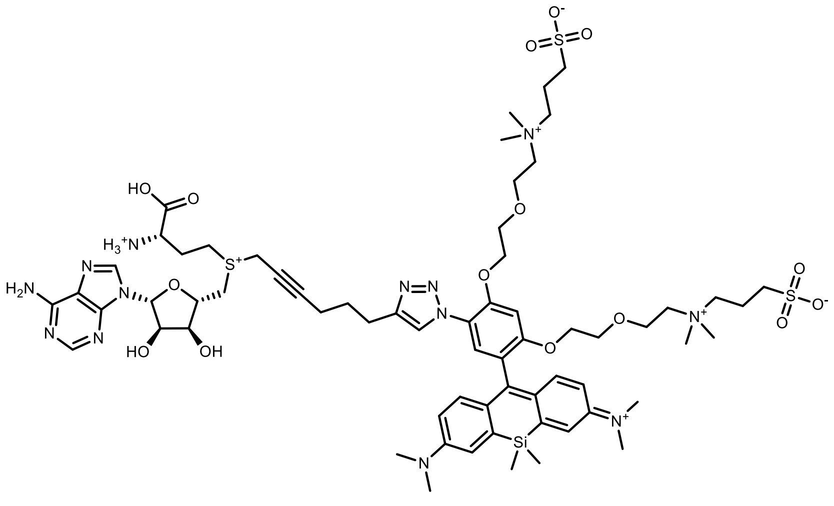

CalFluor647-Ado

Reaction was performed following general synthesis procedure D from 100 µl of 10mM **Octadiynyl-Ado** solution and 100 µl of 10 mM **CalFluor647-N$_3$** (BroadFarm Cat. No. BP-28103) in 25 mM ammonium formate buffer pH = 3.5. 20 µl of fresly prepared copper (I) containing solution was used. The product was purified by preparative HPLC using Eurospher II 100-5 C18 Column 250x30 mm, 5 µm (Knauer, Cat. No. 25ME181E2J) column, flow rate: 35 ml/min, solvent A: 10 mM ammonium formate (pH 3.5) and solvent B: MeOH at room temperature. The following gradient was used: A:B – 0-2 min 80:20 isocratic, 2-12 min 80:20 to 0:100 gradient, 12-15min 100:0 isocratic, 15-17 min to 80:20 gradient and 17-24 min isocratic 80:20). Collected fractions of pure product were concentrated and concentrations of the obtained solutions were determined by their UV absorption ($\varepsilon_{650}$ = 95 000 L·mol$^{-1}$·cm$^{-1}$). Obtained 0.5 ml, 0.4 mM of cofactor CalFluor647-Ado, 20% yield.

HRMS (ESI) calcd for $C_{65}H_{94}N_{13}O_{15}S_3Si^+$ $[M+H]^+$ 1421.807, $C_{65}H_{95}N_{13}O_{15}S_3Si^{2+}$ $[M+2H]^{2+}$ 710.799, $C_{65}H_{96}N_{13}O_{15}S_3Si^{3+}$ $[M+3H]^{3+}$ 474.202

HRMS found $[M+2H]^{2+}$ 710.7988, $[M+3H]^{3+}$ 474.2017.

Copy of HRMS:

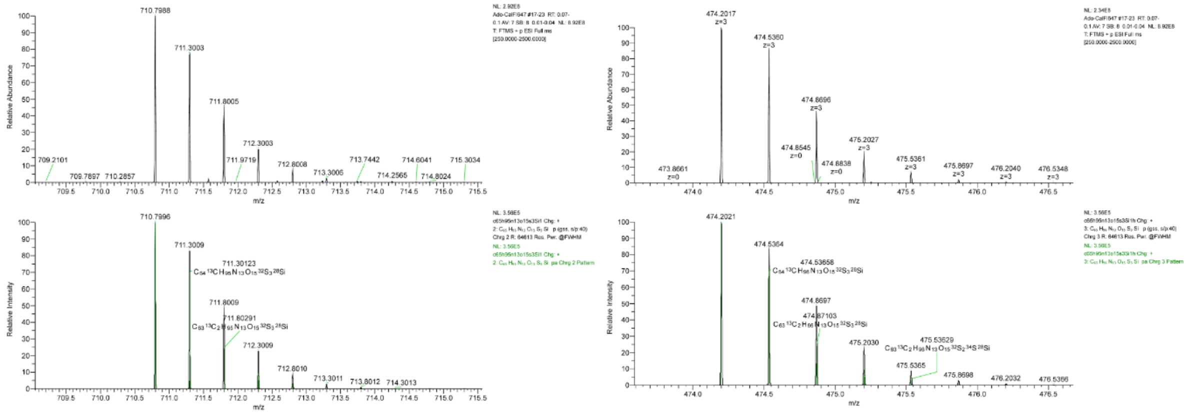

LC/MS:

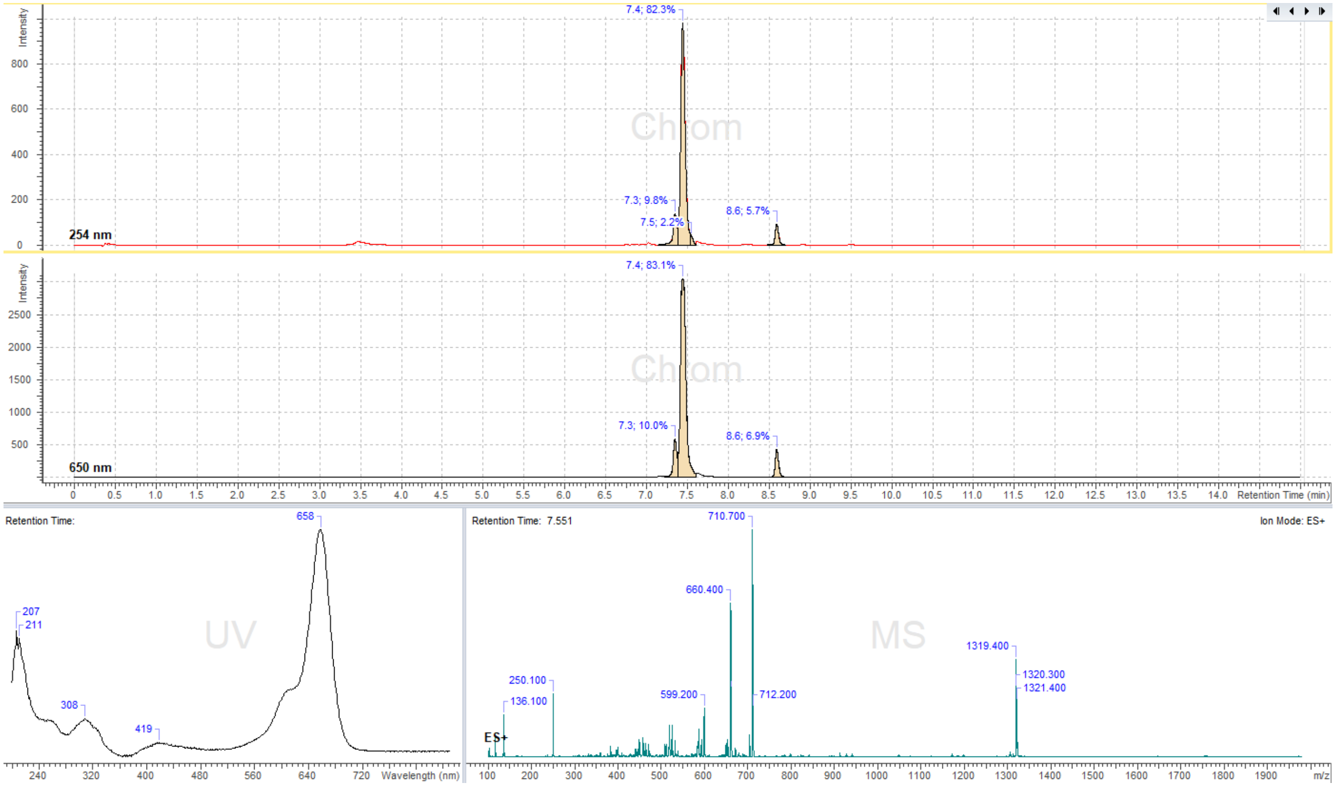

7.4; 82.3%
7.3; 9.8%
7.5; 2.2%
8.6; 5.7%
254 nm
7.4; 83.1%
7.3; 10.0%
8.6; 6.9%
650 nm
Retention Time (min)
Retention Time:
658
207
211
308
419
UV
Wavelength (nm)
Retention Time: 7.551
Ion Mode: ES+
710.700
660.400
1319.400
1320.300
1321.400
250.100
136.100
599.200
712.200
MS
ES+
m/z

### Copies of NMR spectra

$^1H$ spectrum of 6-SiR-PEG$_4$-alkyne:

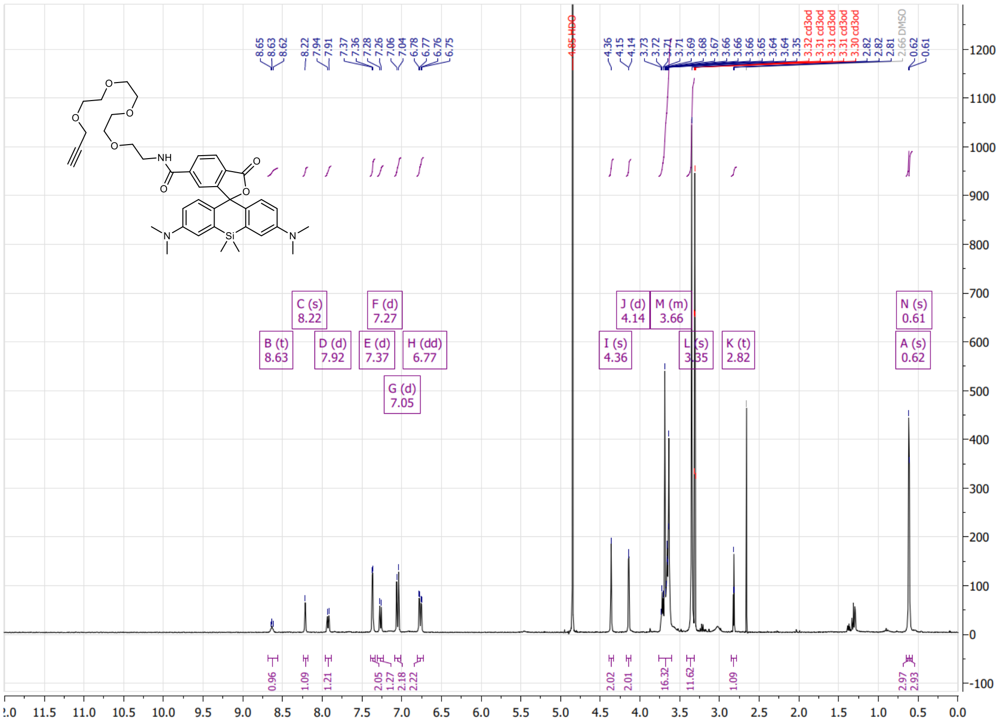

COSY spectrum of **6-SiR-PEG$_4$-alkyne**:

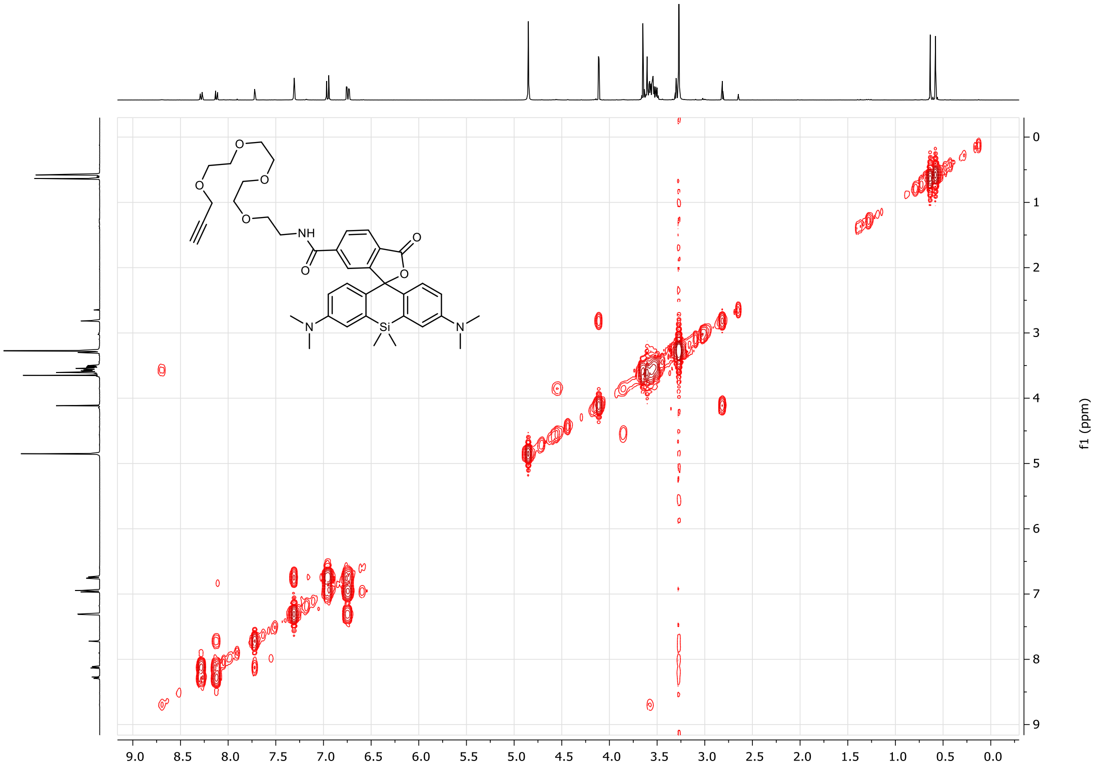

$^1H$ spectrum of 5-HMSiR-PEG$_4$-alkyne:

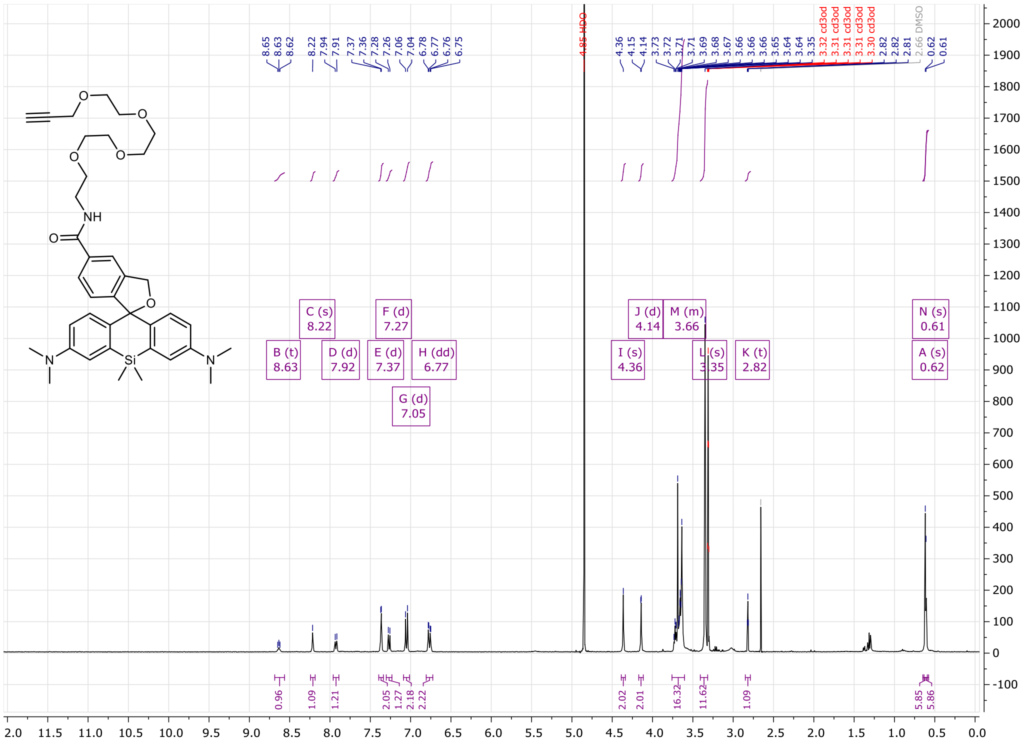

COSY spectrum of 5-HMSiR-PEG$_4$-alkyne:

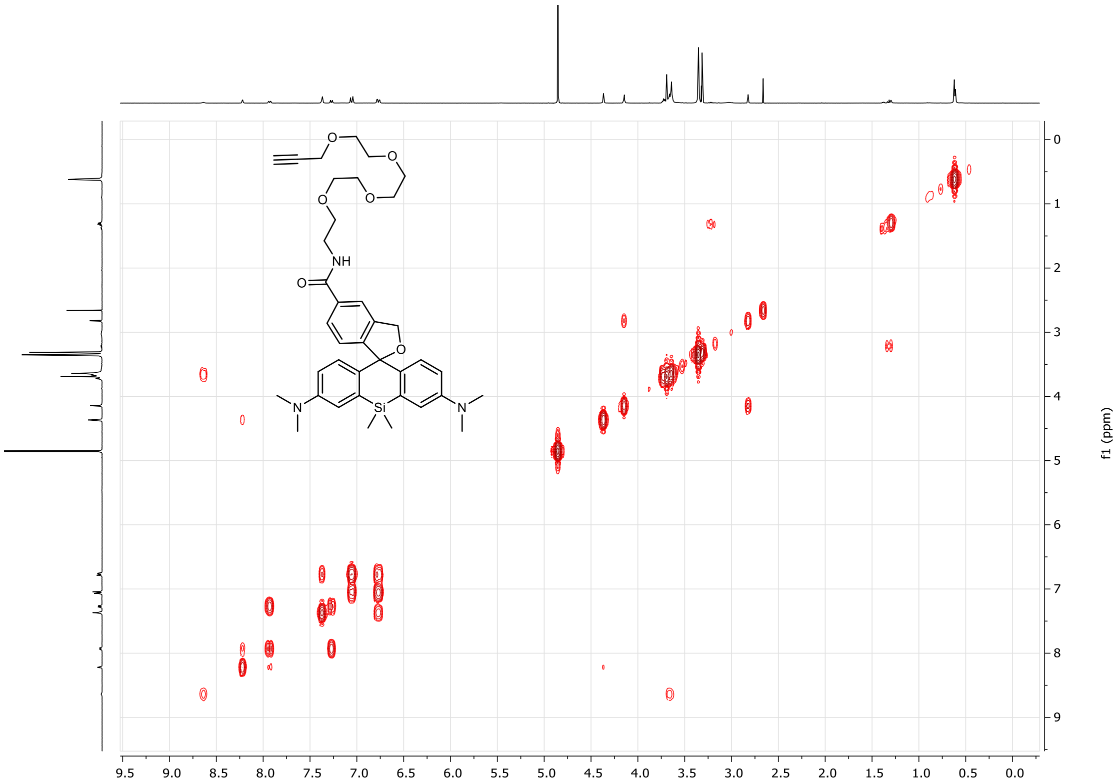

$^{1}H$ spectrum of 4-MeSiR-PEG$_4$-alkyne:

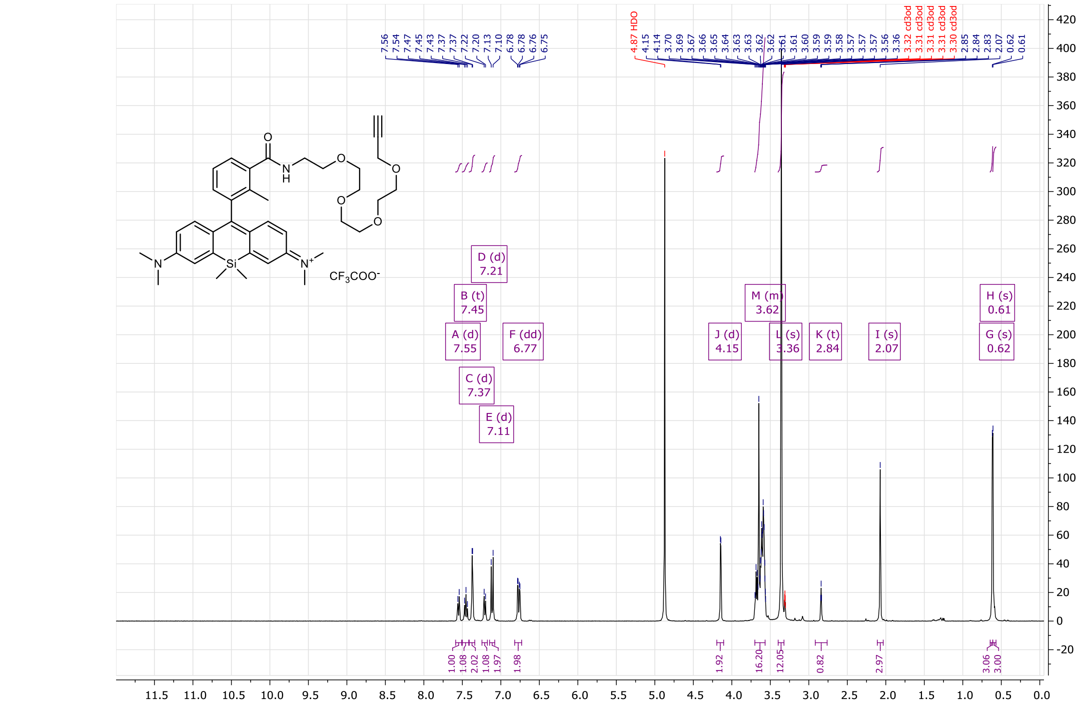

COSY spectrum of 4-MeSiR-PEG$_4$-alkyne:

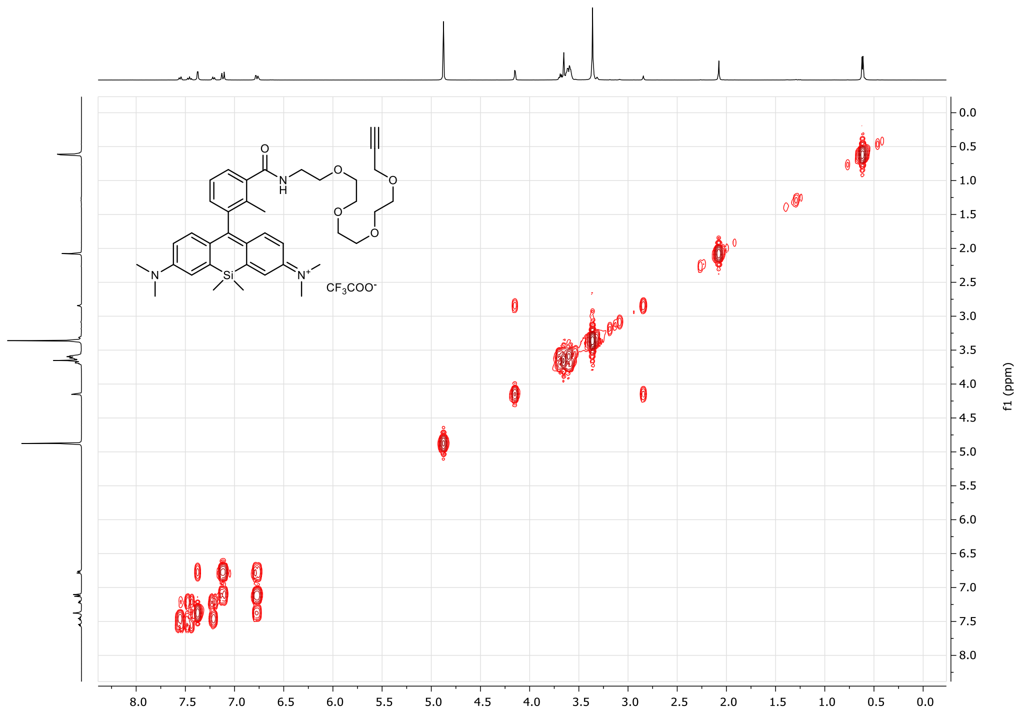

[1]H spectrum of **5-TMR-$(CH_2)_3$-$N_3$**:

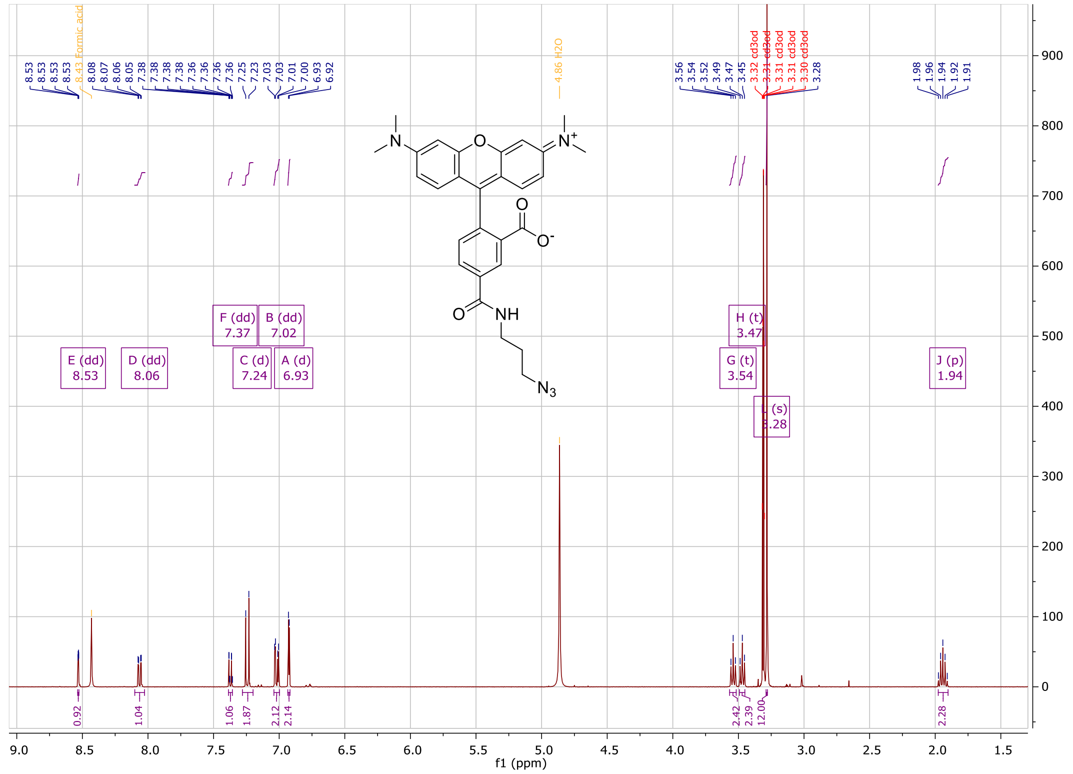

COSY spectrum of 5-TMR-$(CH_2)_3$-$N_3$:

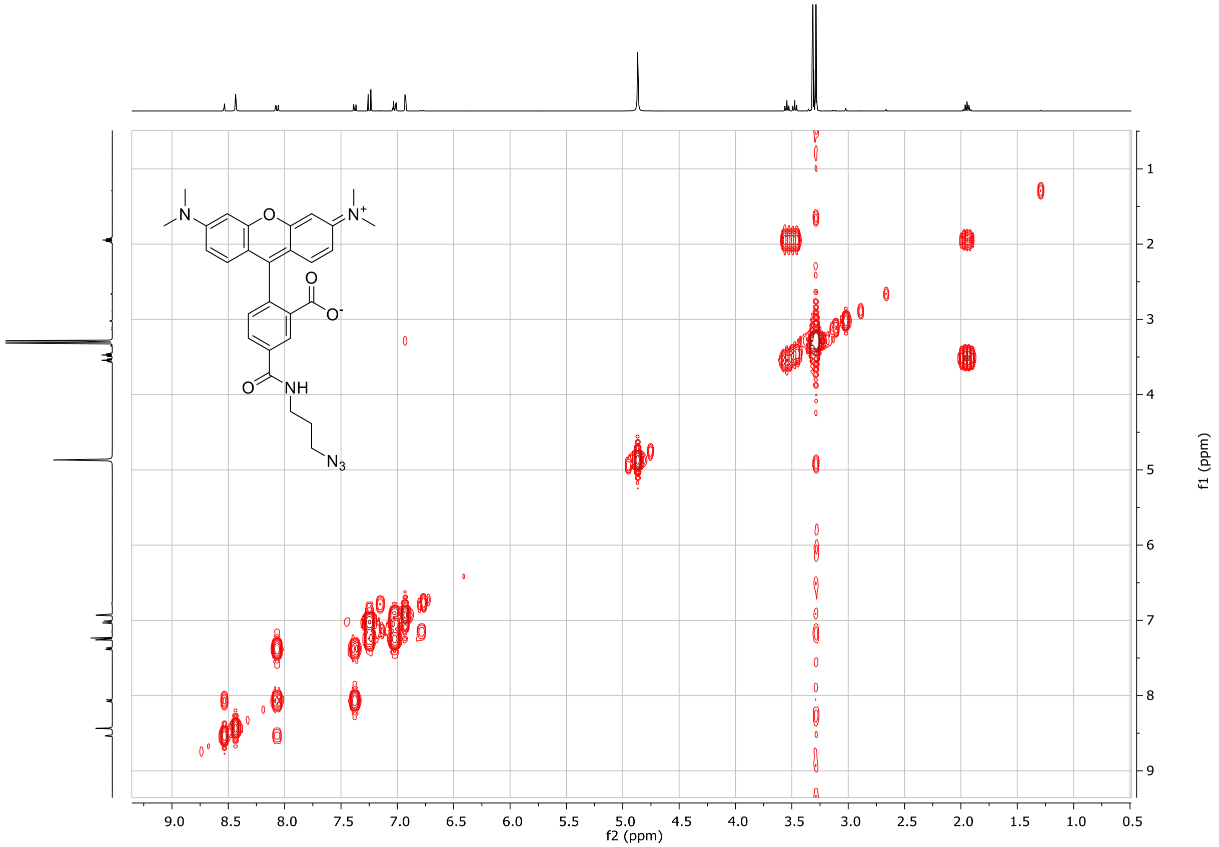

[1]H spectrum of **6-TMR-$(CH_2)_3$-$N_3$**:

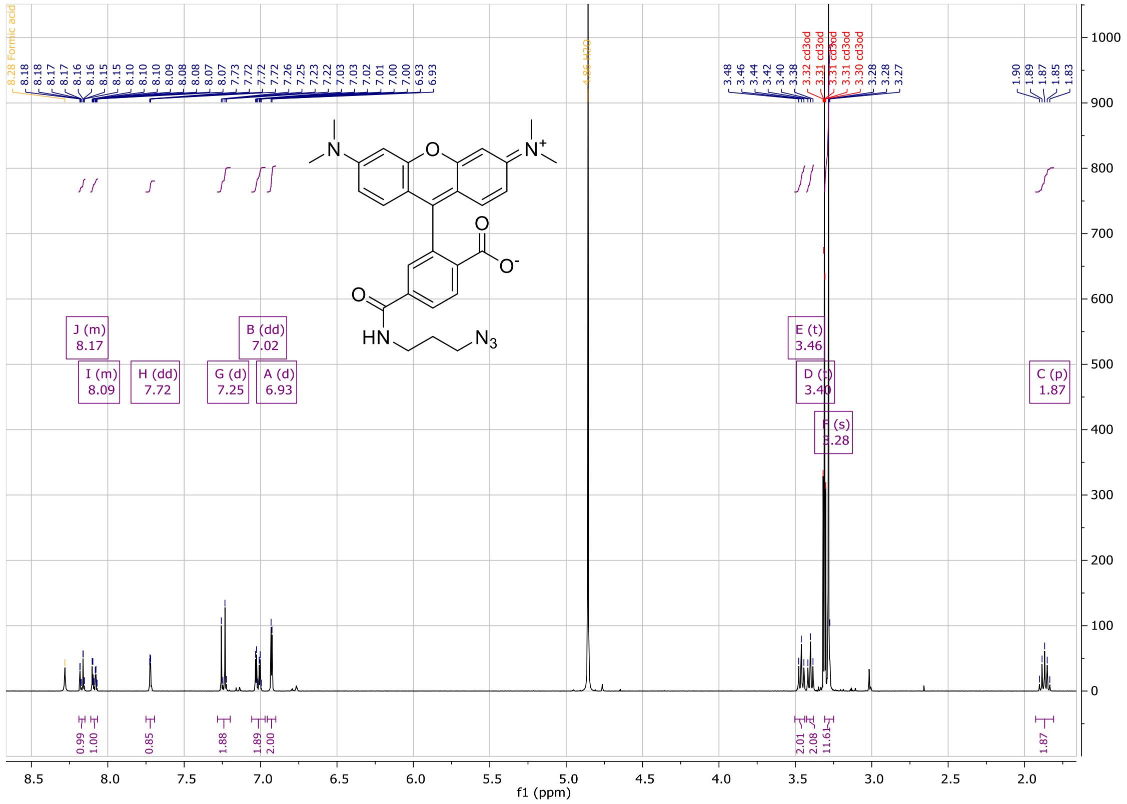

COSY spectrum of 6-TMR-$(CH_2)_3$-$N_3$:

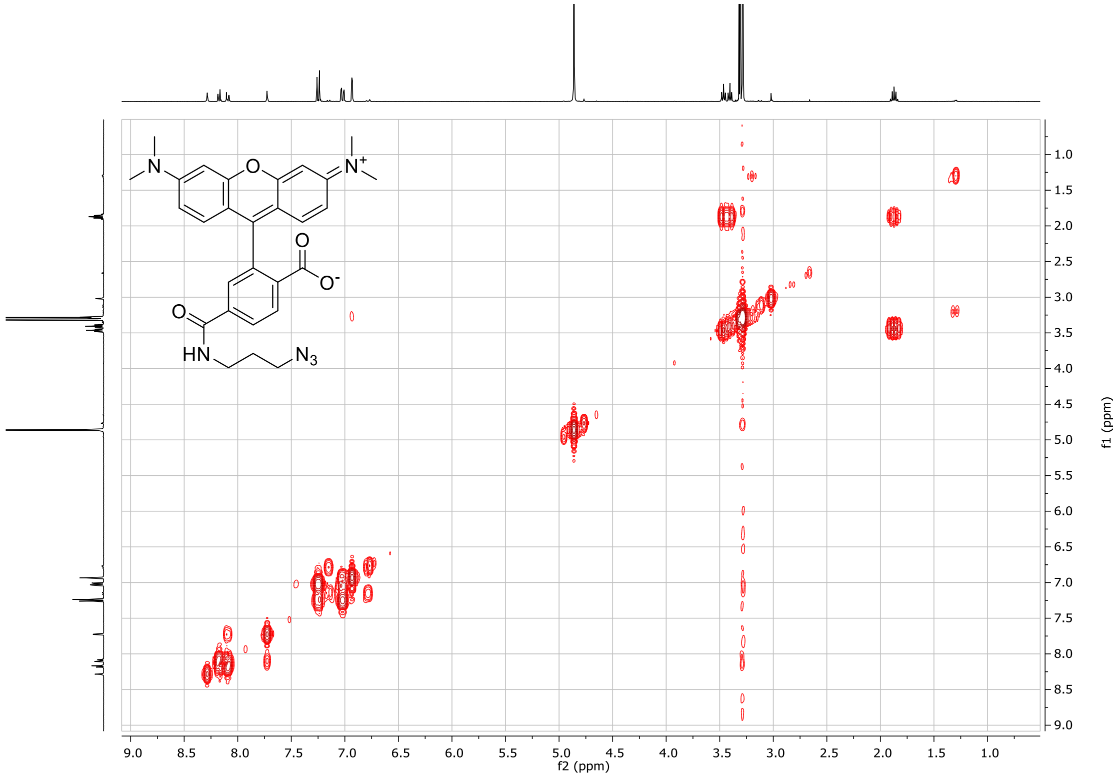

$^1$H spectrum of **Cy3B-(CH$_2$)$_3$-N$_3$**:

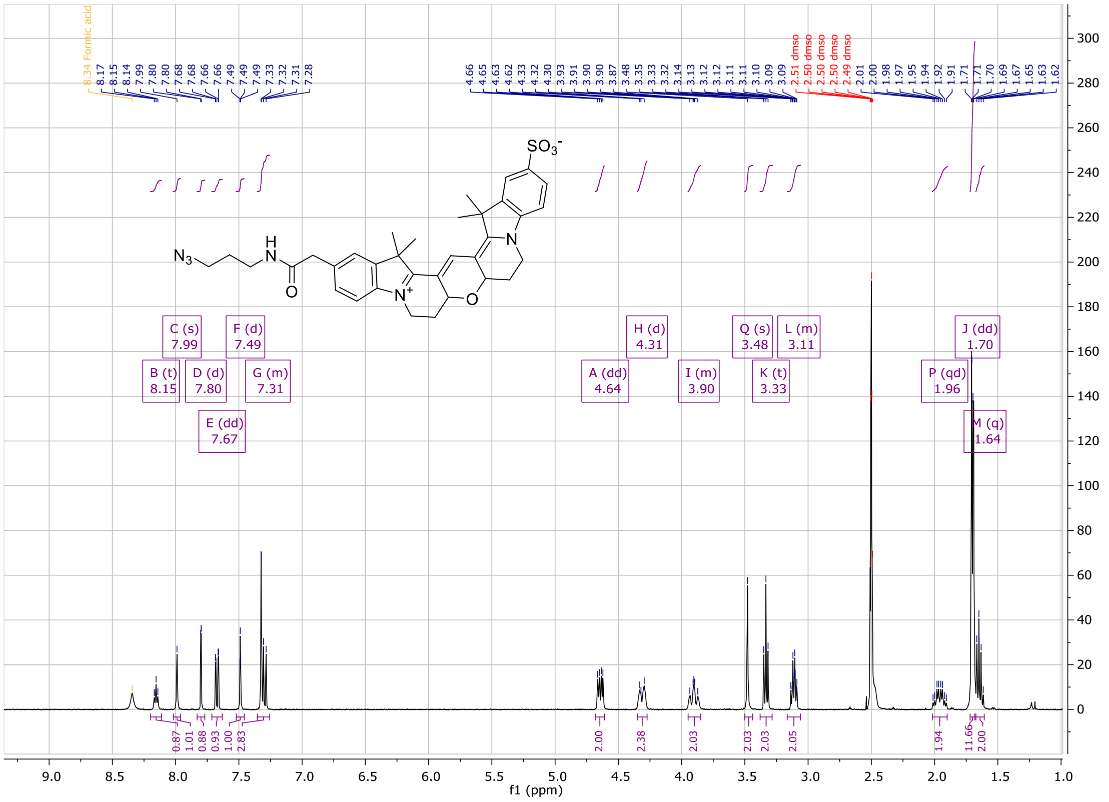

COSY spectrum of **Cy3B-(CH$_2$)$_3$-N$_3$**:

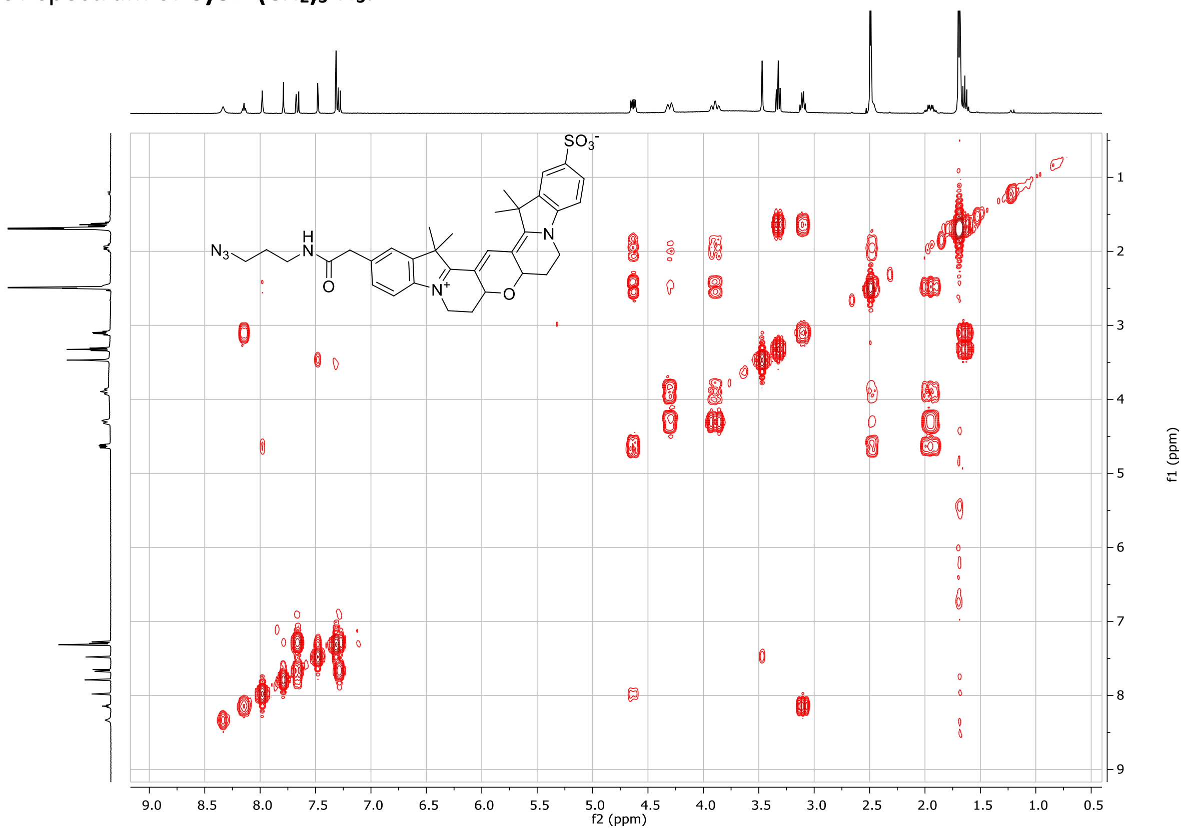

[1]H spectrum of **5-TMR-PEG$_3$-N$_3$**:

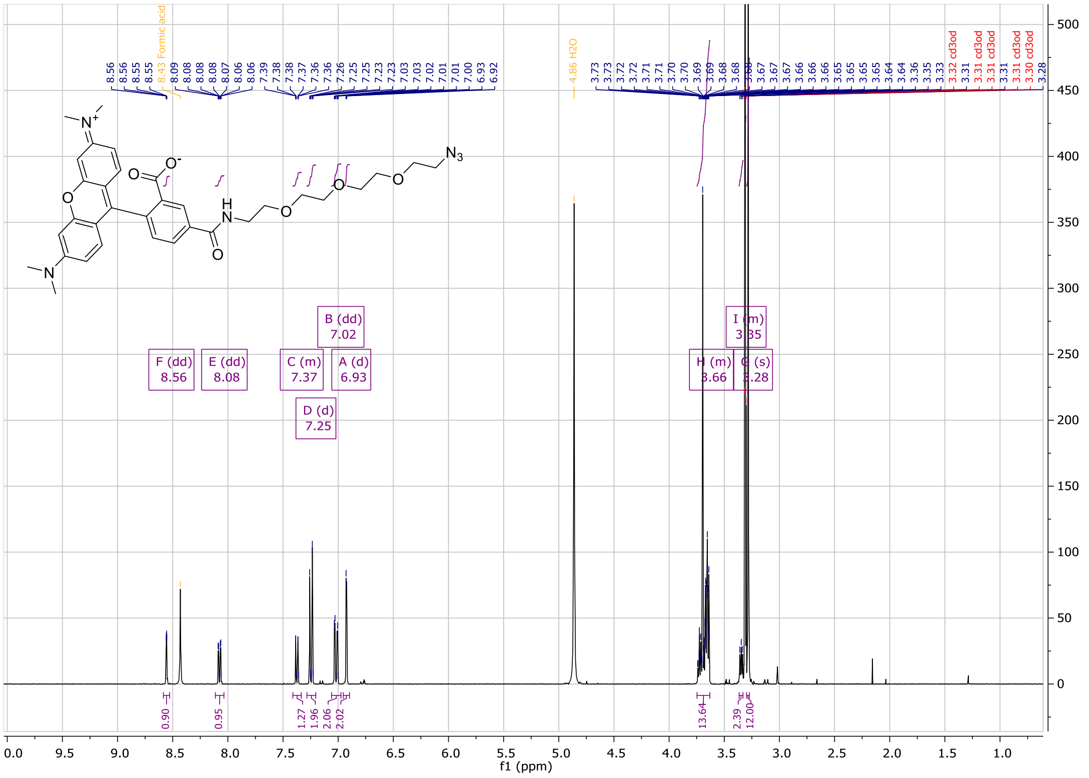

COSY spectrum of **5-TMR-PEG$_3$-N$_3$:**

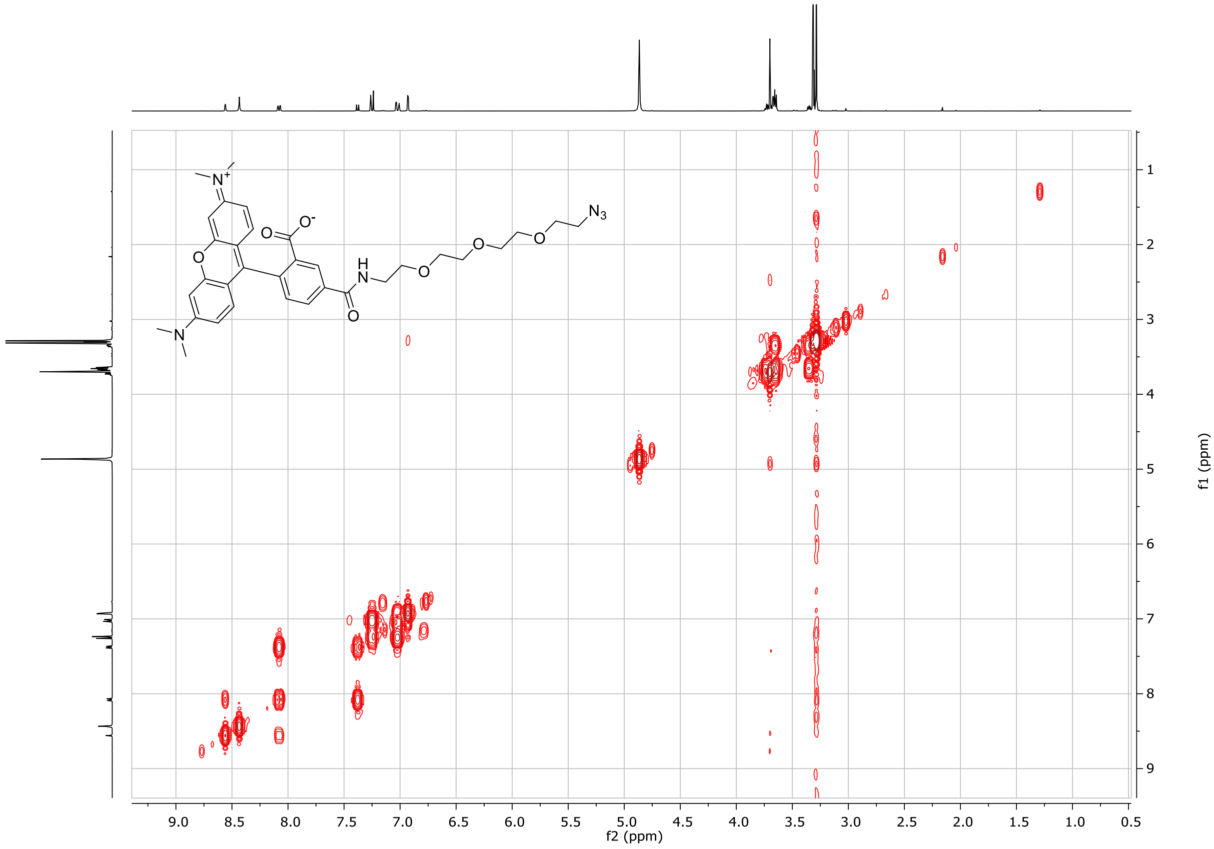

$^1$H spectrum of **6-TMR-PEG$_3$-N$_3$**:

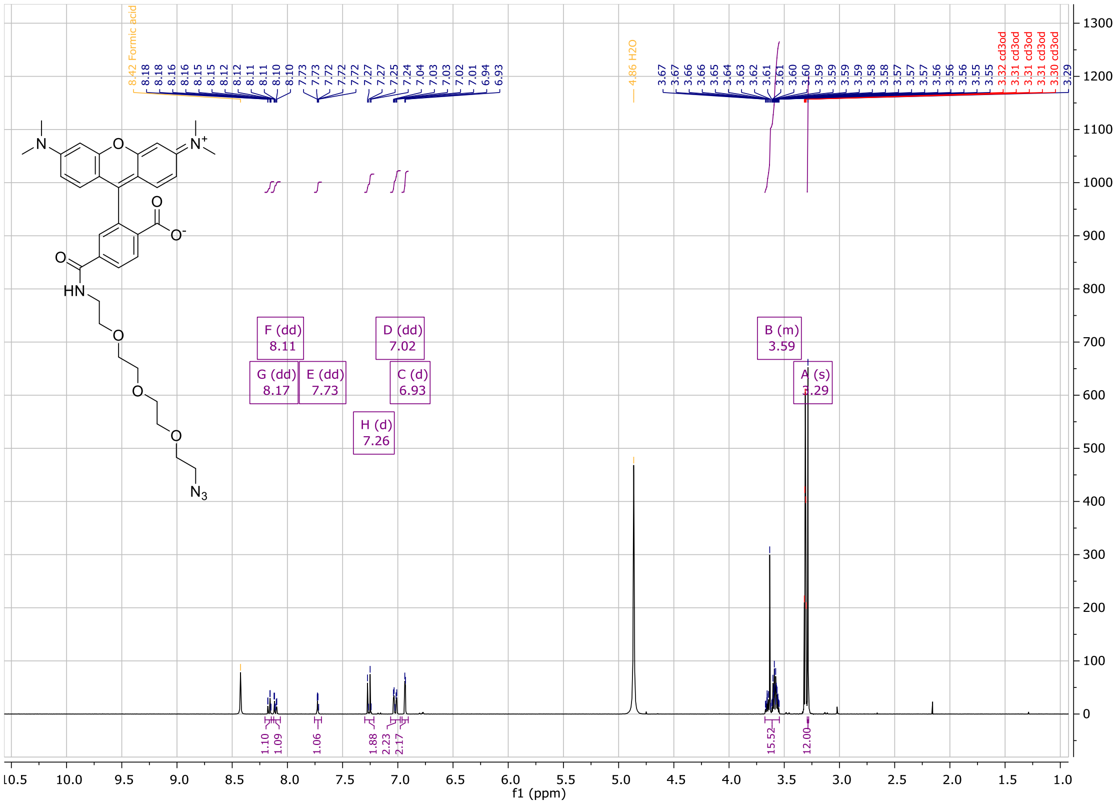

COSY spectrum of **6TMR-PEG$_3$-N$_3$:**

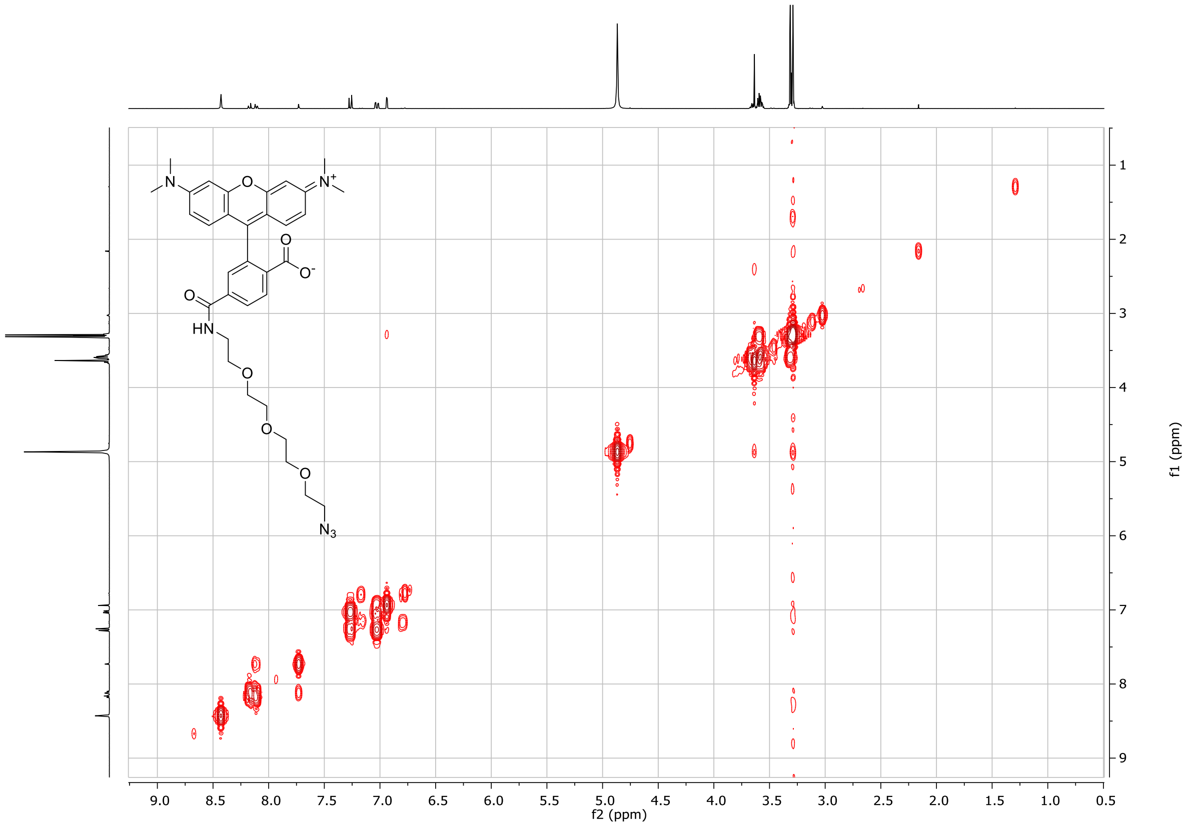

[1]H spectrum of **Cy3B-PEG$_3$-N$_3$**:

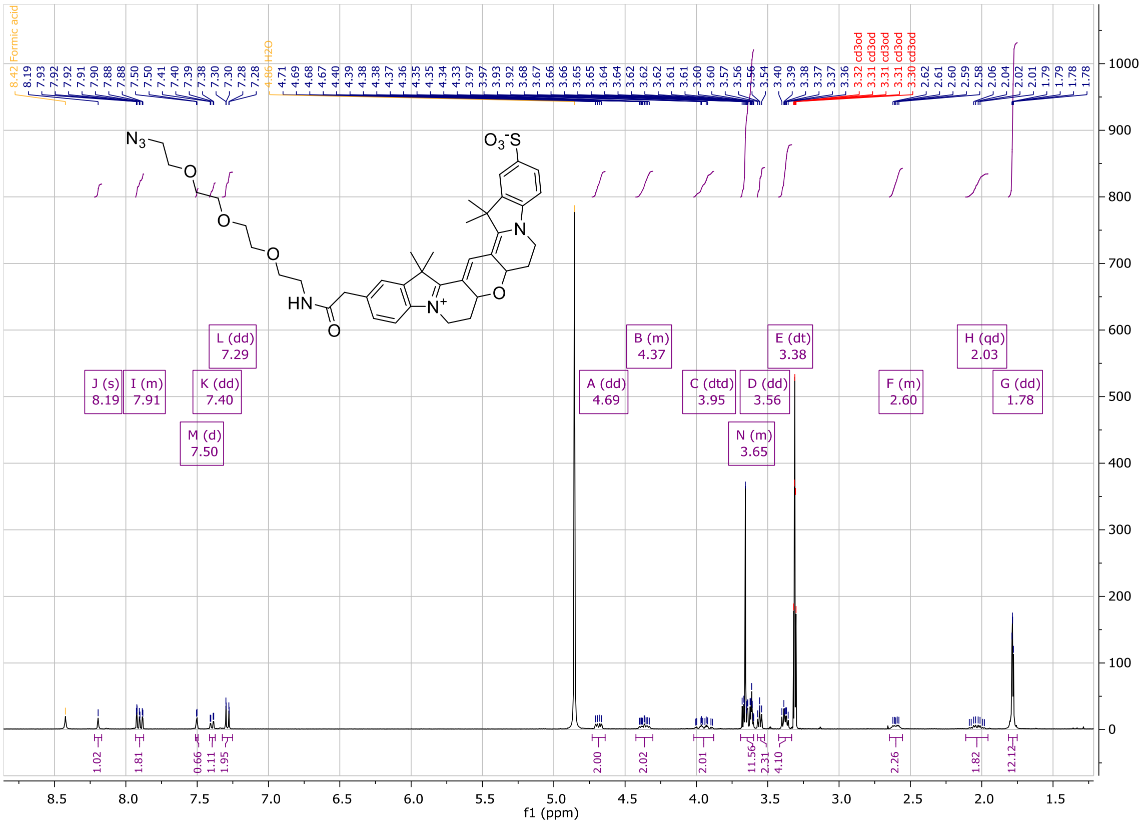

COSY spectrum of **Cy3B-PEG$_3$-N$_3$:**

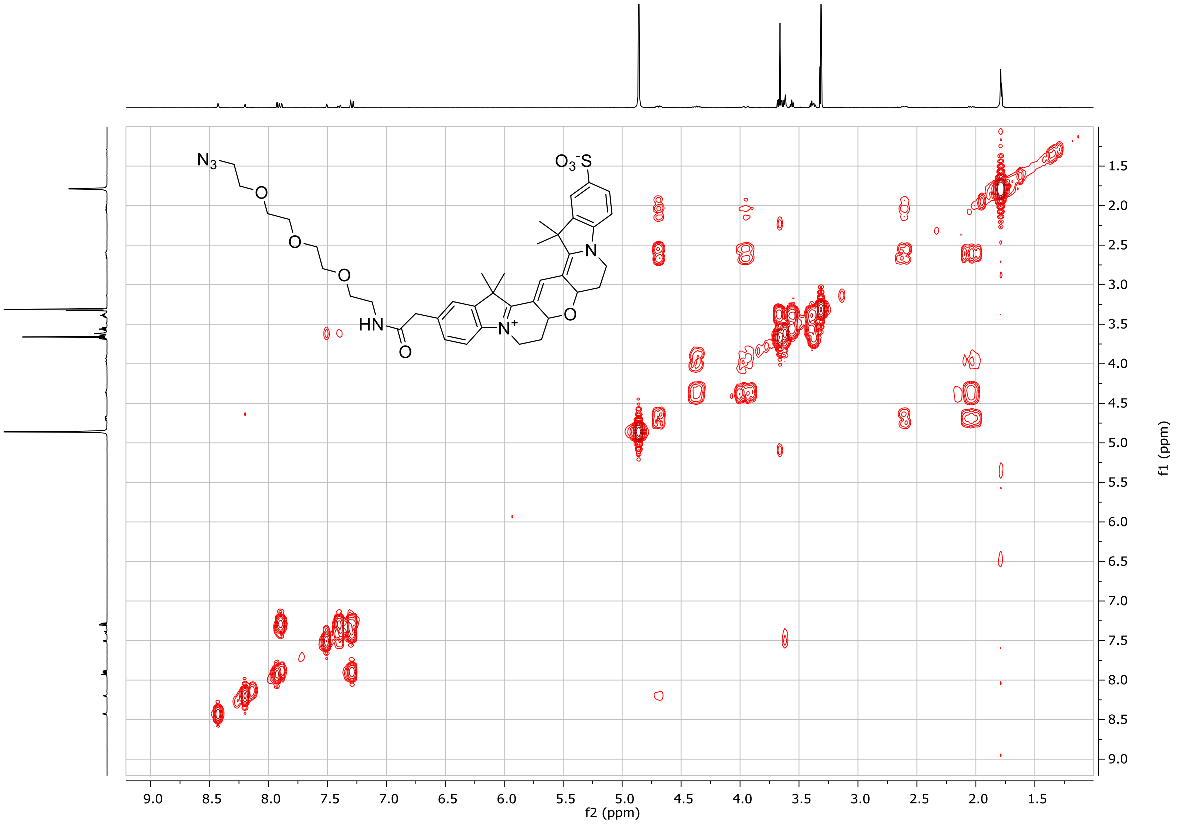

$^{1}$H spectrum of **5-TMR-PEG$_{10}$-N$_3$**:

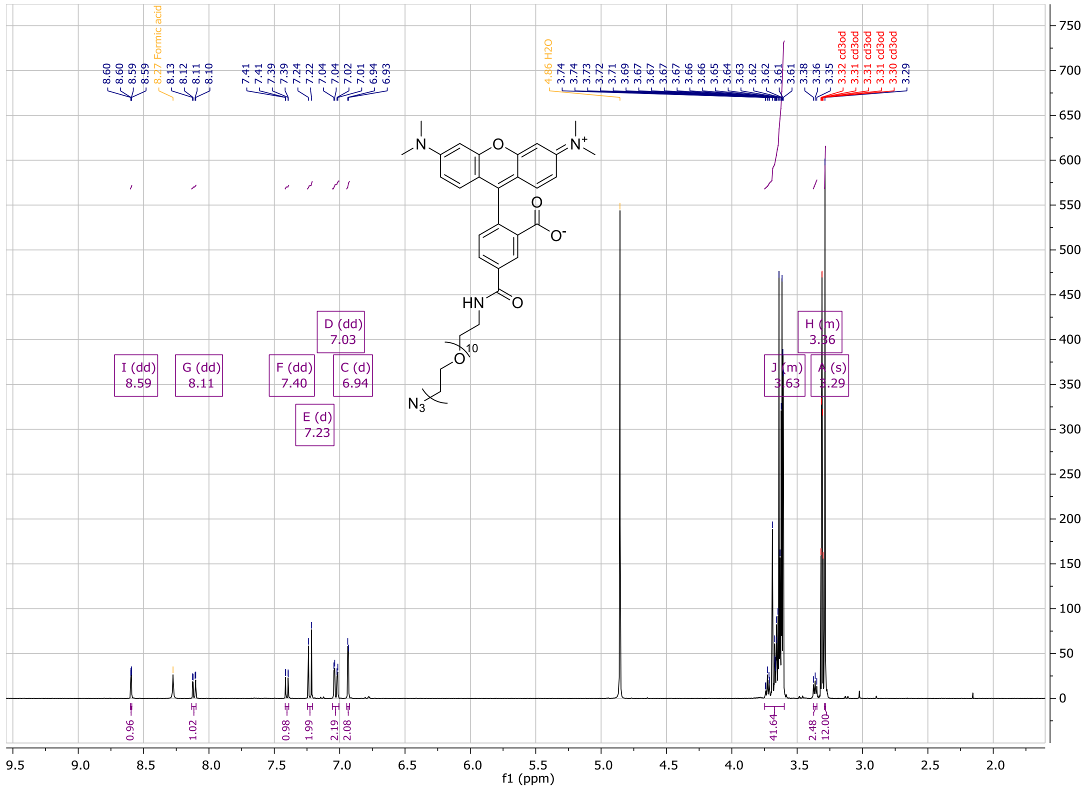

COSY spectrum of **5-TMR-PEG$_{10}$-N$_3$:**

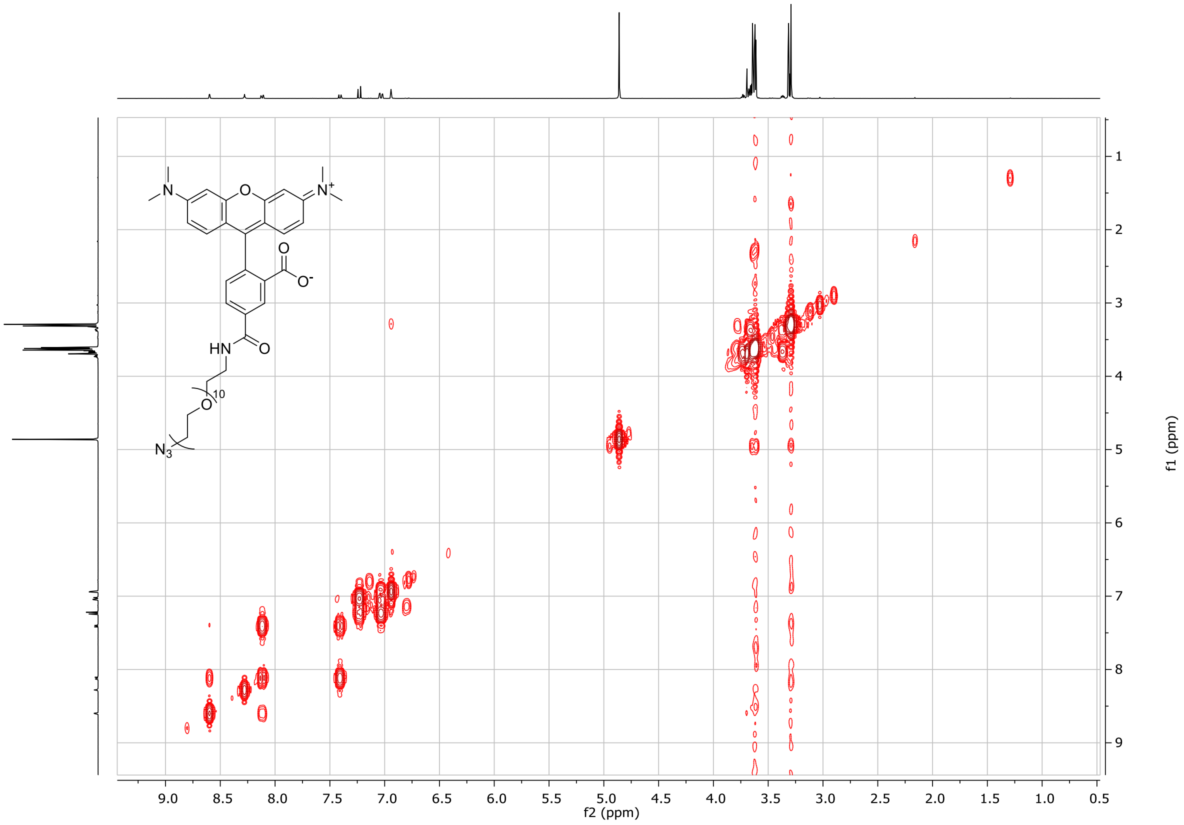

[1]H spectrum of **6-TMR-PEG$_{10}$-N$_3$**:

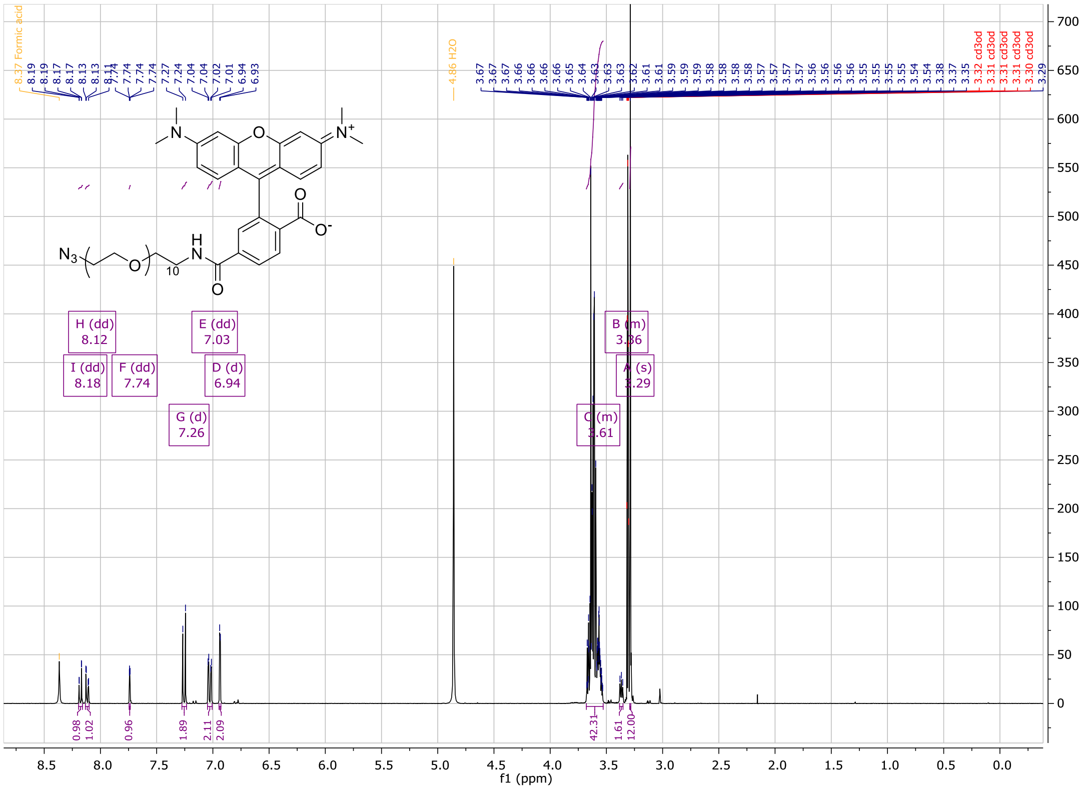

COSY spectrum of **5-TMR-PEG$_{10}$-N$_3$:**

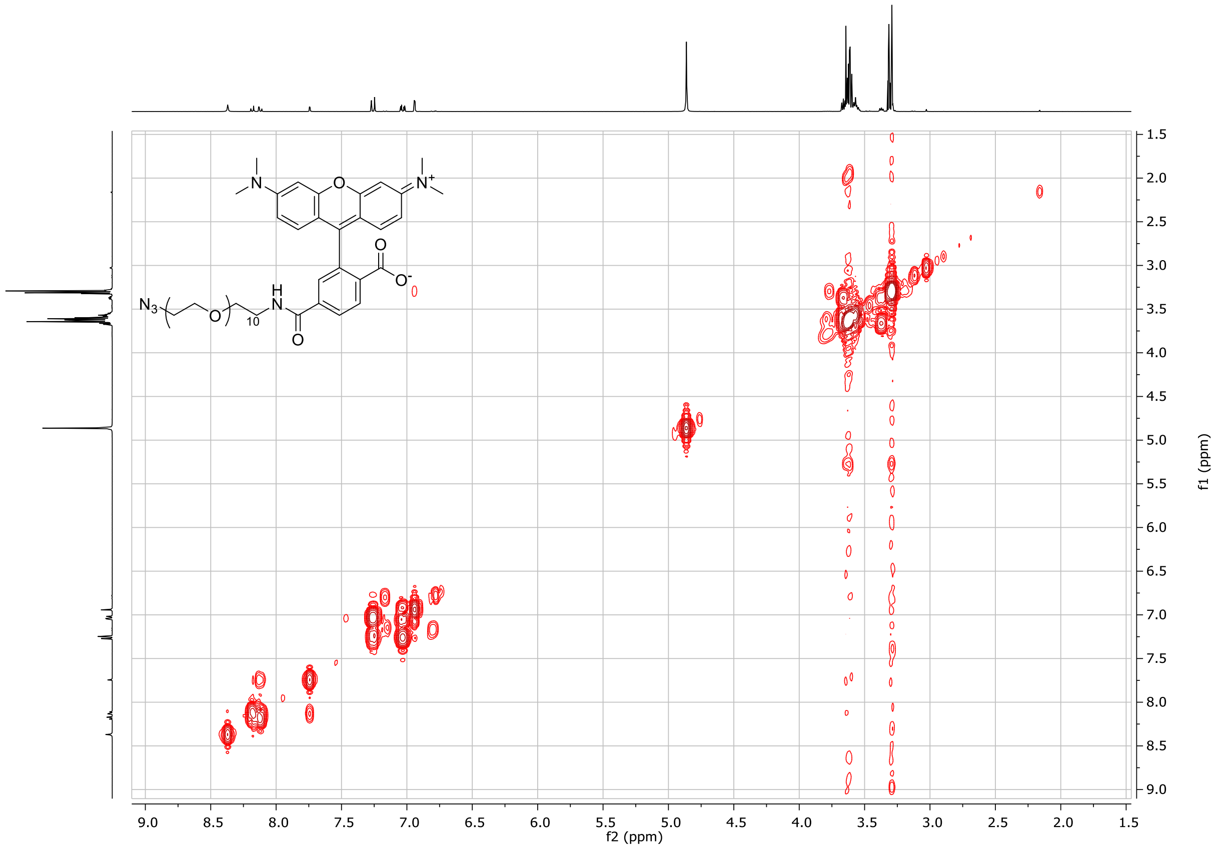

$^{1}$H spectrum of **Cy3B-PEG$_{10}$-N$_3$**:

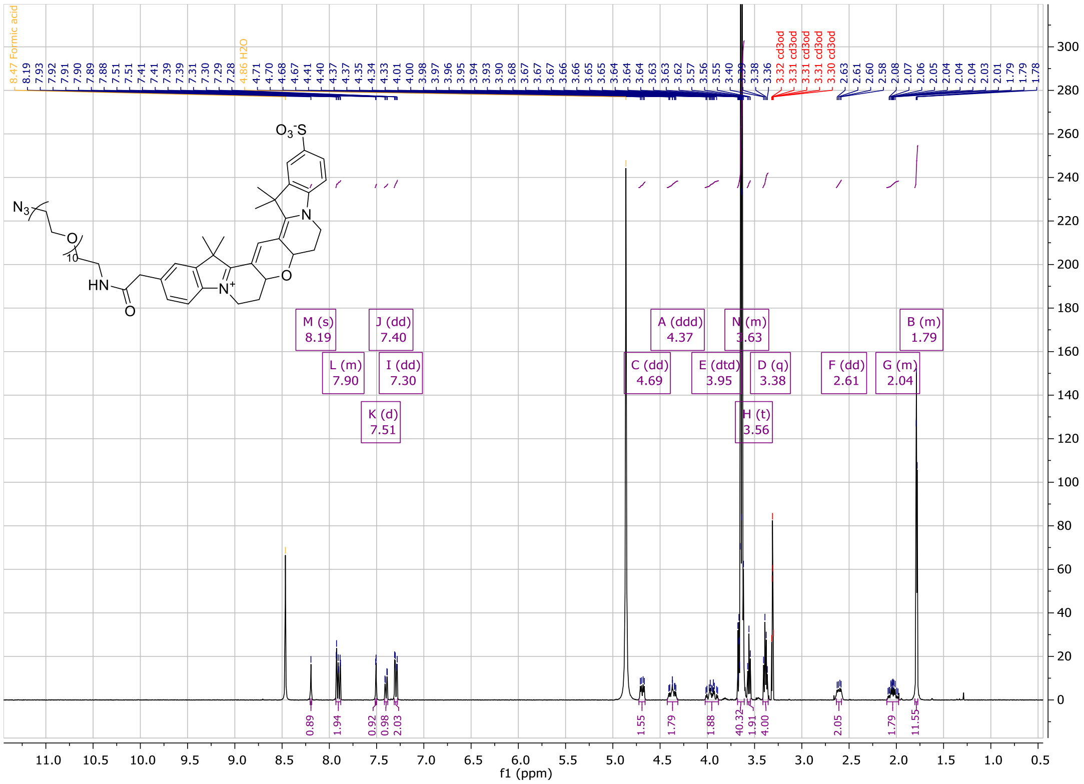

COSY spectrum of **Cy3B-PEG$_{10}$-N$_3$:**

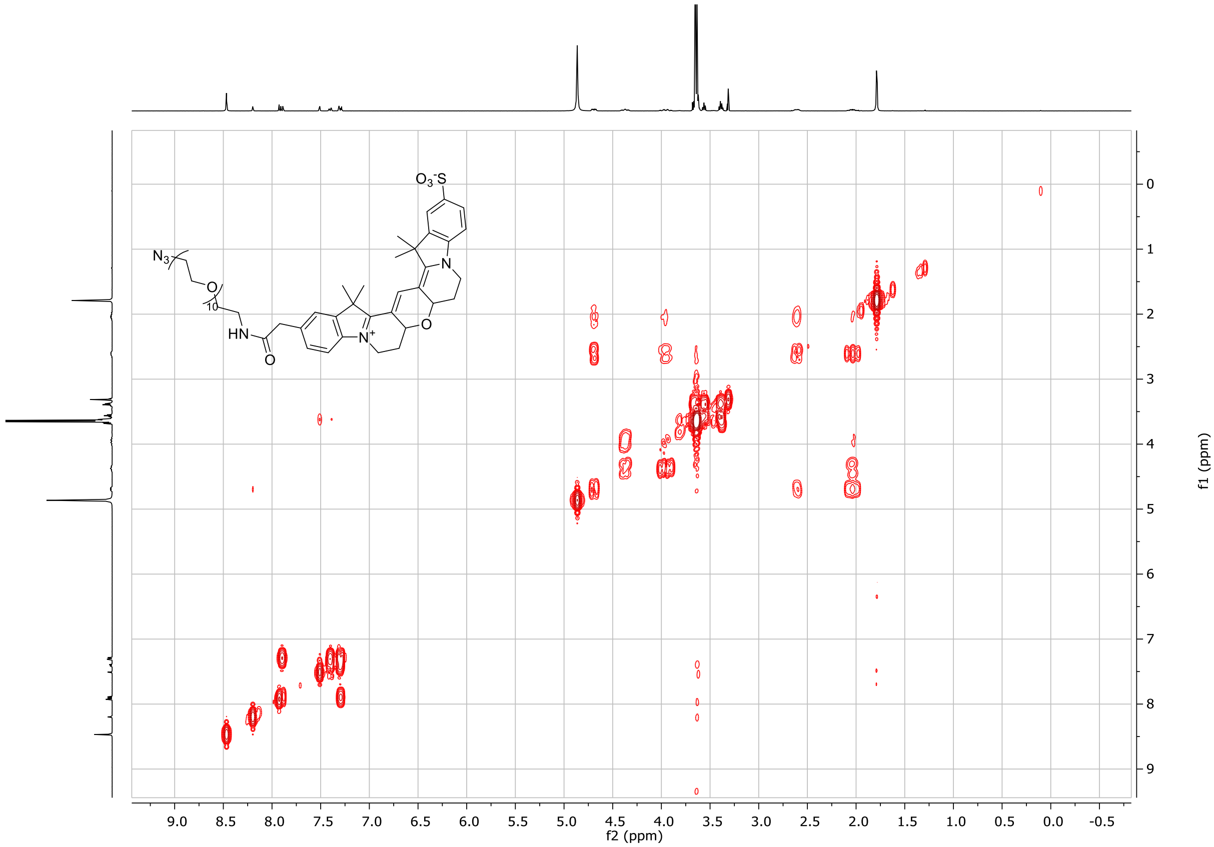
